## Supplementary Code 1 for "Inferring differential subcellular localisation in comparative spatial proteomics using BANDLE": beltran_analysis.rmd

````
---
title: "Beltran24_48_analysis"
author: "Oliver M. Crook"
date: "6 April 2020"
output: html_document
---

```{r, echo = FALSE, out = FALSE, }
require(pRoloc)
require(pRolocdata)
require(ggplot2)
source("bandleFUN.R")
```


```{r,}
data("beltran2016HCMV24")
data("beltran2016HCMV48")
data("beltran2016MOCK24")
data("beltran2016MOCK48")
```

```{r,}
idx <- intersect(rownames(beltran2016MOCK48), rownames(beltran2016HCMV48))

HCMV48_sub <- beltran2016HCMV48[idx, ]
MOCK48_sub <- beltran2016MOCK48[idx, ]


beltrandyn48 <- c(MOCK48_sub, HCMV48_sub) 
beltrandyn48 <- lapply(beltrandyn48, function(x) normalise(x, method = "center.median"))
```

```{r,}
idx <- intersect(rownames(beltran2016MOCK24), rownames(beltran2016HCMV24))

HCMV24_sub <- beltran2016HCMV24[idx, ]
MOCK24_sub <- beltran2016MOCK24[idx, ]


beltrandyn24 <- c(MOCK24_sub, HCMV24_sub) 
beltrandyn24 <- lapply(beltrandyn24, function(x) normalise(x, method = "center.median"))

K <- length(getMarkerClasses(HCMV24_sub))

```


```{r,}
# par(mfrow = c(3,4))
# pc_prior <- matrix(NA, ncol = 3, K)
# pc_prior[seq.int(1:K), ] <- matrix(rep(c(1, 60, 100), each = K), ncol = 3)
# hypres <- lapply(beltrandyn48, function(x) fitGPmaternPC(x, fcol = "markers", nu = 2, materncov = TRUE, hyppar = pc_prior)) 
# save(hypres, file = "beltran48hypers.rda")

```

```{r, echo = FALSE, out = FALSE, print = FALSE}
idx <- intersect(rownames(beltran2016MOCK24), rownames(beltran2016HCMV24))

HCMV24_sub <- beltran2016HCMV24[idx, ]
MOCK24_sub <- beltran2016MOCK24[idx, ]


beltrandyn24 <- c(MOCK24_sub, HCMV24_sub) 
beltrandyn24 <- lapply(beltrandyn24, function(x) normalise(x, method = "center.median"))

K <- length(getMarkerClasses(MOCK24_sub))

dir_prior = diag(rep(1, K)) + matrix(0.01, nrow = K, ncol = K)
prior_pred_dir(object = MOCK24_sub, dir_prior = dir_prior)

```

```{r,}
load("C:/Users/OllyC/Desktop/2019-DiffLoc/bandleres_beltran24.rda")


```

```{r,}
plot2D(beltran2016MOCK24, grid = FALSE, method = "t-SNE")
addLegend(beltran2016MOCK24)
plot2D(beltran2016HCMV24, grid = FALSE, method = "t-SNE")
addLegend(beltran2016HCMV24)

```
```{r,}
setStockcol(paste0(getStockcol(),90))
mycols <- c("blue", "darkgreen", "orange", "purple", "pink", "brown", "gray", "cyan")
mycols <- sapply(mycols, function(x) alpha(x, 0.2))
```


```{r,}
par(mfrow = c(3,4))

for (j in seq_along(getMarkerClasses(beltran2016HCMV24))) {
  for (i in seq_along(beltran2016HCMV24)){
    if (i ==  1) {
      M <- max(bandleres_beltran24[[i]]$weights[j,j,]) + 0.05
      plot(bandleres_beltran24[[i]]$weights[j,j,], col = mycols[[i]], type = "b", pch = 19, ylim = c(0, M), ylab = "mixing proportions")
    } else {
      points(bandleres_beltran24[[i]]$weights[j,j,], col = mycols[[i]], type = "b", pch = 19)
    }    }
}
par(mfrow = c(3,4))
for (j in seq_along(getMarkerClasses(beltran2016HCMV24))) {
  for (i in c(2,3,4,5,6)){
    if (i ==  2) {
      M <- max(bandleres_beltran24[[i]]$weights[j,j,]) + 0.05
      plot(bandleres_beltran24[[i]]$weights[j,j,], col = mycols[[i]], type = "b", pch = 19, ylim = c(0, M), ylab = "mixing proportions")
    } else {
      points(bandleres_beltran24[[i]]$weights[j,j,], col = mycols[[i]], type = "b", pch = 19)
    }  
  }
}


```
```{r,}
require(coda)
r_hat <- vector(mode = "list", length = length(getMarkerClasses(beltran2016HCMV24)))
for (j in seq_along(getMarkerClasses(beltran2016HCMV24))) {
  r_hat[[j]] <- gelman.diag(mcmc.list(mcmc(bandleres_beltran24[[2]]$weights[j,j,]),
                                      mcmc(bandleres_beltran24[[3]]$weights[j,j,])), transform = TRUE)
}
unlist(r_hat)
```
```{r,}
par(mfrow = c(2,3))
rk <- rank(cbind(bandleres_beltran24[[2]]$weights[j,j,],
                 bandleres_beltran24[[3]]$weights[j,j,],
                 bandleres_beltran24[[4]]$weights[j,j,],
                 bandleres_beltran24[[5]]$weights[j,j,],
                 bandleres_beltran24[[6]]$weights[j,j,]))

hist(rk[1:667], col = alpha("blue", 0.2), breaks = seq.int(from = 1, to = length(rk), length.out = 10), main = "rank plot", xlab = "rank")
hist(rk[668:1334], col = alpha("darkgreen", 0.2),breaks = seq.int(from = 1, to = length(rk), length.out = 10), main = "rank plot", xlab = "rank")
hist(rk[1335:2001], col = mycols[3],breaks = seq.int(from = 1, to = length(rk), length.out = 10), main = "rank plot", xlab = "rank")
hist(rk[2001:2668], col = mycols[4],breaks = seq.int(from = 1, to = length(rk), length.out = 10), main = "rank plot", xlab = "rank")
hist(rk[2668:3335], col = mycols[5],breaks = seq.int(from = 1, to = length(rk), length.out = 10), main = "rank plot", xlab = "rank")

rk <- rank(cbind(bandleres_beltran24[[2]]$weights[j,j,],
                 bandleres_beltran24[[3]]$weights[j,j,],
                 bandleres_beltran24[[6]]$weights[j,j,]))
```


```{r,}
par(mfrow = c(1,3))
hist(rk[1:667], col = alpha("blue", 0.2), breaks = seq.int(from = 1, to = length(rk), length.out = 10), main = "rank plot", xlab = "rank")
hist(rk[668:1334], col = alpha("darkgreen", 0.2),breaks = seq.int(from = 1, to = length(rk), length.out = 10), main = "rank plot", xlab = "rank")
hist(rk[1335:2001], col = mycols[3],breaks = seq.int(from = 1, to = length(rk), length.out = 10), main = "rank plot", xlab = "rank")


```

```{r,}
probdir_beltran24 <- probsameorganelle(object = MOCK24_sub, params = bandleres_beltran24[[3]])

```

```{r,}
meanallocationProb1 <- apply(bandleres_beltran24[[3]]$allocprob$cond1,c(1), colMeans)
rownames(meanallocationProb1) <- getMarkerClasses(MOCK24_sub)

maxalloc1 <- apply(meanallocationProb1, 2,which.max)


meanallocationProb2 <- apply(bandleres_beltran24[[3]]$allocprob$cond2,c(1), colMeans)
rownames(meanallocationProb2) <- getMarkerClasses(MOCK24_sub)

maxalloc2 <- apply(meanallocationProb2, 2,which.max)


colnames(meanallocationProb1) <- names(probdir_beltran24)
colnames(meanallocationProb2) <- names(probdir_beltran24)

names(maxalloc1) <- names(maxalloc2) <- rownames(unknownMSnSet(MOCK24_sub))
```


```{r,}
## BANDLE dirichlet
par(mfrow = c(1,1))
y <- 1 - probdir_beltran24[order(probdir_beltran24)]/ncol(bandleres_beltran24[[3]]$allocOut$cond1)

write.csv(y, file = "beltran24transloc.csv")


y_filter <- y[maxalloc1 %in% c(1,2,7) | maxalloc2 %in% c(1,2,7)]

write.csv(y_filter, file = "beltran24translocfitler.csv")


par(mfrow = c(1,1))
plot(y, pch = 19, ylim = c(0,1), ylab = "Differential Localisation Probability", xlab = "Rank", 
     col = getStockcol()[2], main = "Differential localisation rank", cex.lab = 1.5, cex.main = 2, cex.axis = 1.5)
```


```{r,}
require(viridis)
meanallocationProb <- apply(bandleres_beltran24[[3]]$allocprob$cond1,c(1), colMeans)

rownames(meanallocationProb) <- getMarkerClasses(MOCK24_sub)

pheatmap::pheatmap(meanallocationProb1, color = viridis(100, alpha = 0.8, direction = 1), cluster_rows = FALSE, cluster_cols = TRUE, treeheight_col = 0)
pheatmap::pheatmap(meanallocationProb2, color = viridis(100, alpha = 0.8, direction = 1), cluster_rows = FALSE, cluster_cols = TRUE, treeheight_col = 0)
```


```{r,}
require(viridis)
crossalloc <- table(maxalloc1,maxalloc2)
rownames(crossalloc) <- colnames(crossalloc) <- getMarkerClasses(MOCK24_sub)

crossalloc_sub <- crossalloc[-c(1,2,7),-c(1,2,7)]

pheatmap::pheatmap(crossalloc + diag(table(fData(HCMV24_sub)$markers)[1:9]), color = viridis(100, alpha = 0.8, direction = 1), cluster_rows = FALSE, cluster_cols = FALSE, treeheight_col = 0, display_numbers = TRUE, fontsize_number = 12, number_color = "white", number_format = "%g")


pheatmap::pheatmap(crossalloc_sub, color = viridis(100, alpha = 0.8, direction = 1), cluster_rows = FALSE, cluster_cols = FALSE, treeheight_col = 0, display_numbers = TRUE, fontsize_number = 12, number_color = "white", number_format = "%g")
```

```{r,}
crossalloc_conf <- table(maxalloc1[names(y[y > 0.999])], maxalloc2[names(y[y > 0.999])])
#crossalloc_conf <- crossalloc_conf[-c(1,2,7),-c(1,2,7)]
#colnames(crossalloc_conf) <- rownames(crossalloc_sub)
#rownames(crossalloc_conf) <- getMarkerClasses(MOCK24_sub)[c(3,4,5,6,9)]

colnames(crossalloc_conf) <- getMarkerClasses(MOCK24_sub)
rownames(crossalloc_conf) <- getMarkerClasses(MOCK24_sub)[-8]


pheatmap::pheatmap(crossalloc_conf, color = viridis(100, alpha = 0.8, direction = 1), cluster_rows = FALSE, cluster_cols = FALSE, treeheight_col = 0, display_numbers = TRUE, fontsize_number = 12, number_color = "white", number_format = "%g")
```

```{r,}
df_alloc <- diag(diag(crossalloc) + table(fData(HCMV24_sub)$markers)[1:9])
df_alloc[1:7,1:9] <- df_alloc[1:7,1:9] + crossalloc_conf[1:7,1:9]
df_alloc[9,] <- df_alloc[9,] + crossalloc_conf[8,]

rownames(df_alloc) <- colnames(df_alloc) <- getMarkerClasses(MOCK24_sub)

pheatmap::pheatmap(df_alloc, color = viridis(100, alpha = 0.8, direction = 1), cluster_rows = FALSE, cluster_cols = FALSE, treeheight_col = 0, display_numbers = TRUE, fontsize_number = 12, number_color = "white", number_format = "%g")
```

```{r,}
outcyto <- names(maxalloc1[names(y)])[maxalloc1[names(y)] == 1 & maxalloc2[names(y)] != 1]
outcyto[outcyto %in% names(y[y > 0.999])]

write.csv(outcyto[outcyto %in% names(y[y > 0.999])], file = "beltranoutcyto.csv")
write.csv(names(maxalloc1[names(y)][maxalloc1[names(y)] == 1]), file = "beltranbackcyto.csv")


```


```{r,}
outdenscyto <- names(maxalloc1[names(y)])[maxalloc1[names(y)] == 2 & maxalloc2[names(y)] == 3]
outdenscyto[outdenscyto %in% names(y[y > 0.999])]

write.csv(outdenscyto[outdenscyto %in% names(y[y > 0.999])], file = "beltranoutdenscytoER.csv")

outdenscyto <- names(maxalloc1[names(y)])[maxalloc1[names(y)] == 2 & maxalloc2[names(y)] == 4]
outdenscyto[outdenscyto %in% names(y[y > 0.999])]

write.csv(outdenscyto[outdenscyto %in% names(y[y > 0.999])], file = "beltranoutdenscytoGolgi.csv")

outdenscyto <- names(maxalloc1[names(y)])[maxalloc1[names(y)] == 2 & maxalloc2[names(y)] == 9]
outdenscyto[outdenscyto %in% names(y[y > 0.999])]

write.csv(outdenscyto[outdenscyto %in% names(y[y > 0.999])], file = "beltranoutdenscytoPM.csv")

outdenscyto <- names(maxalloc1[names(y)])[maxalloc1[names(y)] == 2 & maxalloc2[names(y)] %in% c(3,4,9)]
outdenscyto[outdenscyto %in% names(y[y > 0.999])]


write.csv(outdenscyto[outdenscyto %in% names(y[y > 0.999])], file = "beltranoutdenssec.csv")


outdenscyto <- names(maxalloc1[names(y)])[maxalloc1[names(y)] == 2 & maxalloc2[names(y)] != 1]
outdenscyto[outdenscyto %in% names(y[y > 0.999])]


write.csv(outdenscyto[outdenscyto %in% names(y[y > 0.999])], file = "beltranoutdenscyto.csv")


```

```{r,}
ERtoCyto <- names(maxalloc1[names(y)])[maxalloc1[names(y)] == 3 & maxalloc2[names(y)] == 2]


write.csv(ERtoCyto, file = "beltranertocyto.csv")


LystoER <- names(maxalloc1[names(y)])[maxalloc1[names(y)] == 5 & maxalloc2[names(y)] == 3]


write.csv(LystoER, file = "beltranlystoER.csv")

CyttoPM <- names(maxalloc1[names(y)])[maxalloc1[names(y)] == 2 & maxalloc2[names(y)] == 9]


write.csv(CyttoPM, file = "beltranCyttoPM.csv")

PMout <- names(maxalloc1[names(y)])[maxalloc1[names(y)] == 9 & maxalloc2[names(y)] != 9]

write.csv(PMout, file = "beltranPMout.csv")
```


```{r,}
cors <- vector(mode = "list", length = nrow(MOCK24_sub))
for (j in seq.int(nrow(MOCK24_sub))) {
  cors[[j]] <- cor.test(exprs(MOCK24_sub[j,]), exprs(HCMV24_sub[j,]))$estimate
}
```


```{r,}
require(openxlsx)
HCMVdegredationdata <- read.xlsx("HCMVdegredationdata.xlsx", sheet = 3)
HCMVviralproteomics <- read.xlsx("HCMVviralproteomicsdata.xlsx", sheet = 2)
HCMVviralproteomics2 <- read.xlsx("HCMVtimecourse2.xlsx", sheet = 1, startRow = 2)
HCMVinteractome1 <- read.xlsx("HCMVinteractome.xlsx", sheet = 1)
HCMVinteractome2 <- read.xlsx("HCMVinteractome.xlsx", sheet = 2)
HCMVacetylome <- read.xlsx("HCMVacetylation.xlsx", sheet = 1, startRow = 2)
```


```{r,}
degradationvalues <- HCMVdegredationdata[,c(3,27:31,36:39,44:47)]
degradationvalues_sub <- degradationvalues[rowSums(is.na.data.frame(degradationvalues)) == 0,]
degradationvalues_sub <- data.frame(apply(degradationvalues_sub, 2, function(x) as.numeric(as.character(x))))[,-c(1,2)]
rownames(degradationvalues_sub) <- HCMVdegredationdata[rowSums(is.na.data.frame(degradationvalues)) == 0,2]

```


```{r,}
acetylationdata <- HCMVacetylome[,c(1,4,13:16)]
acetylationdata_sub <- acetylationdata[rowSums(is.na.data.frame(acetylationdata)) == 0,]
pheatmap::pheatmap(acetylationdata_sub[1:1000, 3:6], cluster_cols = FALSE, scale = "row", show_rownames = FALSE)
rownames(acetylationdata_sub) <- acetylationdata_sub[,1]
```

```{r,}
interactomedata1 <- HCMVinteractome1
hist(interactomedata1[,2])
```
```{r,}
interactomedata2 <- HCMVinteractome2

```

```{r,}
timecoursesub <- HCMVviralproteomics2[rowSums(is.na.data.frame(HCMVviralproteomics2)) == 0,c(1,9:12)]
rownames(timecoursesub) <- HCMVviralproteomics2[rowSums(is.na.data.frame(HCMVviralproteomics2)) == 0,1]
```

```{r,}
acetylationdata <- HCMVacetylome[,c(1,4,13:16)]
acetylationdata_sub <- acetylationdata[rowSums(is.na.data.frame(acetylationdata)) == 0,]
acetylationdata_sub <- aggregate(acetylationdata_sub, by = list(name = acetylationdata_sub$`Primary.Uniprot.Protein.Accession.#`), data =acetylationdata_sub , FUN = mean)
rownames(acetylationdata_sub) <- acetylationdata_sub[,1]
acetylation_overlap <- acetylationdata_sub[rownames(acetylationdata_sub) %in% c(names(y), rownames(markerMSnSet(HCMV24_sub))),]
rownames(acetylation_overlap) <- acetylation_overlap[,1]
boxplot(acetylationdata_sub[names(y[y > 0.999]),4], acetylationdata_sub[names(y[y < 0.999]),4], col = mycols, main = "Acetylation changes 24 hpi", names = c("Differentialy localised", "stable"))
```

```{r,}
combineddata <- cbind(timecoursesub[names(y),2:5], acetylation_overlap[names(y),3:6], degradationvalues_sub[names(y),])
combineddata <- combineddata[rowSums(is.na.data.frame(combineddata)) == 0,]
```

```{r,}
.pca <- prcomp(combineddata, scale = TRUE, center = TRUE)

plotcols <- y[rownames(combineddata)]
plotcols <- 1 * (plotcols > 0.5)
plot(.pca$x[,1], .pca$x[,2], col = mycols[plotcols])
```


```{r,}
timebigchangediff <- apply(timecoursesub[names(y[y > 0.999])[1:200],2:4],1, function(x) abs(max(x) - min(x)))
timebigchangestable <- apply(timecoursesub[names(y[y < 0.999])[1:200],2:4],1, function(x) abs(max(x) - min(x)))
```

```{r,}
degradespatial <- array(NA, c(9,9,12))
dimnames(degradespatial)[[1]] <- dimnames(degradespatial)[[2]] <- getMarkerClasses(MOCK24_sub)
for (j in seq.int(K)){
  for(k in seq.int(K)){
    degradespatial[j, k, ] <- colMeans(degradationvalues_sub[names(maxalloc1[maxalloc1 == j & maxalloc2 == k]),], na.rm = TRUE)
  }
}

for (j in seq.int(12)) {
  degradespatial[, , j][is.na(degradespatial[, , j])] <- 0
}

breaksList = seq(0, 1.2, by = 0.02)

pheatmap::pheatmap(degradespatial[,,1], color = viridis(61, alpha = 0.8, direction = 1), cluster_rows = FALSE, cluster_cols = FALSE, treeheight_col = 0, display_numbers = TRUE, fontsize_number = 12, number_color = "white", number_format = "%.2f", breaks = breaksList, main = "mean MG132 recovery ratio 12h")
```

```{r,}
degradespatialpvalue <- matrix(NA, 9, 9)
colnames(degradespatialpvalue) <- rownames(degradespatialpvalue) <- getMarkerClasses(object = MOCK24_sub)

for (j in seq.int(K)){
  for(k in seq.int(K)){
    if (sum(maxalloc1 == j & maxalloc2 == k) < 3 ){
      degradespatialpvalue[j,k] <- 1
    } else {
    degradespatialpvalue[j, k] <- t.test(degradationvalues_sub[names(maxalloc1[maxalloc1 == j & maxalloc2 == k]),1], degradationvalues_sub[,1])$p.value
    }
  }
}

pheatmap::pheatmap(-log10(degradespatialpvalue) + 0, color = inferno(50, alpha = 0.8, direction = 1), cluster_rows = FALSE, cluster_cols = FALSE, treeheight_col = 0, display_numbers = TRUE, fontsize_number = 12, number_color = "white", number_format = "%.2f", breaks = seq.int(40), main = "pvalue MG132 recovery ratio 12h")


```


```{r,}
breaksList = seq(0, 1.2, by = 0.02)

pheatmap::pheatmap(degradespatial[,,3], color = viridis(61, alpha = 0.8, direction = 1), cluster_rows = FALSE, cluster_cols = FALSE, treeheight_col = 0, display_numbers = TRUE, fontsize_number = 12, number_color = "white", number_format = "%.2f", breaks = breaksList, main = "mean Leupeptin recovery ratio 12h")
```

```{r,}
degradespatial2pvalue <- matrix(NA, 9, 9)
colnames(degradespatial2pvalue) <- rownames(degradespatial2pvalue) <- getMarkerClasses(object = MOCK24_sub)

for (j in seq.int(K)){
  for(k in seq.int(K)){
    if (sum(maxalloc1 == j & maxalloc2 == k) < 2 ){
      degradespatial2pvalue[j, k] <- 1
    } else {
    degradespatial2pvalue[j, k] <- t.test(degradationvalues_sub[names(maxalloc1[maxalloc1 == j & maxalloc2 == k]),3], degradationvalues_sub[,3])$p.value
    }
  }
}

pheatmap::pheatmap(-log10(degradespatial2pvalue) + 0, color = inferno(12, alpha = 0.8, direction = 1), cluster_rows = FALSE, cluster_cols = FALSE, treeheight_col = 0, display_numbers = TRUE, fontsize_number = 12, number_color = "white", number_format = "%.2f", breaks = seq.int(10), main = "pvalue leupeptin recovery ratio 12h")


```


```{r,}
pheatmap::pheatmap(degradespatial[,,5], color = viridis(61, alpha = 0.8, direction = 1), cluster_rows = FALSE, cluster_cols = FALSE, treeheight_col = 0, display_numbers = TRUE, fontsize_number = 12, number_color = "white", number_format = "%.2f", breaks = breaksList, main = "mean MG132 recovery ratio 18h")

```

```{r,}
degradespatialmg18hpvalue <- matrix(NA, 9, 9)
colnames(degradespatialmg18hpvalue) <- rownames(degradespatialmg18hpvalue) <- getMarkerClasses(object = MOCK24_sub)

for (j in seq.int(K)){
  for(k in seq.int(K)){
    if (sum(maxalloc1 == j & maxalloc2 == k) < 3 ){
      degradespatialmg18hpvalue[j,k] <- 1
    } else {
    degradespatialmg18hpvalue[j, k] <- t.test(degradationvalues_sub[names(maxalloc1[maxalloc1 == j & maxalloc2 == k]),5], degradationvalues_sub[,5])$p.value
    }
  }
}

pheatmap::pheatmap(-log10(degradespatialmg18hpvalue) + 0, color = inferno(12, alpha = 0.8, direction = 1), cluster_rows = FALSE, cluster_cols = FALSE, treeheight_col = 0, display_numbers = TRUE, fontsize_number = 12, number_color = "white", number_format = "%.2f", breaks = seq.int(10), main = "pvalue MG132 recovery ratio 18h")


```


```{r,}
pheatmap::pheatmap(degradespatial[,,9], color = viridis(61, alpha = 0.8, direction = 1), cluster_rows = FALSE, cluster_cols = FALSE, treeheight_col = 0, display_numbers = TRUE, fontsize_number = 12, number_color = "white", number_format = "%.2f", breaks = breaksList, main = "mean MG132 recovery ratio 24h")

```


```{r,}
degradespatialmg24hpvalue <- matrix(NA, 9, 9)
colnames(degradespatialmg24hpvalue) <- rownames(degradespatialmg24hpvalue) <- getMarkerClasses(object = MOCK24_sub)

for (j in seq.int(K)){
  for(k in seq.int(K)){
    if (sum(maxalloc1 == j & maxalloc2 == k) < 3 ){
      degradespatialmg24hpvalue[j,k] <- 1
    } else {
    degradespatialmg24hpvalue[j, k] <- t.test(degradationvalues_sub[names(maxalloc1[maxalloc1 == j & maxalloc2 == k]),9], degradationvalues_sub[,9])$p.value
    }
  }
}

pheatmap::pheatmap(-log10(degradespatialmg24hpvalue) + 0, color = inferno(12, alpha = 0.8, direction = 1), cluster_rows = FALSE, cluster_cols = FALSE, treeheight_col = 0, display_numbers = TRUE, fontsize_number = 12, number_color = "white", number_format = "%.2f", breaks = seq.int(10), main = "-log10(p-value) MG132 rescue ratio 24h", fontsize = 14)

```


```{r,}
breaksList = seq(0, 1.2, by = 0.02)

pheatmap::pheatmap(degradespatial[,,7], color = viridis(61, alpha = 0.8, direction = 1), cluster_rows = FALSE, cluster_cols = FALSE, treeheight_col = 0, display_numbers = TRUE, fontsize_number = 12, number_color = "white", number_format = "%.2f", breaks = breaksList, main = "mean leupeptin recovery ratio 18h")
```


```{r,}
degradespatialleu18hpvalue <- matrix(NA, 9, 9)
colnames(degradespatialleu18hpvalue) <- rownames(degradespatialleu18hpvalue) <- getMarkerClasses(object = MOCK24_sub)

for (j in seq.int(K)){
  for(k in seq.int(K)){
    if (sum(maxalloc1 == j & maxalloc2 == k) < 3 ){
      degradespatialleu18hpvalue[j,k] <- 1
    } else {
    degradespatialleu18hpvalue[j, k] <- t.test(degradationvalues_sub[names(maxalloc1[maxalloc1 == j & maxalloc2 == k]),7], degradationvalues_sub[,7])$p.value
    }
  }
}

pheatmap::pheatmap(-log10(degradespatialleu18hpvalue) + 0, color = inferno(12, alpha = 0.8, direction = 1), cluster_rows = FALSE, cluster_cols = FALSE, treeheight_col = 0, display_numbers = TRUE, fontsize_number = 12, number_color = "white", number_format = "%.2f", breaks = seq.int(10), main = "pvalue leupeptin recovery ratio 18h")

```


```{r,}
breaksList = seq(0, 1.2, by = 0.02)

pheatmap::pheatmap(degradespatial[,,11], color = viridis(61, alpha = 0.8, direction = 1), cluster_rows = FALSE, cluster_cols = FALSE, treeheight_col = 0, display_numbers = TRUE, fontsize_number = 12, number_color = "white", number_format = "%.2f", breaks = breaksList, main = "mean leupeptin recovery ratio 24h")
```


```{r,}
degradespatialleu24hpvalue <- matrix(NA, 9, 9)
colnames(degradespatialleu24hpvalue) <- rownames(degradespatialleu24hpvalue) <- getMarkerClasses(object = MOCK24_sub)

for (j in seq.int(K)){
  for(k in seq.int(K)){
    if (sum(maxalloc1 == j & maxalloc2 == k) < 3 ){
      degradespatialleu24hpvalue[j,k] <- 1
    } else {
    degradespatialleu24hpvalue[j, k] <- t.test(degradationvalues_sub[names(maxalloc1[maxalloc1 == j & maxalloc2 == k]),11], degradationvalues_sub[,11])$p.value
    }
  }
}

pheatmap::pheatmap(-log10(degradespatialleu24hpvalue) + 0, color = inferno(12, alpha = 0.8, direction = 1), cluster_rows = FALSE, cluster_cols = FALSE, treeheight_col = 0, display_numbers = TRUE, fontsize_number = 12, number_color = "white", number_format = "%.2f", breaks = seq.int(10), main = "-log(pvalue) leupeptin rescue ratio 24h", fontsize = 14)

```

```{r,}
par(mfrow = c(1,2))
boxplot(degradationvalues_sub[y > 0.999,11], degradationvalues_sub[y < 0.999,11] , names = c("Differentially localised", "Unperturbed"), main = "Degradation Distributions Leupeptin 24 hpi", ylab = "Rescue ratio", col = mycols)

boxplot(degradationvalues_sub[y > 0.999,9], degradationvalues_sub[y < 0.999,9] , names = c("Differentially localised", "Unperturbed"), main = "Degradation Distributions MG132 24 hpi", ylab = "Rescue ratio", col = mycols)

```

```{r,}
boxplot(degradationvalues_sub[names(maxalloc1)[maxalloc1 == 1 & maxalloc2 == 2],11], degradationvalues_sub[,11] , names = c("Cytosol to Dense Cytosol", "Global"), main = "Degradation Distributions Leupteptin 24 hpi", ylab = "Rescue ratio", col = mycols)

```


## Temporal viromics data
```{r,}
temporalspatial <- array(NA, c(9, 9, 4))
dimnames(temporalspatial)[[1]] <- dimnames(temporalspatial)[[2]] <- getMarkerClasses(MOCK24_sub)
for (j in seq.int(K)){
  for(k in seq.int(K)){
    temporalspatial[j, k, ] <- colMeans(timecoursesub[names(maxalloc1[maxalloc1 == j & maxalloc2 == k]),2:5], na.rm = TRUE)
  }
}

for (j in seq.int(4)) {
  temporalspatial[, , j][is.na(temporalspatial[, , j])] <- 0
}

breaksList = seq(0, 1.2, by = 0.02)

pheatmap::pheatmap(temporalspatial[,,1], color = viridis(61, alpha = 0.8, direction = 1), cluster_rows = FALSE, cluster_cols = FALSE, treeheight_col = 0, display_numbers = TRUE, fontsize_number = 12, number_color = "white", number_format = "%.2f", main = "mean log2 fold changes in abudance 24 hpi", fontsize = 14)
```
```{r,}
temporalspatial24hpvalue <- matrix(NA, 9, 9)
colnames(temporalspatial24hpvalue) <- rownames(temporalspatial24hpvalue) <- getMarkerClasses(object = MOCK24_sub)

for (j in seq.int(K)){
  for(k in seq.int(K)){
    if (sum(!is.na(timecoursesub[names(maxalloc1[maxalloc1 == j & maxalloc2 == k]),2])) < 3 ){
      temporalspatial24hpvalue[j,k] <- 1
    } else {
    temporalspatial24hpvalue[j, k] <- t.test(timecoursesub[names(maxalloc1[maxalloc1 == j & maxalloc2 == k]),2], timecoursesub[,2])$p.value
    }
  }
}

pheatmap::pheatmap(-log10(temporalspatial24hpvalue) + 0, color = inferno(40, alpha = 0.8, direction = 1), cluster_rows = FALSE, cluster_cols = FALSE, treeheight_col = 0, display_numbers = TRUE, fontsize_number = 12, number_color = "white", number_format = "%.2f", breaks = seq(from = 0, to = 5, length.out = 30), main = "-log10 p-values for fold changes in abundance 24 hpi")
```
```{r,}
boxplot(timecoursesub[names(y[y > 0.999]),2], timecoursesub[names(y[y < 0.999]),2] , names = c("Differentially localised", "Unperturbed"), main = "log2 abundance distributions 24 hpi", ylab = "lo2 change in abundance", col = mycols)


boxplot(timecoursesub[names(maxalloc1[maxalloc1 == 3 & maxalloc2 == 2]),2], timecoursesub[names(maxalloc1),2] , names = c("ER to dense Cytosol", "global"), main = "log2 abundance distributions 24 hpi", ylab = "lo2 change in abundance", col = mycols)


```


```{r,}

matplot(x = c(24,48,72,96), t(timecoursesub[names(maxalloc1[maxalloc1 == 1 & maxalloc2 == 4]),2:5]),
type = "b", lwd = 6, col = mycols[1], pch = 19, lty = 1, ylab = "log2 fold change", xlab = "hpi")

matplot(x = c(24,48,72,96), t(timecoursesub[names(maxalloc1[maxalloc1 == 3 & maxalloc2 == 2]),2:5]),
type = "b", lwd = 6, col = mycols[1], pch = 19, lty = 1, ylab = "log2 fold change", xlab = "hpi")

matplot(x = c(24,48,72,96), t(timecoursesub[names(maxalloc1[maxalloc1 == 3 & maxalloc2 == 4]),2:5]),
type = "b", lwd = 6, col = mycols[1], pch = 19, lty = 1, ylab = "log2 fold change", xlab = "hpi")

matplot(x = c(24,48,72,96), t(timecoursesub[names(maxalloc1[maxalloc1 == 3 & maxalloc2 == 3]),2:5]),
type = "b", lwd = 6, col = mycols[1], pch = 19, lty = 1, ylab = "log2 fold change", xlab = "hpi")


matplot(x = c(24,48,72,96), t(timecoursesub[names(maxalloc1[maxalloc1 == 4 & maxalloc2 == 5]),2:5]),
type = "b", lwd = 6, col = mycols[1], pch = 19, lty = 1, ylab = "log2 fold change", xlab = "hpi", ylim = c(-0.4,0.4), main = "Q92520 temporal abundance")

```


## Temporal viromics acetylation data
```{r,}
acetylationspatial <- array(NA, c(9, 9, 4))
dimnames(acetylationspatial)[[1]] <- dimnames(acetylationspatial)[[2]] <- getMarkerClasses(MOCK24_sub)
for (j in seq.int(K)){
  for(k in seq.int(K)){
    acetylationspatial[j, k, ] <- colMeans(acetylationdata_sub[acetylationdata_sub[,1] %in% names(maxalloc1[maxalloc1 == j & maxalloc2 == k]),4:7], na.rm = TRUE)
  }
}

for (j in seq.int(4)) {
  acetylationspatial[, , j][is.na(acetylationspatial[, , j])] <- 0
}

breaksList = seq(0, 1.2, by = 0.02)

pheatmap::pheatmap(acetylationspatial[,,1], color = viridis(61, alpha = 0.8, direction = 1), cluster_rows = FALSE, cluster_cols = FALSE, treeheight_col = 0, display_numbers = TRUE, fontsize_number = 12, number_color = "white", number_format = "%.2f", main = "mean log2 fold changes in acetylation 24 hpi", fontsize = 14)
```

```{r,}
acetylation24hpvalue <- matrix(NA, 9, 9)
colnames(acetylation24hpvalue) <- rownames(acetylation24hpvalue) <- getMarkerClasses(object = MOCK24_sub)

for (j in seq.int(K)){
  for(k in seq.int(K)){
    if (sum(!(is.na((acetylationdata_sub[names(maxalloc1[maxalloc1 == j & maxalloc2 == k]) ,4])))) < 3 ){
      acetylation24hpvalue[j,k] <- 1
    } else {
      acetylation24hpvalue[j, k] <- t.test(acetylationdata_sub[names(maxalloc1[maxalloc1 == j & maxalloc2 == k]), 4], acetylationdata_sub[, 4])$p.value
    }
  }
}

pheatmap::pheatmap(-log10(acetylation24hpvalue) + 0, color = inferno(40, alpha = 0.8, direction = 1), cluster_rows = FALSE, cluster_cols = FALSE, treeheight_col = 0, display_numbers = TRUE, fontsize_number = 12, number_color = "white", number_format = "%.2f", breaks = seq(from = 0, to = 2, length.out = 30), main = "-log10(p-value) acetylation changes 24 hpi")
```


```{r,}
acnames <- acetylation_overlap[acetylation_overlap[,1] %in% names(y[y > 0.999]), 1]

table(maxalloc1[acnames], maxalloc2[acnames])

pheatmap::pheatmap(table(maxalloc1[acnames], maxalloc2[acnames]), color = viridis(61, alpha = 0.8, direction = 1), cluster_rows = FALSE, cluster_cols = FALSE, treeheight_col = 0, display_numbers = TRUE, fontsize_number = 12, number_color = "white", number_format = "%.2f")

```


```{r,}
matplot(x = c(24, 48, 72, 96), t(acetylationdata_sub[names(maxalloc1[maxalloc1 == 1 & maxalloc2 == 2]), 4:7]), type = "b", 
        lwd = 6, col = mycols[1], pch = 19, lty = 1, ylab = "log2 fold change")

matplot(x = c(24, 48, 72, 96), t(acetylationdata_sub[names(maxalloc1[maxalloc1 == 2 & maxalloc2 == 1]), 4:7]), type = "b", 
        lwd = 6, col = mycols[1], pch = 19, lty = 1, ylab = "log2 fold change", main = "Temporal acetylation profiles for Dense Cytosol to Cytosol", xlab = "hours post infection")

matplot(x = c(24, 48, 72, 96), t(acetylationdata_sub[names(maxalloc1[maxalloc1 == 1 & maxalloc2 == 2]), 4:7]), type = "b", 
        lwd = 6, col = mycols[1], pch = 19, lty = 1, ylab = "log2 fold change")

matplot(x = c(24, 48, 72, 96), t(acetylationdata_sub[names(maxalloc1[maxalloc1 == 1 & maxalloc2 == 2]), 4:7]), type = "b", 
        lwd = 6, col = mycols[1], pch = 19, lty = 1, ylab = "log2 fold change")


```

## HCMV interactomics

```{r,}
nmitact <- vector(mode = "list", length(unique(interactomedata2$Bait)))
zscoreitact <- vector(mode = "list", length(unique(interactomedata2$Bait)))
allocitact <- vector(mode = "list", length(unique(interactomedata2$Bait)))
allocdiff <- vector(mode = "list", length(unique(interactomedata2$Bait)))
allocconfintact <- vector(mode = "list", length(unique(interactomedata2$Bait)))
allocconfdiff <- vector(mode = "list", length(unique(interactomedata2$Bait)))


for (j in seq_along(nmitact)) {
  nmitact[[j]] <- interactomedata2[interactomedata2$Bait == unique(interactomedata2$Bait)[j], 6]
  zscoreitact[[j]] <- interactomedata2[interactomedata2$Bait == unique(interactomedata2$Bait)[j], 9]
  allocitact[[j]] <- table(maxalloc1[nmitact[[j]]], maxalloc2[nmitact[[j]]])
  allocconfintact[[j]] <- table(maxalloc1[nmitact[[j]][zscoreitact[[j]] > 10]], maxalloc2[nmitact[[j]][zscoreitact[[j]] > 10]])
  allocdiff[[j]] <- c(sum(maxalloc1[nmitact[[j]]] == maxalloc2[nmitact[[j]]], na.rm = TRUE), sum(maxalloc1[nmitact[[j]]] != maxalloc2[nmitact[[j]]], na.rm = TRUE))
  allocconfdiff[[j]] <- c(sum(maxalloc1[nmitact[[j]][zscoreitact[[j]] > 10]] == maxalloc2[nmitact[[j]][zscoreitact[[j]] > 10]], na.rm = TRUE),
                          sum(maxalloc1[nmitact[[j]][zscoreitact[[j]] > 10]] != maxalloc2[nmitact[[j]][zscoreitact[[j]] > 10]], na.rm = TRUE))
}

difflocintact <- matrix(unlist(allocdiff), ncol = 2, byrow = TRUE)
difflocinactconf <- matrix(unlist(allocconfdiff), ncol = 2, byrow = TRUE)

names(allocitact) <- unique(interactomedata2$Bait)

pheatmap::pheatmap(allocitact[[169]], color = magma(61, alpha = 0.8, direction = 1), cluster_rows = FALSE, cluster_cols = FALSE, treeheight_col = 0, display_numbers = TRUE, fontsize_number = 12, number_color = "darkgreen", number_format = "%g")


pheatmap::pheatmap(allocitact$UL8, color = magma(61, alpha = 0.8, direction = 1), cluster_rows = FALSE, cluster_cols = FALSE, treeheight_col = 0, display_numbers = TRUE, fontsize_number = 12, number_color = "darkgreen", number_format = "%g", main = "UL8 interactome", fontsize = 14)

pheatmap::pheatmap(allocitact$UL70, color = magma(61, alpha = 0.8, direction = 1), cluster_rows = FALSE, cluster_cols = FALSE, treeheight_col = 0, display_numbers = TRUE, fontsize_number = 12, number_color = "darkgreen", number_format = "%g", main = "UL70 interactome", fontsize = 14)

pheatmap::pheatmap(allocitact$UL148A, color = viridis(61, alpha = 0.8, direction = 1), cluster_rows = FALSE, cluster_cols = FALSE, treeheight_col = 0, display_numbers = TRUE, fontsize_number = 12, number_color = "darkgreen", number_format = "%g", main = "UL148A interactome", fontsize = 14)

```


```{r,}
allocintactactome <- vector(mode = "list", length(unique(interactomedata2$Bait)))
names(allocintactactome) <- unique(interactomedata2$Bait)


for (j in seq_along(allocintactactome)) {
  allocintactactome[[j]] <- cbind(getMarkerClasses(HCMV24_sub)[maxalloc1[nmitact[[j]]]], getMarkerClasses(HCMV24_sub)[maxalloc2[nmitact[[j]]]])
  colnames(allocintactactome[[j]]) <- c("MOCK", "HCMV")
  rownames(allocintactactome[[j]]) <- nmitact[[j]]
}


write.xlsx(allocintactactome, file="allocintactactome.xlsx", append = TRUE, row.names = TRUE)

```


```{r,}
size <- sapply(allocitact, sum)
pval <- vector(mode = "numeric", length = length(size))

for (j in seq_along(size))  {
  multisamps <- rmultinom(5000, size= size[j], prob = c(df_alloc))[c(1:9)^2,]
  pval[j] <- sum(colSums(multisamps) > difflocintact[j,1])/5000
}

rownames(difflocintact) <- unique(interactomedata2$Bait)

names(size) <- names(pval) <- unique(interactomedata2$Bait)

```

```{r,}
par(mfrow = c(1,3))
hist(colSums(rmultinom(5000, size= size["UL8"], prob = c(df_alloc))[c(1:9)^2,]), col = mycols[1], xlim = c(0, 15), main = "Simulated distribution UL8", xlab = "number of proteins")
abline(v =  difflocintact["UL8", 1], col = mycols[3], lwd = 4)

hist(colSums(rmultinom(5000, size= size["UL70"], prob = c(df_alloc))[c(1:9)^2,]), col = mycols[1], xlim = c(0, 15), main = "Simulated distribution UL70", xlab = "number of proteins")
abline(v =  difflocintact["UL70",1], col = mycols[3], lwd = 4)

hist(colSums(rmultinom(5000, size= size["UL148A"], prob = c(df_alloc))[c(1:9)^2,]), col = mycols[1], xlim = c(0, 30), main = "Simulated distribution UL148A", xlab = "number of proteins")
abline(v =  difflocintact["UL148A",1], col = mycols[3], lwd = 4)

```


```{r,}
interestingproteins <- size[size > 10 & pval < 0.05]

for (j in seq_along(interestingproteins)){
  colnames(allocitact[names(interestingproteins)][[j]]) <- getMarkerClasses(MOCK24_sub)[as.numeric(colnames(allocitact[names(interestingproteins)][[j]]))]
  rownames(allocitact[names(interestingproteins)][[j]]) <- getMarkerClasses(MOCK24_sub)[as.numeric(rownames(allocitact[names(interestingproteins)][[j]]))]
}

```


```{r,}
pheatmap::pheatmap(allocitact[names(interestingproteins)[[1]]][[1]], color = viridis(61, alpha = 0.8, direction = 1), cluster_rows = FALSE, cluster_cols = FALSE, treeheight_col = 0, display_numbers = TRUE, fontsize_number = 12, number_color = "lightblue", number_format = "%g", main = paste0(names(interestingproteins)[[1]]))

```


```{r,}
interestinglargeinter <- size[size > 10 & pval > 0.05]

for (j in seq_along(interestinglargeinter)){
  colnames(allocitact[names(interestinglargeinter)][[j]]) <- getMarkerClasses(MOCK24_sub)[as.numeric(colnames(allocitact[names(interestinglargeinter)][[j]]))]
  rownames(allocitact[names(interestinglargeinter)][[j]]) <- getMarkerClasses(MOCK24_sub)[as.numeric(rownames(allocitact[names(interestinglargeinter)][[j]]))]
}

```

```{r,}
pheatmap::pheatmap(allocitact[names(interestinglargeinter)[[1]]][[1]], color = viridis(61, alpha = 0.8, direction = 1), cluster_rows = FALSE, cluster_cols = FALSE, treeheight_col = 0, display_numbers = TRUE, fontsize_number = 12, number_color = "lightblue", number_format = "%g", main = paste0(names(interestinglargeinter)[[1]]))


names(nmitact) <- unique(interactomedata2$Bait)

write.csv(nmitact$UL148A, file = "UL148Ainteract.csv")

```


```{r,}

allocmatrix <- data.frame(cbind(getMarkerClasses(MOCK24_sub)[maxalloc1],
as.numeric(meanallocationProb1[cbind(maxalloc1,1:length(maxalloc1))]),
getMarkerClasses(MOCK24_sub)[maxalloc2],
as.numeric(meanallocationProb2[cbind(maxalloc2,1:length(maxalloc2))])))

rownames(allocmatrix) <- names(maxalloc1)
colnames(allocmatrix) <- c("Allocation Mock 24hpi", "localisation Prob MoCK", "Allocation HCMV infected 24hpi", "localisation prob HCMV" )

write.csv(allocmatrix, "allocationprobsHCMV24hpi.csv")

write.csv(fData(markerMSnSet(HCMV24_sub)), "HCMVstableprotiens.csv")


```

## pca plots for figure
```{r,}
plot2D(MOCK24_sub[,1:6], dims = c(1,4), grid = FALSE, main = "MOCK 24hpi")
plot2D(HCMV24_sub[,1:6], dims = c(1,4), grid = FALSE, main = "HCMV 24hpi")

```
```{r,}
require(dplyr)
allocproborg <- meanallocationProb1
markerdata <- fData(markerMSnSet(MOCK24_sub))$markers
allocprobmarker <- matrix(0, K, length(markerdata))
object <- MOCK24_sub

for (j in seq.int(K)) {
  
  allocprobmarker[j, markerdata == getMarkerClasses(object)[j]] <- 1
  
}
colnames(allocprobmarker) <- names(markerdata)
allocproborg[,rowMeans(bandleres_beltran24[[2]]$allocprob$cond1) < 0.95] <- 0 

allocproborg <- cbind(allocproborg, allocprobmarker)
colnames(allocproborg) <- c(colnames(meanallocationProb1), rownames(markerMSnSet(MOCK24_sub)))

gg1 <- spatial2D(object = MOCK24_sub, allocproborg = allocproborg, dims = c(1,4))


allocproborg <- meanallocationProb2
markerdata <- fData(markerMSnSet(HCMV24_sub))$markers
allocprobmarker <- matrix(0, K, length(markerdata))
object <- HCMV24_sub

for (j in seq.int(K)) {
  
  allocprobmarker[j, markerdata == getMarkerClasses(object)[j]] <- 1
  
}
colnames(allocprobmarker) <- names(markerdata)
allocproborg[,rowMeans(bandleres_beltran24[[2]]$allocprob$cond2) < 0.95] <- 0 

allocproborg <- cbind(allocproborg, allocprobmarker)
colnames(allocproborg) <- c(colnames(meanallocationProb1), rownames(markerMSnSet(HCMV24_sub)))

gg2 <- spatial2D(object = HCMV24_sub, allocproborg = allocproborg, dims = c(1,4))


.pca1 <- plot2D(MOCK24_sub[,1:6], dims = c(1,4), grid = FALSE, main = "MOCK 24hpi")
.pca2 <- plot2D(HCMV24_sub[,1:6], dims = c(1,4), grid = FALSE, main = "HCMV 24hpi")
gg1 <- gg1 + geom_point(data = data.frame(x = .pca1["Q8WTV0",1], y = .pca1["Q8WTV0",2]),
                   aes(x = x, y = y, z = 1), colour = "black", size = 3, shape = 18) + labs(title = "Mock 24hpi") + 
  annotate(geom="text", x = .pca1["Q8WTV0",1] + 0.5, y = .pca1["Q8WTV0",2], label= "Q8WTV0" ,
              color="purple3")
gg2 <- gg2 + geom_point(data = data.frame(x = .pca2["Q8WTV0",1], y = .pca2["Q8WTV0",2]),
                   aes(x = x, y = y, z = 1), colour = "black", size = 3, shape = 18) + labs(title = "HCMV 24hpi") + 
  annotate(geom="text", x = .pca2["Q8WTV0",1] + 0.5, y = .pca2["Q8WTV0",2], label= "Q8WTV0" ,
              color="purple3")

gg1/gg2

```


## Biological process enrichment
```{r,}
BPenrich <- read.table(file = "enrichment.Process.tsv", sep = '\t', header = FALSE)

## immune proccesses
immproc <- BPenrich[BPenrich$V2 %in% c("neutrophil degranulation", "granulocyte activation", "neutrophil mediated immunity", "leukocyte degranulation",
                          "leukocyte mediated immunity", "immune response", "antigen processing and presentation of exogenous peptide antigen via MHC class II",
                          "positive regulation of natural killer cell degranulation"),c(2,3:5)]

levels(immproc$V2) <- gsub("of", "of \n", levels(immproc$V2))

ggplot(data = immproc, aes(x = V2, y = -log10(V5))) + geom_bar(stat = "identity", fill = mycols[1]) + theme_bw() + theme(legend.position = "none", panel.grid.major = element_blank(), panel.grid.minor = element_blank())+ coord_flip() + ggtitle("Enriched Immune Proccesses") + xlab("GO Term") + ylab("-log10(p-value)")

## Transport
transport <- BPenrich[BPenrich$V2 %in% c("transport","protein transport","vesicle-mediated transport", "regulation of vesicle-mediated transport",
                          "transferrin transport", "endosomal transport"), c(2,3:5)]

levels(transport$V2) <- gsub("of", "of \n", levels(transport$V2))

ggplot(data = transport, aes(x = V2, y = -log10(V5))) + geom_bar(stat = "identity", fill = mycols[2]) + theme_bw() + theme(legend.position = "none", panel.grid.major = element_blank(), panel.grid.minor = element_blank())+ coord_flip() + ggtitle("Enriched Transport Terms") + xlab("GO Term") + ylab("-log10(p-value)")

## Transcription and translation,
transctranl <- BPenrich[BPenrich$V2 %in% c("cotranslational protein targeting to membrane", "SRP-dependent cotranslational protein targeting to membrane",
                          "translation", "mRNA catabolic process", "mRNA metabolic process"), c(2,3:5)]

levels(transctranl$V2) <- gsub("protein", "protein \n", levels(transctranl$V2))

ggplot(data = transctranl, aes(x = V2, y = -log10(V5))) + geom_bar(stat = "identity", fill = mycols[3]) + theme_bw() + theme(legend.position = "none", panel.grid.major = element_blank(), panel.grid.minor = element_blank())+ coord_flip() + ggtitle("Enriched Translation and Transcription Terms") + xlab("GO Term") + ylab("-log10(p-value)")


## viral process
viralprco <- BPenrich[BPenrich$V2 %in% c("viral process", "viral life cycle", "positive regulation of viral release from host cell", "positive regulation by host of viral process",
                          "regulation of viral life cycle", "regulation of viral process"), c(2,3:5)]

levels(viralprco$V2) <- gsub("of", "of \n", levels(viralprco$V2))

ggplot(data = viralprco, aes(x = V2, y = -log10(V5))) + geom_bar(stat = "identity", fill = mycols[4]) + theme_bw() + theme(legend.position = "none", panel.grid.major = element_blank(), panel.grid.minor = element_blank())+ coord_flip() + ggtitle("Enriched Viral Terms") + xlab("GO Term") + ylab("-log10(p-value)")


```

## Reactome pathways
```{r,}
reactome <- read.table(file = "enrichment.RCTM.tsv", sep = '\t', header = FALSE)

reactomedf <- reactome[reactome$V2 %in% c("Translation", "Viral mRNA Translation", "Vesicle-mediated transport",
                   "Membrane Trafficking", "Innate Immune System", "MHC class II antigen presentation",
                   "Cargo recognition for clathrin-mediated endocytosis", "Influenza Life Cycle", "Infectious disease", "Immune System", "Antigen processing-Cross presentation",
                   "Host Interactions of HIV factors", "HIV Infection", "Extracellular matrix organization", "Rab regulation of trafficking"),]

ggplot(data = reactomedf , aes(x = V2, y = -log10(V5))) + geom_bar(stat = "identity", fill = mycols[6]) + theme_bw() + theme(legend.position = "none", panel.grid.major = element_blank(), panel.grid.minor = element_blank())+ coord_flip() + ggtitle("Enriched Reactome Pathways") + xlab("Pathway") + ylab("-log10(p-value)")

write.csv(reactome, "HCMVreactome.csv")


```
````
