## Supplementary Code 1 for "Inferring differential subcellular localisation in comparative spatial proteomics using BANDLE": daviesb_analysis.Rmd

````
---
title: "AP-4 Knockout Davies"
author: "Oliver M. Crook"
date: "21 January 2020"
output: html_document
---

```{r,}
require(pRoloc)
require(pRolocdata)
require(ggplot2)
source("bandleFUN.R")
```

```{r,}
load("data/davies2018wt.rda")
load("data/davies2018ap4b1.rda")

```

```{r,}
davies2018wt <- filterNA(davies2018wt)
davies2018ap4b1 <- filterNA(davies2018ap4b1)
```

```{r,}
exprs(davies2018wt) <- log2(exprs(davies2018wt))
exprs(davies2018ap4b1) <- log2(exprs(davies2018ap4b1))
```


```{r,}
par(mfrow = c(2,3))
plot2D(davies2018wt[,1:5], dims = c(1,2))
plot2D(davies2018wt[,6:10], dims = c(1,2))
plot2D(davies2018ap4b1, dims = c(1,2))
getMarkerClasses(davies2018wt, fcol = "markers")

par(mfrow = c(1,1))
pRoloc::plotDist(davies2018wt, fcol = "markers")

exprs(davies2018wt)

subactin <- fData(davies2018wt)$markers == "Actin binding proteins"
fData(davies2018wt)[rownames(davies2018wt[subactin]), "markers"] <- "unknown"


subactin <- fData(davies2018ap4b1)$markers == "Actin binding proteins"
fData(davies2018ap4b1)[rownames(davies2018ap4b1[subactin]), "markers"] <- "unknown"

subnpc <- fData(davies2018ap4b1)$markers == "Nuclear pore complex"
fData(davies2018ap4b1)[rownames(davies2018ap4b1[subnpc]), "markers"] <- "unknown"

subnpc <- fData(davies2018wt)$markers == "Nuclear pore complex"
fData(davies2018wt)[rownames(davies2018wt[subnpc]), "markers"] <- "unknown"


K <- length(getMarkerClasses(davies2018wt))
dir_prior = diag(rep(1, K)) + matrix(0.02, nrow = K, ncol = K)
prior_pred_dir(object = davies2018wt, dir_prior = dir_prior)

pc_prior <- matrix(NA, ncol = 3, K)
pc_prior[seq.int(1:K), ] <- matrix(rep(c(0.05, 60, 100), each = K), ncol = 3)
```
```{r,}
setStockcol(paste0(getStockcol(),90))
mycols <- c("blue", "darkgreen", "orange", "purple", "pink", "brown", "gray", "cyan")
mycols <- sapply(mycols, function(x) alpha(x, 0.2))
```

```{r,}
plot2D(davies2018wt[,1:5], main = "HeLa Wild Type", dims = c(1,2), cex.lab = 1.5, cex.main = 1.5, grid = FALSE, mirrorX = FALSE, mirrorY = TRUE)
addLegend(davies2018wt, cex = 1.2, where = "topleft")

```
```{r,}
plot2D(davies2018ap4b1[,1:5], main = "HeLa AP-4 Knock Out", dims = c(1,2), cex.lab = 1.5, cex.main = 1.5, grid = FALSE)
addLegend(davies2018ap4b1[,1:5], cex = 1.2, where = "topleft")

```


```{r,}
## Load davies data
load("C:/Users/OllyC/Desktop/2019-DiffLoc/bandleres_daviesb4.rda")

par(mfrow = c(3,4))

mycols <- c("blue", "darkgreen", "orange", "purple", "pink", "brown", "gray", "cyan")
mycols <- sapply(mycols, function(x) alpha(x, 0.2))

par(mfrow = c(3,4))
for (j in seq_along(getMarkerClasses(davies2018ap4b1))) {
  for (i in seq_along(bandleres_daviesb4)){
    if (i ==  1) {
      M <- max(bandleres_daviesb4[[i]]$weights[j,j,]) + 0.05
      plot(bandleres_daviesb4[[i]]$weights[j,j,], col = mycols[[i]], type = "b", pch = 19, ylim = c(0, M), ylab = "mixing proportions")
    } else {
      points(bandleres_daviesb4[[i]]$weights[j,j,], col = mycols[[i]], type = "b", pch = 19)
    }    }
}
par(mfrow = c(3,4))
for (j in seq_along(getMarkerClasses(davies2018ap4b1))) {
  for (i in c(2,3,4,5,6)){
    if (i ==  2) {
      M <- max(bandleres_daviesb4[[i]]$weights[j,j,]) + 0.05
      plot(bandleres_daviesb4[[i]]$weights[j,j,], col = mycols[[i]], type = "b", pch = 19, ylim = c(0, M), ylab = "mixing proportions")
    } else {
      points(bandleres_daviesb4[[i]]$weights[j,j,], col = mycols[[i]], type = "b", pch = 19)
    }  
  }
}
```
```{r,}
require(coda)
r_hat <- vector(mode = "list", length = length(getMarkerClasses(davies2018ap4b1)))
for (j in seq_along(getMarkerClasses(davies2018ap4b1))) {
  r_hat[[j]] <- gelman.diag(mcmc.list(mcmc(bandleres_daviesb4[[2]]$weights[j,j,]),
                                      mcmc(bandleres_daviesb4[[3]]$weights[j,j,])), transform = TRUE)
}
unlist(r_hat)
```

```{r,}
par(mfrow = c(2,3))
rk <- rank(cbind(bandleres_daviesb4[[2]]$weights[j,j,],
                 bandleres_daviesb4[[3]]$weights[j,j,],
                 bandleres_daviesb4[[4]]$weights[j,j,],
                 bandleres_daviesb4[[5]]$weights[j,j,],
                 bandleres_daviesb4[[6]]$weights[j,j,]))

hist(rk[1:667], col = alpha("blue", 0.2), breaks = seq.int(from = 1, to = length(rk), length.out = 10), main = "rank plot", xlab = "rank")
hist(rk[668:1334], col = alpha("darkgreen", 0.2),breaks = seq.int(from = 1, to = length(rk), length.out = 10), main = "rank plot", xlab = "rank")
hist(rk[1335:2001], col = mycols[3],breaks = seq.int(from = 1, to = length(rk), length.out = 10), main = "rank plot", xlab = "rank")
hist(rk[2001:2668], col = mycols[4],breaks = seq.int(from = 1, to = length(rk), length.out = 10), main = "rank plot", xlab = "rank")
hist(rk[2668:3335], col = mycols[5],breaks = seq.int(from = 1, to = length(rk), length.out = 10), main = "rank plot", xlab = "rank")

rk <- rank(cbind(bandleres_daviesb4[[2]]$weights[j,j,],
                 bandleres_daviesb4[[3]]$weights[j,j,],
                 bandleres_daviesb4[[6]]$weights[j,j,]))
```


```{r,}
par(mfrow = c(1,3))
hist(rk[1:667], col = alpha("blue", 0.2), breaks = seq.int(from = 1, to = length(rk), length.out = 10), main = "rank plot", xlab = "rank")
hist(rk[668:1334], col = alpha("darkgreen", 0.2),breaks = seq.int(from = 1, to = length(rk), length.out = 10), main = "rank plot", xlab = "rank")
hist(rk[1335:2001], col = mycols[3],breaks = seq.int(from = 1, to = length(rk), length.out = 10), main = "rank plot", xlab = "rank")


```
```{r,}
probdir_daviesb4 <- probsameorganelle(object = davies2018ap4b1, params = bandleres_daviesb4[[2]])
#probdir_daviesb4_3 <- probsameorganelle(object = davies2018ap4b1, params = bandleres_daviesb4[[3]])
#probdir_daviesb4_6 <- probsameorganelle(object = davies2018ap4b1, params = bandleres_daviesb4[[6]])

#probdir_daviesb4 <- (probdir_daviesb4_1 + probdir_daviesb4_3 + probdir_daviesb4_6)/3

```


```{r,}
## filter out from marker that were removed
markersub <- c(rownames(davies2018wt[subactin]), rownames(davies2018wt[subnpc]))

## BANDLE dirichlet
par(mfrow = c(1,1))
y <- 1 - probdir_daviesb4[order(probdir_daviesb4)]/ncol(bandleres_daviesb4[[1]]$allocOut$cond11)
y <- y[!(names(y) %in% markersub)]


par(mfrow = c(1,1))
plot(y, pch = 19, ylim = c(0,1), ylab = "Differential Localisation Probability", xlab = "Rank", 
     col = getStockcol()[2], main = "Differential localisation rank", cex.lab = 1.5, cex.main = 2, cex.axis = 1.5)
```

```{r,}
par(mfrow = c(2,1))
.pca <- plot2D(object = davies2018wt, dims = c(1,2), grid = FALSE)
addLegend(where = "bottomright", object = davies2018wt, ncol = 2)
highlightOnPlot(object = .pca, foi = names(y[1:25]), col = "purple3", cex = 1.5, pch = 19, labels = TRUE)
.pca <- plot2D(object = davies2018ap4b1, dims = c(1,2))
highlightOnPlot(object = .pca, foi = names(y[1:25]), col = "purple3", cex = 1.5, pch = 19, labels = TRUE)

rownames(bandleres_daviesb4[[3]]$alloc$cond11) <- rownames(unknownMSnSet(davies2018wt))
rownames(bandleres_daviesb4[[3]]$alloc$cond21) <- rownames(unknownMSnSet(davies2018wt))

alloc1 <- apply(bandleres_daviesb4[[3]]$alloc$cond11[names(y[1:50]),],1, table)
alloc2 <- apply(bandleres_daviesb4[[3]]$alloc$cond21[names(y[1:50]),],1, table)

movers <- names(y[1:50])

write.csv(y, file = "davies_all2.csv")
```
```{r,}
library(UniProt.ws)
require(stringr)
require(dplyr)

species <- 9606
columns <- c("SEQUENCE", "SUBCELLULAR-LOCATIONS", "PROTEIN-NAMES")

up <- UniProt.ws(taxId=species)

keytype <- "UNIPROTKB"

# example keys
keys <- str_sub(rownames(davies2018wt), start = 1, end = 6)

protein_seq <- UniProt.ws::select(up, keys, columns, keytype) %>%
  mutate(motif=grepl(pattern='Y..[AVILMPFYW]', SEQUENCE))  

uniprotlocal <- protein_seq[,c(1,3)]
which.nuc <- grepl(uniprotlocal$`SUBCELLULAR-LOCATIONS`, pattern = " Nucle")
which.Nuc <- grepl(uniprotlocal$`SUBCELLULAR-LOCATIONS`, pattern = " nucle")
which.act <- grepl(uniprotlocal$`SUBCELLULAR-LOCATIONS`, pattern = " actin")
names(which.nuc) <- uniprotlocal[,1]
names(which.act) <- uniprotlocal[,1]
names(which.Nuc) <- uniprotlocal[,1]
to.filteruni <- names(which.nuc)[which.nuc]
to.filterNuc <- names(which.nuc)[which.Nuc]
to.filteract <- names(which.act)[which.act]

```

```{r,}
candmoves <- substr(names(y[1:80]), start = 1, stop = 6)

write.csv(candmoves[!(candmoves %in% c(to.filteruni, to.filterNuc, to.filteract))], file = "filtermoversdaviesuniprot.csv")

filteredmovers <- names(y[1:80][!(candmoves %in% c(to.filteruni, to.filteract))])
```

```{r,}
daviesallrank <- substr(names(y), start = 1, stop = 6)

write.csv(daviesallrank[!(candmoves %in% c(to.filteruni, to.filterNuc, to.filteract))], file = "daviesallrank.csv")

```


```{r,}
par(mfrow = c(1,1))
bootresdir <- bootstrapdiffloc(object = davies2018wt, params = bandleres_daviesb4[[2]], top = 25, Bootsample = 1000)
boxplot(t(1 - bootresdir/ncol(bandleres_daviesb4[[2]]$allocOut[[1]])), las = 2, col = mycols[3])

require(reshape2)
df <- t(1 - bootresdir/ncol(bandleres_daviesb4[[2]]$allocOut[[1]]))
df <- df[,!(colnames(df) %in% c(markersub, to.filteract, to.filterNuc, to.filteruni))]
colnames(df) <- str_sub(colnames(df), 1, 6)
df_long <- melt(df)
colnames(df_long) <- c("bootsample", "Protein", "probability")
df_long$diff.loc <- rep(0, nrow(df_long))


require(ggplot2)
p <- ggplot(df_long, aes(x = Protein, y = probability, fill = Protein)) + geom_boxplot(fill = mycols[4]) #+ scale_fill_manual(fill = rep(mycols[3],30))
p <- p + scale_y_continuous(limits = c(0.95, 1)) + theme(legend.position = "none")
p <- p + theme_classic() + theme(axis.text.x = element_text(angle = 90, size = 16), axis.text.y = element_text(size = 16),
                                 axis.title=element_text(size=20), plot.title = element_text(size = 22))  + theme(legend.position = "none")
p <- p + labs(title = "Rank plot of top differentially localised proteins",
              x = "Protein",
              y = "Differential Localisation Probability")
p 


```


```{r,}
par(mfrow = c(4,4))
for (j in seq_along(filteredmovers)) {
  mydata <- t(rbind(exprs(davies2018wt[filteredmovers[j],1:5]),
                    exprs(davies2018wt[filteredmovers[j],6:10])))
  M <- max(mydata) + 0.5
  m <- min(mydata) - 0.5
  matplot(mydata, col = mycols[1], pch = 19, lwd = 3, lty = 1,
          ylab = "abundance", type = "b", ylim = c(m, M), main = paste0(filteredmovers[j]))
  matplot(t(rbind(exprs(davies2018ap4b1[filteredmovers[j],1:5]),
                  exprs(davies2018ap4b1[filteredmovers[j],6:10]))), col = mycols[2], pch = 19, lwd = 3, lty = 1,
          ylab = "abundance", type = "b", add = TRUE)
}

```

```{r,}
require(viridis)
cols <- viridis(5)
par(mfrow = c(3,1))
for (j in seq_along(filteredmovers)[c(8,12,19)]) {
  mydata <- t(rbind(exprs(davies2018wt[filteredmovers[j],1:5]),
                    exprs(davies2018wt[filteredmovers[j],6:10])))
  M <- max(mydata) + 0.8
  m <- min(mydata) - 0.5
  matplot(mydata, col = cols[1], pch = 19, lwd = 3, lty = 1,
          ylab = "Normalised abundance", type = "b", ylim = c(m, M), main = paste0(filteredmovers[j]), xlab = "Fraction")
  matplot(t(rbind(exprs(davies2018ap4b1[filteredmovers[j],1:5]),
                  exprs(davies2018ap4b1[filteredmovers[j],6:10]))), col = cols[4], pch = 19, lwd = 5, lty = 1,
          ylab = "abundance", type = "b", add = TRUE, xlab = "Fraction")
}

```

```{r,}

cor.mat <- matrix(NA, 20, 20)
colnames(cor.mat) <- rownames(cor.mat) <- stringr::str_sub(filteredmovers[1:20], 1, 6)

cor.mat2 <- matrix(NA, 20, 20)
colnames(cor.mat2) <- rownames(cor.mat2) <- stringr::str_sub(filteredmovers[1:20], 1, 6)

for (j in 1:20){
  for (k in 1:20){
    cor.mat[j,k] <- cor.test(exprs(davies2018ap4b1[filteredmovers[j], 1:10]), exprs(davies2018ap4b1[filteredmovers[k], 1:10]), method = "spearman")$estimate
  }
}

for (j in 1:20){
  for (k in 1:20){
    cor.mat2[j,k] <- cor.test(c(exprs(davies2018wt[filteredmovers[j], 1:10]), exprs(davies2018ap4b1[filteredmovers[j], 1:10])),
                              c(exprs(davies2018wt[filteredmovers[j], 1:10]),exprs(davies2018ap4b1[filteredmovers[k], 1:10])), method = "spearman")$estimate
  }
}

df2 <- exprs(davies2018wt[filteredmovers[1:20], 1:10])
df <- exprs(davies2018ap4b1[filteredmovers[1:20], 1:10])
rownames(df) <- stringr::str_sub(filteredmovers[1:20], 1, 6)
rownames(df2) <- stringr::str_sub(filteredmovers[1:20], 1, 6)
colnames(df) <- paste0(rep("Fraction ", 10), rep(1:5, times = 2))
pmap <- pheatmap::pheatmap(df, cluster_cols = FALSE, clustering_distance_rows = "euclidean", scale = "row", color = viridis(100, alpha = 0.8, direction = 1))
pmap2 <- pheatmap::pheatmap(exdf2, cluster_cols = FALSE, clustering_distance_rows = "euclidean", scale = "row")

clust <- cutree(pmap$tree_row, k = 4)

cmat <- pheatmap::pheatmap(cor.mat, color = viridis(100, alpha = 0.8, direction = 1))
clust <- cutree(cmat$tree_row, k = 4)
```


```{r,}
spatial2D <- function(object,
                      allocproborg = allocproborg,
                      dims = c(1, 2),
                      cov.function = wendland.cov,
                      theta = 2,
                      derivative = 2,
                      k = 1,
                      breaks = c(0.99, 0.95, 0.9, 0.85, 0.8, 0.75),
                      aspect = 0.5,
                      mirrorX = FALSE,
                      mirrorY = FALSE) {
  
  # generate pca plot and create data from with probabilties
  .pca <- plot2D(object, dims = dims, plot = FALSE, mirrorX = mirrorX, mirrorY = mirrorY)
  probs <- data.frame(x = .pca[, 1], y = .pca[, 2], mcmc.prob = t(allocproborg)[rownames(object),])
  colnames(probs) <- c(c("x", "y"), getMarkerClasses(object))
  eigs <- colnames(.pca)
  
  # put data in appropriate long format
  probs.lst <- list()
  for(j in getMarkerClasses(object)) {
    probs.lst[[j]] <- probs[, c("x", "y", j)]
    colnames(probs.lst[[j]]) <- c("x", "y", "probability")
  }
  probs.lst.df <- plyr::ldply(probs.lst, .fun = function(x) x, .id = "organelle")
  
  # Create storage
  coords <- list()
  locations <- list()
  df <- list()
  # Create appropriate spatial grid
  for (j in getMarkerClasses(object)) {
    idxOrg <- c(probs.lst.df$organelle == j)
    coords[[j]] <- akima::interp(x = probs.lst.df$x[idxOrg],
                                 y = probs.lst.df$y[idxOrg],
                                 z = probs.lst.df$probability[idxOrg],
                                 extrap=FALSE, linear = TRUE, duplicate = TRUE) # interpolate onto appropriate grid
    coords[[j]]$z[is.na(coords[[j]]$z)] <- 0 # NaNs beyond data set to 0
    locations[[j]] <- cbind(rep(coords[[j]]$x, 40), rep(coords[[j]]$y, each = 40)) # get grid
    smoothedprobs <- fields::smooth.2d(coords[[j]]$z, x = locations[[j]],
                                       cov.function = cov.function,
                                       theta = theta,
                                       derivative = derivative, k = k) # spatial smoothing of probabilities
    # normalisation and formatting
    zmat <- matrix(smoothedprobs$z, ncol = 1)
    zmat <- zmat/max(zmat)
    df[[j]] <- data.frame(x = rep(smoothedprobs$x, 64), y = rep(smoothedprobs$y, each = 64), z = zmat)
  }
  # format data
  df.lst <- plyr::ldply(df, .fun = function(x) x, .id = "organelle") 
  df.lst <- df.lst %>%
    mutate(organelle = factor(organelle)) 
  K <- length(getMarkerClasses(object))
  cols <- getStockcol()[1:K] # get appropriate colours
  
  
  gg <- ggplot(
    data = df.lst,
    aes(x = x, y = y, z = z, color = organelle)) +
    coord_fixed() + 
    geom_contour(breaks = breaks, size = 1.2, aes(alpha = stat(level))) + 
    geom_point(alpha = 0) + 
    xlab(paste0(eigs[1])) + 
    ylab(paste0(eigs[2])) +
    scale_alpha(guide = "none") + 
    theme(legend.position = "right", 
          text = element_text(size = 12)) +
    scale_color_manual(values = cols) +
    scale_fill_manual(values = cols) +
    theme_minimal() + 
    theme(panel.grid.major = element_blank(), panel.grid.minor = element_blank(),
          aspect.ratio = aspect,
          panel.border = element_rect(colour = "black", fill = NA, size = 1),
          plot.title = element_text(hjust = 0.5, size = 20),
          legend.text=element_text(size = 14)) +
    ggtitle(label = "Spatial variation of localisation probabilities") 
  gg
  return(gg)
}
```

```{r,}
mcmcsampleprobs <- bandleres_daviesb4[[2]]$allocprob$cond11
dimnames(mcmcsampleprobs)[[1]] <- rownames(unknownMSnSet(davies2018wt))
dimnames(mcmcsampleprobs)[[3]] <- getMarkerClasses(davies2018wt)
markerdata <- fData(markerMSnSet(davies2018wt))$markers

allocproborg <- apply(bandleres_daviesb4[[2]]$allocprob$cond11, c(1), function(x) colMeans(x))
allocprobmarker <- matrix(0, K, length(markerdata))
object <- davies2018wt

for (j in seq.int(K)) {
  
  allocprobmarker[j, markerdata == getMarkerClasses(object)[j]] <- 1
  
}
colnames(allocprobmarker) <- names(markerdata)
allocproborg[,rowMeans(bandleres_daviesb4[[2]]$allocprob$cond11) < 0.95] <- 0 

allocproborg <- cbind(allocproborg, allocprobmarker)
colnames(allocproborg) <- c(dimnames(mcmcsampleprobs)[[1]], rownames(markerMSnSet(davies2018wt)))


.pca <- plot2D(davies2018wt[,1:5], dims = c(1,2), plot = FALSE, mirrorX = FALSE, mirrorY = TRUE)
gg <- spatial2D(object = davies2018wt[,1:5], allocproborg = allocproborg, dims = c(1,2), mirrorX = FALSE, mirrorY = TRUE, breaks = c(0.99, 0.95, 0.9, 0.85, 0.8, 0.75, 0.7,0.65))
gg <- gg + geom_point(data = data.frame(x = .pca[filteredmovers[c(8,12,19)],1], y = .pca[filteredmovers[c(8,12,19)],2]),
                   aes(x = x, y = y, z = 1), colour = "black", size = 3, shape = 18) + labs(title = "HeLa Wild Type") + 
  annotate(geom="text", x = .pca[filteredmovers[c(8,12,19)],1] + 0.5, y = .pca[filteredmovers[c(8,12,19)],2], label= c(filteredmovers[c(8,12,19)]),
              color="purple3")

mcmcsampleprobs <- bandleres_daviesb4[[2]]$allocprob$cond21
dimnames(mcmcsampleprobs)[[1]] <- rownames(unknownMSnSet(davies2018wt))
dimnames(mcmcsampleprobs)[[3]] <- getMarkerClasses(davies2018wt)
markerdata <- fData(markerMSnSet(davies2018wt))$markers

allocproborg <- apply(bandleres_daviesb4[[2]]$allocprob$cond21, c(1), function(x) colMeans(x))
allocprobmarker <- matrix(0, K, length(markerdata))
object <- davies2018wt

for (j in seq.int(K)) {
  
  allocprobmarker[j, markerdata == getMarkerClasses(object)[j]] <- 1
  
}
colnames(allocprobmarker) <- names(markerdata)
allocproborg[,rowMeans(bandleres_daviesb4[[2]]$allocprob$cond21) < 0.95] <- 0 

allocproborg <- cbind(allocproborg, allocprobmarker)
colnames(allocproborg) <- c(dimnames(mcmcsampleprobs)[[1]], rownames(markerMSnSet(davies2018wt)))


.pca <- plot2D(davies2018ap4b1[,1:5], dims = c(1,2), plot = FALSE)
gg2 <- spatial2D(object = davies2018ap4b1[,1:5],allocproborg = allocproborg, dims = c(1,2))
gg2 <- gg2 + geom_point(data = data.frame(x = .pca[filteredmovers[c(8,12,19)],1], y = .pca[filteredmovers[c(8,12,19)],2]),
                   aes(x = x, y = y, z = 1), colour = "black", size = 3, shape = 18)  + labs(title = "HeLa AP4 Knockout") + 
  annotate(geom="text", x = .pca[filteredmovers[c(8,12,19)],1] + 0.5, y = .pca[filteredmovers[c(8,12,19)],2] + c(0,0,-0.5), label= c(filteredmovers[c(8,12,19)]),
              color="purple3")
gg/gg2
```
````
