## Supplementary Code 1 for "Inferring differential subcellular localisation in comparative spatial proteomics using BANDLE": dynamicItzhak2016analysis.R

```
data(itzhak2016helaCtrl)
data(itzhak2016helaEgf)


helaCtrl <- filterNA(helaCtrl)
helaEgf <- filterNA(helaEgf)

idx <- intersect(rownames(helaCtrl), rownames(helaEgf))

helaCtrl_sub <- helaCtrl[idx, ]
helaEgf_sub <- helaEgf[idx, ]

itzhakdyn <- list(helaCtrl_sub[, 1:5],
                  helaCtrl_sub[, 6:10],
                  helaCtrl_sub[, 11:15],
                  helaEgf_sub[, 1:5],
                  helaEgf_sub[, 6:10],
                  helaEgf_sub[, 11:15])

mrIthzakres <- mrMethod(mydata = itzhakdyn)

plot(mrIthzakres$Mscore, mrIthzakres$Rscore)
text(mrIthzakres$Mscore, mrIthzakres$Rscore, labels=names(mrIthzakres$Mscore), cex= 1)
abline(h = 0.9, col = "blue", lwd = 2)
abline(v = 2, col = "purple", lwd = 2)

names(mrIthzakres$Mscore)[mrIthzakres$Mscore > 2 & mrIthzakres$Rscore > 0.9]

bigMover <- c("P49753", "Q09666", "P00533", "P62993", "Q16513", "P29353", "Q6NUQ4")

par(mfrow = c(1,2))
.pca <- plot2D(helaCtrl_sub, dims = c(1,3))
highlightOnPlot(.pca, foi = bigMover, labels = TRUE)
highlightOnPlot(.pca, foi = names(Mscore[Mscore > 0.6 & Rscore > 0.7]), labels = TRUE)

.pca <- plot2D(helaEgf_sub, dims = c(1,3))
highlightOnPlot(.pca, foi = bigMover, labels = TRUE)

K <- length(getMarkerClasses(helaCtrl_sub))
dir_prior = diag(rep(1, K)) + matrix(0.02, nrow = K, ncol = K)
prior_pred_dir(object = helaCtrl_sub, dir_prior = dir_prior)


pgIthzak <- diffLoc(object_cond1 = c(helaCtrl_sub[, 1:5], helaCtrl_sub[, 6:10], helaCtrl_sub[, 11:15]),
                    object_cond2 = c(helaEgf_sub[, 1:5], helaEgf_sub[, 6:10], helaEgf_sub[, 11:15]),
                    numIter = 5000, burnin = 2500, pg = FALSE, dir_prior = dir_prior, hypLearn = "MH")

itzres <- probsameorganelle(object = helaCtrl_sub, params = pgIthzak)

par(mfrow = c(1,1))
itzDiff <- 1 - itzres[order(itzres)]/ncol(pgIthzak$allocOut[[1]])
plot(itzDiff[1:20], pch = 24, ylim = c(0.8,1.1))
text(itzDiff[1:20], cand, srt = 45)
abline(h = 0.99, col = "blue", lwd = 2)
abline(h = 0.95, col = "purple", lwd = 2)
cand <- names(itzDiff[itzDiff > 0.99])

par(mfrow = c(1,2))
.pca1 <- plot2D(helaCtrl_sub, dims = c(1,2), method = "t-SNE")
highlightOnPlot(object = .pca1, foi = cand, labels = TRUE)
.pca2 <- plot2D(helaEgf_sub, dims = c(1,2), method = "t-SNE")
highlightOnPlot(object = .pca2, foi = cand, labels = TRUE)
addLegend(where = "bottomright", helaCtrl_sub)

par(mfrow = c(1,1))
bootresitz <- bootstrapdiffloc(object = helaCtrl_sub, params = pgIthzak, top = 100, Bootsample = 1000)
boxplot(t(1 - bootresitz/ncol(pgIthzak$allocOut[[1]])), las = 2, col = getStockcol()[3])
```
