## Supplementary Code 1 for "Inferring differential subcellular localisation in comparative spatial proteomics using BANDLE": ithzakprocess.R

```
## Analysis of Bandle of dynamics Itzhak data

data(itzhak2016helaCtrl)
data(itzhak2016helaEgf)
load("data/itzhak2016helaCtrl.rda")
load("data/itzhak2016helaEgf.rda")

helaCtrl <- itzhak2016helaCtrl
helaEgf <- itzhak2016helaEgf

helaCtrl <- filterNA(helaCtrl)
helaEgf <- filterNA(helaEgf)

idx <- intersect(rownames(helaCtrl), rownames(helaEgf))

helaCtrl_sub <- helaCtrl[idx, ]
helaEgf_sub <- helaEgf[idx, ]

itzhakdyn <- list(helaCtrl_sub[, 1:5],
                  helaCtrl_sub[, 6:10],
                  helaCtrl_sub[, 11:15],
                  helaEgf_sub[, 1:5],
                  helaEgf_sub[, 6:10],
                  helaEgf_sub[, 11:15])


## load mcmc run
load("C:/Users/OllyC/Desktop/2019-DiffLoc/bandleres_Itzhak.rda")
load("bandleres_Itzhak2.rda")

dim(bandleres_Itzhak[[1]]$allocprob$cond11)

par(mfrow = c(3,4))

mycols <- c("blue", "darkgreen", "orange", "purple", "pink", "brown", "gray", "cyan")
mycols <- sapply(mycols, function(x) alpha(x, 0.2))

for (j in seq_along(getMarkerClasses(helaCtrl))) {
  for (i in seq_along(bandleres_Itzhak)){
    if (i ==  1) {
    M <- max(bandleres_Itzhak[[i]]$weights[j,j,]) + 0.05
    plot(bandleres_Itzhak[[i]]$weights[j,j,], col = mycols[[i]], type = "b", pch = 19, ylim = c(0, M), xlab = "mixing proportions")
    } else {
    points(bandleres_Itzhak[[i]]$weights[j,j,], col = mycols[[i]], type = "b", pch = 19)
    }  
  }
}

par(mfrow = c(3,4))
for (j in seq_along(getMarkerClasses(helaCtrl))) {
  for (i in c(1,2,5,6)){
    if (i ==  1) {
      M <- max(bandleres_Itzhak[[i]]$weights[j,j,]) + 0.05
      plot(bandleres_Itzhak[[i]]$weights[j,j,], col = mycols[[i]], type = "b", pch = 19, ylim = c(0, M), xlab = "mixing proportions")
    } else {
      points(bandleres_Itzhak[[i]]$weights[j,j,], col = mycols[[i]], type = "b", pch = 19)
    }  
  }
}

par(mfrow = c(3,4))
for (j in seq_along(getMarkerClasses(helaCtrl))) {
  for (i in c(2,6)){
    if (i ==  2) {
      M <- max(bandleres_Itzhak[[i]]$weights[j,j,]) + 0.05
      plot(bandleres_Itzhak[[i]]$weights[j,j,], col = mycols[[i]], type = "b", pch = 19, ylim = c(0, M), xlab = "mixing proportions")
    } else {
      points(bandleres_Itzhak[[i]]$weights[j,j,], col = mycols[[i]], type = "b", pch = 19)
    }  
  }
}

par(mfrow = c(3,4))

mycols <- c("blue", "darkgreen", "orange", "purple", "pink", "brown", "gray", "cyan")
mycols <- sapply(mycols, function(x) alpha(x, 0.2))

for (j in seq_along(getMarkerClasses(helaCtrl))) {
  for (i in seq_along(bandleres_Itzhak2)){
    if (i ==  1) {
      M <- max(bandleres_Itzhak2[[i]]$weights[j,j,]) + 0.05
      plot(bandleres_Itzhak2[[i]]$weights[j,j,], col = mycols[[i]], type = "b", pch = 19, ylim = c(0, M), ylab = "mixing proportions",
           xlab = "mcmc iteration")
    } else {
      points(bandleres_Itzhak2[[i]]$weights[j,j,], col = mycols[[i]], type = "b", pch = 19)
    }  
  }
}

par(mfrow = c(3,4))
for (j in seq_along(getMarkerClasses(helaCtrl))) {
  for (i in c(5,8)){
    if (i ==  5) {
      M <- max(bandleres_Itzhak2[[i]]$weights[1,j,]) + .05
      plot(bandleres_Itzhak2[[i]]$weights[1,j,], col = mycols[[i]], type = "b", pch = 19, ylim = c(0, M), xlab = "mixing proportions")
    } else {
      points(bandleres_Itzhak2[[i]]$weights[1,j,], col = mycols[[i]], type = "b", pch = 19)
    }  
  }
}

r_hat <- vector(mode = "list", length = length(getMarkerClasses(helaCtrl)))
for (j in seq_along(getMarkerClasses(helaCtrl))){
 r_hat[[j]] <- gelman.diag(mcmc.list(mcmc(bandleres_Itzhak2[[5]]$weights[1,j,]), mcmc(bandleres_Itzhak2[[8]]$weights[1,j,] )), transform = TRUE)
}
unlist(r_hat)

probdir <- probsameorganelle(object = helaCtrl_sub, params = bandleres_Itzhak[[8]])

## BANDLE dirichlet
par(mfrow = c(1,1))
y <- 1 - probdir[order(probdir)]/ncol(bandleres_Itzhak[[8]]$allocOut$cond11)
plot(y, pch = 19, ylim = c(0,1), ylab = "Differential Localisation Probability", xlab = "Rank", 
     col = getStockcol()[2], main = "Rank plot from BANDLE", cex.lab = 1.5, cex.main = 2, cex.axis = 1.5)

par(mfrow = c(1,1))
bootresdir <- bootstrapdiffloc(object = tan2009rep[[1]], params = resdyndir, top = 30, Bootsample = 1000)
boxplot(t(1 - bootresdir/ncol(resdyndir$allocOut[[1]])), las = 2, col = getStockcol()[3])

par(mfrow = c(2,1))
.pca <- plot2D(object = helaCtrl_sub, dims = c(1,3))
addLegend(where = "topleft", object = helaCtrl_sub, ncol = 2)
highlightOnPlot(object = .pca, foi = names(y[1:25]), col = "purple3", cex = 1.5, pch = 19, labels = TRUE)
highlightOnPlot(object = .pca, foi = "P04920", col = "purple3", cex = 1.5, pch = 19, labels = TRUE)
.pca <- plot2D(object = helaEgf_sub, dims = c(1,3))
highlightOnPlot(object = .pca, foi = "P04920", col = "purple3", cex = 1.5, pch = 19, labels = TRUE)


## look at changes in phosphoralation
## hypothesis: proteins with greatest increase in phosphorlaytion more likely to be differentially localised

ovlap <- combined_replicates$UniprotAc[combined_replicates$UniprotAc %in% names(y)]
diffprob <- y[ovlap]
phos <- combined_replicates[combined_replicates$UniprotAc %in% ovlap,c(1,4:6)]
phos <- phos %>% distinct(UniprotAc, .keep_all = TRUE)
ovlp <- phos$UniprotAc

mrresitz <- mrMethod(itzhakdyn)

par(mfrow = c(1,3))
plot(diffprob[phos$UniprotAc] ~ abs(phos$ave_max_diff))
plot(diffprob[phos$UniprotAc] ~ phos$max_diff.x)
plot(diffprob[phos$UniprotAc] ~ phos$max_diff.y)
cor(abs(phos$ave_max_diff), diffprob[phos$UniprotAc], method = "kendall")
abline(lm(diffprob[phos$UniprotAc] ~ phos$ave_max_diff))
plot(mrresitz$Mscore[phos$UniprotAc] ~ abs(phos$ave_max_diff))
cor(phos$ave_max_diff, mrresitz$Mscore[ovlap], method = "kendall")
abline(lm(mrresitz$Mscore[ovlap] ~ phos$ave_max_diff))
plot(mrresitz$Rscore[ovlap] ~ phos$ave_max_diff)

par(mfrow = c(1,2))
boxplot(phos$ave_max_diff[diffprob < 0.99], phos$ave_max_diff[diffprob > 0.99], col = alpha(getStockcol(), 0.5))
boxplot(phos$ave_max_diff[mrresitz$Mscore[ovlap] < 2 | mrresitz$Mscore[ovlap] < 0.9], 
        phos$ave_max_diff[mrresitz$Mscore[ovlap] > 2 & mrresitz$Rscore[ovlap] > 0.9], col = alpha(getStockcol(), 0.5))
boxplot(phos$ave_max_diff[mrresitz$Rscore[ovlap] < 0], phos$ave_max_diff[mrresitz$Mscore[ovlap] > 0], col = alpha(getStockcol(), 0.5))
t.test(phos$ave_max_diff[diffprob < 0.999], phos$ave_max_diff[diffprob > 0.999])
t.test(phos$ave_max_diff[mrresitz$Mscore[ovlap] < 2], phos$ave_max_diff[mrresitz$Mscore[ovlap] > 2])
t.test(phos$ave_max_diff[mrresitz$Rscore[ovlap] < 0], phos$ave_max_diff[mrresitz$Mscore[ovlap] > 0])

B <- 10000
cor1 <- cor2 <- rep(0, B)
for(i in 1:B){
  samp <- sample(length(diffprob), length(diffprob), TRUE)  
  cor1[i] <- cor(phos$ave_max_diff[samp], diffprob[samp], method = "kendall")
  cor2[i] <- cor(phos$ave_max_diff[samp], mrresitz$Mscore[ovlap][samp], method = "kendall")
}
par(mfrow = c(1,1))
hist(cor1, col = alpha(getStockcol()[1], 0.5), breaks = 30, ylim = c(0,2000))
hist(cor2, col = alpha(getStockcol()[2], 0.5), add = TRUE, breaks = 30)
.hist <- hist(cor1 - cor2, col = alpha(getStockcol()[3], 0.5), breaks = 30, ylim = c(0,1000))
quantile(cor1 - cor2, c(0.025,0.925))


require(reshape2)
df <- t(1 - bootresdir/ncol(resdyndir$allocOut[[1]]))
df_long <- melt(df)
colnames(df_long) <- c("bootsample", "Protein", "probability")
df_long$diff.loc <- rep(0, nrow(df_long))
df_long$diff.loc[df_long$Protein %in% perm1_names] <- 1

fill.col <- colnames(df) %in% perm1_names
fill.col[fill.col == TRUE] <- getStockcol()[4]
fill.col[fill.col == FALSE] <- getStockcol()[2]

require(ggplot2)
p <- ggplot(df_long, aes(x = Protein, y = probability, fill = Protein)) + geom_boxplot() + scale_fill_manual(values = fill.col)
p <- p + scale_y_continuous(limits = c(0.6, 1)) + theme(legend.position = "none")
p <- p + theme_classic() + theme(axis.text.x = element_text(angle = 90, colour = fill.col, size = 16), axis.text.y = element_text(size = 16),
                                 axis.title=element_text(size=20), plot.title = element_text(size = 22))  + theme(legend.position = "none")
p <- p + labs(title = "Rank plot of top 30 differentially localised proteins",
              x = "Protein",
              y = "Differential Localisation Probability")
p 


## analysis with new prior
load("C:/Users/OllyC/Desktop/2019-DiffLoc/bandleres_Itzhak3.rda")

mycols <- c("blue", "darkgreen", "orange", "purple", "pink", "brown", "gray", "cyan")
mycols <- sapply(mycols, function(x) alpha(x, 0.2))

par(mfrow = c(3,4))
for (j in seq_along(getMarkerClasses(helaCtrl))) {
  for (i in seq_along(bandleres_Itzhak3)){
    if (i ==  1) {
      M <- max(bandleres_Itzhak3[[i]]$weights[j,j,]) + 0.05
      plot(bandleres_Itzhak3[[i]]$weights[j,j,], col = mycols[[i]], type = "b", pch = 19, ylim = c(0, M), ylab = "mixing proportions",
           xlab = "mcmc iteration")
    } else {
      points(bandleres_Itzhak3[[i]]$weights[j,j,], col = mycols[[i]], type = "b", pch = 19)
    }  
  }
}


par(mfrow = c(3,4))
for (j in seq_along(getMarkerClasses(helaCtrl))) {
  for (i in c(1,3,4,7,8)){
    if (i ==  1) {
      M <- max(bandleres_Itzhak3[[i]]$weights[j,j,]) + .05
      plot(bandleres_Itzhak3[[i]]$weights[j,j,], col = mycols[[i]], type = "b", pch = 19, ylim = c(0, M), xlab = "mixing proportions")
    } else {
      points(bandleres_Itzhak3[[i]]$weights[j,j,], col = mycols[[i]], type = "b", pch = 19)
    }  
  }
}

par(mfrow = c(3,4))
for (j in seq_along(getMarkerClasses(helaCtrl))) {
  for (i in c(1,3,7)){
    if (i ==  1) {
      M <- max(bandleres_Itzhak3[[i]]$weights[j,j,]) + .05
      plot(bandleres_Itzhak3[[i]]$weights[j,j,], col = mycols[[i]], type = "b", pch = 19, ylim = c(0, M), xlab = "mixing proportions")
    } else {
      points(bandleres_Itzhak3[[i]]$weights[j,j,], col = mycols[[i]], type = "b", pch = 19)
    }  
  }
}


par(mfrow = c(3,4))
for (j in seq_along(getMarkerClasses(helaCtrl))) {
  for (i in c(4,8)){
    if (i ==  4) {
      M <- max(bandleres_Itzhak3[[i]]$weights[j,j,]) + .05
      plot(bandleres_Itzhak3[[i]]$weights[j,j,], col = mycols[[i]], type = "b", pch = 19, ylim = c(0, M), xlab = "mixing proportions")
    } else {
      points(bandleres_Itzhak3[[i]]$weights[j,j,], col = mycols[[i]], type = "b", pch = 19)
    }  
  }
}

r_hat <- vector(mode = "list", length = length(getMarkerClasses(helaCtrl)))
for (j in seq_along(getMarkerClasses(helaCtrl))){
  r_hat[[j]] <- gelman.diag(mcmc.list(mcmc(bandleres_Itzhak3[[3]]$weights[j,j,]),
                                      mcmc(bandleres_Itzhak3[[7]]$weights[j,j,])), transform = TRUE)
}
unlist(r_hat)

probdir <- probsameorganelle(object = helaCtrl_sub, params = bandleres_Itzhak3[[3]])

## BANDLE dirichlet
par(mfrow = c(1,1))
y <- 1 - probdir[order(probdir)]/ncol(bandleres_Itzhak3[[1]]$allocOut$cond11)
plot(y, pch = 19, ylim = c(0,1), ylab = "Differential Localisation Probability", xlab = "Rank", 
     col = getStockcol()[2], main = "Rank plot from BANDLE", cex.lab = 1.5, cex.main = 2, cex.axis = 1.5)

par(mfrow = c(1,1))
bootresdir <- bootstrapdiffloc(object = tan2009rep[[1]], params = resdyndir, top = 30, Bootsample = 1000)
boxplot(t(1 - bootresdir/ncol(resdyndir$allocOut[[1]])), las = 2, col = getStockcol()[3])

par(mfrow = c(2,1))
.pca <- plot2D(object = helaCtrl_sub, dims = c(1,3))
addLegend(where = "topleft", object = helaCtrl_sub, ncol = 2)
highlightOnPlot(object = .pca, foi = names(y[1:41]), col = "purple3", cex = 1.5, pch = 19, labels = TRUE)
highlightOnPlot(object = .pca, foi = "P04920", col = "purple3", cex = 1.5, pch = 19, labels = TRUE)
.pca <- plot2D(object = helaEgf_sub, dims = c(1,3))
highlightOnPlot(object = .pca, foi = names(y[1:41]), col = "purple3", cex = 1.5, pch = 19, labels = TRUE)


par(mfrow = c(3,4))
res <- lapply(itzhakdyn, function(x) fitGPmaternPC(x, fcol = "markers", nu = 2, hyppar = pc_prior)) 

par(mfrow = c(1,2))
plot2D(helaCtrl_sub, dims = c(1,3))
plot2D(helaEgf_sub, dims = c(1,3))

qsepres <- pRoloc::QSep(object = helaCtrl_sub)
heatmap(qsep(qsepres), Rowv = NA, Colv = NA)
min(qsep(qsepres)) # golgi and peroxisome

subgol <- fData(helaCtrl_sub)$markers == "Golgi"
fData(helaCtrl_sub)[rownames(helaCtrl_sub[subgol]), "markers"] <- "unknown"
subper <- fData(helaCtrl_sub)$markers == "Peroxisome"
fData(helaCtrl_sub)[rownames(helaCtrl_sub[subper]), "markers"] <- "unknown"
qsepres <- pRoloc::QSep(object = helaCtrl_sub)
heatmap(qsep(qsepres), Rowv = NA, Colv = NA)
min(qsep(qsepres)) # golgi and peroxisome


par(mfrow = c(3,4))
plot.new()
# use visual posterior predictive check
res <- lapply(itzhakdyn, function(x) fitGPmaternPC(x, fcol = "markers", nu = 2, hyppar = pc_prior))
itzhak2016hypers2 <- res
save(itzhak2016hypers2, file = "itzhak2016hypers2.rda")

## bandle attempt 4 itzhak
load("C:/Users/OllyC/Desktop/2019-DiffLoc/bandleres_Itzhak4.rda")

mycols <- c("blue", "darkgreen", "orange", "purple", "pink", "brown", "gray", "cyan")
mycols <- sapply(mycols, function(x) alpha(x, 0.2))

par(mfrow = c(3,4))
for (j in seq_along(getMarkerClasses(helaCtrl_sub))) {
  for (i in seq_along(bandleres_Itzhak4)){
    if (i ==  1) {
      M <- max(bandleres_Itzhak4[[i]]$weights[j,j,]) + 0.05
      plot(bandleres_Itzhak4[[i]]$weights[j,j,], col = mycols[[i]], type = "b", pch = 19, ylim = c(0, M), ylab = "mixing proportions",
           xlab = "mcmc iteration")
    } else {
      points(bandleres_Itzhak4[[i]]$weights[j,j,], col = mycols[[i]], type = "b", pch = 19)
    }  
  }
}


par(mfrow = c(3,4))
for (j in seq_along(getMarkerClasses(helaCtrl_sub))) {
  for (i in c(1,3,4,7,8)){
    if (i ==  1) {
      M <- max(bandleres_Itzhak4[[i]]$weights[j,j,]) + .05
      plot(bandleres_Itzhak4[[i]]$weights[j,j,], col = mycols[[i]], type = "b", pch = 19, ylim = c(0, M), xlab = "mixing proportions")
    } else {
      points(bandleres_Itzhak4[[i]]$weights[j,j,], col = mycols[[i]], type = "b", pch = 19)
    }  
  }
}

par(mfrow = c(3,4))
for (j in seq_along(getMarkerClasses(helaCtrl_sub))) {
  for (i in c(1,3,7)){
    if (i ==  1) {
      M <- max(bandleres_Itzhak4[[i]]$weights[j,j,]) + .05
      plot(bandleres_Itzhak4[[i]]$weights[j,j,], col = mycols[[i]], type = "b", pch = 19, ylim = c(0, M), xlab = "mixing proportions")
    } else {
      points(bandleres_Itzhak4[[i]]$weights[j,j,], col = mycols[[i]], type = "b", pch = 19)
    }  
  }
}


par(mfrow = c(3,4))
for (j in seq_along(getMarkerClasses(helaCtrl_sub))) {
  for (i in c(4,8)){
    if (i ==  4) {
      M <- max(bandleres_Itzhak4[[i]]$weights[j,j,]) + .05
      plot(bandleres_Itzhak4[[i]]$weights[j,j,], col = mycols[[i]], type = "b", pch = 19, ylim = c(0, M), xlab = "mixing proportions")
    } else {
      points(bandleres_Itzhak4[[i]]$weights[j,j,], col = mycols[[i]], type = "b", pch = 19)
    }  
  }
}

require(coda)
r_hat <- vector(mode = "list", length = length(getMarkerClasses(helaCtrl_sub)))
for (j in seq_along(getMarkerClasses(helaCtrl_sub))) {
  r_hat[[j]] <- gelman.diag(mcmc.list(mcmc(bandleres_Itzhak4[[3]]$weights[j,j,]),
                                      mcmc(bandleres_Itzhak4[[7]]$weights[j,j,])), transform = TRUE)
}
unlist(r_hat)

par(mfrow = c(2,3))
rk <- rank(cbind(bandleres_Itzhak4[[1]]$weights[j,j,],
                 bandleres_Itzhak4[[2]]$weights[j,j,],
                 bandleres_Itzhak4[[3]]$weights[j,j,],
                 bandleres_Itzhak4[[4]]$weights[j,j,],
                 bandleres_Itzhak4[[5]]$weights[j,j,],
                 bandleres_Itzhak4[[7]]$weights[j,j,]))

hist(rk[1:1000], col = alpha("blue", 0.2), breaks = seq.int(from = 1, to = length(rk), length.out = 10), main = "rank plot")
hist(rk[1001:2000], col = alpha("darkgreen", 0.2),breaks = seq.int(from = 1, to = length(rk), length.out = 10), main = "rank plot")
hist(rk[2001:3000], col = mycols[3],breaks = seq.int(from = 1, to = length(rk), length.out = 10), main = "rank plot")
hist(rk[3001:4000], col = mycols[4],breaks = seq.int(from = 1, to = length(rk), length.out = 10), main = "rank plot")
hist(rk[4001:5000], col = mycols[5],breaks = seq.int(from = 1, to = length(rk), length.out = 10), main = "rank plot")
hist(rk[5001:6000], col = mycols[6],breaks = seq.int(from = 1, to = length(rk), length.out = 10), main = "rank plot")


probdir <- probsameorganelle(object = helaCtrl_sub, params = bandleres_Itzhak4[[3]])

## BANDLE dirichlet
par(mfrow = c(1,1))
y <- 1 - probdir[order(probdir)]/ncol(bandleres_Itzhak4[[1]]$allocOut$cond11)
plot(y, pch = 19, ylim = c(0,1), ylab = "Differential Localisation Probability", xlab = "Rank", 
     col = getStockcol()[2], main = "Rank plot from BANDLE", cex.lab = 1.5, cex.main = 2, cex.axis = 1.5)

par(mfrow = c(2,1))
.pca <- plot2D(object = helaCtrl_sub, dims = c(1,3))
addLegend(where = "topleft", object = helaCtrl_sub, ncol = 2)
highlightOnPlot(object = .pca, foi = names(y[1:25]), col = "purple3", cex = 1.5, pch = 19, labels = TRUE)
highlightOnPlot(object = .pca, foi = "P04920", col = "purple3", cex = 1.5, pch = 19, labels = TRUE)
.pca <- plot2D(object = helaEgf_sub, dims = c(1,3))
highlightOnPlot(object = .pca, foi = names(y[1:25]), col = "purple3", cex = 1.5, pch = 19, labels = TRUE)


## look at changes in phosphoralation
## hypothesis: proteins with greatest increase in phosphorlaytion more likely to be differentially localised

ovlap <- combined_replicates$UniprotAc[combined_replicates$UniprotAc %in% names(y)]
diffprob <- y[ovlap]
phos <- combined_replicates[combined_replicates$UniprotAc %in% ovlap,c(1,4:6)]
phos <- phos %>% distinct(UniprotAc, .keep_all = TRUE)
ovlp <- phos$UniprotAc


y[ovlap]
rep1 <- data.frame(rep1)
rep1_sub <- rep1[rep1$UniprotAc %in% names(table(rep1$UniprotAc)[table(rep1$UniprotAc) > 1]),]
rep2 <- data.frame(rep2)
rep2_sub <- rep2[rep2$UniprotAc %in% names(table(rep2$UniprotAc)[table(rep2$UniprotAc) > 1]),]
matplot(t(rep2_sub[rep2_sub$UniprotAc == "P00533",4:7]), type = "l", col = mycols[[1]], pch = 19, lwd = 3, lty = 1)
plot(colMeans(rep1[rep1$UniprotAc == "P00533",4:7]))

matplot(t(rep1[rep1$UniprotAc == "P04920",4:7]), type = "l", col = mycols[[1]], pch = 19, lwd = 3, lty = 1)
plot(colMeans(rep1[rep1$UniprotAc == "P04920",4:7]))


par(mfrow = c(1,2))
matplot(t(rep1[rep1$UniprotAc %in% names(diffprob[phos$UniprotAc][diffprob[phos$UniprotAc] > 0 ]),4:7]),
         type = "l", col = mycols[[1]], pch = 19, lwd = 3, lty = 1)
matplot(t(rep2[rep1$UniprotAc %in% names(diffprob[phos$UniprotAc][diffprob[phos$UniprotAc] > 0 ]),4:7]),
         type = "l", col = mycols[[1]], pch = 19, lwd = 3, lty = 1)

par(mfrow = c(1,2))
matplot(t(rep1[rep1$UniprotAc == "P00533", 4:7]), type = "l", col = mycols[[1]], pch = 19, lwd = 3, lty = 1)
matplot(t(rep2[rep2$UniprotAc == "P00533", 4:7]), type = "l", col = mycols[[1]], pch = 19, lwd = 3, lty = 1)

par(mfrow = c(1,2))
plot(colMeans(rep1[rep1$UniprotAc == "P00533", 4:7]), type = "l", col = mycols[[1]], pch = 19, lwd = 3, lty = 1)
plot(colMeans(rep2[rep2$UniprotAc == "P00533", 4:7]), type = "l", col = mycols[[1]], pch = 19, lwd = 3, lty = 1)

par(mfrow = c(1,2))
matplot(t(rep1[rep1$UniprotAc == "Q6NZI2", 4:7]), type = "l", col = mycols[[1]], pch = 19, lwd = 3, lty = 1)
matplot(t(rep2[rep2$UniprotAc == "Q6NZI2", 4:7]), type = "l", col = mycols[[1]], pch = 19, lwd = 3, lty = 1)

par(mfrow = c(1,2))
plot(colMeans(rep1[rep1$UniprotAc == "Q09666", 4:7]), type = "l", col = mycols[[1]], pch = 19, lwd = 3, lty = 1)
plot(colMeans(rep2[rep2$UniprotAc == "Q09666", 4:7]), type = "l", col = mycols[[1]], pch = 19, lwd = 3, lty = 1)


par(mfrow = c(1,3))

rep1_sub$UniprotAc

plot(diffprob[phos$UniprotAc][diffprob[phos$UniprotAc] > 0 ] ~ phos$ave_max_diff[tosub])
plot(diffprob[phos$UniprotAc] ~ phos$max_diff.x)
plot(diffprob[phos$UniprotAc] ~ phos$max_diff.y)
cor(abs(phos$ave_max_diff), diffprob[phos$UniprotAc], method = "kendall")
abline(lm(diffprob[phos$UniprotAc] ~ phos$ave_max_diff))
plot(mrresitz$Mscore[phos$UniprotAc] ~ abs(phos$ave_max_diff))
cor(phos$ave_max_diff, mrresitz$Mscore[ovlap], method = "kendall")
abline(lm(mrresitz$Mscore[ovlap] ~ phos$ave_max_diff))
plot(mrresitz$Rscore[ovlap] ~ phos$ave_max_diff)

par(mfrow = c(1,2))
boxplot(phos$ave_max_diff[diffprob == 0], phos$ave_max_diff[diffprob > 0], col = alpha(getStockcol(), 0.5))
boxplot(phos$ave_max_diff[mrresitz$Mscore[ovlap] < 2 | mrresitz$Mscore[ovlap] < 0.9], 
        phos$ave_max_diff[mrresitz$Mscore[ovlap] > 2 & mrresitz$Rscore[ovlap] > 0.9], col = alpha(getStockcol(), 0.5))
boxplot(phos$ave_max_diff[mrresitz$Rscore[ovlap] < 0], phos$ave_max_diff[mrresitz$Mscore[ovlap] > 0], col = alpha(getStockcol(), 0.5))
t.test(phos$ave_max_diff[diffprob < 0.999], phos$ave_max_diff[diffprob > 0.999])
t.test(phos$ave_max_diff[mrresitz$Mscore[ovlap] < 2], phos$ave_max_diff[mrresitz$Mscore[ovlap] > 2])

## Look at different phosophorlyation dataset
egfPhosphotc <- read.csv("C:/Users/OllyC/Desktop/2019-DiffLoc/EGFphosphotimecoursedata.csv")
egf.summary <- egfPhosphotc[,c(1,2,39:45)]
egf.summary[,3:9] <- as.matrix(egf.summary[,3:9])
colnames(egf.summary)[3:9] <- c(2, 4, 8, 16, 32, 64, 128)

par(mfrow = c(2,2))
plot(as.numeric(egf.summary[1,3:9]), x = c(2, 4, 8, 16, 32, 64, 128), type = "b", pch = 19, col = mycols[2], lwd = 4, 
     ylab = "log 2 fold change", xlab = "mins after egf stimulation")
plot(as.numeric(egf.summary[2,3:9]), x = c(2, 4, 8, 16, 32, 64, 128), type = "b", pch = 19, col = mycols[2], lwd = 4, 
     ylab = "log 2 fold change", xlab = "mins after egf stimulation")
plot(as.numeric(egf.summary[3,3:9]), x = c(2, 4, 8, 16, 32, 64, 128), type = "b", pch = 19, col = mycols[2], lwd = 4, 
     ylab = "log 2 fold change", xlab = "mins after egf stimulation")
plot(as.numeric(egf.summary[4,3:9]), x = c(2, 4, 8, 16, 32, 64, 128), type = "b", pch = 19, col = mycols[2], lwd = 4, 
     ylab = "log 2 fold change", xlab = "mins after egf stimulation")

# Compute largest difference in fold change
diff.fold.change <- vector(mode = "numeric", length = nrow(egf.summary))

diff.fold.change <- apply(egf.summary, 1, function(z) max(as.numeric(z[3:9]) - min(as.numeric(z[3:9]))))
var.fold.change <- apply(egf.summary, 1, function(z) var(z[3:9]))
mean.fold.change <- apply(egf.summary, 1, function(z) mean(as.numeric(z[3:9])))
max.fold.change <- apply(egf.summary, 1, function(z) max(as.numeric(z[3:9])))

hist(diff.fold.change)
hist(var.fold.change)
hist(mean.fold.change)
hist(max.fold.change)

protein.namestc <- str_sub(egf.summary[,2], start = 1, end = 6)
protein.namessp <- str_sub(names(y), start = 1, end = 6)

egf.pvalues <- egfPhosphotc$min.tukey.pval.firstprev
names(egf.pvalues) <- protein.namestc

egf.pvalues

# 195 protein matches
name.match <- protein.namessp %in% protein.namestc
matched.proteins <- protein.namessp[name.match]
sum(protein.namessp %in% protein.namestc)

egf.pvalues.matched <- egf.pvalues[matched.proteins]

y.shortname <- y
names(y.shortname) <- protein.namessp

names(diff.fold.change) <- protein.namestc
names(var.fold.change) <- protein.namestc
names(mean.fold.change) <- protein.namestc
names(max.fold.change) <- protein.namestc
y.shortname[matched.proteins]

plot(y.shortname[matched.proteins] ~ diff.fold.change[matched.proteins])

Mscore <- mrresitz$Mscore
names(Mscore) <- str_sub(names(Mscore), start = 1, end = 6)
plot(rank(Mscore[matched.proteins]) ~ diff.fold.change[matched.proteins])

MRscore <- mrresitz$MRscore
names(MRscore) <- str_sub(names(MRscore), start = 1, end = 6)
plot(MRscore[matched.proteins] ~ diff.fold.change[matched.proteins])

par(mfrow = c(1,2))
plot(y.shortname[matched.proteins] ~ var.fold.change[matched.proteins])
plot(MRscore[matched.proteins] ~ var.fold.change[matched.proteins])


plot(rank(y.shortname[matched.proteins]) ~ max.fold.change[matched.proteins])
plot(rank(MRscore[matched.proteins]) ~ max.fold.change[matched.proteins])

r1 <- cor(y.shortname[matched.proteins], diff.fold.change[matched.proteins], method = "spearman")
r2 <- cor(MRscore[matched.proteins], diff.fold.change[matched.proteins], method = "spearman")

r1.var <- cor(y.shortname[matched.proteins], var.fold.change[matched.proteins], method = "spearman")
r2.var <- cor(MRscore[matched.proteins], var.fold.change[matched.proteins], method = "spearman")

r1 <- cor(y.shortname[matched.proteins], mean.fold.change[matched.proteins], method = "spearman")
r2 <- cor(MRscore[matched.proteins], mean.fold.change[matched.proteins], method = "spearman")

r1 <- cor(y.shortname[matched.proteins], max.fold.change[matched.proteins], method = "spearman")
r2 <- cor(MRscore[matched.proteins], max.fold.change[matched.proteins], method = "spearman")

upabovehalfrnak <- rank(y.shortname[matched.proteins]) > quantile(rank(y.shortname[matched.proteins]), prob = 0.9)

boxplot(diff.fold.change[names(upabovehalfrnak)[upabovehalfrnak]], 
        diff.fold.change[names(upabovehalfrnak)[!upabovehalfrnak]])

par(mfrow = c(1,2))
plot(y.shortname[matched.proteins],diff.fold.change[matched.proteins])
plot(MRscore[matched.proteins],diff.fold.change[matched.proteins])


B <- 5000
cor1 <- cor2 <- rep(0, B)
n <- length(diff.fold.change[matched.proteins])
for(i in 1:B){
  samp <- sample(n, n, TRUE)  
  cor1[i] <- cor(diff.fold.change[matched.proteins][samp], y.shortname[matched.proteins][samp], method = "spearman")
  cor2[i] <- cor(diff.fold.change[matched.proteins][samp], MRscore[matched.proteins][samp], method = "spearman")
}

hist(cor1, col = mycols[1], breaks = 30, ylim = c(0,1000))
hist(cor2, col = mycols[2], add = TRUE, breals = 30)
hist(cor1 - cor2, col = mycols[3])

B <- 5000
cor1 <- cor2 <- rep(0, B)
n <- length(diff.fold.change[matched.proteins])
for(i in 1:B){
  samp <- sample(n, n, TRUE)  
  cor1[i] <- cor(var.fold.change[matched.proteins][samp], y.shortname[matched.proteins][samp], method = "spearman")
  cor2[i] <- cor(var.fold.change[matched.proteins][samp], MRscore[matched.proteins][samp], method = "spearman")
}

hist(cor1, col = mycols[1], breaks = 30, ylim = c(0,2000))
hist(cor2, col = mycols[2], add = TRUE, breals = 30)
hist(cor1 - cor2, col = mycols[3])
quantile(cor1 - cor2, prob = c(0.025, 0.975))


r1.prime <- log((1+r1)/(1 - r1))/2
r2.prime <- log((1+r2)/(1 - r2))/2

n1 <- length(y.shortname[matched.proteins])
n2 <- length(MRscore[matched.proteins])

S <- sqrt(1/(n1 -3) + 1/(n2 - 3))

z <- (r1.prime - r2.prime)/S

## have a look at top 20 ranked by methods
top10 <- names(y.shortname[matched.proteins][1:10])
top10itz <- names(MRscore[matched.proteins][1:10])

par(mfrow = c(1, 2))
plot(diff.fold.change[top10], y.shortname[top10])
plot(diff.fold.change[top10itz], MRscore[top10itz])

r1 <- cor(y.shortname[top10], diff.fold.change[top10], method = "spearman")
r2 <- cor(MRscore[top10itz], diff.fold.change[top10itz], method = "spearman")

B <- 10000
cor1 <- cor2 <- rep(0, B)
n <- length(diff.fold.change[top10])
for(i in 1:B){
  samp <- sample(n, n, TRUE)  
  cor1[i] <- cor(y.shortname[top10][samp], diff.fold.change[top10][samp], method = "spearman")
  cor2[i] <- cor(MRscore[top10itz][samp], diff.fold.change[top10itz][samp], method = "spearman")
}

hist(cor1, col = mycols[1], breaks = 30, ylim = c(0,2000))
hist(cor2, col = mycols[2], add = TRUE, breaks = 30)
hist(cor1 - cor2, col = mycols[3])
quantile(cor1 - cor2, prob = c(0.025, 0.975), na.rm = TRUE)

sum(egf.pvalues.matched[top10] < 0.05)
sum(egf.pvalues.matched[names(y.shortname[matched.proteins][-c(1:10)])] > 0.05)

cttble <- matrix(c(7,71,3,114), ncol = 2, byrow = TRUE)


sum(egf.pvalues.matched[top10itz] > 0.05)
sum(egf.pvalues.matched[names(MRscore[matched.proteins][-c(1:10)])] < 0.05)

cor.test(y.shortname[top10], diff.fold.change[top10], method = "spearman")
cor.test(MRscore[top10itz], diff.fold.change[top10itz], method = "spearman")
par(mfrow = c(1,1))
hist(cor1, col = mycols[1], breaks = 30, ylim = c(0,3000))
hist(cor2, col = mycols[2], add = TRUE, breaks = 30)
```
