## Supplementary Code 1 for "Inferring differential subcellular localisation in comparative spatial proteomics using BANDLE": Itzhak2016Process.Rmd

````
---
title: "EGF Stimulation Itzhak"
author: "Oliver M. Crook"
date: "21 January 2020"
output: html_document
---

```{r,}
require(pRoloc)
require(pRolocdata)
require(ggplot2)
source("bandleFUN.R")
```

```{r,}
load("C:/Users/OllyC/Desktop/2019-DiffLoc/bandleres_Itzhak5.rda")
```


```{r,}
data(itzhak2016helaCtrl)
data(itzhak2016helaEgf)
load("data/itzhak2016helaCtrl.rda")
load("data/itzhak2016helaEgf.rda")

helaCtrl <- itzhak2016helaCtrl
helaEgf <- itzhak2016helaEgf

helaCtrl <- filterNA(helaCtrl)
helaEgf <- filterNA(helaEgf)

idx <- intersect(rownames(helaCtrl), rownames(helaEgf))

helaCtrl_sub <- helaCtrl[idx, ]
helaEgf_sub <- helaEgf[idx, ]

itzhakdyn <- list(helaCtrl_sub[, 1:5],
                  helaCtrl_sub[, 6:10],
                  helaCtrl_sub[, 11:15],
                  helaEgf_sub[, 1:5],
                  helaEgf_sub[, 6:10],
                  helaEgf_sub[, 11:15])

subgol <- fData(helaCtrl_sub)$markers == "Golgi"
fData(helaCtrl_sub)[rownames(helaCtrl_sub[subgol]), "markers"] <- "unknown"
subper <- fData(helaCtrl_sub)$markers == "Peroxisome"
fData(helaCtrl_sub)[rownames(helaCtrl_sub[subper]), "markers"] <- "unknown"
subnuc <- fData(helaCtrl_sub)$markers == "Nuclear pore complex"
fData(helaCtrl_sub)[rownames(helaCtrl_sub[subnuc]), "markers"] <- "unknown"

subgol <- fData(helaEgf_sub)$markers == "Golgi"
fData(helaEgf_sub)[rownames(helaEgf_sub[subgol]), "markers"] <- "unknown"
subper <- fData(helaEgf_sub)$markers == "Peroxisome"
fData(helaEgf_sub)[rownames(helaEgf_sub[subper]), "markers"] <- "unknown"
subnuc <- fData(helaEgf_sub)$markers == "Nuclear pore complex"
fData(helaEgf_sub)[rownames(helaEgf_sub[subnuc]), "markers"] <- "unknown"


```

```{r,}
mycols <- c("blue", "darkgreen", "orange", "purple", "pink", "brown", "gray", "cyan")
mycols <- sapply(mycols, function(x) alpha(x, 0.2))
```


```{r,}
par(mfrow = c(3,4))
for (j in seq_along(getMarkerClasses(helaCtrl_sub))) {
  for (i in seq_along(bandleres_Itzhak5)){
    if (i ==  1) {
      M <- max(bandleres_Itzhak5[[i]]$weights[j,j,]) + 0.05
      plot(bandleres_Itzhak5[[i]]$weights[j,j,], col = mycols[[i]], type = "b", pch = 19, ylim = c(0, M), ylab = "mixing proportions",
           xlab = "mcmc iteration")
    } else {
      points(bandleres_Itzhak5[[i]]$weights[j,j,], col = mycols[[i]], type = "b", pch = 19)
    }  
  }
}


par(mfrow = c(3,4))
for (j in seq_along(getMarkerClasses(helaCtrl_sub))) {
  for (i in c(1,3,4,7,8)){
    if (i ==  1) {
      M <- max(bandleres_Itzhak5[[i]]$weights[j,j,]) + .05
      plot(bandleres_Itzhak5[[i]]$weights[j,j,], col = mycols[[i]], type = "b", pch = 19, ylim = c(0, M), ylab = "mixing proportions")
    } else {
      points(bandleres_Itzhak5[[i]]$weights[j,j,], col = mycols[[i]], type = "b", pch = 19)
    }  
  }
}

par(mfrow = c(3,4))
for (j in seq_along(getMarkerClasses(helaCtrl_sub))) {
  for (i in c(1,3,7)){
    if (i ==  1) {
      M <- max(bandleres_Itzhak5[[i]]$weights[j,j,]) + .05
      plot(bandleres_Itzhak5[[i]]$weights[j,j,], col = mycols[[i]], type = "b", pch = 19, ylim = c(0, M), ylab = "mixing proportions")
    } else {
      points(bandleres_Itzhak5[[i]]$weights[j,j,], col = mycols[[i]], type = "b", pch = 19)
    }  
  }
}


par(mfrow = c(3,4))
for (j in seq_along(getMarkerClasses(helaCtrl_sub))) {
  for (i in c(4,8)){
    if (i ==  4) {
      M <- max(bandleres_Itzhak5[[i]]$weights[j,j,]) + .05
      plot(bandleres_Itzhak5[[i]]$weights[j,j,], col = mycols[[i]], type = "b", pch = 19, ylim = c(0, M), ylab = "mixing proportions")
    } else {
      points(bandleres_Itzhak5[[i]]$weights[j,j,], col = mycols[[i]], type = "b", pch = 19)
    }  
  }
}
```


```{r,}
require(coda)
r_hat <- vector(mode = "list", length = length(getMarkerClasses(helaCtrl_sub)))
for (j in seq_along(getMarkerClasses(helaCtrl_sub))) {
  r_hat[[j]] <- gelman.diag(mcmc.list(mcmc(bandleres_Itzhak5[[1]]$weights[j,j,]),
                                      mcmc(bandleres_Itzhak5[[7]]$weights[j,j,])), transform = TRUE)
}
unlist(r_hat)
```

```{r,}
par(mfrow = c(2,3))
rk <- rank(cbind(bandleres_Itzhak5[[1]]$weights[j,j,],
                 bandleres_Itzhak5[[2]]$weights[j,j,],
                 bandleres_Itzhak5[[3]]$weights[j,j,],
                 bandleres_Itzhak5[[4]]$weights[j,j,],
                 bandleres_Itzhak5[[5]]$weights[j,j,],
                 bandleres_Itzhak5[[7]]$weights[j,j,]))

hist(rk[1:1000], col = alpha("blue", 0.2), breaks = seq.int(from = 1, to = length(rk), length.out = 10), xlab = "rank", main = "rank plot chain 1")
hist(rk[1001:2000], col = alpha("darkgreen", 0.2),breaks = seq.int(from = 1, to = length(rk), length.out = 10), xlab = "rank", main = "rank plot chain 2")
hist(rk[2001:3000], col = mycols[3],breaks = seq.int(from = 1, to = length(rk), length.out = 10), xlab = "rank", main = "rank plot chain 3")
hist(rk[3001:4000], col = mycols[4],breaks = seq.int(from = 1, to = length(rk), length.out = 10), xlab = "rank", main = "rank plot chain 4")
hist(rk[4001:5000], col = mycols[5],breaks = seq.int(from = 1, to = length(rk), length.out = 10), xlab = "rank", main = "rank plot chain 5")
hist(rk[5001:6000], col = mycols[6],breaks = seq.int(from = 1, to = length(rk), length.out = 10), xlab = "rank", main = "rank plot chain 7")
```

```{r,}
probdir <- probsameorganelle(object = helaCtrl_sub, params = bandleres_Itzhak5[[7]])
```

```{r,}
## filter out markers that were removed
markersub <- c(rownames(helaCtrl_sub[subgol]), rownames(helaCtrl_sub[subper]), rownames(helaCtrl_sub[subnuc]))


## BANDLE dirichlet
par(mfrow = c(1,1))
y <- 1 - probdir[order(probdir)]/ncol(bandleres_Itzhak5[[1]]$allocOut$cond11)
y <- y[!(names(y) %in% markersub)]
plot(y, pch = 19, ylim = c(0,1), ylab = "Differential Localisation Probability", xlab = "Rank", 
     col = getStockcol()[2], main = "Differential Localisation Rank", cex.lab = 1.5, cex.main = 2, cex.axis = 1.5)
```

```{r,}
write.csv(names(y)[1:11], file = "itzhak2016hit.csv")
```


```{r,}
par(mfrow = c(1,1))
bootresdir <- bootstrapdiffloc(object = helaCtrl_sub, params = bandleres_Itzhak5[[7]], top = 30, Bootsample = 1000)
boxplot(t(1 - bootresdir/ncol(bandleres_Itzhak5[[7]]$allocOut[[1]])), las = 2, col = mycols[3])

require(reshape2)
df <- t(1 - bootresdir/ncol(bandleres_Itzhak5[[7]]$allocOut[[1]]))
df <- df[,!(colnames(df) %in%markersub)]
df_long <- melt(df)
colnames(df_long) <- c("bootsample", "Protein", "probability")
df_long$diff.loc <- rep(0, nrow(df_long))


require(ggplot2)
p <- ggplot(df_long, aes(x = Protein, y = probability, fill = Protein)) + geom_boxplot(fill = mycols[4]) #+ scale_fill_manual(fill = rep(mycols[3],30))
p <- p + scale_y_continuous(limits = c(0.5, 1)) + theme(legend.position = "none")
p <- p + theme_classic() + theme(axis.text.x = element_text(angle = 90, size = 16), axis.text.y = element_text(size = 16),
                                 axis.title=element_text(size=20), plot.title = element_text(size = 22))  + theme(legend.position = "none")
p <- p + labs(title = "Rank plot of top differentially localised proteins",
              x = "Protein",
              y = "Differential Localisation Probability")
p 


```
```{r,}
rownames(bandleres_Itzhak5[[7]]$alloc$cond11) <- rownames(unknownMSnSet(helaCtrl_sub))
rownames(bandleres_Itzhak5[[7]]$alloc$cond21) <- rownames(unknownMSnSet(helaEgf_sub))

alloc1 <- apply(bandleres_Itzhak5[[7]]$alloc$cond11[names(y[1:11]),],1, table)
alloc2 <- apply(bandleres_Itzhak5[[7]]$alloc$cond21[names(y[1:11]),],1, table)

```

```{r,}
library(UniProt.ws)
require(stringr)
require(dplyr)

species <- 9606
columns <- c("SEQUENCE", "SUBCELLULAR-LOCATIONS", "PROTEIN-NAMES")

up <- UniProt.ws(taxId=species)

keytype <- "UNIPROTKB"

# example keys
keys <- str_sub(rownames(helaCtrl_sub), start = 1, end = 6)

protein_seq <- UniProt.ws::select(up, keys, columns, keytype) %>%
  mutate(motif=grepl(pattern='Y..[AVILMPFYW]', SEQUENCE))  

uniprotlocal <- protein_seq[,c(1,3)]
which.nuc <- grepl(uniprotlocal$`SUBCELLULAR-LOCATIONS`, pattern = " nucle")
names(which.nuc) <- uniprotlocal[,1]
to.filteruni <- names(which.nuc)[which.nuc]
```

```{r,}
#y.filter <- names(alloc2)[sapply(alloc2, function(x) length(names(x))) != sapply(alloc2, function(x) sum(names(x) %in% "9"))]
y.filter <- names(y[1:11])
```


```{r,}
par(mfrow = c(2,1))
.pca <- plot2D(object = helaCtrl_sub, dims = c(1,3))
addLegend(where = "topleft", object = helaCtrl_sub, ncol = 2)
highlightOnPlot(object = .pca, foi = y.filter, col = "purple3", cex = 1.5, pch = 19, labels = TRUE)
.pca <- plot2D(object = helaEgf_sub, dims = c(1,3))
highlightOnPlot(object = .pca, foi = y.filter, col = "purple3", cex = 1.5, pch = 19, labels = TRUE)
```

```{r,}
source('C:/Users/OllyC/Desktop/2019-DiffLoc/MRmethod.R')
mrresitz <- mrMethod(itzhakdyn)

```
```{r,}
mrcols <- rep(mycols[1], length(mrresitz$Rscore))
names(mrcols) <- names(mrresitz$MRscore)
mrcols[names(mrresitz$Mscore[mrresitz$Mscore > 2 & mrresitz$Rscore > 0.9])] <- mycols[3]              
plot(mrresitz$Mscore, mrresitz$Rscore, col = mrcols, pch = 19, xlab = "M Score", ylab = "R Score", main = "MR Method HeLa EGF")
abline(h = 0.9, col = mycols[2], lwd = 4)
abline(v = 2, col = mycols[2], lwd = 4)
```

```{r,}
par(mfrow = c(2,1))
.pca <- plot2D(object = helaCtrl_sub, dims = c(1,3))
addLegend(where = "topleft", object = helaCtrl_sub, ncol = 2)
highlightOnPlot(object = .pca, foi = names(mrresitz$Mscore[mrresitz$Mscore > 2 & mrresitz$Rscore > 0.9]), col = "purple3", cex = 1.5, pch = 19, labels = TRUE)
.pca <- plot2D(object = helaEgf_sub, dims = c(1,3))
highlightOnPlot(object = .pca, foi = names(mrresitz$Mscore[mrresitz$Mscore > 2 & mrresitz$Rscore > 0.9]), col = "purple3", cex = 1.5, pch = 19, labels = TRUE)
```

```{r,}
par(mfrow = c(3,4))
for (j in seq_along(names(y[1:10]))) {
  mydata <- t(rbind(exprs(helaCtrl_sub[names(y)[j],1:5]),
                    exprs(helaCtrl_sub[names(y)[j],6:10]),
                    exprs(helaCtrl_sub[names(y)[j],11:15])))
  M <- max(mydata) + 0.5
  m <- min(mydata) - 0.5
  matplot(mydata, col = mycols[1], pch = 19, lwd = 3, lty = 1,
          ylab = "abundance", type = "b", ylim = c(m, M), main = paste0(names(y)[j]))
  matplot(t(rbind(exprs(helaEgf_sub[names(y)[j],1:5]),
                    exprs(helaEgf_sub[names(y)[j],6:10]),
                    exprs(helaEgf_sub[names(y)[j],11:15]))), col = mycols[2], pch = 19, lwd = 3, lty = 1,
          ylab = "abundance", type = "b", add = TRUE)
}


```
```{r,}
mrnames <- names(mrresitz$Mscore[mrresitz$Mscore > 2 & mrresitz$Rscore > 0.9])
par(mfrow = c(2,3))
for (j in seq_along(mrnames)) {
  mydata <- t(rbind(exprs(helaCtrl_sub[mrnames[j],1:5]),
                    exprs(helaCtrl_sub[mrnames[j],6:10]),
                    exprs(helaCtrl_sub[mrnames[j],11:15])))
  M <- max(mydata) + 0.5
  m <- min(mydata) - 0.5
  matplot(mydata, col = mycols[1], pch = 19, lwd = 3, lty = 1,
          ylab = "abundance", type = "b", ylim = c(m, M), main = paste0(mrnames[j]))
  matplot(t(rbind(exprs(helaEgf_sub[mrnames[j],1:5]),
                    exprs(helaEgf_sub[mrnames[j],6:10]),
                    exprs(helaEgf_sub[mrnames[j],11:15]))), col = mycols[2], pch = 19, lwd = 3, lty = 1,
          ylab = "abundance", type = "b", add = TRUE)
}


```


```{r,}
## Look at different phosophorlyation dataset
egfPhosphotc <- read.csv("C:/Users/OllyC/Desktop/2019-DiffLoc/EGFphosphotimecoursedata.csv")
egf.summary <- egfPhosphotc[,c(1,2,39:45)]
egf.summary[,3:9] <- as.matrix(egf.summary[,3:9])
colnames(egf.summary)[3:9] <- c(2, 4, 8, 16, 32, 64, 128)

par(mfrow = c(2,2))
plot(as.numeric(egf.summary[1,3:9]), x = c(2, 4, 8, 16, 32, 64, 128), type = "b", pch = 19, col = mycols[2], lwd = 4, 
     ylab = "log 2 fold change", xlab = "mins after egf stimulation", main = paste0(egf.summary[1,2]))
plot(as.numeric(egf.summary[2,3:9]), x = c(2, 4, 8, 16, 32, 64, 128), type = "b", pch = 19, col = mycols[2], lwd = 4, 
     ylab = "log 2 fold change", xlab = "mins after egf stimulation", main = paste0(egf.summary[2,2]))
plot(as.numeric(egf.summary[3,3:9]), x = c(2, 4, 8, 16, 32, 64, 128), type = "b", pch = 19, col = mycols[2], lwd = 4, 
     ylab = "log 2 fold change", xlab = "mins after egf stimulation", main = paste0(egf.summary[3,2]))
plot(as.numeric(egf.summary[4,3:9]), x = c(2, 4, 8, 16, 32, 64, 128), type = "b", pch = 19, col = mycols[2], lwd = 4, 
     ylab = "log 2 fold change", xlab = "mins after egf stimulation", main = paste0(egf.summary[4,2]))

```

```{r,}
# Compute largest difference in fold change
diff.fold.change <- vector(mode = "numeric", length = nrow(egf.summary))

diff.fold.change <- apply(egf.summary, 1, function(z) max(as.numeric(z[3:7]) - min(as.numeric(z[3:7]))))

```


```{r,}
# 195 protein matches
protein.namestc <- str_sub(egf.summary[,2], start = 1, end = 6)
protein.namessp <- str_sub(names(y), start = 1, end = 6)
name.match <- protein.namessp %in% protein.namestc
matched.proteins <- protein.namessp[name.match]
sum(protein.namessp %in% protein.namestc)
MRscore <- mrresitz$MRscore

y.shortname <- y
names(y.shortname) <- protein.namessp

names(diff.fold.change) <- protein.namestc
y.shortname[matched.proteins]


## have a look at top 10 ranked by methods
top10 <- names(y.shortname[matched.proteins][1:10])
top10itz <- names(MRscore[names(MRscore) %in% matched.proteins][1:10])

par(mfrow = c(1, 2))
plot(diff.fold.change[top10], y.shortname[top10])
plot(diff.fold.change[top10itz], MRscore[top10itz])

r1 <- cor(y.shortname[top10], diff.fold.change[top10], method = "spearman")
r2 <- cor(MRscore[top10itz], diff.fold.change[top10itz], method = "spearman")

B <- 5000
cor1 <- cor2 <- rep(0, B)
n <- length(diff.fold.change[top10])
for(i in 1:B){
  samp <- sample(n, n, TRUE)  
  cor1[i] <- cor(y.shortname[top10][samp], diff.fold.change[top10][samp], method = "spearman")
  cor2[i] <- cor(MRscore[top10itz][samp], diff.fold.change[top10itz][samp], method = "spearman")
}

hist(cor1, col = mycols[1], breaks = 30, ylim = c(0,2000))
hist(cor2, col = mycols[2], add = TRUE, breaks = 30)
hist(cor1 - cor2, col = mycols[3])
quantile(cor1 - cor2, prob = c(0.025, 0.975), na.rm = TRUE)


cor.test(y.shortname[top10], diff.fold.change[top10], method = "spearman")
cor.test(MRscore[top10itz], diff.fold.change[top10itz], method = "spearman")
par(mfrow = c(1,1))
hist(cor1, col = mycols[1], breaks = 30, ylim = c(0,1000))
hist(cor2, col = mycols[2], add = TRUE, breaks = 30)

cor <- cbind(cor1,cor2)
colnames(cor) <- c("BANDLE", "MR")
boxplot(cbind(cor), col = c(mycols[1],mycols[2]), main = "Boostrap distribution of correlations")

quantile(cor1, probs = c(0.025,0.5, 0.975), na.rm = TRUE)
quantile(cor2, probs = c(0.025,0.5, 0.975), na.rm = TRUE)
```
```{r,}
which(protein.namestc %in% top10)
par(mfrow = c(2,2))
plot(as.numeric(egf.summary[87,3:9]), x = c(2, 4, 8, 16, 32, 64, 128), type = "b", pch = 19, col = mycols[2], lwd = 4, 
     ylab = "log 2 fold change", xlab = "mins after egf stimulation")
plot(as.numeric(egf.summary[96,3:9]), x = c(2, 4, 8, 16, 32, 64, 128), type = "b", pch = 19, col = mycols[2], lwd = 4, 
     ylab = "log 2 fold change", xlab = "mins after egf stimulation")
plot(as.numeric(egf.summary[110,3:9]), x = c(2, 4, 8, 16, 32, 64, 128), type = "b", pch = 19, col = mycols[2], lwd = 4, 
     ylab = "log 2 fold change", xlab = "mins after egf stimulation")
plot(as.numeric(egf.summary[141,3:9]), x = c(2, 4, 8, 16, 32, 64, 128), type = "b", pch = 19, col = mycols[2], lwd = 4, 
     ylab = "log 2 fold change", xlab = "mins after egf stimulation")

par(mfrow = c(2,2))
plot(as.numeric(egf.summary[172,3:9]), x = c(2, 4, 8, 16, 32, 64, 128), type = "b", pch = 19, col = mycols[2], lwd = 4, 
     ylab = "log 2 fold change", xlab = "mins after egf stimulation")
plot(as.numeric(egf.summary[176,3:9]), x = c(2, 4, 8, 16, 32, 64, 128), type = "b", pch = 19, col = mycols[2], lwd = 4, 
     ylab = "log 2 fold change", xlab = "mins after egf stimulation")
plot(as.numeric(egf.summary[179,3:9]), x = c(2, 4, 8, 16, 32, 64, 128), type = "b", pch = 19, col = mycols[2], lwd = 4, 
     ylab = "log 2 fold change", xlab = "mins after egf stimulation")
plot(as.numeric(egf.summary[186,3:9]), x = c(2, 4, 8, 16, 32, 64, 128), type = "b", pch = 19, col = mycols[2], lwd = 4, 
     ylab = "log 2 fold change", xlab = "mins after egf stimulation")


```
# Phospho results
# Projected PCA of movers
```{r,}
par(mfrow = c(2,1))
.pca <- plot2D(object = helaCtrl_sub[,6:10], dims = c(1,3))
addLegend(where = "topleft", object = helaCtrl_sub, ncol = 2)
highlightOnPlot(object = .pca, foi = y.filter, col = "purple3", cex = 1.5, pch = 19, labels = TRUE)
.pca <- plot2D(object = helaEgf_sub[,6:10], dims = c(1,3))
highlightOnPlot(object = .pca, foi = y.filter, col = "purple3", cex = 1.5, pch = 19, labels = TRUE)


```


```{r,}
## Some visulisation
object <- helaCtrl_sub
params <- bandleres_Itzhak5
K <- length(getMarkerClasses(object))

rownames(params[[7]]$alloc$cond11) <- names(probdir)
markerdata <- as.character(fData(markerMSnSet(object))$markers)
names(markerdata) <- rownames(markerMSnSet(object))
maxallocprob <- apply(params[[7]]$allocprob$cond11, c(1), function(x) max(colMeans(x)))
allocprob <- c(rep(1, length(markerdata)), maxallocprob)
mostprob <- apply(params[[7]]$alloc$cond11, 1, function(x) which.max(tabulate(x)))
allocorg <- getMarkerClasses(object)[mostprob]
names(allocprob) <- names(allocorg)


mcmcsampleprobs <- params[[7]]$allocprob$cond11
dimnames(mcmcsampleprobs)[[1]] <- names(mostprob)
dimnames(mcmcsampleprobs)[[3]] <- getMarkerClasses(object)

allocproborg <- apply(params[[7]]$allocprob$cond11, c(1), function(x) colMeans(x))
allocprobmarker <- matrix(0, K, length(markerdata))


for (j in seq.int(K)) {
  
  allocprobmarker[j, markerdata == getMarkerClasses(object)[j]] <- 1
  
}
colnames(allocprobmarker) <- names(markerdata)
allocproborg[,rowMeans(params[[7]]$allocOut$cond11) < 0.99] <- 0 

allocproborg <- cbind(allocproborg, allocprobmarker)
colnames(allocproborg) <- c(names(mostprob), names(markerdata))


spatial2D <- function(object,
                      allocproborg = allocproborg,
                      dims = c(1, 2),
                      cov.function = wendland.cov,
                      theta = 2,
                      derivative = 2,
                      k = 1,
                      breaks = c(0.99, 0.95, 0.9, 0.85, 0.8, 0.75),
                      aspect = 0.5) {
  
  # generate pca plot and create data from with probabilties
  .pca <- plot2D(object, dims = dims, plot = FALSE)
  probs <- data.frame(x = .pca[, 1], y = .pca[, 2], mcmc.prob = t(allocproborg)[rownames(object),])
  colnames(probs) <- c(c("x", "y"), getMarkerClasses(object))
  eigs <- colnames(.pca)
  
  # put data in appropriate long format
  probs.lst <- list()
  for(j in getMarkerClasses(object)) {
    probs.lst[[j]] <- probs[, c("x", "y", j)]
    colnames(probs.lst[[j]]) <- c("x", "y", "probability")
  }
  probs.lst.df <- plyr::ldply(probs.lst, .fun = function(x) x, .id = "organelle")
  
  # Create storage
  coords <- list()
  locations <- list()
  df <- list()
  # Create appropriate spatial grid
  for (j in getMarkerClasses(object)) {
    idxOrg <- c(probs.lst.df$organelle == j)
    coords[[j]] <- akima::interp(x = probs.lst.df$x[idxOrg],
                                 y = probs.lst.df$y[idxOrg],
                                 z = probs.lst.df$probability[idxOrg],
                                 extrap=FALSE, linear = TRUE, duplicate = TRUE) # interpolate onto appropriate grid
    coords[[j]]$z[is.na(coords[[j]]$z)] <- 0 # NaNs beyond data set to 0
    locations[[j]] <- cbind(rep(coords[[j]]$x, 40), rep(coords[[j]]$y, each = 40)) # get grid
    smoothedprobs <- fields::smooth.2d(coords[[j]]$z, x = locations[[j]],
                                       cov.function = cov.function,
                                       theta = theta,
                                       derivative = derivative, k = k) # spatial smoothing of probabilities
    # normalisation and formatting
    zmat <- matrix(smoothedprobs$z, ncol = 1)
    zmat <- zmat/max(zmat)
    df[[j]] <- data.frame(x = rep(smoothedprobs$x, 64), y = rep(smoothedprobs$y, each = 64), z = zmat)
  }
  # format data
  df.lst <- plyr::ldply(df, .fun = function(x) x, .id = "organelle") 
  df.lst <- df.lst %>%
    mutate(organelle = factor(organelle)) 
  K <- length(getMarkerClasses(object))
  cols <- getStockcol()[1:K] # get appropriate colours
  
  
  gg <- ggplot(
    data = df.lst,
    aes(x = x, y = y, z = z, color = organelle)) +
    coord_fixed() + 
    geom_contour(breaks = breaks, size = 1.2, aes(alpha = stat(level))) + 
    geom_point(alpha = 0) + 
    xlab(paste0(eigs[1])) + 
    ylab(paste0(eigs[2])) +
    scale_alpha(guide = "none") + 
    theme(legend.position = "right", 
          text = element_text(size = 12)) +
    scale_color_manual(values = cols) +
    scale_fill_manual(values = cols) +
    theme_minimal() + 
    theme(panel.grid.major = element_blank(), panel.grid.minor = element_blank(),
          aspect.ratio = aspect,
          panel.border = element_rect(colour = "black", fill = NA, size = 1),
          plot.title = element_text(hjust = 0.5, size = 20),
          legend.text=element_text(size = 14)) +
    ggtitle(label = "Spatial variation of localisation probabilities") 
  gg
  return(gg)
}
```

```{r,}
mcmcsampleprobs <- params[[7]]$allocprob$cond11
dimnames(mcmcsampleprobs)[[1]] <- names(mostprob)
dimnames(mcmcsampleprobs)[[3]] <- getMarkerClasses(object)

allocproborg <- apply(params[[7]]$allocprob$cond11, c(1), function(x) colMeans(x))
allocprobmarker <- matrix(0, K, length(markerdata))


for (j in seq.int(K)) {
  
  allocprobmarker[j, markerdata == getMarkerClasses(object)[j]] <- 1
  
}
colnames(allocprobmarker) <- names(markerdata)
allocproborg[,rowMeans(params[[7]]$allocOut$cond11) < 0.99] <- 0 

allocproborg <- cbind(allocproborg, allocprobmarker)
colnames(allocproborg) <- c(names(mostprob), names(markerdata))


.pca <- plot2D(helaCtrl_sub[,6:10], dims = c(1,3), plot = FALSE)
gg <- spatial2D(object = helaCtrl_sub[,6:10], allocproborg = allocproborg, dims = c(1,3))
gg <- gg + geom_point(data = data.frame(x = .pca[y.filter[c(1,4,7)],1], y = .pca[y.filter[c(1,4,7)],2]),
                   aes(x = x, y = y, z = 1), colour = "black", size = 3, shape = 18) + labs(title = "HeLa Control") + 
  annotate(geom="text", x = .pca[y.filter[c(1,4,7)],1] + 0.5, y = .pca[y.filter[c(1,4,7)],2], label= c(y.filter[c(1,4,7)]),
              color="purple3")

mcmcsampleprobs <- params[[7]]$allocprob$cond21
dimnames(mcmcsampleprobs)[[1]] <- names(mostprob)
dimnames(mcmcsampleprobs)[[3]] <- getMarkerClasses(object)

allocproborg <- apply(params[[7]]$allocprob$cond21, c(1), function(x) colMeans(x))
allocprobmarker <- matrix(0, K, length(markerdata))


for (j in seq.int(K)) {
  
  allocprobmarker[j, markerdata == getMarkerClasses(object)[j]] <- 1
  
}
colnames(allocprobmarker) <- names(markerdata)
allocproborg[,rowMeans(params[[7]]$allocOut$cond21) < 0.99] <- 0 

allocproborg <- cbind(allocproborg, allocprobmarker)
colnames(allocproborg) <- c(names(mostprob), names(markerdata))


.pca <- plot2D(helaEgf_sub[,6:10], dims = c(1,3), plot = FALSE)
gg2 <- spatial2D(object = helaEgf_sub[,6:10],allocproborg = allocproborg, dims = c(1,3))
gg2 <- gg2 + geom_point(data = data.frame(x = .pca[y.filter[c(1,4,7)],1], y = .pca[y.filter[c(1,4,7)],2]),
                   aes(x = x, y = y, z = 1), colour = "black", size = 3, shape = 18)  + labs(title = "HeLa EGF Stimulated") + 
  annotate(geom="text", x = .pca[y.filter[c(1,4,7)],1] + 0.5, y = .pca[y.filter[c(1,4,7)],2] + c(0,0,-0.5), label= c(y.filter[c(1,4,7)]),
              color="purple3")
gg/gg2
```


```{r,}
mcmcsampleprobs2 <- params[[7]]$allocprob$cond21
dimnames(mcmcsampleprobs2)[[1]] <- names(mostprob)
dimnames(mcmcsampleprobs2)[[3]] <- getMarkerClasses(object)
par(mfrow = c(1,2))
mcmc_plot_probs(param = mcmcsampleprobs, fname = y.filter[2], bw = 0.3)
mcmc_plot_probs(param = mcmcsampleprobs2, fname = y.filter[2], bw = 0.3)
```

```{r,}
plot2D(helaCtrl_sub[,1:5], main = "HeLa Control", dims = c(1,3), cex.lab = 1.5, cex.main = 1.5, grid = FALSE)
addLegend(helaCtrl_sub, cex = 1.2, where = "topleft")

```
```{r,}
plot2D(helaEgf_sub[,1:5], main = "HeLa EGF", dims = c(1,3), cex.lab = 1.5, cex.main = 1.5, grid = FALSE)
addLegend(helaEgf_sub, cex = 1.2, where = "topleft")

```
````
