## Supplementary Code 1 for "Inferring differential subcellular localisation in comparative spatial proteomics using BANDLE": methodtesting2.R

```
## method Testing 2, using K-NN + resdiual bootstrapping to create replicates

# Make required data
require(pRoloc)
require(pRolocdata)
data("tan2009r1")

## Perform K-NN with K = 10 classification to get clustering
tan2009r1 <- knnClassification(object = tan2009r1, k = 10)
plot2D(tan2009r1, fcol = "knn")
fcol <- "markers"

## Compute means of each organelle just using markers
meanOrganelle <- function(mydata, fcol = "markers"){
  
  M <- V <- matrix(NA, nrow = length(getMarkerClasses(mydata, fcol = fcol)), ncol = ncol(mydata))
  rownames(M) <- rownames(V) <-  getMarkerClasses(mydata, fcol = fcol)
  for (j in getMarkerClasses(mydata, fcol = fcol)) {
    M[j, ] <- colMeans(exprs(mydata)[fData(mydata)[, fcol] == j,])
    V[j, ] <- apply(exprs(mydata)[fData(mydata)[, fcol] == j,], 2, var)
  }

  return(list(M = M, V = V))
}

orgM <- meanOrganelle(mydata = tan2009r1)$M


## Use means as fitted function for regression and compute residuals
residual_tan <- matrix(NA, nrow = nrow(tan2009r1), ncol = ncol(tan2009r1))
residual_tan <- exprs(tan2009r1) - orgM[fData(tan2009r1)$knn, ]

## Bootstrap residuals
numRep <- 6 # number of additional datasets

tan2009rep <- list()
K <- length(getMarkerClasses(tan2009r1, fcol = fcol))
#nu <- 1
#nu <- rep(2.5, length(fData(tan2009r1)$knn))
nu <- runif(K, 1,2)[fData(tan2009r1)$knn] # cluster specific residual inflated noise

for (i in seq.int(numRep)) {
  
  idxBoot <- matrix(sample.int(4, size = 4 * nrow(tan2009r1), replace = TRUE), nrow = nrow(tan2009r1), ncol(tan2009r1))
  bootres <- matrix(NA,  nrow = nrow(tan2009r1), ncol(tan2009r1))
  for (k in seq.int(nrow(tan2009r1))) {
    bootres[k, ] <- residual_tan[k, idxBoot[k, ]]
  }
  tan2009rep[[i]] <- tan2009r1
  newExprs <- orgM[fData(tan2009r1)$knn, ] + nu * bootres ## add boostraps
  colnames(newExprs) <- colnames(tan2009r1)
  rownames(newExprs) <- rownames(tan2009r1)
  exprs(tan2009rep[[i]]) <- newExprs
  
}

par(mfrow = c(2,3))
lapply(tan2009rep, plot2D)
plot2D(tan2009r1)

par(mfrow = c(1,1))

## simulate translocation events
mtan2009 <- unknownMSnSet(tan2009r1)
perm1 <- sample.int(nrow(mtan2009), size = 20, replace = TRUE)
perm1_names <- rownames(mtan2009)[perm1]

replicateStats <- lapply(tan2009rep, FUN = meanOrganelle)

# Simulate dynamics # gahh need to use correct names

for (i in perm1_names) {
    K <- length(getMarkerClasses(tan2009r1))
    r <- sample.int(K, size = 1)
  for (j in 4:6) {
    exprs(tan2009rep[[j]])[i, ] <- replicateStats[[j]]$M[r, ] + rnorm(ncol(tan2009r1), mean = 0, sd = sqrt(replicateStats[[j]]$V[r, ]))
  }
}


# exprs(tan2009rep[[4]]) <- t(apply(exprs(tan2009rep[[4]]), 1, function(x) x + c(0.3,0,0,0)))
# exprs(tan2009rep[[5]]) <- t(apply(exprs(tan2009rep[[5]]), 1, function(x) x + c(0,0.3,0,0)))
# exprs(tan2009rep[[6]]) <- t(apply(exprs(tan2009rep[[6]]), 1, function(x) x + c(0,0,0.3,0)))
#tan2009rep[[4]] <- normalise(object = tan2009rep[[4]], method = "sum")
#tan2009rep[[5]] <- normalise(object = tan2009rep[[5]], method = "sum")
#tan2009rep[[6]] <- normalise(object = tan2009rep[[6]], method = "sum")
par(mfrow = c(1,3))
plot2D(tan2009rep[[4]])
#fraction permuting
exprs(tan2009rep[[4]]) <- exprs(tan2009rep[[4]])[,c(2,1,3,4)]
exprs(tan2009rep[[5]]) <- exprs(tan2009rep[[5]])[,c(1,2,4,3)]
exprs(tan2009rep[[6]]) <- exprs(tan2009rep[[6]])[,c(3,2,1,4)]


  
par(mfrow = c(1,2))
plot2D(object = tan2009rep[[4]][,c(2,1,3,4)])
highlightOnPlot(object = tan2009rep[[4]], foi = perm1_names, TRUE)
plot2D(object = tan2009r1)
highlightOnPlot(object = tan2009r1, foi = perm1_names, labels = TRUE)

mrres <- mrMethod(c(tan2009rep))

resdynpg <- diffLoc(object_cond1 = c(tan2009rep[1:3]), object_cond2 = tan2009rep[4:6], numIter = 5000, burnin = 1000, pg = TRUE)
resdyndir <- diffLoc(object_cond1 = c(tan2009rep[1:3]), object_cond2 = tan2009rep[4:6], numIter = 5000, burnin = 1000, pg = FALSE)


probsameorganelle <- function(object , params) {
  
  res <- vector(mode = "numeric", length = nrow(unknownMSnSet(object)))
  for (i in 1:nrow(unknownMSnSet(object))) {
    res[i] <- sum(params$alloc[[1]][i, ] == params$alloc[[2]][i, ])
  }
  names(res) <- rownames(unknownMSnSet(object))
  
  return(res)
}

probpg <- probsameorganelle(object = tan2009r1, params = resdynpg)
probdir <- probsameorganelle(object = tan2009r1, params = resdyndir)

require(pROC)
outcome <- matrix(0, ncol = 1, nrow = length(probpg))
names(outcome) <- names(probpg)
outcome[perm1_names] <- 1
prediction <- probpg/ncol(resdynpg$allocOut[[1]])
par(mfrow = c(1,1))

ROC_bayes <- roc(outcome, 1 - prediction)
ROC_mr <- roc(outcome, mrres$MRscore[names(outcome)], direction = "<")
plot(ROC_bayes, type = "b", col = "darkgreen", lwd = 1, pch = 19, asp = NA, legacy.axes = T)
lines(ROC_mr, type = "b", col = "blue", lwd = 1, pch = 19, asp = NA)

outcome2 <- matrix(0, ncol = 1, nrow = length(probdir))
names(outcome2) <- names(probdir)
outcome2[perm1_names] <- 1
prediction2 <- probdir/ncol(resdyndir$allocOut[[1]])
ROC_dir <- roc(outcome2, 1 - prediction2)
plot(ROC_dir,type = "b", col = "orange", lwd = 1, pch = 19, asp = NA)
legend("bottomright", c("MR", "Dirichlet", "Polya-Gamma"), col = c("blue", "orange", "darkgreen"), pch = 19)


# Dirichlet
y <- 1 - probdir[order(probdir)]/ncol(resdyndir$allocOut[[1]])
plot(y[1:20], col = outcome[order(probdir)] + 2, pch = 19, ylim = c(0.5,1), ylab = "Differential Localisation Probability", xlab = "Rank")

#Polya-Gamma
pgy <- 1 - probpg[order(probpg)]/ncol(resdynpg$allocOut[[1]])
points(pgy, col = outcome[order(probpg)] + 2, pch = 24)
abline(h = 0.99, col = "blue", lwd = 2)
abline(h = 0.95, col = "purple", lwd = 2)


sum(names(probdir[order(probdir)])[(1 - probdir[order(probdir)]/ncol(resdyndir$allocOut[[1]])) > 0.999] %in% perm1_names)
sum(names(probpg[order(probpg)])[(1 - probpg[order(probpg)]/ncol(resdynpg$allocOut[[1]])) > 0.999] %in% perm1_names)
names(mrres$Mscore[(mrres$Mscore > 2) & mrres$Rscore > 0.9 ] ) %in% perm1_names


plot(mrres$Mscore, mrres$Rscore, col = outcome[names(mrres$Mscore)] + 2, pch = 19, ylim = c(-1,1))
abline(h = 0.9, col = "blue", lwd = 2)
abline(v = 2, col = "purple", lwd = 2)


par(mfrow = c(1,2))
plot2D(object = tan2009r1)
highlightOnPlot(object = tan2009r1, foi = "Q9VFV9", labels = TRUE)
plot2D(object = tan2009rep[[4]])
highlightOnPlot(object = tan2009rep[[4]], foi = "Q9VFV9", labels = TRUE)


# prior predictive checks for dirichlet prior

priornotAlloc <- vector(length = 5000)
nkknown <- table(getMarkers(tan2009r1, verbose = FALSE))[getMarkerClasses(tan2009r1)]
for (i in 1:5000) {
  
  concentration <- diag(rep(1, K)) + diag(nkknown) + matrix(0.05, nrow = K, ncol = K)
  currentweights <- t(sampleDirichlet(K^2, c(concentration)))
  
  priornotAlloc[i] <- sum(currentweights[1, -c(12 * seq.int(1:11) - 11)])
}
hist(priornotAlloc)
# average number of differential localisations
mean(priornotAlloc) * nrow(unknownMSnSet(tan2009r1))

# probability of having greater than 15 
sum((priornotAlloc * nrow(unknownMSnSet(tan2009r1))) > 15)/5000


# prior predictive checks for Polya-Gamma prior


# expressions from data and important summaries
exprs_tan <- exprs(tan2009r1)
M <- colMeans(exprs_tan)
V <- cov(exprs_tan)

tau <- 0.2 # 0.3 works well
  
sigma1 <- covOrganelle(mydata = tan2009rep[[1]])
#sigma1 <- tau * (sigma1)
#sigma1[sigma1 == 0] <- -tau
sigma2 <- covOrganelle(mydata = tan2009rep[[4]])
#sigma2 <- tau * (sigma2)
#sigma2[sigma2 == 0] <- - tau
w <- rep(1, K^2)

n_vec <- c(diag(nkknown))
n_cum <- sum(n_vec) - c(0, cumsum(n_vec[1:(K^2 - 1)]))
kappa <- n_vec - 1/2
#sigma_post <- solve(diag(w) + solve(diag(rep(0.01, K^2)))) # inefficient return later
sigma_post <- solve(diag(w) + tau * kronecker(solve(t(sigma1) + diag(0.01, K)), solve(t(sigma2) + diag(0.01, K))))
heatmap(sigma_post)


mu_prior <- rep(-9, K^2)
mu_prior[c(12 * seq.int(1:11) - 11)] <- mu_prior[c(12 * seq.int(1:11) - 11)] + 1

mu_post <- sigma_post %*% (kappa + tau * kronecker(solve(t(sigma1) + diag(0.01, K)), solve(t(sigma2) + diag(0.01, K))) %*% mu_prior)
#mu_post <- sigma_post %*% (kappa + tau * kronecker(solve(t(sigma1) + diag(0.01, K)), solve(t(sigma2) + diag(0.01, K))) %*% mu_prior)

pg_res <- prior_pred_pg(object = tan2009rep, tau = tau, lambda = 0.01, iter = 10000, q = 15)
plot(currentweights)
hist(pg_res$priornotAllocpg)
# average number of differential localisations
mean(pg_res$priornotAllocpg) * nrow(unknownMSnSet(tan2009r1))
var(pg_res$priornotAllocpg)

# probability of having greater than 15 
sum((pg_res$priornotAllocpg * nrow(unknownMSnSet(tan2009r1))) > 15)/10000


## Bootstrap Probabilities

bootstrapdiffloc <- function(object, params, top = 20, Bootsample = 5000, decreasing = FALSE) {
  
  res <- matrix(NA, ncol = Bootsample, nrow = top)
  probs <- probsameorganelle(object = object, params = params)
  
  prBootnames <- names(probs[order(probs, decreasing = decreasing)][seq.int(top)])
  rownames(params$alloc[[1]]) <- rownames(params$alloc[[2]]) <- rownames(unknownMSnSet(object))
  
  rownames(res) <- prBootnames
  for (t in seq.int(Bootsample)){
    bootidx <- sample.int(ncol(params$alloc[[1]]), replace = TRUE)
    for (i in prBootnames) {
      res[i,t] <- sum(params$alloc[[1]][i, bootidx] == params$alloc[[2]][i, bootidx])
    }
  }
  
  return(res)
}
bootresdir <- bootstrapdiffloc(object = tan2009r1, params = resdyndir, top = 50, decreasing = FALSE)
bootrespg <- bootstrapdiffloc(object = tan2009r1, params = resdynpg, top = 30)

par(mfrow = c(1,1))
boxplot(t(1 - bootresdir/800)[,1:50], las = 2, col = 1 * (rownames(bootresdir) %in% perm1_names) + 2)
boxplot(t(1 - bootrespg/800), las = 2, col = 1 * (rownames(bootrespg) %in% perm1_names) + 2)
abline(h = 0.995, col = "blue", lwd = 2)

## Compute KL-divergence

kldirpg <- function(sigma, mu, alpha) {
  
  D <- length(mu)
  entpg <- -log((2*pi*exp(1))^D)/2 - sum(log(eigen(sigma)$values))/2 # compute determinant from eigenvalues
  entdir <- exp(sum(lgamma(alpha)) - lgamma(sum(alpha)))
  entdirpg_1 <- c(alpha) * (log(1 + exp(-mu)) + exp(mu) * diag(sigma)/(1 + exp(mu))^2)
  entdirpg_2 <- c(alpha) * (log(1 + exp(mu)) - exp(mu) * diag(sigma)/(1 + exp(mu))^2)
  entdirpgsum_1 <- sum(entdirpg_1)[-D]
  entdirpgsum_2 <- sum(seq.int(1, D - 1) * entdirpg_2[2:D])
  
  res <- entpg + entdir + entdirpgsum_1 + entdirpgsum_2
  
  return(res)
}

kldirpg(sigma = sigma_post, mu = mu_post, alpha = concentration)

nkknown <- table(getMarkers(tan2009r1, verbose = FALSE))[getMarkerClasses(tan2009r1)]
concentration <- diag(rep(1, K)) + diag(nkknown) + matrix(0.01, nrow = K, ncol = K)

klres <- matrix(NA, nrow = 10, ncol = 14)
for (j in seq.int(1:10)) {
  for (i in seq.int(14)) {
    
    x <- c(0.01, 0.02, 0.05,0.06, 0.07, 0.1, 0.2, 0.3, 0.4, 0.5, 1, 2, 3, 10)
    # expressions from data and important summaries
    exprs_tan <- exprs(tan2009r1)
    M <- colMeans(exprs_tan)
    V <- cov(exprs_tan)
    
    tau <- x[i]
    
    sigma1 <- covOrganelle(mydata = tan2009rep[[1]])
    #sigma1 <- tau * (sigma1)
    #sigma1[sigma1 == 0] <- -tau
    sigma2 <- covOrganelle(mydata = tan2009rep[[4]])
    #sigma2 <- tau * (sigma2)
    #sigma2[sigma2 == 0] <- - tau
    w <- rep(1, K^2)
    
    n_vec <- c(diag(nkknown))
    n_cum <- sum(n_vec) - c(0, cumsum(n_vec[1:(K^2 - 1)]))
    kappa <- n_vec - 1/2
    #sigma_post <- solve(diag(w) + solve(diag(rep(0.01, K^2)))) # inefficient return later
    sigma_post <- solve(diag(w) + tau * kronecker(solve(t(sigma1) + diag(0.01, K)), solve(t(sigma2) + diag(0.01, K))))
    #heatmap(sigma_post)
    
    
    mu_prior <- rep(-j, K^2)
    mu_prior[c(12 * seq.int(1:11) - 11)] <- mu_prior[c(12 * seq.int(1:11) - 11)] + 1
    
    mu_post <- sigma_post %*% (kappa + tau * kronecker(solve(t(sigma1) + diag(0.01, K)), solve(t(sigma2) + diag(0.01, K))) %*% mu_prior)
    #mu_post <- sigma_post %*% (kappa + tau * kronecker(solve(t(sigma1) + diag(0.01, K)), solve(t(sigma2) + diag(0.01, K))) %*% mu_prior)
  
    klres[j, i] <- kldirpg(sigma_post, mu_post, concentration)
  }
}
rownames(klres) <- c(-seq.int(1:10))
colnames(klres) <- x


## The following plot show how are far away different Polya-Gamma priors are from a fixed chosen Dirichlet prior. That is
## assumed to be well calibrated - we observe most models sit around 50 normalised KL. We might choose to disregard
## models that are far away form the Dirichlet model. The normalisation of KL divergence corresponds to its relationship
## with the fisher information. See Simpson et al. 2017 section 4.1 Though the link is not explicit because the models
## are not parameterised by the same parameters.
matplot(log10(x), t(sqrt(2 * klres)), type = "b", lwd = 4, col = getStockcol()[as.factor(rownames(klres))], pch = 19, lty = 1,
        main = "standardised KL divergence between induced priors" ,ylab = "Normalised KL", xlab = "log of tau")
legend(x = 0.7, y = 270, legend = as.factor(rownames(klres)), fill = getStockcol()[as.factor(rownames(klres))])


## How far away are my priors? The plot shows given a particular normalised KL divergence what is the probability that another
## prior has lower distance. That is given we believe we have a well calibrated prior, how surprising are these other priors?
## We assume and exponenial distribution on the normalised KL divergence, which assumes the relative distance between the models
## doesn't depend on the distance itself, it also posits that deviations from
## this assume different decay rates in different part of the parameter space, which maybe appropriate for some priors

klprobs <- pexp(c(0, 50, 100, 150, 200, 250), rate = 1/50, lower.tail = TRUE)
plot(c(0, 50, 100, 150, 200, 250), klprobs, xlab = "normalised KL distance", ylab = "probability of model with lower distance",
     type = "b", lwd = 2, col = getStockcol()[2], pch = 19, lty = 1)


## optim mu is gives around 50
optimres <- optim(par = rep(-6, 121), fn = kldirpg, sigma = sigma_post, alpha = concentration)


## epsilon local circular sensitivity
dir_prior_jef <- matrix(0.5, K, K)
dir_prior_sym <- matrix(0.501, K, K)
dir_prior <- diag(rep(1, K)) + matrix(0.05, K, K)
dirpriorcheck <- prior_pred_dir(tan2009r1, dir_prior = dir_prior_jef, iter = 800)
resdyndir <- diffLoc(object_cond1 = c(tan2009rep[1:3]), object_cond2 = tan2009rep[4:6], numIter = 5000, burnin = 1000,
                     pg = FALSE, dir_prior = dir_prior_jef)

neqmargpost <- density(colSums(resdyndir$alloc$cond11 != resdyndir$alloc$cond21)/ncol(resdyndir$allocOut[[1]]), from = 0, to = 0.3, bw = 0.01)
newmargprior <- density(dirpriorcheck$priornotAlloc, from = 0, to = 0.3, bw = 0.01)

plot(neqmargpost$y)
points(newmargprior$y, col = "blue")

## Compute "importance ratio"
impratio <- neqmargpost$y/newmargprior$y

alpha <- seq(0.05, 10, by = 0.05)
kldireps <- vector(mode = "numeric", length = length(alpha))
for (i in seq.int(length(alpha))) {
  dir_prior_sym <- matrix(alpha[i], K, K)
  kldireps[i] <- kldir(alpha = c(dir_prior_sym), beta = c(dir_prior_jef))
}
plot(alpha, kldireps)

dirpriorcheck <- prior_pred_dir(tan2009r1, dir_prior = dir_prior_jef, iter = 800)

dirsymnotalloc <- matrix(NA, nrow = length(alpha) , ncol = 800)
for (i in seq.int(length(alpha))){
  dir_prior_sym <- matrix(alpha[i], K, K)
  dirsymnotalloc[i, ] <- prior_pred_dir(tan2009r1, dir_prior = dir_prior_sym, iter = 800)$priornotAlloc
}

sens_res <- apply(dirsymnotalloc,1, function(x) density(x, from = 0, to = 0.3)$y *impratio)

plot(kldireps, colMeans(sens_res))


## Some visulisation
rownames(resdyndir$alloc$cond11) <- names(probdir)
markerdata <- as.character(fData(markerMSnSet(tan2009r1))$markers)
names(markerdata) <- rownames(markerMSnSet(tan2009r1))
maxallocprob <- apply(resdyndir$allocprob$cond11, c(1), function(x) max(colMeans(x)))
allocprob <- c(rep(1, length(markerdata)), maxallocprob)
mostprob <- apply(resdyndir$alloc$cond11, 1, function(x) which.max(tabulate(x)))
allocorg <- getMarkerClasses(tan2009r1)[mostprob]
names(allocprob) <- names(allocorg)


mcmcsampleprobs <- resdyndir$allocprob$cond11
dimnames(mcmcsampleprobs)[[1]] <- names(mostprob)
dimnames(mcmcsampleprobs)[[3]] <- getMarkerClasses(tan2009r1)

allocproborg <- apply(resdyndir$allocprob$cond1, c(1), function(x) colMeans(x))
allocprobmarker <- matrix(0, K, length(markerdata))

mcmc_plot_probs(param = mcmcsampleprobs, fname = "P50461")

for (j in seq.int(K)) {
  
  allocprobmarker[j, markerdata == getMarkerClasses(tan2009r1)[j]] <- 1
  
}
colnames(allocprobmarker) <- names(markerdata)
allocproborg[,rowMeans(resdyndir$allocOut$cond11) < 0.99] <- 0 

allocproborg <- cbind(allocproborg, allocprobmarker)
colnames(allocproborg) <- c(names(mostprob), names(markerdata))


spatial2D <- function(object,
                      dims = c(1, 2),
                      cov.function = wendland.cov,
                      theta = 2,
                      derivative = 2,
                      k = 1,
                      breaks = c(0.99, 0.95, 0.9, 0.85, 0.8, 0.75, 0.7),
                      aspect = 0.5) {
  
  # generate pca plot and create data from with probabilties
  .pca <- plot2D(object, dims = dims, plot = FALSE)
  probs <- data.frame(x = .pca[, 1], y = .pca[, 2], mcmc.prob = t(allocproborg)[rownames(object),])
  colnames(probs) <- c(c("x", "y"), getMarkerClasses(object))
  eigs <- colnames(.pca)
  
  # put data in appropriate long format
  probs.lst <- list()
  for(j in getMarkerClasses(object)) {
    probs.lst[[j]] <- probs[, c("x", "y", j)]
    colnames(probs.lst[[j]]) <- c("x", "y", "probability")
  }
  probs.lst.df <- plyr::ldply(probs.lst, .fun = function(x) x, .id = "organelle")
  
  # Create storage
  coords <- list()
  locations <- list()
  df <- list()
  # Create appropriate spatial grid
  for (j in getMarkerClasses(object)) {
    idxOrg <- c(probs.lst.df$organelle == j)
    coords[[j]] <- akima::interp(x = probs.lst.df$x[idxOrg],
                                 y = probs.lst.df$y[idxOrg],
                                 z = probs.lst.df$probability[idxOrg],
                                 extrap=FALSE, linear = TRUE, duplicate = TRUE) # interpolate onto appropriate grid
    coords[[j]]$z[is.na(coords[[j]]$z)] <- 0 # NaNs beyond data set to 0
    locations[[j]] <- cbind(rep(coords[[j]]$x, 40), rep(coords[[j]]$y, each = 40)) # get grid
    smoothedprobs <- fields::smooth.2d(coords[[j]]$z, x = locations[[j]],
                                       cov.function = cov.function,
                                       theta = theta,
                                       derivative = derivative, k = k) # spatial smoothing of probabilities
    # normalisation and formatting
    zmat <- matrix(smoothedprobs$z, ncol = 1)
    zmat <- zmat/max(zmat)
    df[[j]] <- data.frame(x = rep(smoothedprobs$x, 64), y = rep(smoothedprobs$y, each = 64), z = zmat)
  }
  # format data
  df.lst <- plyr::ldply(df, .fun = function(x) x, .id = "organelle") 
  df.lst <- df.lst %>%
    mutate(organelle = factor(organelle)) 
  K <- length(getMarkerClasses(object))
  cols <- getStockcol()[1:K] # get appropriate colours
  
  
  gg <- ggplot(
    data = df.lst,
    aes(x = x, y = y, z = z, color = organelle)) +
    coord_fixed() + 
    geom_contour(breaks = breaks, size = 1.2, aes(alpha = stat(level))) + 
    geom_point(alpha = 0) + 
    xlab(paste0(eigs[1])) + 
    ylab(paste0(eigs[2])) +
    scale_alpha(guide = "none") + 
    theme(legend.position = "right", 
          text = element_text(size = 12)) +
    scale_color_manual(values = cols) +
    scale_fill_manual(values = cols) +
    theme_minimal() + 
    theme(panel.grid.major = element_blank(), panel.grid.minor = element_blank(),
          aspect.ratio = aspect,
          panel.border = element_rect(colour = "black", fill = NA, size = 1),
          plot.title = element_text(hjust = 0.5, size = 20),
          legend.text=element_text(size = 14)) +
    ggtitle(label = "Spatial variation of localisation probabilities") 
  gg
  return(gg)
}
```
