## Supplementary Code 1 for "Inferring differential subcellular localisation in comparative spatial proteomics using BANDLE": methodtesting3.R

```
## Testing 
require(pROC)
ROCresBayes <- list()
ROCresDir <- list()
ROCresmr <- list()
ROCrespg1 <- list()
probpgres <- list()
probpgres1 <- list()
probdirres <- list()
mrreslist <- list()
perm1_nameslist <- list()

for (l in seq.int(10)) {
  # Make required data
  data("tan2009r1")
  
  ## Perform K-NN with K = 10 classification to get clustering
  tan2009r1 <- knnClassification(object = tan2009r1, k = 10)
  plot2D(tan2009r1, fcol = "knn")
  fcol <- "markers"
  
  ## Compute means of each organelle just using markers
  meanOrganelle <- function(mydata, fcol = "markers"){
    
    M <- V <- matrix(NA, nrow = length(getMarkerClasses(mydata, fcol = fcol)), ncol = ncol(mydata))
    rownames(M) <- rownames(V) <-  getMarkerClasses(mydata, fcol = fcol)
    for (j in getMarkerClasses(mydata, fcol = fcol)) {
      M[j, ] <- colMeans(exprs(mydata)[fData(mydata)[, fcol] == j,])
      V[j, ] <- apply(exprs(mydata)[fData(mydata)[, fcol] == j,], 2, var)
    }
    
    return(list(M = M, V = V))
  }
  
  orgM <- meanOrganelle(mydata = tan2009r1)$M
  
  
  ## Use means as fitted function for regression and compute residuals
  residual_tan <- matrix(NA, nrow = nrow(tan2009r1), ncol = ncol(tan2009r1))
  residual_tan <- exprs(tan2009r1) - orgM[fData(tan2009r1)$knn, ]
  
  ## Bootstrap residuals
  numRep <- 6 # number of additional datasets
  
  tan2009rep <- list()
  K <- length(getMarkerClasses(tan2009r1, fcol = fcol))
  #nu <- 1
  #nu <- rep(2.5, length(fData(tan2009r1)$knn))
  nu <- runif(K, 1, 2)[fData(tan2009r1)$knn] # cluster specific residual inflated noise
  
  for (i in seq.int(numRep)) {
    
    idxBoot <- matrix(sample.int(4, size = 4 * nrow(tan2009r1), replace = TRUE), nrow = nrow(tan2009r1), ncol(tan2009r1))
    bootres <- matrix(NA,  nrow = nrow(tan2009r1), ncol(tan2009r1))
    for (k in seq.int(nrow(tan2009r1))) {
      bootres[k, ] <- residual_tan[k, idxBoot[k, ]]
    }
    tan2009rep[[i]] <- tan2009r1
    newExprs <- orgM[fData(tan2009r1)$knn, ] + nu * bootres ## add boostraps
    colnames(newExprs) <- colnames(tan2009r1)
    rownames(newExprs) <- rownames(tan2009r1)
    exprs(tan2009rep[[i]]) <- newExprs
    
  }
  
  par(mfrow = c(2,3))
  lapply(tan2009rep, plot2D)
  plot2D(tan2009r1)
  
  par(mfrow = c(1,1))
  
  ## simulate translocation events
  mtan2009 <- unknownMSnSet(tan2009r1)
  perm1 <- sample.int(nrow(mtan2009), size = 20, replace = TRUE)
  perm1_names <- rownames(mtan2009)[perm1]
  
  replicateStats <- lapply(tan2009rep, FUN = meanOrganelle)
  
  # Simulate dynamics # gahh need to use correct names
  
  for (i in perm1_names) {
    K <- length(getMarkerClasses(tan2009r1))
    r <- sample.int(K, size = 1)
    for (j in 4:6) {
      exprs(tan2009rep[[j]])[i, ] <- replicateStats[[j]]$M[r, ] + rnorm(ncol(tan2009r1), mean = 0, sd = sqrt(replicateStats[[j]]$V[r, ]))
    }
  }
  
  # computational methods
  mrres <- mrMethod(c(tan2009rep))
  par(mfrow = c(3,4))
  resdynpg <- diffLoc(object_cond1 = c(tan2009rep[1:3]), object_cond2 = tan2009rep[4:6], numIter = 5000, burnin = 1000,
                      pg = TRUE, hypLearn = "LBFGS", tau = 0.2)
  par(mfrow = c(3,4))
  resdynpg1 <- diffLoc(object_cond1 = c(tan2009rep[1:3]), object_cond2 = tan2009rep[4:6], numIter = 5000, burnin = 1000,
                      pg = TRUE, hypLearn = "LBFGS", tau = 1)
  par(mfrow = c(3,4))
  resdyndir <- diffLoc(object_cond1 = c(tan2009rep[1:3]), object_cond2 = c(tan2009rep[4:6]),
                       numIter = 5000, burnin = 1000, pg = FALSE, hypLearn = "LFBGS")
  
  probpg <- probsameorganelle(object = tan2009r1, params = resdynpg)
  probpg1 <- probsameorganelle(object = tan2009r1, params = resdynpg1)
  probdir <- probsameorganelle(object = tan2009r1, params = resdyndir)
  
  outcome <- matrix(0, ncol = 1, nrow = length(probpg))
  names(outcome) <- names(probpg)
  outcome[perm1_names] <- 1
  prediction <- probpg/ncol(resdynpg$allocOut[[1]])
  par(mfrow = c(1,1))
  
  ROC_bayes <- roc(outcome, 1 - prediction)
  ROC_mr <- roc(outcome, mrres$MRscore[names(outcome)], direction = "<")
  plot(ROC_bayes, type = "b", col = "darkgreen", lwd = 1, pch = 19, asp = NA, legacy.axes = T)
  lines(ROC_mr, type = "b", col = "blue", lwd = 1, pch = 19, asp = NA)
  
  outcome2 <- matrix(0, ncol = 1, nrow = length(probdir))
  names(outcome2) <- names(probdir)
  outcome2[perm1_names] <- 1
  prediction2 <- probdir/ncol(resdyndir$allocOut[[1]])
  ROC_dir <- roc(outcome2, 1 - prediction2)
  lines(ROC_dir,type = "b", col = "orange", lwd = 1, pch = 19, asp = NA)

  outcome3 <- matrix(0, ncol = 1, nrow = length(probpg1))
  names(outcome3) <- names(probpg1)
  outcome3[perm1_names] <- 1
  prediction3 <- probpg1/ncol(resdynpg1$allocOut[[1]])
  ROC_pg1 <- roc(outcome3, 1 - prediction3)
  lines(ROC_pg1,type = "b", col = "purple", lwd = 1, pch = 19, asp = NA)
  legend("bottomright", c("MR", "Dirichlet", "Polya-Gamma", "Polya-Gamma unshrunk"), col = c("blue", "orange", "darkgreen", "purple"), pch = 19)
  
  
  ROCresBayes[[l]] <- ROC_bayes
  ROCresDir[[l]] <- ROC_dir
  ROCresmr[[l]] <- ROC_mr
  ROCrespg1[[l]] <- ROC_pg1
  probpgres[[l]] <- probpg
  probdirres[[l]] <- probdir
  probpgres1[[l]] <- probpg1
  mrreslist[[l]] <- mrres
  perm1_nameslist[[l]] <- perm1_names

}

auc <- matrix(unlist(lapply(c(ROCresBayes, ROCresDir, ROCresmr, ROCrespg1), function(x)x$auc)), ncol = 4, byrow = FALSE)
colnames(auc) <- c("PG", "Dir", "MR", "PG unshrunk")
boxplot(auc[,c(3,2,1,4)], ylim = c(0.5,1), col = c("blue", "orange", "darkgreen", "darkorchid4"))

auc_long <- as.data.frame(melt(auc[,c(3,2,1,4)]))
colnames(auc_long) <- c("Iteration", "Method", "AUC")

gg <- ggplot(data = auc_long, aes(x = Method, y = AUC, fill = Method)) + geom_boxplot(outlier.colour="black", outlier.shape=16,
                                                                       outlier.size=2, notch=FALSE)
gg <- gg + theme_classic() + scale_fill_manual(values=c("blue", "orange", "darkgreen", "darkorchid4"))
gg <- gg + theme(legend.position="none") + scale_y_continuous(limits = c(0.5,1))
gg <- gg + ggtitle("Area under curve for different methods across multiple random datasets") + xlab("Method") + ylab("AUC")
gg <- gg + scale_x_discrete(labels = c("MR", "Dirichlet", "Polya-Gamma", "Polya-Gamma Unshrunk"))
gg


plot(ROC_bayes, type = "b", col = "darkgreen", lwd = 1.5, pch = 19, asp = NA, legacy.axes = T, 
     xlab = "False Positive Rate",
     ylab = "True Positive Rate",
     main = "ROC curve demonstrating proposed methods outperform current approach")
lines(ROC_mr, type = "b", col = "blue", lwd = 1.5, pch = 19, asp = NA)
lines(ROC_dir,type = "b", col = "orange", lwd = 1.5, pch = 19, asp = NA)
lines(ROC_pg1,type = "b", col = "darkorchid", lwd = 1.5, pch = 19, asp = NA)
legend("bottomright", c("MR", "Dirichlet", "Polya-Gamma", "Polya-Gamma unshrunk"), col = c("blue", "orange", "darkgreen", "darkorchid4"), pch = 19)

dirnumtrue <- dirnumfalse <- pgnumtrue <- pgnumfalse <- mrnumtrue <- mrnumfalse <- pg1numtrue <- pg1numfalse <- vector(length = 10)


for (l in seq.int(10)){
  
  probdir <- probdirres[[l]]
  probpg <- probpgres[[l]]
  probpg1 <- probpgres1[[l]]
  mrres <- mrreslist[[l]]
  perm1_names <- perm1_nameslist[[l]]
  
  dirnumtrue[l] <- sum(names(probdir[order(probdir)])[(1 - probdir[order(probdir)]/ncol(resdyndir$allocOut[[1]])) > 0.999] %in% perm1_names)
  dirnumfalse[l] <- sum(!names(probdir[order(probdir)])[(1 - probdir[order(probdir)]/ncol(resdyndir$allocOut[[1]])) > 0.999] %in% perm1_names)
  pgnumtrue[l] <- sum(names(probpg[order(probpg)])[(1 - probpg[order(probpg)]/ncol(resdynpg$allocOut[[1]])) > 0.999] %in% perm1_names)
  pgnumfalse[l] <- sum(!names(probpg[order(probpg)])[(1 - probpg[order(probpg)]/ncol(resdynpg$allocOut[[1]])) > 0.999] %in% perm1_names)
  pg1numtrue[l] <- sum(names(probpg1[order(probpg1)])[(1 - probpg[order(probpg1)]/ncol(resdynpg1$allocOut[[1]])) > 0.999] %in% perm1_names)
  pg1numfalse[l] <- sum(!names(probpg1[order(probpg1)])[(1 - probpg[order(probpg1)]/ncol(resdynpg1$allocOut[[1]])) > 0.999] %in% perm1_names)
  mrnumtrue[l] <- sum(names(mrres$Mscore[(mrres$Mscore > 2) & mrres$Rscore > 0.9 ] ) %in% perm1_names)
  mrnumfalse[l] <- sum(!names(mrres$Mscore[(mrres$Mscore > 2) & mrres$Rscore > 0.9 ] ) %in% perm1_names)
}

numtrue <- cbind(dirnumtrue, pgnumtrue, mrnumtrue, pg1numtrue)

true_long <- as.data.frame(melt(numtrue[,c(3,1,2,4)]))
colnames(true_long) <- c("Iteration", "Method", "Relocalisations")

gg2 <- ggplot(data = true_long, aes(x = Method, y = Relocalisations, fill = Method)) + geom_boxplot(outlier.colour="black", outlier.shape=16,
                                                                                      outlier.size=2, notch=FALSE)
gg2 <- gg2 + theme_classic() + scale_fill_manual(values=c("blue", "orange", "darkgreen", "darkorchid4"))
gg2 <- gg2 + theme(legend.position="none")
gg2 <- gg2 + ggtitle("Correctly identified relocalisation events at 1% FDR") + xlab("Method") + ylab("Number of Relocalisations")
gg2 <- gg2 + scale_x_discrete(labels = c("MR", "Dirichlet", "Polya-Gamma","Polya-Gamma unshrunk"))
gg2


numfalse <- cbind(dirnumfalse, pgnumfalse, mrnumfalse, pg1numfalse)
abline(v = 0.99, col = "purple", lwd = 2)


par(mfrow = c(1,2))

plot2D(tan2009rep[[1]], grid = FALSE, main = "Example control spatial proteomics dataset")
plot2D(tan2009rep[[4]], grid = FALSE, main = "Example spatial proteomics dataset after cellular perturbation")
addLegend(where = "bottomright", tan2009rep[[4]], ncol = 3, cex = 0.9, xpd = TRUE)

dfmr <- as.data.frame(cbind(M = mrres$Mscore, R = mrres$Rscore))

gg3 <- ggplot(dfmr, aes(x = M , y = R)) + geom_point(col = "blue", alpha = 0.5, size = 4)
gg3 <- gg3 + theme_classic() 
gg3 <- gg3 + theme(axis.text.x=element_text(size=20), axis.text.y=element_text(size=20), axis.title=element_text(size=14))
gg3 <- gg3 + ggtitle("MR plot") + xlab("Movement Score") + ylab("Reproducibility Score")
gg3 <- gg3 + geom_hline(yintercept=0.9, color = "darkred",size=2) + geom_vline(xintercept = 2, color = "darkred",size=2)
gg3

probdir <- probsameorganelle(object = tan2009r1, params = resdyndir)
y <- 1 - probdir[order(probdir)]/ncol(resdyndir$allocOut[[1]])
par(mfrow = c(1, 1))
plot(y, pch = 19, ylim = c(0,1), ylab = "Differential Localisation Probability", xlab = "Rank", cex.axis = 1.2,
     col = "darkgreen", main = "Rank plot of differential localisation probability")

df <- data.frame(rank = seq.int(length(y)), y = y)
gg4 <- ggplot(df , aes(x = rank, y = y)) + geom_point(col = "darkgreen", alpha = 0.7, size = 3)
gg4 <- gg4 + theme_classic() + theme(axis.text.x = element_text(angle = 90), axis.text.y = element_text(size = 16),
                                     axis.title=element_text(size=14))
gg4 <- gg4 +  ggtitle("Rank plot of differential localisation probability") + xlab("Rank")
gg4 <- gg4 + ylab("Differential Localisation Probability")
gg4


par(mfrow = c(1,1))
bootresdir <- bootstrapdiffloc(object = tan2009r1, params = resdyndir, top = 100, Bootsample = 1000)
boxplot(t(1 - bootresdir/ncol(resdyndir$allocOut[[1]])), las = 2, col = getStockcol()[3])

df <- t(1 - bootresdir[1:40,]/ncol(resdyndir$allocOut[[1]]))
df_long <- melt(df)
colnames(df_long) <- c("bootsample", "Protein", "probability")

p <- ggplot(df_long, aes(x = Protein, y = probability)) + geom_boxplot(fill = getStockcol()[3])
p <- p + scale_y_continuous(limits = c(0.5, 1))
p <- p + theme_classic() + theme(axis.text.x = element_text(angle = 90), axis.text.y = element_text(size = 16),
                                 axis.title=element_text(size=14))
p <- p + labs(title = "Rank plot of top 40 differentially localised proteins",
              x = "Protein",
              y = "Differential Localisation Probability")
p 

require(patchwork)

gg4 + p

## form allocation probs
mcmcsampleprobs <- resdyndir$allocprob$cond11
dimnames(mcmcsampleprobs)[[1]] <- names(mostprob)
dimnames(mcmcsampleprobs)[[3]] <- getMarkerClasses(tan2009r1)

allocproborg <- apply(resdyndir$allocprob$cond1, c(1), function(x) colMeans(x))
allocprobmarker <- matrix(0, K, length(markerdata))

for (j in seq.int(K)) {
  
  allocprobmarker[j, markerdata == getMarkerClasses(tan2009r1)[j]] <- 1
  
}
colnames(allocprobmarker) <- names(markerdata)
allocproborg[,rowMeans(resdyndir$allocOut$cond11) < 0.99] <- 0 

allocproborg <- cbind(allocproborg, allocprobmarker)
colnames(allocproborg) <- c(names(mostprob), names(markerdata))


spatial2D <- function(object,
                      dims = c(1, 2),
                      cov.function = wendland.cov,
                      theta = 2,
                      derivative = 2,
                      k = 1,
                      breaks = c(0.99, 0.95, 0.9, 0.85, 0.8, 0.75, 0.7),
                      aspect = 0.5) {
  
  # generate pca plot and create data from with probabilties
  .pca <- plot2D(object, dims = dims, plot = FALSE)
  probs <- data.frame(x = .pca[, 1], y = .pca[, 2], mcmc.prob = t(allocproborg)[rownames(object),])
  colnames(probs) <- c(c("x", "y"), getMarkerClasses(object))
  eigs <- colnames(.pca)
  
  # put data in appropriate long format
  probs.lst <- list()
  for(j in getMarkerClasses(object)) {
    probs.lst[[j]] <- probs[, c("x", "y", j)]
    colnames(probs.lst[[j]]) <- c("x", "y", "probability")
  }
  probs.lst.df <- plyr::ldply(probs.lst, .fun = function(x) x, .id = "organelle")
  
  # Create storage
  coords <- list()
  locations <- list()
  df <- list()
  # Create appropriate spatial grid
  for (j in getMarkerClasses(object)) {
    idxOrg <- c(probs.lst.df$organelle == j)
    coords[[j]] <- akima::interp(x = probs.lst.df$x[idxOrg],
                                 y = probs.lst.df$y[idxOrg],
                                 z = probs.lst.df$probability[idxOrg],
                                 extrap=FALSE, linear = TRUE, duplicate = TRUE) # interpolate onto appropriate grid
    coords[[j]]$z[is.na(coords[[j]]$z)] <- 0 # NaNs beyond data set to 0
    locations[[j]] <- cbind(rep(coords[[j]]$x, 40), rep(coords[[j]]$y, each = 40)) # get grid
    smoothedprobs <- fields::smooth.2d(coords[[j]]$z, x = locations[[j]],
                                       cov.function = cov.function,
                                       theta = theta,
                                       derivative = derivative, k = k) # spatial smoothing of probabilities
    # normalisation and formatting
    zmat <- matrix(smoothedprobs$z, ncol = 1)
    zmat <- zmat/max(zmat)
    df[[j]] <- data.frame(x = rep(smoothedprobs$x, 64), y = rep(smoothedprobs$y, each = 64), z = zmat)
  }
  # format data
  df.lst <- plyr::ldply(df, .fun = function(x) x, .id = "organelle") 
  df.lst <- df.lst %>%
    mutate(organelle = factor(organelle)) 
  K <- length(getMarkerClasses(object))
  cols <- getStockcol()[1:K] # get appropriate colours
  
  
  gg <- ggplot(
    data = df.lst,
    aes(x = x, y = y, z = z, color = organelle)) +
    coord_fixed() + 
    geom_contour(breaks = breaks, size = 1.2, aes(alpha = stat(level))) + 
    geom_point(alpha = 0) + 
    xlab(paste0(eigs[1])) + 
    ylab(paste0(eigs[2])) +
    scale_alpha(guide = "none") + 
    theme(legend.position = "right", 
          text = element_text(size = 12)) +
    scale_color_manual(values = cols) +
    scale_fill_manual(values = cols) +
    theme_minimal() + 
    theme(panel.grid.major = element_blank(), panel.grid.minor = element_blank(),
          aspect.ratio = aspect,
          panel.border = element_rect(colour = "black", fill = NA, size = 1),
          plot.title = element_text(hjust = 0.5, size = 20),
          legend.text=element_text(size = 14)) +
    ggtitle(label = "Spatial variation of localisation probabilities") 
  gg
  return(gg)
}


.pca <- plot2D(tan2009rep[[1]], dims = dims, plot = FALSE)
gg <- gg + geom_point(data = data.frame(x = .pca[names(y[1]),1], y = .pca[names(y[1]),2]),
                aes(x = x, y = y, z = 1), colour = "black", size = 3, shape = 18)
gg

## form allocation probs for dataset 2
mcmcsampleprobs <- resdyndir$allocprob$cond21
dimnames(mcmcsampleprobs)[[1]] <- names(mostprob)
dimnames(mcmcsampleprobs)[[3]] <- getMarkerClasses(tan2009r1)

allocproborg <- apply(resdyndir$allocprob$cond1, c(1), function(x) colMeans(x))
allocprobmarker <- matrix(0, K, length(markerdata))

for (j in seq.int(K)) {
  
  allocprobmarker[j, markerdata == getMarkerClasses(tan2009r1)[j]] <- 1
  
}
colnames(allocprobmarker) <- names(markerdata)
allocproborg[,rowMeans(resdyndir$allocOut$cond21) < 0.99] <- 0 

allocproborg <- cbind(allocproborg, allocprobmarker)
colnames(allocproborg) <- c(names(mostprob), names(markerdata))


spatial2D <- function(object,
                      dims = c(1, 2),
                      cov.function = wendland.cov,
                      theta = 2,
                      derivative = 2,
                      k = 1,
                      breaks = c(0.99, 0.95, 0.9, 0.85, 0.8, 0.75, 0.7),
                      aspect = 0.5) {
  
  # generate pca plot and create data from with probabilties
  .pca <- plot2D(object, dims = dims, plot = FALSE)
  probs <- data.frame(x = .pca[, 1], y = .pca[, 2], mcmc.prob = t(allocproborg)[rownames(object),])
  colnames(probs) <- c(c("x", "y"), getMarkerClasses(object))
  eigs <- colnames(.pca)
  
  # put data in appropriate long format
  probs.lst <- list()
  for(j in getMarkerClasses(object)) {
    probs.lst[[j]] <- probs[, c("x", "y", j)]
    colnames(probs.lst[[j]]) <- c("x", "y", "probability")
  }
  probs.lst.df <- plyr::ldply(probs.lst, .fun = function(x) x, .id = "organelle")
  
  # Create storage
  coords <- list()
  locations <- list()
  df <- list()
  # Create appropriate spatial grid
  for (j in getMarkerClasses(object)) {
    idxOrg <- c(probs.lst.df$organelle == j)
    coords[[j]] <- akima::interp(x = probs.lst.df$x[idxOrg],
                                 y = probs.lst.df$y[idxOrg],
                                 z = probs.lst.df$probability[idxOrg],
                                 extrap=FALSE, linear = TRUE, duplicate = TRUE) # interpolate onto appropriate grid
    coords[[j]]$z[is.na(coords[[j]]$z)] <- 0 # NaNs beyond data set to 0
    locations[[j]] <- cbind(rep(coords[[j]]$x, 40), rep(coords[[j]]$y, each = 40)) # get grid
    smoothedprobs <- fields::smooth.2d(coords[[j]]$z, x = locations[[j]],
                                       cov.function = cov.function,
                                       theta = theta,
                                       derivative = derivative, k = k) # spatial smoothing of probabilities
    # normalisation and formatting
    zmat <- matrix(smoothedprobs$z, ncol = 1)
    zmat <- zmat/max(zmat)
    df[[j]] <- data.frame(x = rep(smoothedprobs$x, 64), y = rep(smoothedprobs$y, each = 64), z = zmat)
  }
  # format data
  df.lst <- plyr::ldply(df, .fun = function(x) x, .id = "organelle") 
  df.lst <- df.lst %>%
    mutate(organelle = factor(organelle)) 
  K <- length(getMarkerClasses(object))
  cols <- getStockcol()[1:K] # get appropriate colours
  
  
  gg2 <- ggplot(
    data = df.lst,
    aes(x = x, y = y, z = z, color = organelle)) +
    coord_fixed() + 
    geom_contour(breaks = breaks, size = 1.2, aes(alpha = stat(level))) + 
    geom_point(alpha = 0) + 
    xlab(paste0(eigs[1])) + 
    ylab(paste0(eigs[2])) +
    scale_alpha(guide = "none") + 
    theme(legend.position = "right", 
          text = element_text(size = 12)) +
    scale_color_manual(values = cols) +
    scale_fill_manual(values = cols) +
    theme_minimal() + 
    theme(panel.grid.major = element_blank(), panel.grid.minor = element_blank(),
          aspect.ratio = aspect,
          panel.border = element_rect(colour = "black", fill = NA, size = 1),
          plot.title = element_text(hjust = 0.5, size = 20),
          legend.text=element_text(size = 14)) +
    ggtitle(label = "Spatial variation of localisation probabilities") 
  gg2
  return(gg)
}

.pca <- plot2D(tan2009rep[[4]], dims = dims, plot = FALSE)
gg2 <- gg2 + geom_point(data = data.frame(x = .pca[names(y[1]),1], y = .pca[names(y[1]),2]),
                   aes(x = x, y = y, z = 1), colour = "black", size = 3, shape = 18)


gg + gg2
```
