## Supplementary Code 1 for "Inferring differential subcellular localisation in comparative spatial proteomics using BANDLE": methodtesting4.R

```
##
data("lopitdcU2OS2018")

simres <- sim_dynamic(object = lopitdcU2OS2018[,c(1:10)], subnum = 1000)
lopitDCrep <- simres$lopitrep
lapply(lopitDCrep, plot2D)
perm1_names <- simres$perm1_names

mrres <- mrMethod(c(lopitDCrep))

resdynpg <- diffLoc(object_cond1 = c(tan2009rep[1:3]), object_cond2 = tan2009rep[4:6], numIter = 5000, burnin = 1000, pg = TRUE)
resdyndir <- diffLoc(object_cond1 = c(lopitDCrep[1:3]), object_cond2 = c(lopitDCrep[4:6]),
                     numIter = 5000, burnin = 1000, pg = FALSE, fcol = "markers")
probpg <- probsameorganelle(object = lopitDCrep, params = resdynpg)
probdir <- probsameorganelle(object = simres$lopitrep[[1]], params = resdyndir)


outcome <- matrix(0, ncol = 1, nrow = length(probdir))
names(outcome) <- names(probdir)
outcome[perm1_names] <- 1
prediction <- probdir/ncol(resdyndir$allocOut[[1]])
par(mfrow = c(1,1))
plot(mrres$Mscore, mrres$Rscore, col = outcome[names(mrres$Mscore)] + 2, pch = 19, ylim = c(-1,1))
abline(h = 0.9, col = "blue", lwd = 2)
abline(v = 2, col = "purple", lwd = 2)

ROC_dir <- roc(outcome, 1 - prediction)
ROC_mr <- roc(outcome, mrres$MRscore[names(outcome)], direction = "<")
plot(ROC_dir, type = "b", col = "darkgreen", lwd = 1, pch = 19, asp = NA, legacy.axes = T)
lines(ROC_mr, type = "b", col = "blue", lwd = 1, pch = 19, asp = NA)


priornotAlloc <- vector(length = 5000)
nkknown <- table(getMarkers(lopitdcU2OS2018, verbose = FALSE))[getMarkerClasses(lopitdcU2OS2018)]
for (i in 1:5000) {
  
  concentration <- diag(rep(10, K)) + diag(nkknown) + matrix(0.1, nrow = K, ncol = K)
  currentweights <- t(sampleDirichlet(K^2, c(concentration)))
  
  priornotAlloc[i] <- sum(currentweights[1, -c((K+1) * seq.int(1:K) - K)])
}
hist(priornotAlloc)
# average number of differential localisations
mean(priornotAlloc) * nrow(unknownMSnSet(lopitdcU2OS2018))

# probability of having greater than 15 
sum((priornotAlloc * nrow(unknownMSnSet(lopitdcU2OS2018))) > 15)/5000
```
