## Supplementary Code 1 for "Inferring differential subcellular localisation in comparative spatial proteomics using BANDLE": methodtesting.R

```
## Method testing

data("tan2009r1")
test1 <- tan2009r1
test2 <- tan2009r1

mtan2009 <- unknownMSnSet(tan2009r1)
perm1 <- sample.int(nrow(mtan2009), size = 20, replace = TRUE)
perm1_names <- rownames(mtan2009)[perm1]

M <- matrix(NA, nrow = length(getMarkerClasses(tan2009r1)), ncol = ncol(tan2009r1))
V <- matrix(NA, nrow = length(getMarkerClasses(tan2009r1)), ncol = ncol(tan2009r1))
rownames(M) <- rownames(V) <- getMarkerClasses(tan2009r1)

for (j in getMarkerClasses(tan2009r1)) {
  M[j, ] <- colMeans(exprs(tan2009r1)[fData(tan2009r1)$markers == j,])
  V[j, ] <- apply(exprs(tan2009r1)[fData(tan2009r1)$markers == j,], 2, var)
}
sigma1 <- cov(t(M))

M2 <- matrix(NA, nrow = length(getMarkerClasses(test2)), ncol = ncol(test2))
rownames(M2) <- getMarkerClasses(test2)
for (j in getMarkerClasses(test2)) {
  M2[j, ] <- colMeans(exprs(test2)[fData(test2)$markers == j,])
}
sigma2 <- cov(t(M2))


# Simulate dynamics
for (i in perm1) {
  
  K <- length(getMarkerClasses(tan2009r1))
  r <- sample.int(K, size = 1)
  exprs(mtan2009)[i, ] <- M[r,] + rnorm(ncol(tan2009r1), mean = 0, sd = mean(sqrt(V)))
}

exprs(test2)[perm1_names,] <- exprs(mtan2009)[perm1_names, ]
par(mfrow = c(1,2))
plot2D(object = test2)
highlightOnPlot(object = test2, foi = perm1_names, TRUE)
plot2D(object = test1)
highlightOnPlot(object = test1, foi = perm1_names, labels = TRUE)
highlightOnPlot(object = test1, foi = "Q9VDD1")


test1b <- test1
test1c <- test1
test2b <- test2
test2c <- test2
# exprs(test1b) <- exprs(test1) + rt(ncol(test1) * nrow(test1), df = 20)#+ rnorm(n = ncol(test1) * nrow(test1), mean = 0, sd = 0.15)
# exprs(test1c) <- exprs(test1) + rt(ncol(test1) * nrow(test1), df = 20)#+ rnorm(n = ncol(test1) * nrow(test1), mean = 0, sd = 0.15)
# exprs(test2b) <- exprs(test2) + rt(ncol(test1) * nrow(test1), df = 20)#+ rnorm(n = ncol(test2) * nrow(test2), mean = 0, sd = 0.15)
# exprs(test2c) <- exprs(test2) + rt(ncol(test1) * nrow(test1), df = 20)#+ rnorm(n = ncol(test2) * nrow(test2), mean = 0, sd = 0.15)
exprs(test1b) <- exprs(test1) + rnorm(n = ncol(test1) * nrow(test1), mean = 0, sd = 2*min(sqrt(V)))
exprs(test1c) <- exprs(test1) + rnorm(n = ncol(test1) * nrow(test1), mean = 0, sd = 2*min(sqrt(V)))
exprs(test2b) <- exprs(test2) + rnorm(n = ncol(test2) * nrow(test2), mean = 0, sd = 2*min(sqrt(V)))
exprs(test2c) <- exprs(test2) + rnorm(n = ncol(test2) * nrow(test2), mean = 0, sd = 2*min(sqrt(V)))
exprs(test2) <- exprs(test2) + rnorm(n = ncol(test2) * nrow(test2), mean = 0, sd = 2*min(sqrt(V)))

plot2D(test1b)
plot2D(test1)
plot2D(test2)

restempdyn <- diffLoc(object_cond1 = c(test1, test1b, test1c), object_cond2 = c(test2, test2b, test2c), numIter = 5000, burnin = 1000, pg = TRUE)
restempdyn2 <- diffLoc(object_cond1 = c(test1, test1b, test1c), object_cond2 = c(test2, test2b, test2c), numIter = 5000, burnin = 1000, pg = FALSE)
#restempdyn <- diffLoc(object_cond1 = c(test1), object_cond2 = c(test2), numIter = 10, burnin = 1, thin = 1)


probsameorganelle <- vector(mode = "numeric", length = nrow(unknownMSnSet(tan2009r1)))
for (i in 1:nrow(unknownMSnSet(tan2009r1))) {
  probsameorganelle[i] <- sum(restempdyn$alloc[[1]][i, ] == restempdyn$alloc[[2]][i, ])
}
names(probsameorganelle) <- rownames(unknownMSnSet(tan2009r1))

probsameorganelle2 <- vector(mode = "numeric", length = nrow(unknownMSnSet(tan2009r1)))
for (i in 1:nrow(unknownMSnSet(tan2009r1))) {
  probsameorganelle2[i] <- sum(restempdyn2$alloc[[1]][i, ] == restempdyn2$alloc[[2]][i, ])
}
names(probsameorganelle2) <- rownames(unknownMSnSet(tan2009r1))


names(probsameorganelle)[probsameorganelle/1000 < 0.9]

names(probsameorganelle)[probsameorganelle/ncol(restempdyn$allocOut[[1]]) < 0.5] %in% perm1_names
sum(names(probsameorganelle)[probsameorganelle/ncol(restempdyn$allocOut[[1]]) < 0.5] %in% perm1_names)

which(rownames(unknown_cmb$cond1) == "Q9VDD1")

r <- ncol(restempdyn$allocOut[[1]])
ROC_bayes <- matrix(NA, nrow = r, ncol = 2)
PrecRecall_bayes <- matrix(NA, nrow = r, ncol = 2)
for (i in seq.int(r)) {
  
  TP <- sum(names(probsameorganelle)[probsameorganelle < i] %in% perm1_names)
  FN <- sum(!(perm1_names %in% names(probsameorganelle)[probsameorganelle < i]))
  TN <- sum(!(names(probsameorganelle)[probsameorganelle > i] %in% perm1_names))
  FP <- sum(!(names(probsameorganelle)[probsameorganelle < i]  %in% perm1_names))
  
  ROC_bayes[i, 1] <- 1 - (TN/(TN + FP))
  ROC_bayes[i, 2] <- TP/(TP + FN)
  
  PrecRecall_bayes[i, 1] <- TP/(TP + FN) # recall
  PrecRecall_bayes[i, 2] <- TP/(TP + FP) # precision
  
  
}

require(pROC)
plot(ROC_bayes, type = "b", col = "darkgreen", lwd = 1, ylim = c(0,1), xlim = c(0,1), pch = 19)
points(ROC, type = "b", col = "blue", lwd = 1, ylim = c(0,1), xlim = c(0,1), pch = 19) #From MR method
points(seq.int(100)/100, seq.int(100)/100, type = "b", col = "red", pch = 19)

outcome <- matrix(0, ncol = 1, nrow = length(probsameorganelle))
names(outcome) <- names(probsameorganelle)
outcome[perm1_names] <- 1
prediction <- probsameorganelle/ncol(restempdyn$allocOut[[1]])
par(mfrow = c(1,1))

ROC_bayes <- roc(outcome, 1 - prediction)
ROC_mr <- roc(outcome, MRscore[names(outcome)], direction = "<")
plot(ROC_bayes, type = "b", col = "darkgreen", lwd = 1, pch = 19, asp = NA, legacy.axes = T)
lines(ROC_mr, type = "b", col = "blue", lwd = 1, pch = 19, asp = NA)

outcome2 <- matrix(0, ncol = 1, nrow = length(probsameorganelle2))
names(outcome2) <- names(probsameorganelle2)
outcome2[perm1_names] <- 1
prediction2 <- probsameorganelle2/ncol(restempdyn2$allocOut[[1]])
ROC_dir <- roc(outcome2, 1 - prediction2)
lines(ROC_dir,type = "b", col = "orange", lwd = 1, pch = 19, asp = NA)
legend("bottomright", c("MR", "Dirichlet", "Polya-Gamma"), col = c("blue", "orange", "darkgreen"), pch = 19)

plot(roc(outcome, MRscore[names(outcome)]),  type = "b", col = "darkgreen", lwd = 1, pch = 19, asp = NA, legacy.axes = TRUE, direction = ">")

tmp2 <- coords(tmp)

# Dirichlet
y <- 1 - probsameorganelle2[order(probsameorganelle2)]/ncol(restempdyn2$allocOut[[1]])
plot(y[1:20], col = outcome[order(probsameorganelle2)] + 2, pch = 19, ylim = c(0.5,1), ylab = "Differential Localisation Probability", xlab = "Rank")

#Polya-Gamma
pgy <- 1 - probsameorganelle[order(probsameorganelle)]/ncol(restempdyn$allocOut[[1]])
points(pgy, col = outcome[order(probsameorganelle)] + 2, pch = 24)
abline(h = 0.99, col = "blue", lwd = 2)
abline(h = 0.95, col = "purple", lwd = 2)


names(probsameorganelle2[order(probsameorganelle2)])[(1 - probsameorganelle2[order(probsameorganelle2)]/ncol(restempdyn2$allocOut[[1]])) > 0.99] %in% perm1_names
names(probsameorganelle[order(probsameorganelle)])[(1 - probsameorganelle[order(probsameorganelle)]/ncol(restempdyn$allocOut[[1]])) > 0.99] %in% perm1_names
names(Mscore[(Mscore > 2) & Rscore > 0.9 ] ) %in% perm1_names

plot(Mscore, Rscore, col = outcome[names(Mscore)] + 2, pch = 19, ylim = c(-1,1))
abline(h = 0.9, col = "blue", lwd = 2)
abline(v = 2, col = "purple", lwd = 2)


par(mfrow = c(1,2))
plot2D(test1b)
highlightOnPlot(object = test1b, foi = perm1_names, labels = TRUE)

plot2D(test2b)
highlightOnPlot(object = test2b, foi = perm1_names, labels = TRUE)
```
