## Supplementary Code 1 for "Inferring differential subcellular localisation in comparative spatial proteomics using BANDLE": plottingMetricResults2.rmd

````
---
title: "PlottingSimulation2"
author: "Oliver M. Crook"
date: "25 April 2020"
output: html_document
---

```{r,}
require(reshape2)

load("diffloc_sim_wild_dc.rda") # cluster specific bootstrap

```

```{r,}
auc <- matrix(unlist(lapply(c(sim_results$ROCresBayes,
                              sim_results$ROCresDir, 
                              sim_results$ROCresmr), function(x)x$auc)), ncol = 3, byrow = FALSE)
colnames(auc) <- c("PG", "Dir", "MR")
boxplot(auc[,c(3,2,1)], ylim = c(0.5,1), col = c("blue", "orange", "darkgreen"))

auc_long <- as.data.frame(melt(auc[,c(3,2,1)]))
colnames(auc_long) <- c("Iteration", "Method", "AUC")

gg <- ggplot(data = auc_long, aes(x = Method, y = AUC, fill = Method)) + geom_boxplot(outlier.colour="black", outlier.shape=16,
                                                                                      outlier.size=2, notch=FALSE)
gg <- gg + theme_classic() + scale_fill_manual(values=c("blue", "orange", "darkgreen", "darkorchid4"))
gg <- gg + theme(legend.position="none") + scale_y_continuous(limits = c(0.5,1))
gg <- gg + ggtitle("Cluster specific noise distribution") + xlab("Method") + ylab("AUC")
gg <- gg + scale_x_discrete(labels = c("MR", "Dirichlet", "Polya-Gamma"))
gg

ROC_bayes <- sim_results$ROCresBayes[[1]]
ROC_dir <- sim_results$ROCresDir[[1]]
ROC_mr <- sim_results$ROCresmr[[1]]


plot(ROC_bayes, type = "b", col = "darkgreen", lwd = 1.5, pch = 19, asp = NA, legacy.axes = T, 
     xlab = "False Positive Rate",
     ylab = "True Positive Rate",
     main = "ROC curve demonstrating proposed methods outperform current approach")
lines(ROC_mr, type = "b", col = "blue", lwd = 1.5, pch = 19, asp = NA)
lines(ROC_dir,type = "b", col = "orange", lwd = 1.5, pch = 19, asp = NA)
legend("bottomright", c("MR", "Dirichlet", "Polya-Gamma"), col = c("blue", "orange", "darkgreen"), pch = 19)

dirnumtrue <- dirnumfalse <- pgnumtrue <- pgnumfalse <- mrnumtrue <- mrnumfalse <- vector(length = 10)


for (l in seq.int(10)){
  
  probdir <- sim_results$probdirres[[l]]
  probpg <- sim_results$probpgres[[l]]
  mrres <- sim_results$mrreslist[[l]]
  perm1_names <- sim_results$perm1_nameslist[[l]]
  
  dirnumtrue[l] <- sum(names(probdir[order(probdir)])[(1 - probdir[order(probdir)]/800) > 0.999] %in% perm1_names)
  dirnumfalse[l] <- sum(!names(probdir[order(probdir)])[(1 - probdir[order(probdir)]/800) > 0.999] %in% perm1_names)
  pgnumtrue[l] <- sum(names(probpg[order(probpg)])[(1 - probpg[order(probpg)]/800) > 0.999] %in% perm1_names)
  pgnumfalse[l] <- sum(!names(probpg[order(probpg)])[(1 - probpg[order(probpg)]/800) > 0.999] %in% perm1_names)
  mrnumtrue[l] <- sum(names(mrres$Mscore[(mrres$Mscore > 2) & mrres$Rscore > 0.9 ] ) %in% perm1_names)
  mrnumfalse[l] <- sum(!names(mrres$Mscore[(mrres$Mscore > 2) & mrres$Rscore > 0.9 ] ) %in% perm1_names)
}

numtrue <- cbind(dirnumtrue, pgnumtrue, mrnumtrue)

true_long <- as.data.frame(melt(numtrue[,c(3,1,2)]))
colnames(true_long) <- c("Iteration", "Method", "Relocalisations")

gg2 <- ggplot(data = true_long, aes(x = Method, y = Relocalisations, fill = Method)) + geom_boxplot(outlier.colour="black", outlier.shape=16,
                                                                                                    outlier.size=2, notch=FALSE)
gg2 <- gg2 + theme_classic() + scale_fill_manual(values=c("blue", "orange", "darkgreen", "darkorchid4"))
gg2 <- gg2 + theme(legend.position="none")
gg2 <- gg2 + ggtitle("Cluster specific noise distribution") + xlab("Method") + ylab("Number of Relocalisations")
gg2 <- gg2 + scale_x_discrete(labels = c("MR", "Dirichlet", "Polya-Gamma","Polya-Gamma unshrunk"))
gg2

```

```{r,}
require(reshape2)

load("diffloc_sim_sys_dc.rda") # random batch effects

```


```{r,}
auc <- matrix(unlist(lapply(c(sim_results$ROCresBayes,
                              sim_results$ROCresDir, 
                              sim_results$ROCresmr), function(x)x$auc)), ncol = 3, byrow = FALSE)
colnames(auc) <- c("PG", "Dir", "MR")
boxplot(auc[,c(3,2,1)], ylim = c(0.5,1), col = c("blue", "orange", "darkgreen"))

auc_long <- as.data.frame(melt(auc[,c(3,2,1)]))
colnames(auc_long) <- c("Iteration", "Method", "AUC")

gg1 <- ggplot(data = auc_long, aes(x = Method, y = AUC, fill = Method)) + geom_boxplot(outlier.colour="black", outlier.shape=16,
                                                                                      outlier.size=2, notch=FALSE)
gg1 <- gg1 + theme_classic() + scale_fill_manual(values=c("blue", "orange", "darkgreen", "darkorchid4"))
gg1 <- gg1 + theme(legend.position="none") + scale_y_continuous(limits = c(0.5,1))
gg1 <- gg1 + ggtitle("Systematic batch effects") + xlab("Method") + ylab("AUC")
gg1 <- gg1 + scale_x_discrete(labels = c("MR", "Dirichlet", "Polya-Gamma"))
gg1

ROC_bayes <- sim_results$ROCresBayes[[1]]
ROC_dir <- sim_results$ROCresDir[[1]]
ROC_mr <- sim_results$ROCresmr[[1]]


plot(ROC_bayes, type = "b", col = "darkgreen", lwd = 1.5, pch = 19, asp = NA, legacy.axes = T, 
     xlab = "False Positive Rate",
     ylab = "True Positive Rate",
     main = "ROC curve demonstrating proposed methods outperform current approach")
lines(ROC_mr, type = "b", col = "blue", lwd = 1.5, pch = 19, asp = NA)
lines(ROC_dir,type = "b", col = "orange", lwd = 1.5, pch = 19, asp = NA)
legend("bottomright", c("MR", "Dirichlet", "Polya-Gamma"), col = c("blue", "orange", "darkgreen"), pch = 19)

dirnumtrue <- dirnumfalse <- pgnumtrue <- pgnumfalse <- mrnumtrue <- mrnumfalse <- vector(length = 10)


for (l in seq.int(10)){
  
  probdir <- sim_results$probdirres[[l]]
  probpg <- sim_results$probpgres[[l]]
  mrres <- sim_results$mrreslist[[l]]
  perm1_names <- sim_results$perm1_nameslist[[l]]
  
  dirnumtrue[l] <- sum(names(probdir[order(probdir)])[(1 - probdir[order(probdir)]/800) > 0.999] %in% perm1_names)
  dirnumfalse[l] <- sum(!names(probdir[order(probdir)])[(1 - probdir[order(probdir)]/800) > 0.999] %in% perm1_names)
  pgnumtrue[l] <- sum(names(probpg[order(probpg)])[(1 - probpg[order(probpg)]/800) > 0.999] %in% perm1_names)
  pgnumfalse[l] <- sum(!names(probpg[order(probpg)])[(1 - probpg[order(probpg)]/800) > 0.999] %in% perm1_names)
  mrnumtrue[l] <- sum(names(mrres$Mscore[(mrres$Mscore > 2) & mrres$Rscore > 0.9 ] ) %in% perm1_names)
  mrnumfalse[l] <- sum(!names(mrres$Mscore[(mrres$Mscore > 2) & mrres$Rscore > 0.9 ] ) %in% perm1_names)
}

numtrue <- cbind(dirnumtrue, pgnumtrue, mrnumtrue)

true_long <- as.data.frame(melt(numtrue[,c(3,1,2)]))
colnames(true_long) <- c("Iteration", "Method", "Relocalisations")

gg3 <- ggplot(data = true_long, aes(x = Method, y = Relocalisations, fill = Method)) + geom_boxplot(outlier.colour="black", outlier.shape=16,
                                                                                                    outlier.size=2, notch=FALSE)
gg3 <- gg3 + theme_classic() + scale_fill_manual(values=c("blue", "orange", "darkgreen", "darkorchid4"))
gg3 <- gg3 + theme(legend.position="none")
gg3 <- gg3 + ggtitle("Systematic batch effects") + xlab("Method") + ylab("Number of Relocalisations")
gg3 <- gg3 + scale_x_discrete(labels = c("MR", "Dirichlet", "Polya-Gamma","Polya-Gamma unshrunk"))
gg3

```

```{r,}
require(reshape2)

load("diffloc_sim_perm_dc.rda") # fraaction swapping

```


```{r,}
auc <- matrix(unlist(lapply(c(sim_results$ROCresBayes,
                              sim_results$ROCresDir, 
                              sim_results$ROCresmr), function(x)x$auc)), ncol = 3, byrow = FALSE)
colnames(auc) <- c("PG", "Dir", "MR")
boxplot(auc[,c(3,2,1)], ylim = c(0.5,1), col = c("blue", "orange", "darkgreen"))

auc_long <- as.data.frame(melt(auc[,c(3,2,1)]))
colnames(auc_long) <- c("Iteration", "Method", "AUC")

gg4 <- ggplot(data = auc_long, aes(x = Method, y = AUC, fill = Method)) + geom_boxplot(outlier.colour="black", outlier.shape=16,
                                                                                      outlier.size=2, notch=FALSE)
gg4 <- gg4 + theme_classic() + scale_fill_manual(values=c("blue", "orange", "darkgreen", "darkorchid4"))
gg4 <- gg4 + theme(legend.position="none") + scale_y_continuous(limits = c(0.5,1))
gg4 <- gg4 + ggtitle("Fraction swapping") + xlab("Method") + ylab("AUC")
gg4 <- gg4 + scale_x_discrete(labels = c("MR", "Dirichlet", "Polya-Gamma"))
gg4

ROC_bayes <- sim_results$ROCresBayes[[1]]
ROC_dir <- sim_results$ROCresDir[[1]]
ROC_mr <- sim_results$ROCresmr[[1]]


plot(ROC_bayes, type = "b", col = "darkgreen", lwd = 1.5, pch = 19, asp = NA, legacy.axes = T, 
     xlab = "False Positive Rate",
     ylab = "True Positive Rate",
     main = "ROC curve demonstrating proposed methods outperform current approach")
lines(ROC_mr, type = "b", col = "blue", lwd = 1.5, pch = 19, asp = NA)
lines(ROC_dir,type = "b", col = "orange", lwd = 1.5, pch = 19, asp = NA)
legend("bottomright", c("MR", "Dirichlet", "Polya-Gamma"), col = c("blue", "orange", "darkgreen"), pch = 19)

dirnumtrue <- dirnumfalse <- pgnumtrue <- pgnumfalse <- mrnumtrue <- mrnumfalse <- vector(length = 10)


for (l in seq.int(10)){
  
  probdir <- sim_results$probdirres[[l]]
  probpg <- sim_results$probpgres[[l]]
  mrres <- sim_results$mrreslist[[l]]
  perm1_names <- sim_results$perm1_nameslist[[l]]
  
  dirnumtrue[l] <- sum(names(probdir[order(probdir)])[(1 - probdir[order(probdir)]/800) > 0.999] %in% perm1_names)
  dirnumfalse[l] <- sum(!names(probdir[order(probdir)])[(1 - probdir[order(probdir)]/800) > 0.999] %in% perm1_names)
  pgnumtrue[l] <- sum(names(probpg[order(probpg)])[(1 - probpg[order(probpg)]/800) > 0.999] %in% perm1_names)
  pgnumfalse[l] <- sum(!names(probpg[order(probpg)])[(1 - probpg[order(probpg)]/800) > 0.999] %in% perm1_names)
  mrnumtrue[l] <- sum(names(mrres$Mscore[(mrres$Mscore > 2) & mrres$Rscore > 0.9 ] ) %in% perm1_names)
  mrnumfalse[l] <- sum(!names(mrres$Mscore[(mrres$Mscore > 2) & mrres$Rscore > 0.9 ] ) %in% perm1_names)
}

numtrue <- cbind(dirnumtrue, pgnumtrue, mrnumtrue)

true_long <- as.data.frame(melt(numtrue[,c(3,1,2)]))
colnames(true_long) <- c("Iteration", "Method", "Relocalisations")

gg5 <- ggplot(data = true_long, aes(x = Method, y = Relocalisations, fill = Method)) + geom_boxplot(outlier.colour="black", outlier.shape=16,
                                                                                                    outlier.size=2, notch=FALSE)
gg5 <- gg5 + theme_classic() + scale_fill_manual(values=c("blue", "orange", "darkgreen", "darkorchid4"))
gg5 <- gg5 + theme(legend.position="none")
gg5 <- gg5 + ggtitle("Fraction swapping") + xlab("Method") + ylab("Number of Relocalisations")
gg5 <- gg5 + scale_x_discrete(labels = c("MR", "Dirichlet", "Polya-Gamma","Polya-Gamma unshrunk"))
gg5

```

```{r,}
require(reshape2)

load("diffloc_sim_dc_wild_batch_rand.rda") # random batch effects

```


```{r,}
auc <- matrix(unlist(lapply(c(sim_results$ROCresBayes,
                              sim_results$ROCresDir, 
                              sim_results$ROCresmr), function(x)x$auc)), ncol = 3, byrow = FALSE)
colnames(auc) <- c("PG", "Dir", "MR")
boxplot(auc[,c(3,2,1)], ylim = c(0.5,1), col = c("blue", "orange", "darkgreen"))

auc_long <- as.data.frame(melt(auc[,c(3,2,1)]))
colnames(auc_long) <- c("Iteration", "Method", "AUC")

gg6 <- ggplot(data = auc_long, aes(x = Method, y = AUC, fill = Method)) + geom_boxplot(outlier.colour="black", outlier.shape=16,
                                                                                      outlier.size=2, notch=FALSE)
gg6 <- gg6 + theme_classic() + scale_fill_manual(values=c("blue", "orange", "darkgreen", "darkorchid4"))
gg6 <- gg6 + theme(legend.position="none") + scale_y_continuous(limits = c(0.5,1))
gg6 <- gg6 + ggtitle("Random bathc effects") + xlab("Method") + ylab("AUC")
gg6 <- gg6 + scale_x_discrete(labels = c("MR", "Dirichlet", "Polya-Gamma"))
gg6

ROC_bayes <- sim_results$ROCresBayes[[1]]
ROC_dir <- sim_results$ROCresDir[[1]]
ROC_mr <- sim_results$ROCresmr[[1]]


plot(ROC_bayes, type = "b", col = "darkgreen", lwd = 1.5, pch = 19, asp = NA, legacy.axes = T, 
     xlab = "False Positive Rate",
     ylab = "True Positive Rate",
     main = "ROC curve demonstrating proposed methods outperform current approach")
lines(ROC_mr, type = "b", col = "blue", lwd = 1.5, pch = 19, asp = NA)
lines(ROC_dir,type = "b", col = "orange", lwd = 1.5, pch = 19, asp = NA)
legend("bottomright", c("MR", "Dirichlet", "Polya-Gamma"), col = c("blue", "orange", "darkgreen"), pch = 19)

dirnumtrue <- dirnumfalse <- pgnumtrue <- pgnumfalse <- mrnumtrue <- mrnumfalse <- vector(length = 10)


for (l in seq.int(10)){
  
  probdir <- sim_results$probdirres[[l]]
  probpg <- sim_results$probpgres[[l]]
  mrres <- sim_results$mrreslist[[l]]
  perm1_names <- sim_results$perm1_nameslist[[l]]
  
  dirnumtrue[l] <- sum(names(probdir[order(probdir)])[(1 - probdir[order(probdir)]/800) > 0.999] %in% perm1_names)
  dirnumfalse[l] <- sum(!names(probdir[order(probdir)])[(1 - probdir[order(probdir)]/800) > 0.999] %in% perm1_names)
  pgnumtrue[l] <- sum(names(probpg[order(probpg)])[(1 - probpg[order(probpg)]/800) > 0.999] %in% perm1_names)
  pgnumfalse[l] <- sum(!names(probpg[order(probpg)])[(1 - probpg[order(probpg)]/800) > 0.999] %in% perm1_names)
  mrnumtrue[l] <- sum(names(mrres$Mscore[(mrres$Mscore > 2) & mrres$Rscore > 0.9 ] ) %in% perm1_names)
  mrnumfalse[l] <- sum(!names(mrres$Mscore[(mrres$Mscore > 2) & mrres$Rscore > 0.9 ] ) %in% perm1_names)
}

numtrue <- cbind(dirnumtrue, pgnumtrue, mrnumtrue)

true_long <- as.data.frame(melt(numtrue[,c(3,1,2)]))
colnames(true_long) <- c("Iteration", "Method", "Relocalisations")

gg7 <- ggplot(data = true_long, aes(x = Method, y = Relocalisations, fill = Method)) + geom_boxplot(outlier.colour="black", outlier.shape=16,
                                                                                                    outlier.size=2, notch=FALSE)
gg7 <- gg7 + theme_classic() + scale_fill_manual(values=c("blue", "orange", "darkgreen", "darkorchid4"))
gg7 <- gg7 + theme(legend.position="none")
gg7 <- gg7 + ggtitle("Random batch effect") + xlab("Method") + ylab("Number of Relocalisations")
gg7 <- gg7 + scale_x_discrete(labels = c("MR", "Dirichlet", "Polya-Gamma","Polya-Gamma unshrunk"))
gg7

```

```{r,}
require(reshape2)

load("diffloc_sim_dc_wild_batch_perm.rda") # fraaction swapping

```

```{r,}
auc <- matrix(unlist(lapply(c(sim_results$ROCresBayes,
                              sim_results$ROCresDir, 
                              sim_results$ROCresmr), function(x)x$auc)), ncol = 3, byrow = FALSE)
colnames(auc) <- c("PG", "Dir", "MR")
boxplot(auc[,c(3,2,1)], ylim = c(0.5,1), col = c("blue", "orange", "darkgreen"))

auc_long <- as.data.frame(melt(auc[,c(3,2,1)]))
colnames(auc_long) <- c("Iteration", "Method", "AUC")

gg8 <- ggplot(data = auc_long, aes(x = Method, y = AUC, fill = Method)) + geom_boxplot(outlier.colour="black", outlier.shape=16,
                                                                                      outlier.size=2, notch=FALSE)
gg8 <- gg8 + theme_classic() + scale_fill_manual(values=c("blue", "orange", "darkgreen", "darkorchid4"))
gg8 <- gg8 + theme(legend.position="none") + scale_y_continuous(limits = c(0.5,1))
gg8 <- gg8 + ggtitle("Fraction swapping and batch effects") + xlab("Method") + ylab("AUC")
gg8 <- gg8 + scale_x_discrete(labels = c("MR", "Dirichlet", "Polya-Gamma"))
gg8

ROC_bayes <- sim_results$ROCresBayes[[1]]
ROC_dir <- sim_results$ROCresDir[[1]]
ROC_mr <- sim_results$ROCresmr[[1]]


plot(ROC_bayes, type = "b", col = "darkgreen", lwd = 1.5, pch = 19, asp = NA, legacy.axes = T, 
     xlab = "False Positive Rate",
     ylab = "True Positive Rate",
     main = "ROC curve demonstrating proposed methods outperform current approach")
lines(ROC_mr, type = "b", col = "blue", lwd = 1.5, pch = 19, asp = NA)
lines(ROC_dir,type = "b", col = "orange", lwd = 1.5, pch = 19, asp = NA)
legend("bottomright", c("MR", "Dirichlet", "Polya-Gamma"), col = c("blue", "orange", "darkgreen"), pch = 19)

dirnumtrue <- dirnumfalse <- pgnumtrue <- pgnumfalse <- mrnumtrue <- mrnumfalse <- vector(length = 10)


for (l in seq.int(10)){
  
  probdir <- sim_results$probdirres[[l]]
  probpg <- sim_results$probpgres[[l]]
  mrres <- sim_results$mrreslist[[l]]
  perm1_names <- sim_results$perm1_nameslist[[l]]
  
  dirnumtrue[l] <- sum(names(probdir[order(probdir)])[(1 - probdir[order(probdir)]/800) > 0.999] %in% perm1_names)
  dirnumfalse[l] <- sum(!names(probdir[order(probdir)])[(1 - probdir[order(probdir)]/800) > 0.999] %in% perm1_names)
  pgnumtrue[l] <- sum(names(probpg[order(probpg)])[(1 - probpg[order(probpg)]/800) > 0.999] %in% perm1_names)
  pgnumfalse[l] <- sum(!names(probpg[order(probpg)])[(1 - probpg[order(probpg)]/800) > 0.999] %in% perm1_names)
  mrnumtrue[l] <- sum(names(mrres$Mscore[(mrres$Mscore > 2) & mrres$Rscore > 0.9 ] ) %in% perm1_names)
  mrnumfalse[l] <- sum(!names(mrres$Mscore[(mrres$Mscore > 2) & mrres$Rscore > 0.9 ] ) %in% perm1_names)
}

numtrue <- cbind(dirnumtrue, pgnumtrue, mrnumtrue)

true_long <- as.data.frame(melt(numtrue[,c(3,1,2)]))
colnames(true_long) <- c("Iteration", "Method", "Relocalisations")

gg9 <- ggplot(data = true_long, aes(x = Method, y = Relocalisations, fill = Method)) + geom_boxplot(outlier.colour="black", outlier.shape=16,
                                                                                                    outlier.size=2, notch=FALSE)
gg9 <- gg9 + theme_classic() + scale_fill_manual(values=c("blue", "orange", "darkgreen", "darkorchid4"))
gg9 <- gg9 + theme(legend.position="none")
gg9 <- gg9 + ggtitle("Fraction swapping and batch effects") + xlab("Method") + ylab("Number of Relocalisations")
gg9 <- gg9 + scale_x_discrete(labels = c("MR", "Dirichlet", "Polya-Gamma","Polya-Gamma unshrunk"))
gg9

```


```{r,}
require(patchwork)

gg + gg1 + gg4 + gg6 + gg8

```
````
