## Supplementary Code 1 for "Inferring differential subcellular localisation in comparative spatial proteomics using BANDLE": sensitivityAnalysis.Rmd

````
---
title: "Senstivity Analysis"
author: "Oliver M. Crook"
date: "24 April 2020"
output: html_document
---

```{r,}
source("simDynamic.R")

data("tan2009r1")
simdata <- sim_dynamic(object = tan2009r1)

tan2009rep <- simdata$lopitrep
perm1_names <- simdata$perm1_names
```


```{r,}
par(mfrow = c(3,4))
pc_prior <- matrix(NA, ncol = 3, 11)
pc_prior[seq.int(1:11),] <- matrix(rep(c(0.05,60,100), each = 11), ncol = 3)
simdatares_defaultprior <- diffLoc(object_cond1 = c(tan2009rep[1:3]), object_cond2 = c(tan2009rep[4:6]),
                                    numIter = 5000, burnin = 1000, pg = FALSE, hypLearn = "LFBGS", 
                                    maternCov = TRUE, nu = 2, pc_prior = pc_prior)

save(simdatares_defaultprior, file = "simdatares_defaultprior.rda")

```

```{r,}
par(mfrow = c(3,4))
pc_prior <- matrix(NA, ncol = 3, 11)
pc_prior[seq.int(1:11),] <- matrix(rep(c(0.05,60,100), each = 11), ncol = 3)
simdatares_jefprior <- diffLoc(object_cond1 = c(tan2009rep[1:3]), object_cond2 = c(tan2009rep[4:6]),
                                    numIter = 5000, burnin = 1000, pg = FALSE, hypLearn = "LFBGS", 
                                    maternCov = TRUE, nu = 2, pc_prior = pc_prior, dir_prior = matrix(0.5, 11, 11))

save(simdatares_jefprior, file = "simdatares_jefprior.rda")

```
```{r,}
par(mfrow = c(3,4))
pc_prior <- matrix(NA, ncol = 3, 11)
pc_prior[seq.int(1:11),] <- matrix(rep(c(0.05,60,100), each = 11), ncol = 3)
simdatares_symprior <- diffLoc(object_cond1 = c(tan2009rep[1:3]), object_cond2 = c(tan2009rep[4:6]),
                                    numIter = 5000, burnin = 1000, pg = FALSE, hypLearn = "LFBGS", 
                                    maternCov = TRUE, nu = 2, pc_prior = pc_prior, dir_prior = matrix(1, 11, 11))

save(simdatares_symprior, file = "simdatares_symprior.rda")
```


```{r,}
probdefault <- probsameorganelle(object = tan2009r1, params = simdatares_defaultprior)
probjef <- probsameorganelle(object = tan2009r1, params = simdatares_jefprior)
probsym <- probsameorganelle(object = tan2009r1, params = simdatares_symprior)


outcome <- matrix(0, ncol = 1, nrow = length(probdefault))
names(outcome) <- names(probdefault)
outcome[perm1_names] <- 1
prediction <- probdefault/ncol(simdatares_defaultprior$allocOut[[1]])
par(mfrow = c(1,1))

outcome2 <- matrix(0, ncol = 1, nrow = length(probjef))
names(outcome2) <- names(probjef)
outcome2[perm1_names] <- 1
prediction2 <- probjef/ncol(simdatares_jefprior$allocOut[[1]])
par(mfrow = c(1,1))

outcome3 <- matrix(0, ncol = 1, nrow = length(probsym))
names(outcome3) <- names(probsym)
outcome3[perm1_names] <- 1
prediction3 <- probsym/ncol(simdatares_symprior$allocOut[[1]])
par(mfrow = c(1,1))


ROC_default <- roc(outcome, 1 - prediction)
ROC_jef <- roc(outcome2, 1 - prediction2)
ROC_sym <- roc(outcome3, 1 - prediction3)


plot(ROC_default, type = "b", col = mycols[1], lwd = 4, pch = 19, asp = NA, legacy.axes = T, main = "Prior Sensitivity Analysis")
lines(ROC_jef, type = "b", col = mycols[2], lwd = 4, pch = 19, asp = NA)
lines(ROC_sym, type = "b", col = mycols[3], lwd = 4, pch = 19, asp = NA)
```
````
