## Supplementary Code 1 for "Inferring differential subcellular localisation in comparative spatial proteomics using BANDLE": bandleFUN.R

```
require(pRoloc)
require(pRolocdata)


# functions requried
source("diffLoc.R")
source("diffLocUtils.R")
source("fitGPmatern.R")
source("fitGPmaternPC.R")
source("metropolisGP.R")

bandle <- function(object_cond1,
                   object_cond2,
                   fcol = "markers",
                   hypLearn = "MH",
                   numIter = 1000,
                   burnin = 100L,
                   thin = 5L,
                   alpha = 1,
                   u = 2,
                   v = 10,
                   lambda = 1,
                   params = FALSE,
                   hypIter = 20,
                   hypMean = c(0,0,0),
                   hypSd = c(1,1,1),
                   seed = NULL,
                   pg = TRUE,
                   pgPrior = NULL,
                   tau = 0.2,
                   dir_prior = NULL,
                   maternCov = TRUE,
                   PC = TRUE,
                   pc_prior = c(0.5,3,100),
                   nu = 2,
                   propsd = c(0.3,0.1,0.05),
                   numChains = 4L,
                   BPPARAM = BiocParallel::bpparam()){

    ## chains run in parallel, repeating number of iterations
    .res <- bplapply(rep(numIter, numChains),
                     FUN = diffLoc,
                     object_cond1 = object_cond1,
                     object_cond2 = object_cond2,
                     fcol = fcol,
                     hypLearn = hypLearn,
                     #numIter = numIter,
                     burnin = burnin,
                     thin = thin,
                     alpha = alpha,
                     u = u,
                     v = v,
                     lambda = lambda,
                     params = params,
                     hypIter = hypIter,
                     hypMean = hypMean,
                     hypSd = hypSd,
                     seed = seed,
                     pg = pg,
                     pgPrior = pgPrior,
                     tau = tau,
                     dir_prior = dir_prior,
                     maternCov = maternCov,
                     PC = PC,
                     pc_prior = pc_prior,
                     nu = nu,
                     propsd = propsd,
                     BPPARAM = BPPARAM)

    return(.res)
}
```
