## Supplementary Code 1 for "Inferring differential subcellular localisation in comparative spatial proteomics using BANDLE": Bessel2ndKind.cpp

```
#include <RcppArmadillo.h>
#include <boost/math/special_functions/bessel.hpp>
// [[Rcpp::depends(RcppArmadillo)]]
// [[Rcpp::depends(BH)]]
using namespace Rcpp;

// [[Rcpp::export]]
double besselK_boost(double x, double v) {
  return boost::math::cyl_bessel_k(v, x);
}

// [[Rcpp::export]]
arma::mat besselK(arma::mat x,
                  double v) {
  int N = x.n_cols;
  arma::mat B(N, N);
  
for (int i = 0; i < N; i++){
  for (int j = 0; j < N; j ++){
    if (x(i, j) < pow(10, -8)){
      B(i, j) = 0;
    } else {
    B(i, j) = besselK_boost(x(i, j), v); 
    }
  }
}
  return(B);
}

// [[Rcpp::export]]
arma::mat matern(double nu,
                 double a,
                 double rho,
                 arma::vec tau,
                 int D){
  arma::mat S = arma::zeros(D, D);
  arma::mat kappa;
  arma::mat bes(D, D);
  arma::mat cov(D, D);
  
  
  S = S.each_col() + tau;
  kappa = sqrt(8 * nu) * abs(S - S.t())/rho;
  bes = besselK(kappa, nu);
  cov = a * a * (pow(2.0, 1 - nu)/exp(lgamma(nu))) * pow(kappa, nu) % bes;
  cov = cov + a * a * arma::eye(D, D);
  
  return(cov);
}

// [[Rcpp::export]]
List trenchDetcpp(arma::vec c) {
  
  int N = c.size();
  int i;
  arma::vec xi(N-1);
  arma::vec z(N-1);
  arma::vec v(N);
  double beta = 1;
  double alpha;
  double l = 0;
  double logdet;
  
  xi = c.subvec(1, N-1)/c(0);
  z(0) = - xi(0);
  alpha = -xi(0);
  int K = xi.size(); 
  
  for(i = 0; i < (K - 1); i++){
    beta = (1 - pow(alpha, 2)) * beta ;
    l = l + log(beta) ;
    if(i == 0){
      alpha = - (xi(i + 1) + xi(0) * z(0)) / beta ;
      z(0) = z(0) + alpha * z(0) ;
    }else{
      alpha = - (xi(i + 1) + dot(flipud(xi.subvec(0, i)), z.subvec(0, i)) ) / beta ;
      z.subvec(0, i) = z.subvec(0, i) + alpha * flipud(z.subvec(0, i)) ;
    }
    
    z(i+1) = alpha  ;
  }
  
  beta = (1 - pow(alpha, 2)) * beta ;
  l = l + log(beta) ;
  
  logdet = l + N * log(c(0));
  
  v(N-1) = 1 / ((1 + dot(xi, z)) * c(0)) ;
  v.subvec(0,N-2) = v(N-1) * flipud(z.subvec(0, N - 2));
  
  return List::create(Rcpp::Named("logdet") = logdet,
                      Rcpp::Named("z") = z,
                      Rcpp::Named("v") = v);
}

// [[Rcpp::export]]
arma::mat trenchInvcpp(arma::vec v) {
  
  int N = v.size();
  int i;
  int j;
  arma::mat C(N, N);
  arma::mat trenchInv;
  
  
  C.row(0) = flipud(v).t();
  C.col(0) = flipud(v);
  C.row(N - 1) = v.t();
  C.col(N - 1) = v;
  for(i = 1; i < floor( (N - 1) / 2 ) + 1; i++){
    for(j = 1; j < N - i; j++){
      C(i, j) = C(i - 1, j - 1) + (v(N - 1 - j) * v(N - 1 - i) - v(i - 1) * v(j - 1)) / v(N - 1) ;
      C(j, i) = C(i, j);
      C(N - 1 - i , N - 1 - j ) = C(i, j);
      C(N - 1 - j , N - 1 - i ) = C(i, j);
    }
  } 
  
  trenchInv = C;
  return(trenchInv) ; 
  
}

// [[Rcpp::export]]
arma::vec loglikeGPcpp(arma::vec Y,
                       arma::mat Z,
                       arma::mat A,
                       double logcovDet,
                       double sigmak,
                       double nk,
                       double D,
                       double Y2){
  
  arma::vec loglike;
  arma::mat I = arma::eye(D, D);
  arma::mat J = arma::ones(nk, nk);
  
  loglike = - Y2/ (2 * sigmak) + (Y.t() * ((I - Z) * Y))/ (2 * sigmak * nk) 
    -  logcovDet / 2 - (nk * D *  log(2 * M_PI)) / 2 ;
  
  return(loglike); 
}

// [[Rcpp::export]]
double likelihoodGPcpp(arma::mat &Xk,
                       arma::vec tau,
                       arma::vec h,
                       int nk,
                       int D,
                       bool materncov = 0,
                       const double nu = 2
){    
  double sigmak; sigmak = exp(2 * h(2));
  double a; a = exp(2 * h(1));
  double l; l = exp(h(0));
  double amatern; amatern = exp(h(1));
  double rhomatern; rhomatern = exp(h(0));

  double negloglike;
  double logdet;
  double logcovDet;
  double Y2; //keep track of sum of squares
  arma::mat Z(D,D);
  arma::mat S(D,D); S = arma::zeros(D,D);
  arma::mat A(D,D);
  arma::mat R(D,D);
  arma::vec temp;
  arma::vec v;
  arma::vec Y(D); //keep track of means
  List trenchres = List::create(Rcpp::Named("logdet"),
                                Rcpp::Named("z"),
                                Rcpp::Named("v"));
  
  S = S.each_col() + tau;
  if (materncov == 0) {
    A = a * exp(- pow(S - S.t(), 2)/l);
  } else {
    A = matern(nu, amatern, rhomatern, tau, D);
  }
  
  R = arma::eye(D,D) + (nk * A)/sigmak;
  
  trenchres = trenchDetcpp(R.row(0).t());
  logdet = trenchres["logdet"];
  v = as<arma::vec>(trenchres["v"]);
  
  
  Z = trenchInvcpp(v);
  logcovDet = (nk * D) * log(sigmak) + logdet ;
  
  Y2 = arma::accu(pow(Xk, 2));  
  Y = sum(Xk, 1);
  
  
  negloglike = -loglikeGPcpp(Y, Z, A, logcovDet, sigmak, nk, D, Y2)(0,0);
  
  if(ISNAN(logdet)){
    negloglike = R_NegInf;
    return(negloglike);
  } else {
    return(negloglike);
  }
}

// [[Rcpp::export]]
arma::vec gradientrhomatern(arma::vec Y,
                            arma::mat drvrhomatern,
                            double nk,
                            double D,
                            arma::mat Z,
                            arma::mat A,
                            double sigmak){
  
  arma::mat B;
  arma::mat J = arma::ones(nk, nk);
  arma::mat eye = arma::eye(D, D);
  arma::mat C;
  
  arma::vec gradientrhomatern;
  arma::mat invZA = Z * drvrhomatern;
  arma::mat invZdrv = invZA * Z;
  
  B = - (((Y.t() * invZdrv) * Y)) / (2 * sigmak * sigmak) ;
  C = trace(invZA)/2;
  
  gradientrhomatern = B - C;
  
  return(gradientrhomatern); 
}

// [[Rcpp::export]]
arma::vec gradientamatern(arma::vec Y,
                         double amatern,
                         double nk,
                         double D,
                         arma::mat Z,
                         arma::mat A,
                         double sigmak){
  
  arma::mat B;
  arma::mat J = arma::ones(nk, nk);
  arma::mat eye = arma::eye(D, D);
  arma::mat C;
  arma::vec gradientamatern;
  
  arma::mat invZdrv = (Z - Z * Z);
  B = - (((Y.t() * invZdrv) * Y)) / (amatern * sigmak * nk) ;
  C = trace(eye - Z) * sigmak/(2 * nk);
  
  gradientamatern = B - C;
  
  return(gradientamatern); 
}


arma::vec gradientNoise(arma::vec Y,
                        double drv,
                        double nk,
                        double sigmak,
                        arma::mat Z,
                        double D,
                        double Y2){
  
  arma::mat B;
  arma::mat J = arma::ones(nk, nk);
  arma::mat C;
  arma::mat eye = arma::eye(D,D);
  arma::vec gradientNoise;

  B = drv * Y2/(2 * sigmak * sigmak) + drv * Y.t() * ( Z * Z - eye) * Y /(nk * sigmak * sigmak * 2);
  C = drv * ((nk - 1) * D / sigmak + trace(Z)/sigmak)/2;
  
  gradientNoise = B - C;
  
  return(gradientNoise); 
}


// [[Rcpp::export]]
arma::vec gradientGPcppmatern(arma::mat &Xk,
                              arma::vec tau,
                              arma::vec h,
                              int nk,
                              int D,
                              double nu
){
  
  //terms are made unconstrained
  double sigmak; sigmak = exp(2 * h(2));
  double amatern; amatern = exp(h(1));
  double rhomatern; rhomatern = exp(h(0));
  double negloglike;
  double logdet;
  arma::mat Z(D,D);
  arma::mat S(D,D); S = arma::zeros(D,D);
  arma::mat A(D,D);
  arma::mat R(D,D);
  arma::mat B(D,D);
  arma::mat kappa;
  arma::mat covInv;
  arma::mat drvrhomatern;
  double drvsigma;
  arma::vec v;
  arma::vec grdrhomatern;
  arma::vec grdamatern;
  arma::vec grdsigma;
  arma::vec grad(3);
  arma::vec Y(D);
  double Y2;
  List trenchres = List::create(Rcpp::Named("logdet"),
                                Rcpp::Named("z"),
                                Rcpp::Named("v"));
  
  A = matern(nu, amatern, rhomatern, tau, D);
  R = arma::eye(D,D) + (nk * A)/sigmak;
  
  trenchres = trenchDetcpp(R.row(0).t());
  logdet = trenchres["logdet"];
  v = as<arma::vec>(trenchres["v"]);
  
  
  if(R_IsNA(logdet)){
    negloglike = R_NegInf;
    grad.row(0) = negloglike;
    return(grad);
  }
  
  Z = trenchInvcpp(v);
  
  // compute derivative of matern wrt to rho  
  S = S.each_col() + tau;
  kappa = sqrt(8 * nu) * abs(S - S.t())/rhomatern;
  B = - (amatern * amatern * pow(2.0, 1 - nu)/exp(lgamma(nu))) * (pow(kappa, nu + 1)/rhomatern) % besselK(kappa, nu - 1);
  drvrhomatern = - (A * nu/rhomatern) + (kappa * nu/ 2 * rhomatern) % A + B;
  
  // compute derivate of noise
  drvsigma = 2 * sigmak ;
  
  Y2 = arma::accu(pow(Xk, 2));  
  Y = sum(Xk, 1);
  
  grdamatern = gradientamatern(Y, amatern, nk, D, Z, A, sigmak);
  grdrhomatern = gradientrhomatern(Y, drvrhomatern, nk, D, Z, A, sigmak);
  grdsigma = gradientNoise(Y, drvsigma, nk, sigmak, Z , D, Y2);
  
  //return gradient of negative log like
  grad(0) = -grdrhomatern(0);
  grad(1) = -grdamatern(0);
  grad(2) = -grdsigma(0);
  
  return(grad);
}

// [[Rcpp::export]]
List LeapfrogGPcppPC(arma::mat Xk,
                     arma::vec lambda,
                     arma::vec tau,
                     arma::vec p,
                     arma::vec x,
                     arma::vec m,
                     int nk,
                     int D,
                     int L,
                     double delta,
                     double nu){
  
  arma::vec gradprior(3);
  gradprior(0) = 3/2 - lambda(0)*exp(-x(0)/2)/2; gradprior(1) = lambda(1) * exp(x(1)); gradprior(2) = -3 + lambda(2)*exp(x(2));
  
  for (int t=0; t < L; t++){
    //half step for momenta
    p = p - delta * (gradientGPcppmatern(Xk, tau, x , nk, D, nu) + gradprior)/2 ;
    //full step for position
    x = x + delta * p / m  ;
    //another half step for momenta
    p = p - delta * (gradientGPcppmatern(Xk, tau, x , nk, D, nu) + gradprior)/2 ;
  } 
  
  
  return List::create(Rcpp::Named("p") = p,
                      Rcpp::Named("x") = x
  );
  
}  

// [[Rcpp::export]]
arma::vec sampleGPmeanmaterncpp(arma::mat Xk,
                                arma::vec tau,
                                arma::vec h,
                                int nk,
                                int D,
                                double nu
){
  double sigmak; sigmak = exp(2 * h(2));
  double amatern; amatern = exp(h(1));
  double rhomatern; rhomatern = exp(h(0));
  arma::mat Z(D,D);
  arma::mat S(D,D); S = arma::zeros(D,D);
  arma::mat A(D,D);
  arma::mat R(D,D);
  arma::mat J = arma::ones(nk, nk);
  arma::mat invC(D * nk , D * nk);
  arma::mat priorCor(D, D * nk);
  arma::mat postV(D, D);
  arma::mat cholV(D, D);
  arma::vec v;
  arma::vec postM;
  arma::vec GPmean;
  arma::vec eigval;
  arma::mat eigvec;
  arma::mat eye = arma::eye(D, D);
  arma::vec Y(D);
  List trenchres = List::create(Rcpp::Named("logdet"),
                                Rcpp::Named("z"),
                                Rcpp::Named("v"));
  
  A = matern(nu, amatern, rhomatern, tau, D);
  R = arma::eye(D, D) + (nk * A)/sigmak;
  
  trenchres = trenchDetcpp(R.row(0).t());
  v = as<arma::vec>(trenchres["v"]);
  Z = trenchInvcpp(v);

  Y = sum(Xk, 1);
  postM =  ((eye - Z) * Y) / nk ;
  postV = (eye - Z)  * sigmak/nk;
  eig_sym(eigval, eigvec, postV);
  
  for(int i = 0; i < D; i++){
    if(eigval[i] < 0 ){
      eigval[i] = 10e-6;
    }
  }

  GPmean = postM + (eigvec * (eye.each_col() % sqrt(eigval))) * as<arma::vec>(rnorm(D));
  
  return(GPmean);
  
}
arma::mat subset(arma::mat &X,
                 arma::vec &B,
                 int j){
  
  arma::uvec ind_ = find(B==j);
  arma::mat subX ;
  
  subX = X.rows(ind_);
  
  return(subX);
  
}

arma::vec makevector(arma::mat &X){
  
  arma::vec Y;
  Y = arma::vectorise(X);
  return(Y);
  
}

// [[Rcpp::export]]
arma::mat makeComponent(arma::mat &X,
                        arma::vec &BX,
                        arma::mat &Y,
                        arma::vec &BY,
                        int j){
  arma::mat subX;
  arma::mat subY;
  arma::mat component;
  
  subX = subset(X, BX, j);
  subY = subset(Y, BY, j);
  
  component = arma::join_cols(subX, subY);
  
  return(component);
}

// [[Rcpp::export]]
arma::mat normalisedDatamatern(arma::mat &Xknown,
                               arma::vec &BX,
                               arma::mat &Xunknown,
                               arma::vec &BXun,
                               arma::vec &hypers,
                               arma::vec &nk,
                               arma::vec &tau,
                               int D,
                               int j,
                               double nu){
  arma::mat Xk;
  arma::vec sampleGPMean;
  int n = Xunknown.n_rows;
  arma::mat out(n, D);
  
  
  
  Xk = makeComponent(Xknown, BX, Xunknown, BXun, j);
  sampleGPMean = sampleGPmeanmaterncpp(Xk.t(), tau, hypers, nk(j-1), D, nu);
  for (int d = 0; d < D; d++) {
    for (int i = 0; i < n; i++) {
      out(i, d) = Xunknown(i, d) - sampleGPMean(d);
    }
  }

  return(out);
}

// [[Rcpp::export]]
List centeredDatamatern(arma::mat Xknown,
                        arma::vec BX,
                        arma::mat Xunknown,
                        arma::vec BXun,
                        arma::mat hypers,
                        arma::vec nk,
                        arma::vec tau,
                        int D,
                        int K,
                        double nu){
  int j;
  List centereddata(K);
  arma::vec currenthyper(3);
  
  for(j = 0; j < K; j++){
    currenthyper = hypers.row(j).t();
    centereddata[j] = normalisedDatamatern(Xknown,
                                           BX,
                                           Xunknown, 
                                           BXun,
                                           currenthyper, 
                                           nk, tau, D = D, j + 1, nu).t();
  }
  
  
  return(centereddata);
}

// [[Rcpp::export]]
arma::vec componentloglike(const arma::mat &centereddata,
                           double sigmak){
  int d;
  int D = centereddata.n_rows;
  int N = centereddata.n_cols;
  arma::vec likelihood;
  arma::mat likelihoods(N,D);
  arma::vec sumlikelihood;
  
  for(d = 0; d < D; d++){
    likelihood = dnorm(as<NumericVector>(wrap(centereddata.row(d))), 0, sigmak, 1);
    likelihoods.col(d) = likelihood;
  }
  
  sumlikelihood = rowSums(as<NumericMatrix>(wrap(likelihoods)));
  return(sumlikelihood);
  
}

// [[Rcpp::export]]
arma::mat comploglike(const List &centereddata,
                      arma::vec sigmak
){
  
  int K = sigmak.size();
  int N = as<arma::mat>(wrap(centereddata[1])).n_cols;
  int j;
  arma::mat likelihoods(N, K);
  
  for(j = 0; j < K; j++){
    likelihoods.col(j) = componentloglike(centereddata[j], sigmak(j)); 
  }
  
  return(likelihoods);
}

// [[Rcpp::export]]
List comploglikelist(const List &centereddata,
                     List sigmak) {
  
  int numRep = centereddata.size();
  int l;
  arma::mat loglikelihood;
  List loglikelihoods(numRep);
  
  for (l = 0; l < numRep; l++) {
    loglikelihoods[l] = comploglike(centereddata[l], sigmak[l]);
  }
  
  return(loglikelihoods);
  
}
```
