## Supplementary Code 1 for "Inferring differential subcellular localisation in comparative spatial proteomics using BANDLE": diffLoc.R

```
# Testing ground for diff Loc = 

require(Rcpp)
require(RcppArmadillo)
sourceCpp("dmvtCpp.cpp")
source("gradientGP.R")
sourceCpp("gradienthyp.cpp")
source("hamiltonianGP.R")
sourceCpp("leapfrogGPcpp.cpp")
sourceCpp("trenchDetcpp.cpp")
source("likelihoodGP.R")
source("metropolisGP.R")
source("fitGP.R")
sourceCpp("pg_sample.cpp")
sourceCpp("Bessel2ndKind.cpp")

require(lbfgs)

diffLoc <- function(object_cond1,
                    object_cond2,
                    fcol = "markers",
                    hypLearn = "MH",
                    numIter = 1000,
                    burnin = 100L,
                    thin = 5L,
                    alpha = 1,
                    u = 2,
                    v = 10,
                    lambda = 1,
                    params = NULL,
                    hypIter = 20,
                    hypMean = c(0,0,0),
                    hypSd = c(1,1,1),
                    seed = NULL,
                    pg = TRUE,
                    pgPrior = NULL,
                    tau = 0.2,
                    dir_prior = NULL,
                    maternCov = TRUE,
                    PC = TRUE,
                    pc_prior = c(0.5,3,100),
                    nu = 2,
                    propsd = c(0.3,0.1,0.05)){

  
  # Setting seed manually
  if (is.null(seed)) {
    seed <- sample(.Machine$integer.max, 1)
  }
  .seed <- as.integer(seed)  
  set.seed(.seed)
  
  
  # Elementary checks
  stopifnot(length(hypMean)==3)
  stopifnot(length(hypSd)==3)
  stopifnot(sum(hypSd > 0)==3)
  
  ## Samples to be retained as a result of burnin and thinning
  toRetain <- seq.int(burnin + 1L, numIter, thin)
  numRetained <- length(toRetain)
  
  # Normalising data
  hlnorm_cond1 <- lapply(object_cond1,  function(x) normalise(x, method = "center.mean"))
  hlnorm_cond2 <- lapply(object_cond2,  function(x) normalise(x, method = "center.mean"))
  
  # Bring data together
  object_cmb <- c(cond1 = hlnorm_cond1, cond2 = hlnorm_cond2)
  
  # numCond should be 2
  numCond <- 2
  numRepl <- length(object_cond1)
  
  # expressions from data and important summaries
  exprs_cmb <- lapply(object_cmb, exprs)
  M <- lapply(exprs_cmb, colMeans)
  V <- lapply(exprs_cmb, function(x) lambda * diag(diag(cov(x))) + diag(rep(10^{-6}, ncol(x))))

  # Getting data dimensions
  D <- ncol(object_cmb[[1]])
  K <- length(getMarkerClasses(object_cmb[[1]]))
  
  # construct empirical Bayes Polya-Gamma prior
  if (is.null(pgPrior)) {
   pgPrior <- pg_prior(hlnorm_cond1, hlnorm_cond2, K = K, pgPrior = NULL)
  }

  # Fit GPs to markers
  componenthypers <- lapply(object_cmb, function(x) vector(mode = "list", K))
  if (maternCov == FALSE) {
  res <- lapply(object_cmb, function(x) fitGP(x, fcol = fcol))
  } else if ((maternCov == TRUE) & (PC = FALSE) & (is.null(params))) {
  res <- lapply(object_cmb, function(x) fitGPmatern(x, fcol = fcol, nu = nu))
  } else if ((maternCov ==TRUE) & (PC = TRUE) & (is.null(params))) {
  res <- lapply(object_cmb, function(x) fitGPmaternPC(x, fcol = fcol, nu = nu, hyppar = pc_prior))  
  } else {
  res <- params
  }
  
  # seperate data into known and unknown
  unknown_cmb <- lapply(object_cmb, unknownMSnSet)
  exprsUnknown_cmb <- lapply(unknown_cmb, exprs)
  exprsKnown <- lapply(object_cmb, function(x) exprs(markerMSnSet(x, fcol = fcol)))
  
  # separate conditions allocations
  allocKnown <- lapply(object_cmb[c(1, numRepl + 1)], function(x) seq.int(K)[fData(markerMSnSet(x, fcol = fcol))[, fcol]])
  numProtein <- lapply(unknown_cmb[c(1, numRepl + 1)], nrow)
  
  # Some storage
  alloc <- lapply(numProtein, function(x) matrix(0, nrow = x, ncol = numRetained))
  allocOut <- lapply(numProtein, function(x) matrix(0, nrow = x, ncol = numRetained))
  outlierprob <- lapply(numProtein, function(x) matrix(0, nrow = x, ncol = numRetained))
  weights <- array(0, c(K, K, numRetained))
  epsilons <- matrix(0, nrow = 2, ncol = numRetained)
  #allocOutprob <- lapply(numProtein, function(x) array(0, c(x, numRetained, 2)))
  allocprob <- lapply(numProtein, function(x) array(0, c(x, numRetained, K)))
  loglikelihoods <- lapply(rep(numProtein, numRepl), function(x) matrix(0, nrow = x, ncol = K))
  
  #random allocation of unknown Proteins, allocations are condition specific
  alloctemp <- lapply(numProtein, function(x) sample.int(n = K, size = x, replace = TRUE))
  for (i in seq.int(numCond)) {
    
    object_cmb[[i]] <- knnClassification(object_cmb[[i]], k = 10)
    alloc[[i]][, 1] <- fData(object_cmb[[i]][rownames(unknown_cmb[[i]])])$knn
    
    #alloc[[i]][, 1] <- alloctemp[[i]]
  }
  
  # number of proteins allocated to each component
  nkknown <- lapply(object_cmb[c(1, numRepl + 1)], function(x)
                                      table(getMarkers(x, verbose = FALSE))[getMarkerClasses(x)])
  
  #number initial allocated to outlier component
  outlier <- vector(mode = "list", length = 2)
  for (i in seq.int(numCond)) {
    outlier[[i]] <- allocOut[[i]][,1]
  }
  
  sampleGPMean <- lapply(object_cmb, function(x) vector(mode = "list", length = K))
  centereddata <- lapply(object_cmb, function(x) vector(mode = "list", length = K))
  hypers <- lapply(object_cmb, function(x) vector(mode = "list", length = numRetained))
  .hypers <- vector(mode = "list", length = length(object_cmb))

  for (i in seq.int(object_cmb)) {
    hypers[[i]][[1]] <- res[[i]]$params
    .hypers[[i]] <- hypers[[i]][[1]]
  }
  # intialise polya-gamma auxiliary variables
  w <- rep(1, K^2)
  
  for(t in 2:numIter){
    
    # Between data allocation tally
    nk_mat <- diag(nkknown[[1]]) + table(factor(alloctemp[[1]], levels = 1:K),
                                          factor(alloctemp[[2]], levels = 1:K))
    
    # Within data allocation tally
    nk <- list(cond1 = rowSums(nk_mat), cond2 = colSums(nk_mat))
    
    if((t %% 1) ==0){
      print(t)
    }
    
    gpnk_cond1 <- gpnk_cond2 <- vector(mode = "numeric", length = K)
    gpnk <- list(gpnk_cond1 = gpnk_cond1, gpnk_cond2 = gpnk_cond2)
    
    for (i in seq.int(object_cmb)) {
      
     j <- ceiling(i/numRepl) # allocs don't change across replicates
     for (l in seq.int(K)) {
       gpnk[[j]][l] <- sum(alloctemp[[j]]*outlier[[j]] == l) + nkknown[[1]][l]
     }
      if (maternCov == FALSE) {
        centereddata[[i]] <- centeredData(Xknown = exprsKnown[[i]],
                                         BX = allocKnown[[j]],
                                         Xunknown = exprsUnknown_cmb[[i]], 
                                         BXun = alloctemp[[j]]*outlier[[j]],
                                         hypers = .hypers[[i]], 
                                         nk = gpnk[[j]], tau = seq.int(D), D = D, K)
      } else if (maternCov == TRUE) {  
        centereddata[[i]] <- centeredDatamatern(Xknown = exprsKnown[[i]],
                                                BX = allocKnown[[j]],
                                                Xunknown = exprsUnknown_cmb[[i]], 
                                                BXun = alloctemp[[j]]*outlier[[j]],
                                                hypers = .hypers[[i]], 
                                                nk = gpnk[[j]], tau = seq.int(D), D = D, K, nu = nu)
      }
    }

    # Polya-Gamma Sampler or Dirichlet weights
    if (pg == TRUE) {
      pgres <- sample_weights_pg(nk_mat = nk_mat, pgPrior = pgPrior, K = K, w = w, tau = tau)
      currentweights <- pgres$currentweights
      w <- pgres$w

    } else {
     if (is.null(dir_prior)){
       dir_prior <- diag(rep(1, K)) + matrix(0.05, nrow = K, ncol = K)
     }
      currentweights <- sample_weights_dir(nk_mat = nk_mat, dir_prior = dir_prior)
    }
    
    currentweights <- matrix(currentweights, ncol = K, nrow = K)
    # if((t %% 2) ==0){
    #   print(currentweights)
    #   print(nk_mat[3,3])
    #   print(w)
    # }
    # print(nk_mat)

    #extract noise component from hypers
    sigmak <- lapply(hypers, function(x) exp(2 * x[[1]][, 3])) # fixed for the moment
    sigmasqrt <- lapply(sigmak, sqrt)
    loglikelihoods <- comploglikelist(centereddata, sigmasqrt)
    
    resAllocCond1 <- proteinAllocation(loglikelihoods = loglikelihoods[1:numRepl],
                                   currentweights = currentweights, alloctemp = alloctemp, cond = 1)
    resAllocCond2 <- proteinAllocation(loglikelihoods = loglikelihoods[(1+numRepl):(numRepl*numCond)],
                                   currentweights = currentweights, alloctemp = alloctemp, cond = 2)
    
    alloctemp <- list(resAllocCond1$alloctemp, resAllocCond2$alloctemp)
    loglikelihoods_cond1 <- resAllocCond1$loglikelihoods_comb
    loglikelihoods_cond2 <- resAllocCond2$loglikelihoods_comb
    
    # sample epsilon
    tau1 <- lapply(outlier, function(x) sum(x == 1) + nrow(exprsKnown[[1]]))
    tau2 <- lapply(outlier, function(x) sum(x == 0))
    epsilon <- rbeta(n = 2, shape1 = u + unlist(tau2), shape2 = v + unlist(tau1))
    
    
    allocnotOutprob <- vector(mode = "list")
    allocOutprob <- vector(mode = "list")
    # Sample outlier allocations first condition 1 then condition 2
    # outlier likelihood
    outlierlikelihood <- vector(mode = "list")
    for (i in seq.int(exprsUnknown_cmb)) {
      outlierlikelihood[[i]] <- dmvtCpp(exprsUnknown_cmb[[i]], mu_ = M[[i]], sigma_ = V[[i]], df_ = 4, log_ = TRUE, isChol_ = F)
    }
    
    resOutCond1 <- outlierAllocationProbs(outlierlikelihood = outlierlikelihood[1:numRepl],
                                          loglikelihoods = loglikelihoods_cond1, epsilon = epsilon, alloctemp = alloctemp, cond = 1)
    resOutCond2 <- outlierAllocationProbs(outlierlikelihood = outlierlikelihood[(1+numRepl):(numRepl*numCond)],
                                          loglikelihoods = loglikelihoods_cond2, epsilon = epsilon, alloctemp = alloctemp, cond = 2)

    allocnotOutprob[[1]] <- resOutCond1[[1]]
    allocOutprob[[1]] <- resOutCond1[[2]]

    allocnotOutprob[[2]] <- resOutCond2[[1]]
    allocOutprob[[2]] <- resOutCond2[[2]]
    
    
    # Sample outliers
    allocoutlierprob <- cbind(allocnotOutprob[[1]], allocOutprob[[1]])
    outlier[[1]] <- sampleOutlier(allocoutlierprob = allocoutlierprob)
    
    allocoutlierprob <- cbind(allocnotOutprob[[2]], allocOutprob[[2]])
    outlier[[2]] <- sampleOutlier(allocoutlierprob = allocoutlierprob)
    
    #update hypers
    if((t %% hypIter) == 0){
    if (maternCov == FALSE) {  
      if(hypLearn == "LBFGS"){
      }else if(hypLearn == "MH"){
        # Between data allocation tally
        nk_mat <- diag(nkknown[[1]]) + table(factor(alloctemp[[1]], levels = 1:K),
                                             factor(alloctemp[[2]], levels = 1:K))
        
        # Within data allocation tally
        nk <- list(cond1 = rowSums(nk_mat), cond2 = colSums(nk_mat)) 

      for (i in seq.int(object_cmb)) {
        for(j in seq.int(K)){
          
          l <- ceiling(i/numRepl) # allocs don't change across replicates
          Y <- makeComponent(X = exprsKnown[[i]],
                             BX = allocKnown[[l]],
                             Y = exprsUnknown_cmb[[i]],
                             BY = alloctemp[[l]] * outlier[[l]],
                             j = j)
          componenthypers[[i]][[j]] <- metropolisGP(inith = .hypers[[i]][j,],
                                               X = t(Y),
                                               tau = seq.int(D), nk = nk[[l]][j],
                                               D = D,
                                               niter = 1,
                                               hypMean = hypMean,
                                               hypSd = hypSd)$h[ ,2]
          
          
        }
        # stores current hyperparameters invisibily
        .hypers[[i]] <- matrix(unlist(componenthypers[[i]]), ncol = 3, byrow = TRUE)
      }  
      } else if(hypLearn == "HMC"){
        print("HMC step performed")
        # Between data allocation tally
        nk_mat <- diag(nkknown[[1]]) + table(factor(alloctemp[[1]], levels = 1:K),
                                             factor(alloctemp[[2]], levels = 1:K))
        
        # Within data allocation tally
        nk <- list(cond1 = rowSums(nk_mat), cond2 = colSums(nk_mat)) 
        
        for (i in seq.int(object_cmb)) {
          for(j in seq.int(K)){
            l <- ceiling(i/numRepl) # allocs don't change across replicates
            Y <- makeComponent(X = exprsKnown[[i]],
                               BX = allocKnown[[l]],
                               Y = exprsUnknown_cmb[[i]],
                               BY = alloctemp[[l]] * outlier[[l]],
                               j = j)
            componenthypers[[i]][[j]] <- hamiltonianGP(inith = .hypers[[i]][j,],
                                                       tau = seq.int(D),
                                                       X = t(Y),
                                                       nk = nk[[l]][j],
                                                       D = D,
                                                       niter = 1,
                                                       stepsize = c(0.08,0.1),
                                                       steps = 10,
                                                       hypMean = hypMean,
                                                       hypSd = hypSd)$h[,2]
          }
          # stores current hyperparameters invisibily
          .hypers[[i]] <- matrix(unlist(componenthypers[[i]]), ncol = 3, byrow = TRUE)
        }
      }
    } else if ((maternCov == TRUE) & (hypLearn == "MH")) {
      # Between data allocation tally
      nk_mat <- diag(nkknown[[1]]) + table(factor(alloctemp[[1]], levels = 1:K),
                                           factor(alloctemp[[2]], levels = 1:K))
      
      print("here")
      # Within data allocation tally
      #nk <- list(cond1 = rowSums(nk_mat), cond2 = colSums(nk_mat)) 
      #nk <- lapply(tan2009rep, function(x) table(fData(x)[,fcol])[getMarkerClasses(x)])[c(1,4)]
      
      for (i in seq.int(object_cmb)) {
        for(j in seq.int(K)) {
          l <- ceiling(i/numRepl) # allocs don't change across replicates
          Y <- makeComponent(X = exprsKnown[[i]],
                             BX = allocKnown[[l]],
                             Y = exprsUnknown_cmb[[i]],
                             BY = - alloctemp[[l]] * 0,
                             j = j)
          
          hyperstoupdate <- c(exp(.hypers[[i]][j,1:2]), exp(2 * .hypers[[i]][j,3]))
          
          .pc_prior <- pc_prior[j, ]
          #newhypers <- hyperstoupdate
          newhypers <- metropolisGPmatern(inith = hyperstoupdate,
                                          X = t(Y),
                                          tau = seq.int(D),
                                          nk = nk[[l]][j],
                                          D = D,
                                          niter = 1,
                                          nu = nu,
                                          hyppar = .pc_prior,
                                          propsd = propsd)$h[ ,2]
          
          componenthypers[[i]][[j]] <- c(log(newhypers[1:2]), log(newhypers[3])/2)
        }
        # stores current hyperparameters invisibily
        .hypers[[i]] <- matrix(unlist(componenthypers[[i]]), ncol = 3, byrow = TRUE)
      
      }
    }
    }  

    ## Only store iterations that are going to be retained
    if(t %in% toRetain) {
    
      s <- which(t == toRetain) # index of variable to save
      for(i in seq.int(object_cmb)) {
        hypers[[i]][[s]] <- .hypers[[i]]
      }  
      weights[, , s] <- currentweights
      for (i in seq.int(numCond)) {
        alloc[[i]][, s] <- alloctemp[[i]]
        allocOut[[i]][, s] <- outlier[[i]]
        if (i == 1) {
        allocprob[[i]][, s, ] <- resAllocCond1$allocprobtemp
        } else {
        allocprob[[i]][, s, ] <- resAllocCond2$allocprobtemp
        }
      }  
      epsilons[, s] <- epsilon
    }  
  }
  
  return(list(allocOut = allocOut,
              alloc = alloc,
              allocprob = allocprob,
              hypers = hypers,
              weights = weights,
              epsilons = epsilons))
  
  
}

proteinAllocation <- function(loglikelihoods, currentweights, alloctemp, cond) {
  
  K <- ncol(currentweights)
  loglikelihoods_comb <- Reduce("+", loglikelihoods)
  if (cond == 1) {
   logconditionalprior <- t(log(currentweights[, alloctemp[[2]]])) # columns given condition 2
  } else {
   logconditionalprior <- log(currentweights[alloctemp[[1]], ]) # rows given condition 1
  }
  conditionalAlloc <- loglikelihoods_comb + logconditionalprior 
  cc <- apply(conditionalAlloc, 1, max)
  conditionalAlloc <- conditionalAlloc - cc # correct for underflow
  allocprobtemp <- exp(conditionalAlloc)/rowSums(exp(conditionalAlloc))
  #alloctemp <- apply(allocprobtemp, 1, function(x) sample.int(n = K, size = 1, replace = F, prob = x))
  alloctemp <- sampleAlloccpp(allocprobtemp)
  
  return(list(alloctemp = alloctemp, loglikelihoods_comb = loglikelihoods_comb, allocprobtemp = allocprobtemp))
}

outlierAllocationProbs <- function(outlierlikelihood, loglikelihoods, epsilon, alloctemp, cond) {
  
  subset <- cbind(1:length(alloctemp[[cond]]), alloctemp[[cond]])
  outlierlikelihood_comb <- Reduce("+", outlierlikelihood)
  allocnotOutprob <- log(1 - epsilon[cond]) + loglikelihoods[subset]
  allocOutprob <- log(epsilon[cond]) + outlierlikelihood_comb
  
  return(list(allocnotOutprob = allocnotOutprob, allocOutprob = allocOutprob))
  
}

sampleOutlier <- function(allocoutlierprob){
  
  c <- apply(allocoutlierprob , 1, max) 
  allocoutlierprob <- allocoutlierprob - c # correct for underflow
  allocoutlierprob <- exp(allocoutlierprob)/rowSums(exp(allocoutlierprob))
  
  outlier <- sampleOutliercpp(allocoutlierprob) # reversed sample so 2nd entry is prob of 0
  # outlier <- apply(allocoutlierprob, 1, function(z){
  #   sample(x = c(1, 0), 1, prob = z)}
  # ) # reversed sample so 2nd entry is prob of 0
  
  return(outlier)
}


covOrganelle <- function(mydata, fcol = "markers"){
  
  M <- matrix(NA, nrow = length(getMarkerClasses(mydata, fcol = fcol)), ncol = ncol(mydata))
  rownames(M) <- getMarkerClasses(mydata, fcol = fcol)
  for (j in getMarkerClasses(mydata, fcol = fcol)) {
    M[j, ] <- colMeans(exprs(mydata)[fData(mydata)[, fcol] == j,])
  }
  sigma <- cov(t(M))
  
  return(sigma)
}

pg_prior <- function(object_cond1, object_cond2, K, pgPrior = NULL) {
  
  if (is.null(pgPrior)) {
    mu_prior <- rep(-7, K^2)
    mu_prior[c(K+1 * seq.int(1:K) - K)] <- mu_prior[c(K+1 * seq.int(1:K) - K)] + 1
    sigma1 <- covOrganelle(mydata = object_cond1[[1]])
    sigma2 <- covOrganelle(mydata = object_cond2[[1]])
  }
  pgPrior <- list(mu_prior = mu_prior, sigma1 =  sigma1, sigma2 = sigma2)
  return(pgPrior)
}


sample_weights_pg <- function(nk_mat, pgPrior, w, K, tau = tau) {
  
  # Polya-Gamma prior
  mu_prior <- pgPrior$mu_prior
  sigma1 <- pgPrior$sigma1
  sigma2 <- pgPrior$sigma2
  
  # Polya-Gamma Sampler
  n_vec <- c(nk_mat)
  kappa <- n_vec - 1/2
  sigma_post <- solve(diag(w) + tau * kronecker(solve(t(sigma1) + diag(0.01, K)), solve(t(sigma2) + diag(0.01, K))))
  mu_post <- sigma_post %*% (kappa + tau * kronecker(solve(t(sigma1) + diag(0.01, K)), solve(t(sigma2) + diag(0.01, K))) %*% mu_prior)
  phi <- mu_post + chol(sigma_post) %*% rnorm(n = K^2, mean = 0, sd = rep(1, K^2))
  
  #stick breaking construction
  currentweights <- rep(0, length(mu_post))
  stick <- 1
  for (j in seq.int(K^2 - 1)) {
    currentweights[j] <- (1/(1 + exp(-phi[j]))) * stick # Stick-breaking construction
    stick <- stick - currentweights[j]
  }
  currentweights[K^2] <- stick
  
  # Sample Polya-Gamma Variable
  w <- rcpp_pgdraw(rep(1, K^2), phi)

  return(list(currentweights = currentweights, w = w))
}

sample_weights_dir <- function(nk_mat, dir_prior){
  
  #sample weights from dirichlet by normalising gammas
  K <- ncol(nk_mat)
  concentration <- dir_prior + nk_mat
  currentweights <- t(sampleDirichlet(K^2, c(concentration)))
  return(currentweights)
}
```
