## Supplementary Code 1 for "Inferring differential subcellular localisation in comparative spatial proteomics using BANDLE": diffLocUtils.R

```
##' These functions implement helper functions for the bandle method
##' 
##' 
##' @title Compute differential localisation probabilities from ms-based
##' experiments using the bandle method
##' @param params An instance of class `bandleParams`
##' @return  returns a named vector of differential localisation probabilities
##' @md
##' 
##' @rdname bandle
diffLocalisationProb <- function(params) {
  
  # Must be a valid bandleParams object
  stopifnot(class(params) == "bandleParams")
  
  res <- rowSums(1 * (params@chains@chains[[1]]@niche[[1]] - params@chains@chains[[1]]@niche[[2]]) != 0)
  res <- res/ncol(params@chains@chains[[1]]@niche[[1]])

  return(res)
}

##' @title Obtain bootstrap uncertainty estimates for differential localisations
##' probabilities
##' @param params An instance of class `bandleParams`
##' @param top The number of proteins for which to compute bootstrap distributions
##'  default is 20.
##' @param Bootsample Number of Bootstramp samples. Default is 5000
##' @param Descreasing Starting at proteins most likely to be differentially localised.
##'  default is 5000.
##' @return  returns a matrix of size Bootsample * top containing bootstrap 
##' @md
##' 
##' @rdname bandle
bootstrapdiffLocprob <- function(params,
                                 top = 20,
                                 Bootsample = 5000,
                                 decreasing = TRUE) {
  
  res <- matrix(NA, ncol = Bootsample, nrow = top)
  probs <- diffLocalisationProb(params = params)
  
  prBootnames <- names(probs[order(probs, decreasing = decreasing)][seq.int(top)])
  rownames(res) <- prBootnames
  
  for (t in seq.int(Bootsample)){
    bootidx <- sample.int(ncol(params@chains@chains[[1]]@niche[[1]]),
                          replace = TRUE)
    res[,t] <- rowSums(1 * (params@chains@chains[[1]]@niche[[1]][prBootnames, bootidx] - 
                          params@chains@chains[[1]]@niche[[2]][prBootnames, bootidx] != 0))/length(bootidx)
  }
  
  return(res)
}

##' @title Computes Organelle means and variances using markers
##' @param object a instance of class `MSnset`
##' @param fcol a feature column indicating which feature define the markers
##' @return returns a list of means and variances for each 
##' @md
##' 
##' @rdname bandle
meanOrganelle <- function(object, fcol = "markers"){
  
  stopifnot(class(object) == "MSnSet")
  
  M <- V <- matrix(NA, nrow = length(getMarkerClasses(object, fcol = fcol)), ncol = ncol(object))
  rownames(M) <- rownames(V) <-  getMarkerClasses(object, fcol = fcol)
  for (j in getMarkerClasses(object, fcol = fcol)) {
    M[j, ] <- colMeans(exprs(object)[fData(object)[, fcol] == j,])
    V[j, ] <- apply(exprs(object)[fData(object)[, fcol] == j,], 2, var)
  }
  
  return(list(M = M, V = V))
}

##' @title Computes the Kullback-Leiber divergence between Polya-Gamma and 
##' Dirichlet priors
##' @param sigma the sigma parameter of the Polya-Gamma prior
##' @param mu the mu parameter of the Polya-Gamma prior
##' @param alpha the alpha (concentration) parameter of the Dirichlet prior
##' @return returns a numeric indicating the KL divergence
##' @md
##' 
##' @rdname bandle

kldirpg <- function(sigma, mu, alpha) {
  
  D <- length(mu)
  entpg <- -log((2*pi*exp(1))^D)/2 - sum(log(eigen(sigma)$values))/2 # compute determinant from eigenvalues
  entdir <- sum(lgamma(alpha)) - lgamma(sum(alpha))
  entdirpg_1 <- c(alpha) * (log(1 + exp(-mu)) + exp(mu) * diag(sigma)/(1 + exp(mu))^2)
  entdirpg_2 <- c(alpha) * (log(1 + exp(mu)) - exp(mu) * diag(sigma)/(1 + exp(mu))^2)
  entdirpgsum_1 <- sum(entdirpg_1)[-D]
  entdirpgsum_2 <- sum(seq.int(1, D - 1) * entdirpg_2[2:D])
  
  res <- entpg + entdir + entdirpgsum_1 + entdirpgsum_2
  
  return(res)
}

##' @title  Compute the KL divergence between two Dirichlet distributions
##' 
##' @param alpha The concentration parameter of the first Dirichlet distribution
##' @param beta The concentration parameter of the second Dirichlet distribution
##' @return a numeric indicating the KL divergence
##' @md
##' 
##' @rdname bandle
kldir <- function(alpha, beta) {
  
  res <- lgamma(sum(alpha)) - sum(lgamma(alpha)) - lgamma(sum(beta)) + sum(lgamma(beta)) +
            sum((alpha - beta) * (digamma(alpha) - digamma(sum(alpha))))
  
  
  return(res)
  
}

##' @title A function to compute the prior predictive distribution of the 
##' Dirichet prior.
##' 
##' @param object An instance of class `MSnSet`
##' @param iter Number of sample to use from prior predictive distribution.
##'  Default is 5000
##' @param dirPrior The Dirichlet prior used. If NULL (default) will generate a 
##'  a default Dirichlet prior
##' @param q The upper tail value. That is the prior probability of having more 
##'  than q differential localisations. Default is 15. 
##' @return A list contain the prior predictive distribution of
##'   differential localisations, the mean number of differential localised proteins
##'   and the probability than more than q are differentially localised 
##' @md
##' 
##' @rdname bandle     
prior_pred_dir <- function(object,
                           iter = 5000,
                           dirPrior = NULL,
                           q = 15) {

  K <- length(getMarkerClasses(object))
  if (is.null(dirPrior)) {
    dirPrior <- diag(rep(1, K)) + matrix(0.05, nrow = K, ncol = K)
  }
  
  priornotAlloc <- vector(length = iter)
  nkknown <- table(getMarkers(object, verbose = FALSE))[getMarkerClasses(object)]
  for (i in seq.int(iter)) {
    
    concentration <- diag(nkknown) + dirPrior
    currentweights <- t(sampleDirichlet(K^2, c(concentration)))
    priornotAlloc[i] <- sum(currentweights[1, -c((K + 1) * seq.int(1:(K)) - K)])
  }

  # average number of differential localisations
  meannotAlloc <- mean(priornotAlloc) * nrow(unknownMSnSet(object))
  
  # probability of having greater than q
  tailnotAlloc <- sum((priornotAlloc * nrow(unknownMSnSet(object))) > q)/iter

  return(list(priornotAlloc = priornotAlloc,
              meannotAlloc = meannotAlloc,
              tailnotAlloc = tailnotAlloc))
}

##' @title A function to compute the prior predictive distribution of the 
##' Polya-Gamma prior.
##' 
##' @param objectCond1 An instance of class `MSnSet`, usually the control dataset
##' @param objectCond2 An instance of class `MSnSet`, usually the treatment dataset
##' @param tau The `tau` parameter of the Polya-Gamma prior. Default is 0.2.
##' @param lambda The `lambda` ridge parameter used for numerical stability. 
##'  Default is 0.01
##' @param mu_prior The mean of the Polya-Gamma prior. Default is NULL which generates
##'  a default Polya-Gamma prior.  
##' @param iter Number of sample to use from prior predictive distribution.
##'  Default is 10000
##' @param q The upper tail value. That is the prior probability of having more 
##'  than q differential localisations. Default is 15. 
##' @return A list contain the prior predictive distribution of
##'   differential localisations, the mean number of differential localised proteins
##'   and the probability than more than q are differentially localised 
##' @md
##' 
##' @rdname bandle     
prior_pred_pg <- function(objectCond1,
                          objectCond2,
                          tau = 0.2,
                          lambda = 0.01,
                          mu_prior = NULL,
                          iter = 10000,
                          q = 15) {

  stopifnot(class(objectCond1) == "MSnSet")
  stopifnot(class(objectCond2) == "MSnSet")
  
  # Expressions from data and important summaries
  mydata <- exprs(objectCond1)
  M <- colMeans(mydata)
  V <- cov(mydata)
  nkknown <- table(getMarkers(objectCond1, verbose = FALSE))[getMarkerClasses(objectCond1)]
  K <- length(getMarkerClasses(objectCond1))
  
  sigma1 <- covOrganelle(mydata = objectCond1)
  sigma2 <- covOrganelle(mydata = objectCond2)
  w <- rep(1, K^2)
  
  n_vec <- c(diag(nkknown))
  kappa <- n_vec - 1/2
  sigma_post <- solve(diag(w) + tau * kronecker(solve(t(sigma1) + diag(lambda, K)), solve(t(sigma2) + diag(lambda, K))))
  
  if (is.null(mu_prior)) {
    mu_prior <- rep(-9, K^2)
    mu_prior[c((K + 1) * seq.int(1:(K)) - K)] <- mu_prior[c((K + 1) * seq.int(1:(K)) - K)] + 1
  }
  
  
  mu_post <- sigma_post %*% (kappa + tau * kronecker(solve(t(sigma1) + diag(lambda, K)),
                                                     solve(t(sigma2) + diag(lambda, K))) %*% mu_prior)
  
  priornotAllocpg <- vector(length = iter)
  for (i in seq.int(iter)) {
    phi <- mu_post + chol(sigma_post) %*% rnorm(n = K^2, mean = 0, sd = rep(1, K^2))
    
    currentweights <- rep(0, length(mu_post))
    
    stick <- 1
    for (j in seq.int(K^2 - 1)) {
      currentweights[j] <- (1/(1 + exp(-phi[j]))) * stick # Stick-breaking construction
      stick <- stick - currentweights[j]
    }
    currentweights[K^2] <- stick
    
    w <- rcpp_pgdraw(rep(1, K^2), phi)
    priornotAllocpg[i] <- sum(currentweights[-c((K + 1) * seq.int(1:(K)) - K)])
  }
  
  # average number of differential localisations
  meannotAlloc <- mean(priornotAllocpg) * nrow(unknownMSnSet(objectCond1))
  varnotAlloc <- var(priornotAllocpg)
  
  # probability of having greater than 15 
  tailnotAlloc <- sum((priornotAllocpg * nrow(unknownMSnSet(objectCond1))) > q)/iter


return(list(priornotAllocpg = priornotAllocpg,
            meannotAlloc = meannotAlloc,
            varnotAlloc = varnotAlloc,
            tailnotAlloc = tailnotAlloc))
}
```
