## Supplementary Code 1 for "Inferring differential subcellular localisation in comparative spatial proteomics using BANDLE": dmvtCpp.cpp

```
# include <RcppArmadillo.h>
// [[Rcpp::depends(RcppArmadillo)]]

using namespace Rcpp ;
//'  Copyright (C) 2014 Matteo Fasiolo 
//'This program is free software; you can redistribute it and/or
//' modify it under the terms of the GNU General Public License
//' as published by the Free Software Foundation; either version 2
//' of the License, or (at your option) any later version.
//' This program is distributed in the hope that it will be useful,
//' but WITHOUT ANY WARRANTY; without even the implied warranty of
//' MERCHANTABILITY or FITNESS FOR A PARTICULAR PURPOSE.  See the
//' GNU General Public License for more details.
//' (www.gnu.org/copyleft/gpl.html)
//' You should have received a copy of the GNU General Public License
//' along with this program; if not, write to the Free Software
//' Foundation, Inc., 59 Temple Place - Suite 330, Boston, MA  02111-1307,
//' USA. */

//[[Rcpp::export]]
arma::vec mahaInt(arma::mat & X,  
                  arma::vec & mu,  
                  arma::mat & sigma,
                  const bool isChol = false)
{
  using namespace arma;
  
  // Some sanity checks 
  if(mu.n_elem != sigma.n_cols) Rcpp::stop("The mean vector has a different dimensions from the covariance matrix.");
  if(X.n_cols != sigma.n_cols)  Rcpp::stop("The number of columns of X is different from the dimension of the covariance matrix.");
  
  // Calculate transposed cholesky dec. unless sigma is alread a cholesky dec.
  mat cholDec;
  if( isChol == false ) {
    cholDec = trimatl(chol(sigma).t());
  }
  else{
    cholDec = trimatl(sigma.t()); 
    if(any(cholDec.diag() <= 0.0))  Rcpp::stop("The supplied cholesky decomposition has values <= 0.0 on the main diagonal.");
  }
  
  vec D = cholDec.diag();
  
  vec out(X.n_rows);
  
  // Declaring some private variables
  uint32_t d = X.n_cols;
  uint32_t n = X.n_rows;
  
  vec tmp(d);  
  
  double acc;
  uint32_t icol, irow, ii;  
  
  // For each of the "n" random vectors, forwardsolve the corresponding linear system.
  // Forwardsolve because I'm using the lower triangle Cholesky.
  for(icol = 0; icol < n; icol++)
  {
    
    for(irow = 0; irow < d; irow++)
    {
      acc = 0.0;
      
      for(ii = 0; ii < irow; ii++) acc += tmp.at(ii) * cholDec.at(irow, ii);
      
      tmp.at(irow) = ( X.at(icol, irow) - mu.at(irow) - acc ) / D.at(irow);
    }
    
    out.at(icol) = sum(square(tmp)); 
  }
  
return out;
}

//[[Rcpp::export]]
  arma::vec dmvtInt( arma::mat X, 
                     arma::vec mu, 
                     arma::mat cholDec, 
                     bool log, 
                     double df)
{
  using namespace arma;
  
  unsigned int d = X.n_cols;
  
  vec out = mahaInt(X, mu, cholDec, true);
  
  if( df <= 0.0 ){ // Multivariate normal density OR ...
    
    out = - 0.5 * out - ( (d / 2.0) * std::log(2.0 * M_PI) + sum(arma::log(cholDec.diag())) );
    
  } else { // ... multivariate Student-t density
    
      
      uint32_t ii;  
      uint32_t n = X.n_rows;  
      double logDet = sum(arma::log(cholDec.diag())); 
      double c = lgamma((d + df)/2.0) - (lgamma(df/2.0) + logDet + d/2.0 * std::log(M_PI * df));
      
      for(ii = 0; ii < n; ii++)
      {
        out.at(ii) = c - 0.5 * (df + d) * log1p(out.at(ii)/df);
      }
      
    
  }
  
  if (log == false) out = exp(out);
  
  return( out );
  }


//[[Rcpp::export]]
SEXP dmvtCpp( arma::mat X_,  
              arma::vec mu_,  
              arma::mat sigma_, 
              double df_, 
              bool log_,
              bool isChol_) 
{ 
  
  
  try{
    arma::mat X = X_;
    arma::vec mu = mu_;  
    arma::mat sigma = sigma_; 
    double df = df_; 
    bool log = log_; 
    bool isChol = isChol_;
    
    if( X.n_cols != mu.n_elem ) Rcpp::stop("X.n_cols != mu.n_elem"); 
    if( X.n_cols != sigma.n_cols ) Rcpp::stop("X.n_cols != sigma.n_cols"); 
    if( sigma.n_rows != sigma.n_cols ) Rcpp::stop("sigma.n_rows != sigma.n_cols"); 
    
    // Calculate cholesky dec. unless sigma is alread a cholesky dec.
    arma::mat cholDec;
    if( isChol == false ) {
      cholDec = arma::chol(sigma);
    }
    else{
      cholDec = sigma;
    }
    
    // Dropping the dimensionality of the output vector
    Rcpp::NumericVector Rout = Rcpp::wrap( dmvtInt( X, mu, cholDec, log, df) );
    Rout.attr( "dim" ) = R_NilValue;
    
    return Rout;
    
  } catch( std::exception& __ex__){
    forward_exception_to_r(__ex__);
  } catch(...){
    ::Rf_error( "c++ exception (unknown reason)" );
  }
  return wrap(NA_REAL);
  }


  
  
  
```
