## Supplementary Code 1 for "Inferring differential subcellular localisation in comparative spatial proteomics using BANDLE": fitGP.R

```
## Helper function to fit GPs

fitGP <- function(object = object,
                  fcol = "markers") {

componenthypers <- vector(mode = "list", length(getMarkerClasses(object)))

D <- ncol(object)
K <- length(getMarkerClasses(object, fcol = fcol))

# random grid sampling for starting values
initialvalues <- seq(-3, 3, 2)
init <- matrix(0, length(initialvalues), 3)
for(i in seq_along(initialvalues)){
  init[i,] <- initialvalues[sample.int(length(initialvalues), size = 3, replace = T)]
}

idx <- seq.int(1:D)
tau <- seq.int(1:D)

# LBFGS routine to get hypers
for (j in seq.int(K)) {
  
  exprs <- t(exprs(object[fData(object)[, fcol] == getMarkerClasses(object)[j], idx]))
  
  res <- apply(init, 1,function(z){lbfgs(likelihoodGP,
                                         gradientGP,
                                         vars = z,
                                         invisible = 1,
                                         epsilon = 1e-8,
                                         Xk = exprs,
                                         tau =  seq.int(D),
                                         nk = length(exprs)/D,
                                         D = D)})
  componenthypers[[j]] <- res[[which.min(lapply(res, function(x){max(x$value)}))]]$par
  
}

 .hypers <- matrix(unlist(componenthypers), ncol = 3, byrow = TRUE)

lk <- exp(.hypers[,1])
ak <- exp(2 * .hypers[,2])
sigma <- exp(2 * .hypers[,3])
M <- vector(mode = "list", K)
V <- vector(mode = "list", K)
Var <- vector(mode = "list", K)


  for(j in seq.int(K)){
    Orgdata <- t(exprs(object[fData(object)$markers == getMarkerClasses(object)[j],idx]))
     matplot(Orgdata, col = getStockcol()[j],pch = 19, type = "b", lty = 1, lwd = 1.5, main = paste(getMarkerClasses(object)[j]),
            xlab = "Fraction", ylab = "", cex.main = 2, ylim = c(min(Orgdata) - 0.05, max(Orgdata) + 0.05))
    
    nk <- table(fData(object)$markers)[getMarkerClasses(object)][j]
    S <- matrix(rep(1:length(tau), length(tau)), nrow = length(tau))
    params <- .hypers
    sigmak <- sigma[j]
    a <- ak[j]
    l <- lk[j]
    
    
    covA <- a * exp( - (S - t(S))^ 2 / l)
    R <- diag(1, D) + (nk * covA)/sigmak;
    trenchres <- trenchDetcpp(R[1,])
    Z <- trenchInvcpp(trenchres$v)
    invcov <- diag(1, nk*D)/sigmak - kronecker(matrix(1, nk, nk), Z %*% covA)/sigmak^2
    Kstar <- a*exp(-(matrix(rep(tau, nk * D),  nrow = D, byrow = F) - matrix(rep(tau, nk*D),nrow = D, byrow = T))^2/l)
    Kstarstar <- rep(a+sigmak, length(tau))
    M[[j]] <- Kstar %*% invcov %*% as.vector(Orgdata)
    V[[j]] <- sqrt(diag(diag(Kstarstar, length(tau)) - Kstar %*% invcov %*% t(Kstar)))
    Var[[j]] <- diag(rep(a, length(tau))) - Kstar %*% invcov %*% t(Kstar)
    points(seq_along(tau), M[[j]], col = "black", pch = 19, cex = 1.3, type = "b", lwd = 5, lty = 1)
    arrows(seq_along(tau), M[[j]]-1.96*V[[j]], seq_along(tau), M[[j]]+1.96*V[[j]], length=0.1, angle=90, code=3, col = "black", lwd = 3)
  }

.res <- list(M = M, sigma = sigma, params = params)

return(.res)

}
```
