## Supplementary Code 1 for "Inferring differential subcellular localisation in comparative spatial proteomics using BANDLE": fitGPmatern.R

```
## Helper function to fit GPs

fitGPmatern <- function(object = object,
                          fcol = "markers",
                          materncov = TRUE,
                          nu = 2) {
  
  componenthypers <- vector(mode = "list", length(getMarkerClasses(object)))
  
  D <- ncol(object)
  K <- length(getMarkerClasses(object, fcol = fcol))
  
  # random grid sampling for starting values
  initialvalues <- seq(-5, 0, 0.5)
  init <- matrix(0, length(initialvalues), 3)
  for(i in seq_along(initialvalues)){
    init[i,] <- initialvalues[sample.int(length(initialvalues), size = 3, replace = T)]
  }
  
  idx <- seq.int(1:D)
  tau <- seq.int(1:D)
  
  # LBFGS routine to get hypers
  for (j in seq.int(K)) {
    
    exprs <- t(exprs(object[fData(object)[, fcol] == getMarkerClasses(object)[j], idx]))
    
    res <- apply(init, 1,function(z){lbfgs(likelihoodGPmatern,
                                           gradientGPmatern,
                                           vars = z,
                                           invisible = 1,
                                           epsilon = 1e-6,
                                           Xk = exprs,
                                           tau =  seq.int(D),
                                           nk = length(exprs)/D,
                                           D = D,
                                           materncov = materncov,
                                           nu = nu)})
    componenthypers[[j]] <- res[[which.min(lapply(res, function(x){max(x$value)}))]]$par
    
  }
  
  .hypers <- matrix(unlist(componenthypers), ncol = 3, byrow = TRUE)
  
  rhomaternk <- exp(.hypers[,1])
  amaternk <- exp(.hypers[,2])
  sigma <- exp(2 * .hypers[,3])
  M <- vector(mode = "list", K)
  V <- vector(mode = "list", K)
  Var <- vector(mode = "list", K)
  
  
  for(j in seq.int(K)){
    Orgdata <- t(exprs(object[fData(object)$markers == getMarkerClasses(object)[j],idx]))
     matplot(Orgdata, col = getStockcol()[j],pch = 19, type = "b", lty = 1, lwd = 1.5, main = paste(getMarkerClasses(object)[j]),
            xlab = "Fraction", ylab = "", cex.main = 2, ylim = c(min(Orgdata) - 0.05, max(Orgdata) + 0.05))
    
    nk <- table(fData(object)$markers)[getMarkerClasses(object)][j]
    S <- matrix(rep(1:length(tau), length(tau)), nrow = length(tau))
    params <- .hypers
    sigmak <- sigma[j]
    amatern <- amaternk[j]
    rhomatern <- rhomaternk[j]
    
    
    covA <- matern(nu = nu, a = amatern, rho = rhomatern, tau = seq.int(D), D = D)
    R <- diag(1, D) + (nk * covA)/sigmak;
    trenchres <- trenchDetcpp(R[1,])
    Z <- trenchInvcpp(trenchres$v)
    invcov <- diag(1, nk*D)/sigmak - kronecker(matrix(1, nk, nk), Z %*% covA)/sigmak^2
    Kstar <- do.call(cbind, replicate(nk, covA, simplify=FALSE))
    Kstarstar <- rep(amatern^2 + sigmak, length(tau))
    M[[j]] <- Kstar %*% invcov %*% as.vector(Orgdata)
    V[[j]] <- sqrt(diag(diag(Kstarstar, length(tau)) - Kstar %*% invcov %*% t(Kstar)))
    Var[[j]] <- diag(rep(amatern^2, length(tau))) - Kstar %*% invcov %*% t(Kstar)
    points(seq_along(tau), M[[j]], col = "black", pch = 19, cex = 1.3, type = "b", lwd = 5, lty = 1)
    arrows(seq_along(tau), M[[j]]-1.96*V[[j]], seq_along(tau), M[[j]]+1.96*V[[j]], length=0.1, angle=90, code=3, col = "black", lwd = 3)
  }
  
  .res <- list(M = M, sigma = sigma, params = params)
  
  return(.res)
  
}
```
