## Supplementary Code 1 for "Inferring differential subcellular localisation in comparative spatial proteomics using BANDLE": gradientGP.R

```
gradientGP <- function(Xk, tau, h, nk, D){
  
  grad <- gradientGPcpp(Xk, tau, h, nk, D)
  
  return(unlist(grad))
  
}

gradientGPmatern <- function(Xk, tau, h, nk, D, materncov, nu){
  
  grad <- gradientGPcppmatern(Xk, tau, h, nk, D, nu)
  
  return(unlist(grad))
  
}

posteriorgradientGPmatern <- function(Xk, tau, h, nk, D, materncov, nu, hyppar){
  
  grad <- gradientGPcppmatern(Xk, tau, h, nk, D, nu) + gradientlogprior(h, hyppar)
  
  return(unlist(grad))
  
}

gradientlogprior <- function(h, hyppar){
  
  gradrhoprior <- 3/2 - hyppar[1]*exp(- h[1]/2)/2 
  gradaprior <- hyppar[2]*exp(h[2])
  gradsigmaprior <- -3 + hyppar[3]*exp(h[3])
    
  gradprior <- as.matrix(c(gradrhoprior, gradaprior, gradsigmaprior))
  
  return(gradprior)
}
```
