## Supplementary Code 1 for "Inferring differential subcellular localisation in comparative spatial proteomics using BANDLE": gradienthyp.cpp

```
# include <RcppArmadillo.h>
// [[Rcpp::depends(RcppArmadillo)]]
using namespace Rcpp;

arma::vec gradienthyp(arma::vec Y,
                      arma::mat drv,
                      double nk,
                      double D,
                      arma::mat Z,
                      arma::mat A,
                      double sigmak){
  
  arma::mat B;
  arma::mat J = arma::ones(nk, nk);
  arma::mat eye = arma::eye(D, D);
  arma::mat C;
  
  arma::vec gradienthyp;
  
  arma::mat invZdrv = Z * drv * Z;
  
  //B = (((Xk.t() * covInv) * arma::kron(J, drv)) * (covInv * Xk)) / 2 ;
  B = (((Y.t() * invZdrv) * Y)) / (2 * sigmak * sigmak) ;
  //C = arma::accu(arma::kron(J, drv) % covInv)/2; //use trace is sum of entrywise product
  C = nk * trace(drv)/(2 * sigmak) - nk * arma::accu((eye - Z) % drv)/(2 * sigmak);
  
  //gradienthyp = B - C;  
  gradienthyp = B - C;
  
  return(gradienthyp); 
}

arma::vec gradientNoise(arma::vec Y,
                        double drv,
                        double nk,
                        double sigmak,
                        arma::mat Z,
                        double D,
                        double Y2){
  
  arma::mat B;
  arma::mat J = arma::ones(nk, nk);
  arma::mat C;
  arma::mat eye = arma::eye(D,D);
  arma::vec gradientNoise;
  
  
  //B = drv * ((Xk.t() * covInv) * (covInv * Xk))/2 ;
  B = drv * Y2/(2 * sigmak * sigmak) + drv * Y.t() * ( Z * Z - eye) * Y /(nk * sigmak * sigmak * 2);
  //C = drv * trace(covInv)/2; //use trace is sum of entrywise product
  C = drv * ((nk - 1) * D / sigmak + trace(Z)/sigmak)/2;
  
  //gradientNoise = B - C;  
  gradientNoise = B - C;
  
  return(gradientNoise); 
}


List trenchDetcpp(arma::vec c) {
  
  int N = c.size();
  int i;
  arma::vec xi(N-1);
  arma::vec z(N-1);
  arma::vec v(N);
  double beta = 1;
  double alpha;
  double l = 0;
  double logdet;
  
  xi = c.subvec(1, N-1)/c(0);
  z(0) = - xi(0);
  alpha = -xi(0);
  int K = xi.size(); 
  
  for(i = 0; i < (K - 1); i++){
    beta = (1 - pow(alpha, 2)) * beta ;
    l = l + log(beta) ;
    if(i == 0){
      alpha = - (xi(i + 1) + xi(0) * z(0)) / beta ;
      z(0) = z(0) + alpha * z(0) ;
    }else{
      alpha = - (xi(i + 1) + dot(flipud(xi.subvec(0, i)), z.subvec(0, i)) ) / beta ;
      z.subvec(0, i) = z.subvec(0, i) + alpha * flipud(z.subvec(0, i)) ;
    }
    
    z(i+1) = alpha  ;
  }
  
  beta = (1 - pow(alpha, 2)) * beta ;
  l = l + log(beta) ;
  
  logdet = l + N * log(c(0));
  
  v(N-1) = 1 / ((1 + dot(xi, z)) * c(0)) ;
  v.subvec(0,N-2) = v(N-1) * flipud(z.subvec(0, N - 2));
  
  return List::create(Rcpp::Named("logdet") = logdet,
                      Rcpp::Named("z") = z,
                      Rcpp::Named("v") = v);
}


arma::mat trenchInvcpp(arma::vec v) {
  
  int N = v.size();
  int i;
  int j;
  arma::mat C(N, N);
  arma::mat trenchInv;
  
  
  C.row(0) = flipud(v).t();
  C.col(0) = flipud(v);
  C.row(N - 1) = v.t();
  C.col(N - 1) = v;
  for(i = 1; i < floor( (N - 1) / 2 ) + 1; i++){
    for(j = 1; j < N - i; j++){
      C(i, j) = C(i - 1, j - 1) + (v(N - 1 - j) * v(N - 1 - i) - v(i - 1) * v(j - 1)) / v(N - 1) ;
      C(j, i) = C(i, j);
      C(N - 1 - i , N - 1 - j ) = C(i, j);
      C(N - 1 - j , N - 1 - i ) = C(i, j);
    }
  } 
  
  trenchInv = C;
  return(trenchInv) ; 
  
}


// [[Rcpp::export]]
arma::vec gradientGPcpp(arma::mat Xk,
                       arma::vec tau,
                       arma::vec h,
                       int nk,
                       int D
){
  
  //terms are made unconstrained
  double sigmak; sigmak = exp(2 * h(2));
  double a; a = exp(2 * h(1));
  double l; l = exp(h(0));
  double negloglike;
  double logdet;
  arma::mat Z(D,D);
  arma::mat S(D,D); S = arma::zeros(D,D);
  arma::mat A(D,D);
  arma::mat R(D,D);
  arma::mat covInv;
  arma::mat drvl;
  arma::mat drva;
  double drvsigma;
  arma::vec temp;
  arma::vec v;
  arma::vec grdl;
  arma::vec grda;
  arma::vec grdsigma;
  arma::vec grad(3);
  arma::vec Y(D);
  double Y2;
  List trenchres = List::create(Rcpp::Named("logdet"),
                                Rcpp::Named("z"),
                                Rcpp::Named("v"));
  
  S = S.each_col() + tau;
  A = a * exp(- pow(S - S.t(), 2)/l);
  R = arma::eye(D,D) + (nk * A)/sigmak;
  
  trenchres = trenchDetcpp(R.row(0).t());
  logdet = trenchres["logdet"];
  v = as<arma::vec>(trenchres["v"]);
  
  
  if(R_IsNA(logdet)){
    negloglike = R_NegInf;
    grad.row(0) = negloglike;
    return(grad);
  }
  
  Z = trenchInvcpp(v);
  //covInv = covInvcpp(Z, A, sigmak, nk, D);
    
  drvl = A % (pow(S - S.t(), 2)/l);
  drva = 2 * A ;
  drvsigma = 2 * sigmak ;
  
  Y2 = arma::accu(pow(Xk, 2));  
  Y = sum(Xk, 1);
  
  grdl = gradienthyp(Y, drvl, nk, D, Z, A, sigmak);
  grda = gradienthyp(Y, drva, nk, D, Z, A, sigmak);
  grdsigma = gradientNoise(Y, drvsigma, nk, sigmak, Z , D, Y2);
    
 //return gradient of negative log like
 grad(0) = -grdl(0);
 grad(1) = -grda(0);
 grad(2) = -grdsigma(0);
 
 return(grad);
}
```
