## Supplementary Code 1 for "Inferring differential subcellular localisation in comparative spatial proteomics using BANDLE": hamiltonianGP.R

```
# Hamiltonian Monte-Carlo updates for GP hyperparameters


hamiltonianGP <- function(inith,
                         X,
                         tau,
                         nk,
                         D,
                         steps = 30,
                         stepsize = c(0.01,0.02),
                         alpha = 0.01,
                         niter = 100,
                         hypMean = c(0,0,0),
                         hypSd = c(1,1,1)
                        ){
    #number of leapfrog steps
    L <- steps
    h <- matrix(0, ncol = niter + 1, nrow = length(inith))
    h[, 1] <- inith
    ar <- 0
    m <- c(3, 2, 200)
    
    #number of samples
    for(i in seq.int(niter)){
      
     #current hyperparameters
     x <- h[,i]
     currentx <- x
     #sample momenta value using residual momentum
     if(i == 1){
       p <- rnorm(3, mean = rep(0,3), sd = 1/m)
     }else{
       p <- alpha * residualp + sqrt(1 - alpha ^ 2) * rnorm(3, mean = rep(0,3), sd = 1/m)
     }
     currentp <- p
     delta <- runif(1, min = stepsize[1], max = stepsize[2])
    
     #Leapfrog algorithm
     
     leapres <- LeapfrogGPcpp(X, tau, p, x, m, nk, D, L, delta)
     #reverse momenta for symmetric proposal
     p <- -leapres$p
     x <- t(leapres$x)
     
     #evalution of energies at start and end
     currentU <- -likelihoodGPcpp(X, tau, currentx, nk, D) + sum(dnorm(currentx, mean = hypMean, sd = hypSd, log = TRUE))
     proposedU <- -likelihoodGPcpp(X, tau, x, nk, D) + sum(dnorm(x, mean = hypMean, sd = hypSd, log = TRUE))
     currentK <-  sum(currentp ^ 2 / (2 * m))
     proposedK <- sum(p ^ 2 / (2 * m))
    
     #log metropolis ratio
     logmhratio <- proposedU - currentU + proposedK - currentK
    
     if(is.na(logmhratio)){
      h[, i] <- currentx
      print("Error in Metropolis ratio suggestion reduce stepsize")
      return(list(h=h, ar=ar))
     }

     if(logmhratio > log(runif(1, 0, 1))){
      h[, i + 1] <- x
      ar <- ar + 1
     }else{
      h[, i + 1] <- currentx
     }
    
     #allow momentum to continue
     residualp <- t(p)
    }
    ar <- ar/niter
  return(list(h = h, ar = ar))    

}

hamiltonianGPmatern <- function(inith,
                                X,
                                lambda,
                                nu,
                                tau,
                                nk,
                                D,
                                steps = 30,
                                stepsize = c(0.01,0.02),
                                alpha = 0.01,
                                niter = 100
){
  #number of leapfrog steps
  L <- steps
  h <- matrix(0, ncol = niter + 1, nrow = length(inith))
  h[, 1] <- inith
  ar <- 0
  m <- c(3, 1, 200)
  
  #number of samples
  for(i in seq.int(niter)){
    
    #current hyperparameters
    x <- h[,i]
    currentx <- x
    #sample momenta value using residual momentum
    if(i == 1){
      p <- rnorm(3, mean = rep(0,3), sd = 1/m)
    }else{
      p <- alpha * residualp + sqrt(1 - alpha ^ 2) * rnorm(3, mean = rep(0,3), sd = 1/m)
    }
    currentp <- p
    delta <- runif(1, min = stepsize[1], max = stepsize[2])
    
    #Leapfrog algorithm
    
    leapres <- LeapfrogGPcppPC(X, lambda, tau, p, x, m, nk, D, L, delta, nu)
    #reverse momenta for symmetric proposal
    p <- -leapres$p
    x <- t(leapres$x)
    
    #evalution of energies at start and end
    currentU <- -likelihoodGPcpp(X, tau, currentx, nk, D, materncov = TRUE, nu) + PCrhomvar(rho = exp(currentx[1]), a = exp(currentx[2]), lambda1 = lambda[1],
                                                                      lambda2 = lambda[2], log = TRUE) +  
      Gumbel(exp(-2 * currentx[3]), lambda = lambda[3], log = TRUE)
    proposedU <- -likelihoodGPcpp(X, tau, x, nk, D, materncov = TRUE, nu) + PCrhomvar(rho = exp(x[1]), a = exp(x[2]), lambda1 = lambda[1],
                                                                lambda2 = lambda[2], log = TRUE) +  
      Gumbel(exp(-2 * x[3]), lambda = lambda[3], log = TRUE)
    currentK <-  sum(currentp ^ 2 / (2 * m))
    proposedK <- sum(p ^ 2 / (2 * m))
    
    #log metropolis ratio
    logmhratio <- proposedU - currentU + proposedK - currentK
    
    if(is.na(logmhratio)){
      h[, i] <- currentx
      print("Error in Metropolis ratio suggestion reduce stepsize")
      return(list(h=h, ar=ar))
    }
    
    if(logmhratio > log(runif(1, 0, 1))){
      h[, i + 1] <- x
      ar <- ar + 1
    }else{
      h[, i + 1] <- currentx
    }
    
    #allow momentum to continue
    residualp <- t(p)
  }
  ar <- ar/niter
  return(list(h = h, ar = ar))    
  
}
    
```
