## Supplementary Code 1 for "Inferring differential subcellular localisation in comparative spatial proteomics using BANDLE": lfgbsGP.R

```
# LBFGS optimisation procedure for GPs hyperparmeters

# NTS minimising negative log likelihood, since lbfgs minimises
require(MASS)
h <- c(1,0.1, -0.4)
sigmak <- exp(2 * h[3])
a <- exp(2 * h[2])
l <- exp(h[1])
tau = c(1,2,3,4,5)
S <- matrix(rep(1:length(tau), length(tau)), nrow = length(tau))
A <- a * exp( - (S - t(S))^ 2 / l)
X <- mvrnorm(1, rep(0,100), kronecker(matrix(1,20,20), A) + sigmak*diag(1,100))
X <- matrix(X, nrow = 5, byrow = TRUE)
par(mfrow = c(2,2))
X <- exprs(hyperLOPIT2015[fData(hyperLOPIT2015)$markers == getMarkerClasses(hyperLOPIT2015)[10],])
matplot(t(X), pch = 19, col = "black")


par(mfrow = c(3,3))

hlnorm <- normalise(hyperLOPIT2015, method = "quantiles")
hlnorm <- normalise(hyperLOPIT2015, method = "center.mean")
X <- exprs(hlnorm[fData(hlnorm)$markers == getMarkerClasses(hlnorm)[4],idx])
idx <- c(1,11,2,12,3,13,4,14,5,15,6,16,7,17,8,18,9,19,10,20)
X <- X[,idx]
matplot(t(X), pch = 19, col = "black", ylim = c(-0.2, 0.5))


init <- seq(-3,3, 2)
initz <- matrix(0,length(init),3)

for(i in seq_along(init)){
  initz[i,] <- init[sample.int(length(init), size = 3, replace = T)]
}

tau <- seq_along(idx)
nk <- length(X)/max(idx)
D <- 20

res <- apply(initz, 1,function(z){lbfgs(likelihoodGP,
      gradientGP,
      vars = z,
      invisible=1,
      epsilon = 1e-8,
      Xk = t(X),
      tau = seq_along(idx),
      nk = length(X)/max(idx),
      D = 20)})

bestparams <- res[[which.min(lapply(res, function(x){max(x$value)}))]]

S <- matrix(rep(1:length(tau),length(tau)), nrow = length(tau))
sigmak <- exp(2 * bestparams$par[3])
a <- exp(2 * bestparams$par[2])
l <- exp(bestparams$par[1])


covA <- a * exp( - (S - t(S))^ 2 / l)
cov <- kronecker(matrix(1,nk, nk), covA) + sigmak*diag(1, nk*D)
Kstar <- a*exp(-(matrix(rep(tau, nk*D),  nrow = D, byrow = F) - matrix(rep(tau, nk*D),nrow = D, byrow = T))^2/l)
Kstarstar <- rep(a+sigmak, length(tau))


Y <- matrix(X, nrow = 20 , byrow = TRUE)
matplot(t(Y), pch = 19, col = "black", ylim = c(min(X)-1,max(X)+1))

M <- Kstar %*% solve(cov) %*% as.vector(t(X))
V <- sqrt(diag(diag(Kstarstar, length(tau)) - Kstar %*% solve(cov) %*% t(Kstar)))
points(seq_along(tau), M, col = "red", pch = 19, cex = 1.2, type = "b")
arrows(seq_along(tau), M-1.96*V, seq_along(tau), M+2*V, length=0.1, angle=90, code=3, col = "red", lwd = 2)

Orgparamstest <- vector(mode = "list", length(getMarkerClasses(hlnorm)))

system.time(for(j in seq.int(length(getMarkerClasses(hlnorm)))){
 exprs <- t(exprs(hlnorm[fData(hlnorm)$markers == getMarkerClasses(hlnorm)[j],idx]))
 
res <- apply(initz, 1,function(z){lbfgs(likelihoodGP,
                                         gradientGP,
                                         vars = z,
                                         invisible=1,
                                         epsilon = 1e-8,
                                         Xk = exprs,
                                         tau =  seq.int(D),
                                         nk = length(exprs)/D,
                                         D = 20)})
 Orgparamstest[[j]] <- res[[which.min(lapply(res, function(x){max(x$value)}))]]
 
})


#save(Orgparams, file = "lbfgsOrg.rda")

getallpars <- sapply(Orgparams, function(x) {x$par})
colnames(getallpars) <- getMarkerClasses(hlnorm)
rownames(getallpars) <- c("Memory", "Amplitude", "Noise")

par(mfrow = c(4,4))
for(j in 1:14){
Orgdata <- t(exprs(hlnorm[fData(hlnorm)$markers == getMarkerClasses(hlnorm)[j],idx]))
matplot(Orgdata, col = getStockcol()[j],pch = 19, type = "b", lty = 1, lwd = 1.5, main = paste(getMarkerClasses(hlnorm)[j]),
        xlab = "Fraction", ylab = "", cex.main = 2)

nk <- table(fData(hlnorm)$markers)[j]
S <- matrix(rep(1:length(tau),length(tau)), nrow = length(tau))
params <- Orgparams[[j]]$par
sigmak <- exp(2 * params[3])
a <- exp(2 * params[2])
l <- exp(params[1])


covA <- a * exp( - (S - t(S))^ 2 / l)
R <- diag(1,D) + (nk * covA)/sigmak;
trenchres <- trenchDetcpp(R[1,])
Z <- trenchInvcpp(trenchres$v)
invcov <- diag(1, nk*D)/sigmak - kronecker(matrix(1, nk, nk), Z %*% covA)/sigmak^2
Kstar <- a*exp(-(matrix(rep(tau, nk*D),  nrow = D, byrow = F) - matrix(rep(tau, nk*D),nrow = D, byrow = T))^2/l)
Kstarstar <- rep(a+sigmak, length(tau))
M <- Kstar %*% invcov %*% as.vector(Orgdata)
V <- sqrt(diag(diag(Kstarstar, length(tau)) - Kstar %*% invcov %*% t(Kstar)))
points(seq_along(tau), M, col = "black", pch = 19, cex = 1.3, type = "b", lwd = 5, lty = 1)
arrows(seq_along(tau), M-1.96*V, seq_along(tau), M+1.96*V, length=0.1, angle=90, code=3, col = "black", lwd = 3)
}
```
