## Supplementary Code 1 for "Inferring differential subcellular localisation in comparative spatial proteomics using BANDLE": likelihoodGP.R

```
likelihoodGP <- function(Xk, tau, h, nk, D){
  
  negloglike <- likelihoodGPcpp(Xk, tau, h, nk, D)
  
  return(negloglike)
}

likelihoodGPmatern <- function(Xk, tau, h, nk, D, materncov, nu){
  
  negloglike <- likelihoodGPcpp(Xk, tau, h, nk, D, materncov, nu)
  
  return(negloglike)
}

posteriorGPmatern <- function(Xk, tau, h, nk, D, materncov, nu, hyppar){
  
  negloglike <- likelihoodGPcpp(Xk, tau, h, nk, D, materncov, nu) - PCrhomvar(rho = exp(h[1]), a = exp(h[2]), lambda1 = hyppar[1],
                                                                              lambda2 = hyppar[2], log = TRUE) - Gumbel(1/exp(2*h[3]), lambda = hyppar[3], log = TRUE)
  
  return(negloglike)
}

Gumbel <- function(x,
                   lambda,
                   log = TRUE){
  
  if (log == FALSE) {
    gumbel <- (lambda/2) * x^{-3/2} * exp(-lambda * x^{-1/2})
  } else {
    gumbel <- log(lambda/2) - 3*log(x)/2 - lambda * x^{-1/2}
  }
  
  return(gumbel)
}


PCrhomvar <- function(rho,
                      a,
                      lambda1,
                      lambda2,
                      log = TRUE){
  
  if (log == FALSE) {
    res <- (lambda1 * lambda2/2) * rho^{- 3/2} * exp(-lambda1 * rho^{-1/2} - lambda2 * a)
  } else {
    res <- log(lambda1 * lambda2/2) - 3 * log(rho)/2 - lambda1 * rho^{-1/2} - lambda2 * a
  }
  
  return(res) 
  
}
```
