## Supplementary Code 1 for "Inferring differential subcellular localisation in comparative spatial proteomics using BANDLE": metropolisGP.R

```
# Metroplis-Hastings for GP hyperparameters with normal prior


metropolisGP <- function(inith,
                         X,
                         tau,
                         nk,
                         D,
                         niter,
                         hypMean = c(0,0,0),
                         hypSd = c(1,1,1)
                         ){
  
  
  h <- matrix(0, ncol = niter + 1, nrow = length(inith))
  h[, 1] <- inith
  ar <- 0 
  
  for(i in seq.int(niter)){
   xi <- rnorm(length(inith), mean = 0, sd = 0.1) # sample random walk steps
   oldHypers <- h[, i]
   proposedHypers <- h[, i] + xi #random walk proposal
   #compute metropolis ratio, likelihoodGPcpp return negative logliklihood
   mhratio <- -likelihoodGPcpp(X, tau, proposedHypers, nk, D) + likelihoodGPcpp(X, tau, oldHypers, nk, D) + 
     sum(dnorm(proposedHypers, mean = hypMean, sd = hypSd, log = TRUE)) - sum(dnorm(oldHypers, mean = hypMean,
                                                                                    sd = hypSd, log = TRUE))
  
   if(mhratio > log(runif(1, 0, 1))){
    h[, i + 1] <- proposedHypers
    ar <- ar + 1
   }else{
    h[, i + 1] <- oldHypers
   }
   
  }
  ar <- ar/niter
  
  return(list(h = h, ar = ar))
  
}


metropolisGPmatern <- function(inith,
                               X,
                               tau,
                               nk,
                               D,
                               niter,
                               nu = 2,
                               hyppar = c(1, 1, 1),
                               propsd = c(0.3,0.1,0.1)
){
  
  
  h <- matrix(0, ncol = niter + 1, nrow = length(inith))
  h[, 1] <- inith
  ar <- 0 

  propsd <- propsd
  for(i in seq.int(niter)){
    # repeat proposal so proposals are positive, (truncated sampler)
    repeat {
    xi <- rnorm(length(inith), mean = 0, sd = propsd) # sample random walk steps
    
    oldHypers <- h[, i]
    proposedHypers <- h[, i] + xi #random walk proposal
    if ((all((proposedHypers > 0)))){
      break
    }
    }
    #compute metropolis ratio, likelihoodGPcpp return negative logliklihood, densities from PC priors + truncated sampler correction
    mhratiolike <- -likelihoodGPmatern(X, tau, proposedHypers, nk, D, materncov = TRUE, nu = nu) +
                likelihoodGPmatern(X, tau, oldHypers, nk, D, materncov = TRUE, nu = nu)
    mhratioprior <- PCrhomvar(rho = proposedHypers[1], a = proposedHypers[2], lambda1 = hyppar[1],
                              lambda2 = hyppar[2], log = TRUE) + 
                    Gumbel(1/proposedHypers[3], lambda = hyppar[3], log = TRUE) + sum(pnorm(oldHypers, mean = 0,
                                                                                      sd = propsd, log = TRUE)) - 
                    (PCrhomvar(rho = oldHypers[1], a = oldHypers[2], lambda1 = hyppar[1], 
                               lambda2 = hyppar[2], log = TRUE) + Gumbel(1/oldHypers[3], lambda = hyppar[3], log = TRUE) + 
                       sum(pnorm(proposedHypers, mean = 0, sd = propsd, log = TRUE)))
    mhratio <- mhratiolike + mhratioprior
    
    if(mhratio > log(runif(1, 0, 1))){
      h[, i + 1] <- proposedHypers
      ar <- ar + 1
    }else{
      h[, i + 1] <- oldHypers
    }
    
  }
  ar <- ar/niter
  
  # returns parameter values on true scale note that variance rather than precision is returned
  return(list(h = h, ar = ar))
  
}

Gumbel <- function(x,
                   lambda,
                   log = TRUE){
  
  if (log == FALSE) {
  gumbel <- (lambda/2) * x^{-3/2} * exp(-lambda * x^{-1/2})
  } else {
  gumbel <- log(lambda/2) - 3*log(x)/2 - lambda * x^{-1/2}
  }
  
  return(gumbel)
}


PCrhomvar <- function(rho,
                      a,
                      lambda1,
                      lambda2,
                      log = TRUE){
  
  if (log == FALSE) {
  res <- (lambda1 * lambda2/2) * rho^{- 3/2} * exp(-lambda1 * rho^{-1/2} - lambda2 * a)
  } else {
  res <- log(lambda1 * lambda2/2) - 3 * log(rho)/2 - lambda1 * rho^{-1/2} - lambda2 * a
  }
  
  return(res) 
  
}
```
