## Supplementary Code 1 for "Inferring differential subcellular localisation in comparative spatial proteomics using BANDLE": mixGP.R

```
#supervised mixture of gaussian process regression models

mixGP <- function(object,
                  hypLearn = "MH",
                  numIter = 1000,
                  burnin = 100L,
                  thin = 5L,
                  alpha = 1,
                  u = 2,
                  v = 10,
                  hypIter = 20,
                  hypMean = c(0,0,0),
                  hypSd = c(1,1,1),
                  seed = NULL){
  
  if (is.null(seed)) {
    seed <- sample(.Machine$integer.max, 1)
  }
  .seed <- as.integer(seed)  
  set.seed(.seed)
  
  stopifnot(length(hypMean)==3)
  stopifnot(length(hypSd)==3)
  stopifnot(sum(hypSd > 0)==3)
  
  ## on the fly number of samples to be retained
  retained <- seq.int(burnin + 1L, numIter , thin)
  
  hlnorm <- normalise(object, method = "quantiles")
  hlnorm <- normalise(object, method = "center.mean")
  D <- ncol(exprs(hlnorm))
  M <- colMeans(exprs(hlnorm))
  V <- cov(exprs(object))
  K <- length(getMarkerClasses(hlnorm))
  S <- matrix(rep(1:D,D), nrow = D)
  
  componenthypers <- vector(mode = "list", length(getMarkerClasses(hlnorm)))
  
  #random grid sampling for starting values
  initialvalues <- seq(-3,3, 2)
  init <- matrix(0, length(initialvalues), 3)
  for(i in seq_along(initialvalues)){
    init[i,] <- initialvalues[sample.int(length(initialvalues), size = 3, replace = T)]
  }
  
  idx <- seq.int(1:D)
  #LBFGS routine to get hypers
  for(j in seq.int(length(getMarkerClasses(hlnorm)))){
    
    exprs <- t(exprs(hlnorm[fData(hlnorm)$markers == getMarkerClasses(hlnorm)[j],idx]))
    
    res <- apply(init, 1,function(z){lbfgs(likelihoodGP,
                                           gradientGP,
                                           vars = z,
                                           invisible=1,
                                           epsilon = 1e-8,
                                           Xk = exprs,
                                           tau =  seq.int(D),
                                           nk = length(exprs)/D,
                                           D = D)})
    componenthypers[[j]] <- res[[which.min(lapply(res, function(x){max(x$value)}))]]$par
    
  }
  
  .hypers <- matrix(unlist(componenthypers), ncol = 3, byrow = TRUE)
  
  #seperate data into known and unknown
  unknownData <- unknownMSnSet(hlnorm)
  exprsUnknown <- exprs(unknownData)
  exprsKnown <- exprs(markerMSnSet(hlnorm))
  allocKnown <- seq.int(1:K)[fData(markerMSnSet(hlnorm))$markers]
  numProtein <- nrow(unknownData)
  
  #some storage
  alloc <- matrix(0, nrow = numProtein, ncol = numIter)
  allocOut <- matrix(0, nrow = numProtein, ncol = numIter)
  outlierprob <- matrix(0, nrow = numProtein, ncol = numIter)
  weights <- matrix(0, nrow = K, ncol = numIter)
  epsilons <- matrix(0, nrow = 1, ncol = numIter)
  allocOutprob <- array(0, c(nrow(unknownData), numIter, 2))
  allocprob <- array(0, c(nrow(unknownData), numIter, K))
  loglikelihoods <- matrix(0, nrow = numProtein, ncol = K)
  
  #random allocation of unknown Proteins
  alloc[, 1] <- sample.int(n = K, size = numProtein, replace = TRUE)
  
  #number of proteins allocated to each component
  nkknown <- table(getMarkers(hlnorm, verbose = FALSE))[getMarkerClasses(hlnorm)]
  
  #number of proteins allocated to each component 
  nkknown <- nkknown
  
  #number initial allocated to outlier component
  allocOut[, 1] <- 0
  
  sampleGPMean <- vector(mode = "list", length = K)
  centereddata <- vector(mode = "list", length = K)
  hypers <- vector(mode = "list", length = numIter)
  
  hypers[[1]] <- .hypers

  for(t in 2:numIter){
    
    
    alloctemp <- alloc[, t - 1]
    outlier <- allocOut[, t - 1]
    
    nk <- nkknown + tabulate(alloctemp[outlier!=0], nbins = K)
    
    if((t %% 50) ==0){
      print(t)
    }
    
    centereddata <- centeredData(Xknown = exprsKnown,
            BX = allocKnown,
            Xunknown = exprsUnknown, 
            BXun = alloctemp*outlier,
            hypers = matrix(unlist(componenthypers), ncol = 3, byrow = TRUE), 
            nk = nk, tau = seq.int(D), D = D, K)

    #sample weights from dirichlet by normalising gammas
    currentweights <- t(sampleDirichlet(K, alpha/K + nk))
    
    #extract noise component from hypers
    sigmak <- exp(2 * hypers[[t-1]][,3])
    loglikelihoods <- comploglike(centereddata, sqrt(sigmak))
    
    conditionalAlloc <- t(apply(loglikelihoods,1, function(x) x + log(currentweights)))
    cc <- apply(conditionalAlloc, 1, max)
    conditionalAlloc <- conditionalAlloc - cc #correct for underflow
    allocprob[,t,] <- exp(conditionalAlloc)/rowSums(exp(conditionalAlloc))
    alloctemp <- apply(allocprob[,t,], 1, function(x) sample.int(n = K, size = 1, replace = F, prob = x))
    
    #sample epsilon
    tau1 <- sum(allocOut[, t] == 1) + nrow(exprsKnown)
    tau2 <- sum(allocOut[, t] == 0)
    epsilon <- rbeta(n = 1, shape1 = u + tau1, shape2 = v + tau2)
    
    #sample outlier allocations
    allocOutprob[, t, 1] <- log(1 - epsilon) + loglikelihoods[cbind(1:length(alloctemp), alloctemp)] 
    allocOutprob[, t, 2] <- log(epsilon) + dmvtCpp(exprsUnknown, mu_ = M, sigma_ = V, df_ = 4,
                                                   log_ = TRUE, isChol_ = F)
    c <- apply(allocOutprob[,t,], 1, max) 
    allocOutprob[ , t, ] <- allocOutprob[,t,] - c
    allocOutprob[ , t, ] <- exp(allocOutprob[, t, ])/rowSums(exp(allocOutprob[, t, ]))
    outlier <- apply(allocOutprob[,t,], 1, function(z){
                                                  sample(x = c(1,0), 1, prob = z)}
                                                  ) # reversed sample so 2nd entry is prob of 0

    #update hypers
    if((t %% hypIter) == 0){
     if(hypLearn == "LBFGS"){
     }else if(hypLearn == "MH"){
      
       nk <- nkknown + tabulate(alloctemp[outlier!=0], nbins = K)
       for(j in seq.int(K)){
         Y <- makeComponent(X = exprsKnown, BX = allocKnown, Y = exprsUnknown, BY = alloctemp * outlier, j = j)
         componenthypers[[j]] <- metropolisGP(inith = componenthypers[[j]], 
                                             X = t(Y), tau = seq.int(D), nk = nk[j], D = D, niter = 1,
                                             hypMean = hypMean, hypSd = hypSd)$h[,2]
       }
     }else if(hypLearn == "HMC"){
       print("HMC step performed")
       nk <- nkknown + tabulate(alloctemp[outlier!=0], nbins = K)
       for(j in seq.int(K)){
         Y <- makeComponent(X = exprsKnown, BX = allocKnown, Y = exprsUnknown, BY = alloctemp * outlier, j = j)
         componenthypers[[j]] <- hamiltonianGP(componenthypers[[j]],
                                               tau = seq.int(D), X = t(Y), nk = nk[j],
                                               D = D, niter = 1, stepsize = c(0.13,0.13), steps = 10,
                                               hypMean = hypMean, hypSd = hypSd)$h[,2]
       }
     }
    }
    .hypers <- matrix(unlist(componenthypers), ncol = 3, byrow = TRUE)
    
    
    hypers[[t]] <- .hypers
    weights[,t] <- currentweights
    alloc[,t] <- alloctemp
    epsilons[t] <- epsilon
    allocOut[,t] <- outlier
    
  }
  
  
  
  return(list(allocOutprob = allocOutprob[, retained, ],
              allocOut = allocOut[, retained], 
              allocprob = allocprob[, retained, ],
              alloc = alloc[, retained], 
              hypers = hypers[retained],
              weights = weights[, retained],
              epsilons = epsilons[retained]))
  
  
}
```
