## Supplementary Code 1 for "Inferring differential subcellular localisation in comparative spatial proteomics using BANDLE": MRimplementation.R

```
robustMahalanobis <- function(delta) {
  
  # Compute robust covariance
  rCov <- covMcd(delta)
  
  sqMan <- matrix(NA, nrow = nrow(delta), ncol = 1)
  for (i in seq.int(nrow(sqMan))) {
    sqMan[i] <- (delta[i,] - colMeans(delta)) %*% solve(rCov$cov) %*% (delta[i,] - colMeans(delta))
  }
  
  return(sqMan)
}  

reprodScore <- function(x, y) {
  
  Rscore <- matrix(NA, nrow = nrow(x), ncol = 1)
  for (i in seq.int(nrow(x))){
    Rscore[i] <- cor(x[i,], y[i,], method = c("pearson"))
  }
  
  return(Rscore)
}


require(robustbase)

mrMethod <- function(mydata) {
  
  test1 <- mydata[[1]]
  test1b <- mydata[[2]]
  test1c <- mydata[[3]]
  
  test2 <- mydata[[4]]
  test2b <- mydata[[5]]
  test2c <- mydata[[6]]
  
  # Compute delta matrices
  delta1 <- exprs(test1) - exprs(test2)
  delta2 <- exprs(test1b) - exprs(test2b)
  delta3 <- exprs(test1c) - exprs(test2c)
  
  # Compute robust mahalanobis distance
  sqMan1 <- robustMahalanobis(delta = delta1)
  sqMan2 <- robustMahalanobis(delta = delta2)
  sqMan3 <- robustMahalanobis(delta = delta3)
  
  
  # Under null distance is chi squared distributed
  df <- ncol(delta1)
  pchi1 <- pchisq(sqMan1, df = df, lower.tail = FALSE)
  pchi2 <- pchisq(sqMan2, df = df, lower.tail = FALSE)
  pchi3 <- pchisq(sqMan3, df = df, lower.tail = FALSE)
  
  
  # As stated take the maximum p-value from each test
  summarisedPvalue <- apply(cbind(pchi1, pchi2, pchi3), 1 ,max)
  
  # Adjust for multiple testing of the cubed pValue
  summarisedPvalue <- p.adjust(summarisedPvalue^3, method = "BH")
  
  # add Names
  names(summarisedPvalue) <- rownames(test1)
  
  # Compute M-score
  Mscore <- -log10(summarisedPvalue)
  
  # compute reproducibility sore
  Rscore1 <- reprodScore(x = delta1, y = delta2)
  Rscore2 <- reprodScore(x = delta2, y = delta3)
  Rscore3 <- reprodScore(x = delta3, y = delta1)
  
  # take smallest score
  Rscore <- apply(cbind(Rscore1, Rscore2, Rscore3), 1,function(x) min(x, na.rm = TRUE))
  
  # add Names
  names(Rscore) <- names(Mscore) <- rownames(test1)
  
  plot(Mscore, Rscore)
  
  ## Compute ROC 
  MRscore <- ((Rscore - mean(Rscore))/sd(Rscore)) * ((Mscore - mean(Mscore[!is.infinite(Mscore)]))/sd(Mscore[!is.infinite(Mscore)]))
  names(MRscore) <- rownames(test1)
  MRscore[is.infinite(MRscore)] <- max(MRscore[!is.infinite(MRscore)])
  
  return(list(Rscore = Rscore, Mscore = Mscore, MRscore = MRscore))
}
```
