## Supplementary Code 1 for "Inferring differential subcellular localisation in comparative spatial proteomics using BANDLE": MRmethod.R

```
##

robustMahalanobis <- function(delta) {
  
  # Compute robust covariance
  rCov <- covMcd(delta)

  sqMan <- matrix(NA, nrow = nrow(delta), ncol = 1)
  for (i in seq.int(nrow(sqMan))) {
    sqMan[i] <- (delta[i,] - colMeans(delta)) %*% solve(rCov$cov) %*% (delta[i,] - colMeans(delta))
  }
  
  return(sqMan)
}  

reprodScore <- function(x, y) {
  
  Rscore <- matrix(NA, nrow = nrow(x), ncol = 1)
  for (i in seq.int(nrow(x))){
    Rscore[i] <- cor(x[i,], y[i,], method = c("pearson"))
  }

  return(Rscore)
}


require(robustbase)

mrMethod <- function(mydata) {
  
  test1 <- mydata[[1]]
  test1b <- mydata[[2]]
  test1c <- mydata[[3]]
  
  test2 <- mydata[[4]]
  test2b <- mydata[[5]]
  test2c <- mydata[[6]]

# Compute delta matrices
delta1 <- exprs(test1) - exprs(test2)
delta2 <- exprs(test1b) - exprs(test2b)
delta3 <- exprs(test1c) - exprs(test2c)

# Compute robust mahalanobis distance
sqMan1 <- robustMahalanobis(delta = delta1)
sqMan2 <- robustMahalanobis(delta = delta2)
sqMan3 <- robustMahalanobis(delta = delta3)


# Under null distance is chi squared distributed
df <- ncol(delta1)
pchi1 <- pchisq(sqMan1, df = df, lower.tail = FALSE)
pchi2 <- pchisq(sqMan2, df = df, lower.tail = FALSE)
pchi3 <- pchisq(sqMan3, df = df, lower.tail = FALSE)


# As stated take the maximum p-value from each test
summarisedPvalue <- apply(cbind(pchi1, pchi2, pchi3), 1 ,max)

# Adjust for multiple testing of the cubed pValue
summarisedPvalue <- p.adjust(summarisedPvalue^3, method = "BH")

# add Names
names(summarisedPvalue) <- rownames(test1)

# Compute M-score
Mscore <- -log10(summarisedPvalue)

# compute reproducibility sore
Rscore1 <- reprodScore(x = delta1, y = delta2)
Rscore2 <- reprodScore(x = delta2, y = delta3)
Rscore3 <- reprodScore(x = delta3, y = delta1)

# take smallest score
Rscore <- apply(cbind(Rscore1, Rscore2, Rscore3), 1,function(x) min(x, na.rm = TRUE))

# add Names
names(Rscore) <- names(Mscore) <- rownames(test1)

plot(Mscore, Rscore)

## Compute ROC 
MRscore <- ((Rscore - mean(Rscore))/sd(Rscore)) * ((Mscore - mean(Mscore[!is.infinite(Mscore)]))/sd(Mscore[!is.infinite(Mscore)]))
names(MRscore) <- rownames(test1)
MRscore[is.infinite(MRscore)] <- max(MRscore[!is.infinite(MRscore)])

  return(list(Rscore = Rscore, Mscore = Mscore, MRscore = MRscore, sqMan1 = sqMan1, sqMan2 = sqMan2, sqMan3 = sqMan3))
}

# MRscore <- MRscore[names(outcome)]
# MRscore_ord <- MRscore[order(MRscore, decreasing = TRUE)]
# 
# 
# ROC <- matrix(NA, nrow = length(MRscore_ord), ncol = 2)
# PrecRecall <- matrix(NA, nrow = length(MRscore_ord), ncol = 2)
# for (i in seq.int(length(MRscore_ord))) {
#   
#   TP <- sum((names(MRscore_ord[1:i])  %in% perm1_names))
#   FN <- sum(!(perm1_names %in% names(MRscore_ord[1:i])))
#   TN <- sum(!(names(MRscore_ord[-c(1:i)])  %in% perm1_names))
#   FP <- sum(!(names(MRscore_ord[1:i])  %in% perm1_names))
#   
#   ROC[i, 1] <- 1 - (TN/(TN + FP))
#   ROC[i, 2] <- TP/(TP + FN)
#   
#   PrecRecall[i, 1] <- TP/(TP + FN) # recall
#   PrecRecall[i, 2] <- TP/(TP + FP) # precision
#   
#   
# }
# plot(ROC, type = "b", col = "blue", lwd = 1, ylim = c(0,1), xlim = c(0,1))
# points(seq.int(100)/100, seq.int(100)/100, type = "b", col = "red")
# 
# plot(PrecRecall, type = "b", col = "blue", lwd = 1, ylim = c(0,1), xlim = c(0,1))
# points(seq.int(100)/100, rep(length(perm1)/nrow(tan2009r1), 100), type = "b", col = "red")
```
