## Supplementary Code 1 for "Inferring differential subcellular localisation in comparative spatial proteomics using BANDLE": pg_sample.cpp

```
#include <Rcpp.h>
using namespace Rcpp;

// Mathematical constants computed using Wolfram Alpha
#define MATH_PI        3.141592653589793238462643383279502884197169399375105820974
#define MATH_PI_2      1.570796326794896619231321691639751442098584699687552910487
#define MATH_2_PI      0.636619772367581343075535053490057448137838582961825794990
#define MATH_PI2       9.869604401089358618834490999876151135313699407240790626413
#define MATH_PI2_2     4.934802200544679309417245499938075567656849703620395313206
#define MATH_SQRT1_2   0.707106781186547524400844362104849039284835937688474036588
#define MATH_SQRT_PI_2 1.253314137315500251207882642405522626503493370304969158314
#define MATH_LOG_PI    1.144729885849400174143427351353058711647294812915311571513
#define MATH_LOG_2_PI  -0.45158270528945486472619522989488214357179467855505631739
#define MATH_LOG_PI_2  0.451582705289454864726195229894882143571794678555056317392

// FCN prototypes
double samplepg(double);
double exprnd(double);
double tinvgauss(double, double);
double truncgamma();
double randinvg(double);
double aterm(int, double, double);

// [[Rcpp::export]]
NumericVector rcpp_pgdraw(NumericVector b, NumericVector c)
{
  int m = b.size();
  int n = c.size();
  NumericVector y(n);
  
  // Setup
  int i, j, bi = 1;
  if (m == 1)
  {
    bi = b[0];
  }
  
  // Sample
  for (i = 0; i < n; i++)
  {
    if (m > 1)
    {
      bi = b[i];
    }
    
    // Sample
    y[i] = 0;
    for (j = 0; j < (int)bi; j++)
    {
      y[i] += samplepg(c[i]);
    }
  }
  
  return y;
}


// Sample PG(1,z)
// Based on Algorithm 6 in PhD thesis of Jesse Bennett Windle, 2013
// URL: https://repositories.lib.utexas.edu/bitstream/handle/2152/21842/WINDLE-DISSERTATION-2013.pdf?sequence=1
double samplepg(double z)
{
  //  PG(b, z) = 0.25 * J*(b, z/2)
  z = (double)std::fabs((double)z) * 0.5;
  
  // Point on the intersection IL = [0, 4/ log 3] and IR = [(log 3)/pi^2, \infty)
  double t = MATH_2_PI;
  
  // Compute p, q and the ratio q / (q + p)
  // (derived from scratch; derivation is not in the original paper)
  double K = z*z/2.0 + MATH_PI2/8.0;
  double logA = (double)std::log(4.0) - MATH_LOG_PI - z;
  double logK = (double)std::log(K);
  double Kt = K * t;
  double w = (double)std::sqrt(MATH_PI_2);
  
  double logf1 = logA + R::pnorm(w*(t*z - 1),0.0,1.0,1,1) + logK + Kt;
  double logf2 = logA + 2*z + R::pnorm(-w*(t*z+1),0.0,1.0,1,1) + logK + Kt;
  double p_over_q = (double)std::exp(logf1) + (double)std::exp(logf2);
  double ratio = 1.0 / (1.0 + p_over_q); 
  
  double u, X;
  
  // Main sampling loop; page 130 of the Windle PhD thesis
  while(1) 
  {
    // Step 1: Sample X ? g(x|z)
    u = R::runif(0.0,1.0);
    if(u < ratio) {
      // truncated exponential
      X = t + exprnd(1.0)/K;
    }
    else {
      // truncated Inverse Gaussian
      X = tinvgauss(z, t);
    }
    
    // Step 2: Iteratively calculate Sn(X|z), starting at S1(X|z), until U ? Sn(X|z) for an odd n or U > Sn(X|z) for an even n
    int i = 1;
    double Sn = aterm(0, X, t);
    double U = R::runif(0.0,1.0) * Sn;
    int asgn = -1;
    bool even = false;
    
    while(1) 
    {
      Sn = Sn + asgn * aterm(i, X, t);
      
      // Accept if n is odd
      if(!even && (U <= Sn)) {
        X = X * 0.25;
        return X;
      }
      
      // Return to step 1 if n is even
      if(even && (U > Sn)) {
        break;
      }
      
      even = !even;
      asgn = -asgn;
      i++;
    }
  }
  return X;
}

// Generate exponential distribution random variates
double exprnd(double mu)
{
  return -mu * (double)std::log(1.0 - (double)R::runif(0.0,1.0));
}

// Function a_n(x) defined in equations (12) and (13) of
// Bayesian inference for logistic models using Polya-Gamma latent variables
// Nicholas G. Polson, James G. Scott, Jesse Windle
// arXiv:1205.0310
//
// Also found in the PhD thesis of Windle (2013) in equations
// (2.14) and (2.15), page 24
double aterm(int n, double x, double t)
{
  double f = 0;
  if(x <= t) {
    f = MATH_LOG_PI + (double)std::log(n + 0.5) + 1.5*(MATH_LOG_2_PI- (double)std::log(x)) - 2*(n + 0.5)*(n + 0.5)/x;
  }
  else {
    f = MATH_LOG_PI + (double)std::log(n + 0.5) - x * MATH_PI2_2 * (n + 0.5)*(n + 0.5);
  }    
  return (double)exp(f);
}

// Generate inverse gaussian random variates
double randinvg(double mu)
{
  // sampling
  double u = R::rnorm(0.0,1.0);
  double V = u*u;
  double out = mu + 0.5*mu * ( mu*V - (double)std::sqrt(4.0*mu*V + mu*mu * V*V) );
  
  if(R::runif(0.0,1.0) > mu /(mu+out)) {    
    out = mu*mu / out; 
  }    
  return out;
}

// Sample truncated gamma random variates
// Ref: Chung, Y.: Simulation of truncated gamma variables 
// Korean Journal of Computational & Applied Mathematics, 1998, 5, 601-610
double truncgamma()
{
  double c = MATH_PI_2;
  double X, gX;
  
  bool done = false;
  while(!done)
  {
    X = exprnd(1.0) * 2.0 + c;
    gX = MATH_SQRT_PI_2 / (double)std::sqrt(X);
    
    if(R::runif(0.0,1.0) <= gX) {
      done = true;
    }
  }
  
  return X;  
}

// Sample truncated inverse Gaussian random variates
// Algorithm 4 in the Windle (2013) PhD thesis, page 129
double tinvgauss(double z, double t)
{
  double X, u;
  double mu = 1.0/z;
  
  // Pick sampler
  if(mu > t) {
    // Sampler based on truncated gamma 
    // Algorithm 3 in the Windle (2013) PhD thesis, page 128
    while(1) {
      u = R::runif(0.0, 1.0);
      X = 1.0 / truncgamma();
      
      if ((double)std::log(u) < (-z*z*0.5*X)) {
        break;
      }
    }
  }  
  else {
    // Rejection sampler
    X = t + 1.0;
    while(X >= t) {
      X = randinvg(mu);
    }
  }    
  return X;
}
```
