## Supplementary Code 1 for "Inferring differential subcellular localisation in comparative spatial proteomics using BANDLE": trenchDetcpp.cpp

```
# include <RcppArmadillo.h>
// [[Rcpp::depends(RcppArmadillo)]]
using namespace Rcpp;

// [[Rcpp::export]]
List trenchDetcpp(arma::vec c) {
  
  int N = c.size();
  int i;
  arma::vec xi(N-1);
  arma::vec z(N-1);
  arma::vec v(N);
  double beta = 1;
  double alpha;
  double l = 0;
  double logdet;
  
  xi = c.subvec(1, N-1)/c(0);
  z(0) = - xi(0);
  alpha = -xi(0);
  int K = xi.size(); 
  
  for(i = 0; i < (K - 1); i++){
    beta = (1 - pow(alpha, 2)) * beta ;
    l = l + log(beta) ;
    if(i == 0){
      alpha = - (xi(i + 1) + xi(0) * z(0)) / beta ;
      z(0) = z(0) + alpha * z(0) ;
    }else{
      alpha = - (xi(i + 1) + dot(flipud(xi.subvec(0, i)), z.subvec(0, i)) ) / beta ;
      z.subvec(0, i) = z.subvec(0, i) + alpha * flipud(z.subvec(0, i)) ;
    }
    
    z(i+1) = alpha  ;
  }
  
  beta = (1 - pow(alpha, 2)) * beta ;
  l = l + log(beta) ;
  
  logdet = l + N * log(c(0));
  
  v(N-1) = 1 / ((1 + dot(xi, z)) * c(0)) ;
  v.subvec(0,N-2) = v(N-1) * flipud(z.subvec(0, N - 2));
  
  return List::create(Rcpp::Named("logdet") = logdet,
                      Rcpp::Named("z") = z,
                      Rcpp::Named("v") = v);
}

// [[Rcpp::export]]
arma::mat trenchInvcpp(arma::vec v) {
  
  int N = v.size();
  int i;
  int j;
  arma::mat C(N, N);
  arma::mat trenchInv;
  
  
  C.row(0) = flipud(v).t();
  C.col(0) = flipud(v);
  C.row(N - 1) = v.t();
  C.col(N - 1) = v;
  for(i = 1; i < floor( (N - 1) / 2 ) + 1; i++){
    for(j = 1; j < N - i; j++){
     C(i, j) = C(i - 1, j - 1) + (v(N - 1 - j) * v(N - 1 - i) - v(i - 1) * v(j - 1)) / v(N - 1) ;
     C(j, i) = C(i, j);
     C(N - 1 - i , N - 1 - j ) = C(i, j);
     C(N - 1 - j , N - 1 - i ) = C(i, j);
    }
  } 
  
  trenchInv = C;
  return(trenchInv) ; 
  
}

//  arma::vec loglikeGPcpp(arma::vec Xk,
//                         arma::mat Z,
//                         arma::mat A,
//                         double logcovDet,
//                         double sigmak,
//                         double nk,
//                         double D){
//    
//    arma::vec loglike;
//    arma::mat I = arma::eye(D, D);
//    arma::mat J = arma::ones(nk, nk);
//    
//    loglike = - dot(Xk,Xk) / (2 * sigmak) + ((Xk.t() * arma::kron(J, I - Z )) * Xk) / (2 * sigmak * nk) 
//      -  logcovDet / 2 - (nk * D *  log(2 * M_PI)) / 2 ;
    
//    return(loglike); 
//  }

arma::vec loglikeGPcpp(arma::vec Y,
                       arma::mat Z,
                       arma::mat A,
                       double logcovDet,
                       double sigmak,
                       double nk,
                       double D,
                       double Y2){
  
  arma::vec loglike;
  arma::mat I = arma::eye(D, D);
  arma::mat J = arma::ones(nk, nk);

  loglike = - Y2/ (2 * sigmak) + (Y.t() * ((I - Z) * Y))/ (2 * sigmak * nk) 
    -  logcovDet / 2 - (nk * D *  log(2 * M_PI)) / 2 ;
  
  return(loglike); 
}


// [[Rcpp::export]]
double likelihoodGPcpp(arma::mat &Xk,
                       arma::vec tau,
                       arma::vec h,
                       int nk,
                       int D
                      ){
  double sigmak; sigmak = exp(2 * h(2));
  double a; a = exp(2 * h(1));
  double l; l = exp(h(0));
  double negloglike;
  double logdet;
  double logcovDet;
  double Y2; //keep track of sum of squares
  arma::mat Z(D,D);
  arma::mat S(D,D); S = arma::zeros(D,D);
  arma::mat A(D,D);
  arma::mat R(D,D);
  arma::vec temp;
  arma::vec v;
  arma::vec Y(D); //keep track of means
  List trenchres = List::create(Rcpp::Named("logdet"),
                                Rcpp::Named("z"),
                                Rcpp::Named("v"));
  
  S = S.each_col() + tau;
  A = a * exp(- pow(S - S.t(), 2)/l);
  R = arma::eye(D,D) + (nk * A)/sigmak;
  
  trenchres = trenchDetcpp(R.row(0).t());
  logdet = trenchres["logdet"];
  v = as<arma::vec>(trenchres["v"]);
  

  if(R_IsNA(trenchres["logdet"])){
    negloglike = R_NegInf;
    return(negloglike);
  }
  
  Z = trenchInvcpp(v);
  logcovDet = (nk * D) * log(sigmak) + logdet ;
  
  Y2 = arma::accu(pow(Xk, 2));  
  Y = sum(Xk, 1);
  
  negloglike = -loglikeGPcpp(Y, Z, A, logcovDet, sigmak, nk, D, Y2)(0,0);
  
  return(negloglike);
  
}

// [[Rcpp::export]]
arma::vec sampleGPmeancpp(arma::mat Xk,
                          arma::vec tau,
                          arma::vec h,
                          int nk,
                          int D
){
  double sigmak; sigmak = exp(2 * h(2));
  double a; a = exp(2 * h(1));
  double l; l = exp(h(0));
  arma::mat Z(D,D);
  arma::mat S(D,D); S = arma::zeros(D,D);
  arma::mat A(D,D);
  arma::mat R(D,D);
  arma::mat J = arma::ones(nk, nk);
  arma::mat invC(D * nk , D * nk);
  arma::mat priorCor(D, D * nk);
  arma::mat postV(D, D);
  arma::mat cholV(D, D);
  arma::vec v;
  arma::vec postM;
  arma::vec GPmean;
  arma::vec eigval;
  arma::mat eigvec;
  arma::mat eye = arma::eye(D, D);
  arma::vec Y(D);
  List trenchres = List::create(Rcpp::Named("logdet"),
                                Rcpp::Named("z"),
                                Rcpp::Named("v"));
  
  S = S.each_col() + tau;
  A = a * exp(- pow(S - S.t(), 2)/l);
  R = arma::eye(D, D) + (nk * A)/sigmak;
  
  trenchres = trenchDetcpp(R.row(0).t());
  v = as<arma::vec>(trenchres["v"]);
  
  Z = trenchInvcpp(v);
  //invC = (arma::eye(D * nk , D * nk) / sigmak)  - arma::kron(J, eye - Z)/(sigmak * nk);
  //priorCor = kron(arma::ones(1, nk), A);
  //postM = priorCor * (invC * Xk);
  
  Y = sum(Xk, 1);
  postM =  ((eye - Z) * Y) / nk ;

  //postV = A - (nk * (A * A)/sigmak) + nk * (A * (eye - Z) * A)/(sigmak);
  //postV = A - (nk * A * Z * A)/(sigmak)
  postV = (eye - Z)  * sigmak/nk;
  eig_sym(eigval, eigvec, postV);
  
  for(int i = 0; i < D; i++){
    if(eigval[i] < 0 ){
      eigval[i] = 10e-6;
    }
  }
  //cholV = arma::chol(postV);
  GPmean = postM + (eigvec * (eye.each_col() % sqrt(eigval))) * as<arma::vec>(rnorm(D));
  
  return(GPmean);
  
}

arma::mat subset(arma::mat &X,
                 arma::vec &B,
                 int j){
  
  arma::uvec ind_ = find(B==j);
  arma::mat subX ;
  
  subX = X.rows(ind_);

  return(subX);
  
}

arma::vec makevector(arma::mat &X){
  
  arma::vec Y;
  Y = arma::vectorise(X);
  return(Y);

}

// [[Rcpp::export]]
arma::mat makeComponent(arma::mat &X,
                        arma::vec &BX,
                        arma::mat &Y,
                        arma::vec &BY,
                        int j){
  arma::mat subX;
  arma::mat subY;
  arma::mat component;
  
  subX = subset(X, BX, j);
  subY = subset(Y, BY, j);
  
  //vecX = makevector(subX.t());
  //vecY = makevector(subY.t());  
  component = arma::join_cols(subX, subY);
  
  return(component);
}

// [[Rcpp::export]]
arma::mat normalisedData(arma::mat &Xknown,
                         arma::vec &BX,
                         arma::mat &Xunknown,
                         arma::vec &BXun,
                         arma::vec &hypers,
                         arma::vec &nk,
                         arma::vec &tau,
                         int D,
                         int j){
  arma::mat Xk;
  arma::vec sampleGPMean;
  //arma::mat outbefore;
  int n = Xunknown.n_rows;
  arma::mat out(n, D);
  
  
  
  Xk = makeComponent(Xknown, BX, Xunknown, BXun, j);
  sampleGPMean = sampleGPmeancpp(Xk.t(), tau, hypers, nk(j-1), D); //need to correct still
  for (int d = 0; d < D; d++) {
   for (int i = 0; i < n; i++) {
      out(i, d) = Xunknown(i, d) - sampleGPMean(d);
    }
  }
  //outbefore = Xunknown.each_row() - sampleGPMean.t();
  
  return(out);
}

// [[Rcpp::export]]
List centeredData(arma::mat Xknown,
                  arma::vec BX,
                  arma::mat Xunknown,
                  arma::vec BXun,
                  arma::mat hypers,
                  arma::vec nk,
                  arma::vec tau,
                  int D,
                  int K){
  int j;
  List centereddata(K);
  arma::vec currenthyper(3);
  
  for(j = 0; j < K; j++){
    currenthyper = hypers.row(j).t();
    centereddata[j] = normalisedData(Xknown,
                                    BX,
                                    Xunknown, 
                                    BXun,
                                    currenthyper, 
                                    nk, tau, D = D, j + 1).t();
  }
   
    
  return(centereddata);
}

// [[Rcpp::export]]
arma::vec sampleDirichlet(int numSamples,
                          NumericVector alpha
                            ){
  
  int j;
  NumericVector shape = alpha;
  arma::vec gamma(numSamples);
  NumericVector dirichlet;
  double scale = 1;
  
  for(j = 0; j < numSamples; j++){
   gamma(j) = rgamma(1, shape(j), scale)(0); 
  }
  
  dirichlet = gamma/sum(gamma);
  
  return(dirichlet);
}

// [[Rcpp::export]]
arma::vec componentloglike(const arma::mat &centereddata,
                           double sigmak){
  int d;
  int D = centereddata.n_rows;
  int N = centereddata.n_cols;
  arma::vec likelihood;
  arma::mat likelihoods(N,D);
  arma::vec sumlikelihood;

  for(d = 0; d < D; d++){
   likelihood = dnorm(as<NumericVector>(wrap(centereddata.row(d))), 0, sigmak, 1);
   likelihoods.col(d) = likelihood;
  }
  
  sumlikelihood = rowSums(as<NumericMatrix>(wrap(likelihoods)));
  return(sumlikelihood);

}

// [[Rcpp::export]]
arma::mat comploglike(const List &centereddata,
                      arma::vec sigmak
                      ){
  
  int K = sigmak.size();
  int N = as<arma::mat>(wrap(centereddata[1])).n_cols;
  int j;
  arma::mat likelihoods(N, K);
  
  for(j = 0; j < K; j++){
    likelihoods.col(j) = componentloglike(centereddata[j], sigmak(j)); 
  }
  
  return(likelihoods);
}

// [[Rcpp::export]]
List comploglikelist(const List &centereddata,
                     List sigmak) {
  
  int numRep = centereddata.size();
  int l;
  arma::mat loglikelihood;
  List loglikelihoods(numRep);

  for (l = 0; l < numRep; l++) {
    loglikelihoods[l] = comploglike(centereddata[l], sigmak[l]);
    //loglikelihoods[l] = loglikelihood;
  }

  return(loglikelihoods);
  
}


// [[Rcpp::export]]
arma::vec sampleOutliercpp(arma::mat allocoutlierprob){
  
  int i;
  int N = allocoutlierprob.n_rows;
  arma::vec u; 
  arma::vec outlier = arma::zeros(N);
  Rcpp::NumericVector allocoutlierprobi;
  
  for (i = 0; i < N; i++) {
    allocoutlierprobi = allocoutlierprob.row(i);
    
    u = runif(1, 0, 1);
    if (u(0) > allocoutlierprobi(1)) {
      outlier(i) = 1;
    }
  }
  
  return(outlier);
  
}

// [[Rcpp::export]]
arma::vec sampleAlloccpp(arma::mat allocprob) {
  
  int i;
  int N = allocprob.n_rows;
  int k;
  int classNumber = allocprob.n_cols;
  arma::vec u;
  arma::vec u1 = arma::ones(classNumber);
  arma::vec alloc = arma::zeros(N);
  arma::vec allocprobi;
  arma::vec probcum;
  Rcpp::LogicalVector loc;
 
  
  for (i = 0; i < N; i++) {
    allocprobi = allocprob.row(i).t(); 
    probcum = cumsum(allocprobi);
    u = runif(1, 0, 1);
    u1 = (u * u1.t()).t();
    loc = (probcum <= u1);
    k = sum(loc) + 1;
    alloc(i) = k;
    u1 = arma::ones(classNumber); //reset u1
  }

  return(alloc);
}


/*** R
res <- trenchDetcpp(c(2, -1, 0))
res
abs(trenchDetcpp(c(2, -1, 0))$logdet - log(det(matrix(c(2,-1,0,-1,2,-1,0,-1,2), nrow = 3)))) < 10-8
res2 <- trenchDetcpp(c(2, 1, 0, 0))
res2
abs(trenchDetcpp(c(2, 1, 0, 0))$logdet - log(det(matrix(c(2,1,0,0,1,2,1,0,0,1,2,1,0,0,1,2), nrow = 4)))) < 10-8
abs(sum(trenchInvcpp(res$v) - solve(matrix(c(2,-1,0,-1,2,-1,0,-1,2), nrow = 3)))) < 10-8
abs(sum(trenchInvcpp(res2$v) - solve(matrix(c(2,1,0,0,1,2,1,0,0,1,2,1,0,0,1,2), nrow = 4)))) <10-8
*/
```
