## Supplementary Code 1 for "Inferring differential subcellular localisation in comparative spatial proteomics using BANDLE": bandle_beltranvirus.R

```
## Analysis of dynamic HCMV infection
source("bandleFUN.R")
require(pRoloc)
require(pRolocdata)


data("beltran2016HCMV24")
data("beltran2016HCMV48")
load("data/beltranvirushypers.rda")

par(mfrow = c(1,1))
#plot2D(beltran2016HCMV24, method = "t-SNE")
#plot2D(beltran2016HCMV48, method = "t-SNE")

idx <- intersect(rownames(beltran2016HCMV24), rownames(beltran2016HCMV48))

HCMV24_sub <- beltran2016HCMV24[idx, ]
HCMV48_sub <- beltran2016HCMV48[idx, ]


beltrandynv <- c(HCMV24_sub, HCMV48_sub) 
beltrandynv <- lapply(beltrandynv, function(x) normalise(x, method = "center.median"))

K <- length(getMarkerClasses(HCMV24_sub))

dir_prior = diag(rep(1, K)) + matrix(0.01, nrow = K, ncol = K)
prior_pred_dir(object = HCMV24_sub, dir_prior = dir_prior)

par(mfrow = c(3,4))
pc_prior <- matrix(NA, ncol = 3, K)
pc_prior[seq.int(1:K), ] <- matrix(rep(c(10, 60, 250), each = K), ncol = 3)
#hypres <- lapply(beltrandynv, function(x) fitGPmaternPC(x, fcol = "markers", nu = 2, materncov = TRUE, hyppar = pc_prior)) 
#save(hypres, file = "beltranvirushypers.rda")

par(mfrow = c(4,4))
bandleres_beltranvirus <- bandle(object_cond1 = c(beltrandynv[[1]]),
                              object_cond2 = c(beltrandynv[[2]]), params = hypres,
                              numIter = 20000, burnin = 10000, thin = 10, pg = FALSE, hypLearn = "LFBGS", 
                              maternCov = TRUE, nu = 2, pc_prior = pc_prior, numChains = 9, dir_prior = dir_prior)

save(bandleres_beltranvirus, file = "bandleres_beltranvirus.rda")
```
