## Supplementary Code 1 for "Inferring differential subcellular localisation in comparative spatial proteomics using BANDLE": bandle_davies.R

```
source("bandleFUN.R")

load("data/davies2018wt.rda")
load("data/davies2018ap4b1.rda")
load("data/davies2018ap4e1.rda")
load("data/davies_hypers2.rda")


davies2018wt <- filterNA(davies2018wt)
davies2018ap4b1 <- filterNA(davies2018ap4b1)
davies2018ap4e1 <- filterNA(davies2018ap4e1)

exprs(davies2018wt) <- log2(exprs(davies2018wt))
exprs(davies2018ap4b1) <- log2(exprs(davies2018ap4b1))
exprs(davies2018ap4e1) <- log2(exprs(davies2018ap4e1))

par(mfrow = c(2,3))
plot2D(davies2018wt[,1:5], dims = c(1,2))
plot2D(davies2018wt[,6:10], dims = c(1,2))
plot2D(davies2018ap4b1, dims = c(1,2))
plot2D(davies2018ap4e1[,1:5], dims = c(1,2))
getMarkerClasses(davies2018wt, fcol = "markers")

par(mfrow = c(1,1))
pRoloc::plotDist(davies2018wt, fcol = "markers")

exprs(davies2018wt)

subactin <- fData(davies2018wt)$markers == "Actin binding proteins"
fData(davies2018wt)[rownames(davies2018wt[subactin]), "markers"] <- "unknown"


subactin <- fData(davies2018ap4b1)$markers == "Actin binding proteins"
fData(davies2018ap4b1)[rownames(davies2018ap4b1[subactin]), "markers"] <- "unknown"

subnpc <- fData(davies2018ap4b1)$markers == "Nuclear pore complex"
fData(davies2018ap4b1)[rownames(davies2018ap4b1[subnpc]), "markers"] <- "unknown"

subnpc <- fData(davies2018wt)$markers == "Nuclear pore complex"
fData(davies2018wt)[rownames(davies2018wt[subnpc]), "markers"] <- "unknown"


K <- length(getMarkerClasses(davies2018wt))
dir_prior = diag(rep(1, K)) + matrix(0.02, nrow = K, ncol = K)
prior_pred_dir(object = davies2018wt, dir_prior = dir_prior)

pc_prior <- matrix(NA, ncol = 3, K)
pc_prior[seq.int(1:K), ] <- matrix(rep(c(0.05, 60, 100), each = K), ncol = 3)


daviesdyn <- c(davies2018wt[ ,1:5],
               davies2018wt[, 6:10],
               davies2018ap4b1[, 1:5],
               davies2018ap4b1[, 6:10])

daviesdyn <- lapply(daviesdyn, function(x) normalise(x, methed = "center.median"))


par(mfrow = c(4,4))
bandleres_daviesb4 <- bandle(object_cond1 = c(daviesdyn[[1]], daviesdyn[[2]]),
                             object_cond2 = c(daviesdyn[[3]], daviesdyn[[4]]), params = hypres,
                              numIter = 20000, burnin = 10000, thin = 15, pg = FALSE, hypLearn = "LFBGS", 
                              maternCov = TRUE, nu = 2, pc_prior = pc_prior, numChains = 6, dir_prior = dir_prior)

save(bandleres_daviesb4, file = "bandleres_daviesb4.rda")
```
