## Supplementary Code 1 for "Inferring differential subcellular localisation in comparative spatial proteomics using BANDLE": bandle_Itzhak.R

```
source("bandleFUN.R")


data(itzhak2016helaCtrl)
data(itzhak2016helaEgf)
load("data/itzhak2016hypers2.rda")

helaCtrl <- itzhak2016helaCtrl
helaEgf <- itzhak2016helaEgf

helaCtrl <- filterNA(helaCtrl)
helaEgf <- filterNA(helaEgf)

idx <- intersect(rownames(helaCtrl), rownames(helaEgf))

helaCtrl_sub <- helaCtrl[idx, ]
helaEgf_sub <- helaEgf[idx, ]

subgol <- fData(helaCtrl_sub)$markers == "Golgi"
fData(helaCtrl_sub)[rownames(helaCtrl_sub[subgol]), "markers"] <- "unknown"
subper <- fData(helaCtrl_sub)$markers == "Peroxisome"
fData(helaCtrl_sub)[rownames(helaCtrl_sub[subper]), "markers"] <- "unknown"
subnuc <- fData(helaCtrl_sub)$markers == "Nuclear pore complex"
fData(helaCtrl_sub)[rownames(helaCtrl_sub[subnuc]), "markers"] <- "unknown"

subgol <- fData(helaEgf_sub)$markers == "Golgi"
fData(helaEgf_sub)[rownames(helaEgf_sub[subgol]), "markers"] <- "unknown"
subper <- fData(helaEgf_sub)$markers == "Peroxisome"
fData(helaEgf_sub)[rownames(helaEgf_sub[subper]), "markers"] <- "unknown"
subnuc <- fData(helaEgf_sub)$markers == "Nuclear pore complex"
fData(helaEgf_sub)[rownames(helaEgf_sub[subnuc]), "markers"] <- "unknown"

itzhakdyn <- list(helaCtrl_sub[, 1:5],
                  helaCtrl_sub[, 6:10],
                  helaCtrl_sub[, 11:15],
                  helaEgf_sub[, 1:5],
                  helaEgf_sub[, 6:10],
                  helaEgf_sub[, 11:15])


K <- length(getMarkerClasses(helaCtrl_sub))
dir_prior = diag(rep(1, K)) + matrix(0.02, nrow = K, ncol = K)
colnames(dir_prior) <- rownames(dir_prior) <- getMarkerClasses(helaCtrl_sub)
dir_prior["Plasma membrane", "Lysosome"] <- 1

tmp <- prior_pred_dir(object = helaCtrl_sub, dir_prior = dir_prior)

pc_prior <- matrix(NA, ncol = 3, K)
pc_prior[seq.int(1:K), ] <- matrix(rep(c(0.05, 60, 100), each = K), ncol = 3)


bandleres_Itzhak5 <- bandle(object_cond1 = c(helaCtrl_sub[, 1:5], helaCtrl_sub[, 6:10], helaCtrl_sub[, 11:15]),
                           object_cond2 = c(helaEgf_sub[, 1:5], helaEgf_sub[, 6:10], helaEgf_sub[, 11:15]),
                           params = itzhak2016hypers2, numIter = 20000, burnin = 10000, thin = 10, pg = FALSE, hypLearn = "LFBGS", 
                           maternCov = TRUE, nu = 2, pc_prior = pc_prior, numChains = 8, dir_prior = dir_prior)

save(bandleres_Itzhak5, file = "bandleres_Itzhak5.rda")
```
