## Supplementary Code 1 for "Inferring differential subcellular localisation in comparative spatial proteomics using BANDLE": bandleandMRsim.R

```
## Direct comparison of MR and BANDLE on simulated data
## Results simulations
setwd("C:/Users/OllyC/Desktop/2019-DiffLoc")
require(pRoloc)
require(pRolocdata)


# functions requried
source("diffLoc.R")
source("diffLocUtils.R")
source("fitGPmatern.R")
source("fitGPmaternPC.R")
source("simDynamic.R")
source("metropolisGP.R")
source("MRmethod.R")
data("tan2009r1")

pRoloc::setStockcol(NULL)
pRoloc::setStockcol(paste0(pRoloc::getStockcol(), 90))
.seed <- set.seed(314)

simres <- sim_dynamic(object = tan2009r1)

tan2009rep <- simres$lopitrep
perm1_names <- simres$perm1_names

# computational methods
mrres <- mrMethod(c(tan2009rep))
par(mfrow = c(3,4))
pc_prior <- matrix(NA, ncol = 3, 11)
pc_prior[seq.int(1:11),] <- matrix(rep(c(0.05,60,100), each = 11), ncol = 3)

resdynpg <- diffLoc(object_cond1 = c(tan2009rep[1:3]), object_cond2 = tan2009rep[4:6], numIter = 20000, burnin = 10000,
                    pg = TRUE, hypLearn = "LBFGS", tau = 0.3, maternCov = TRUE, nu = 2, pc_prior = pc_prior)
par(mfrow = c(3,4))
resdyndir <- diffLoc(object_cond1 = c(tan2009rep[1:3]), object_cond2 = c(tan2009rep[4:6]),
                     numIter = 20000, burnin = 10000, pg = FALSE, hypLearn = "LFBGS", 
                     maternCov = TRUE, nu = 2, pc_prior = pc_prior)

save(resdynpg, file = "figure2pgdata.rda")
save(resdyndir, file = "figure2dirdata.rda")


probpg <- probsameorganelle(object = tan2009r1, params = resdynpg)
probdir <- probsameorganelle(object = tan2009r1, params = resdyndir)

outcome <- matrix(0, ncol = 1, nrow = length(probpg))
names(outcome) <- names(probpg)
outcome[perm1_names] <- 1
prediction <- probpg/ncol(resdynpg$allocOut[[1]])
par(mfrow = c(1,1))

ROC_bayes <- roc(outcome, 1 - prediction)
ROC_mr <- roc(outcome, mrres$MRscore[names(outcome)], direction = "<")
plot(ROC_bayes, type = "b", col = "darkgreen", lwd = 1, pch = 19, asp = NA, legacy.axes = T)
lines(ROC_mr, type = "b", col = "blue", lwd = 1, pch = 19, asp = NA)

outcome2 <- matrix(0, ncol = 1, nrow = length(probdir))
names(outcome2) <- names(probdir)
outcome2[perm1_names] <- 1
prediction2 <- probdir/ncol(resdyndir$allocOut[[1]])
ROC_dir <- roc(outcome2, 1 - prediction2)
lines(ROC_dir,type = "b", col = "orange", lwd = 1, pch = 19, asp = NA)

legend("bottomright", c("MR", "Dirichlet", "Polya-Gamma"), col = c("blue", "orange", "darkgreen"), pch = 19)


par(mfrow = c(1,2))
plot2D(tan2009rep[[1]], grid = FALSE)
highlightOnPlot(object = tan2009rep[[1]], foi = perm1_names, pch = 18, cex = 1.5)
addLegend(object = tan2009rep[[1]], ncol = 2)
plot2D(tan2009rep[[4]], grid = FALSE)
highlightOnPlot(object = tan2009rep[[4]], foi = perm1_names, pch = 18, cex = 1.5)

par(mfrow = c(1,2))
mycol <- rep(getStockcol()[2], length(mrres$Mscore))
names(mycol) <- names(mrres$Mscore)
mycol[perm1_names] <- getStockcol()[4]
plot(mrres$Mscore, mrres$Rscore, col = mycol, pch = 19, cex = 1.5, xlab = "M-score", ylab = "R-Score",main = "MR-plot")
abline(h = 0.9, col = getStockcol()[7], lwd = 4)
abline(v = 2, col = getStockcol()[7], lwd = 4)

pvalues <- (10^(-mrres$Mscore))^(1/3)
hist(pvalues, breaks = 40, col = getStockcol()[3], xlab = "p-value", freq = FALSE, main = "Histogram of p-values")
abline(h = 1, col = getStockcol()[13], lwd = 4)

require(MASS)
dist.resgamma <- fitdistr(mrres$sqMan1, densfun = "gamma", list(shape = 8, scale = 2))
dist.reschi <- fitdistr(mrres$sqMan1, densfun = "chi-squared", list(df = 4))

hist(qchisq(pvalues, df = 4, lower.tail = FALSE), breaks = 40,  main = "Raw Statistics", xlab = "chi-squared statistic", 
     col = getStockcol()[3], freq = FALSE, ylim = c(0, 0.2), text.cex = 16)
hist(mrres$sqMan1, breaks = 40,  main = "Raw Statistics", xlab = "chi-squared statistic", 
     col = getStockcol()[3], freq = FALSE, ylim = c(0, 0.2))
curve( dchisq(x, df=4), col= getStockcol()[4], add=TRUE, lwd = 5)
curve(dgamma(x, shape = dist.resgamma$estimate[1], scale = dist.resgamma$estimate[2]), col= getStockcol()[2], add=TRUE, lwd = 5)
legend(1, 95, legend=c("Line 1", "Line 2"),
       col=c("red", "blue"), lty=1:2, cex=0.8)

df.dist <- c(-dist.reschi$loglik, -dist.resgamma$loglik)
names(df.dist) <- c("Chi-squared", "Gamma")
barplot(df.dist,
        col= c(getStockcol()[4], getStockcol()[2]), ylab = "negative log-likelihood")

p.values.corrected <- pgamma(mrres$sqMan1, shape = dist.resgamma$estimate[1], scale = dist.resgamma$estimate[2], lower.tail = FALSE)
names(p.values.corrected) <- names(mrres$Mscore)
hist(p.values.corrected, freq = FALSE,
     col = getStockcol()[2], xlab = "p-value", main = "p-values")
abline(h = 1, col = getStockcol()[13], lwd = 4)

par(mfrow = c(2,2))
plot(mrres$Mscore, mrres$Rscore, col = mycol, pch = 19, cex = 1.5, xlab = "M-score", ylab = "R-Score",main = "MR-plot", cex.lab = 1.5)
abline(h = 0.9, col = getStockcol()[7], lwd = 4)
abline(v = 2, col = getStockcol()[7], lwd = 4)

hist(pvalues, breaks = 40, col = getStockcol()[3], xlab = "p-value", freq = FALSE, main = "Histogram of p-values",
     cex.main = 1.5, cex.lab = 1.5)
abline(h = 1, col = getStockcol()[13], lwd = 4)

hist(mrres$sqMan1, breaks = 40,  main = "Raw Statistics", xlab = "statistic", 
     col = getStockcol()[3], freq = FALSE, ylim = c(0, 0.2), cex.lab = 1.5)
curve( dchisq(x, df=4), col= getStockcol()[4], add=TRUE, lwd = 5)
curve(dgamma(x, shape = dist.resgamma$estimate[1], scale = dist.resgamma$estimate[2]), col= getStockcol()[2], add=TRUE, lwd = 5)
legend(70, 0.1, legend=c("Gamma", "Chi-squared"),
       col= c(getStockcol()[2], getStockcol()[4]), lty=1, cex=1.5,lwd = 3)


barplot(df.dist,
        col= c(getStockcol()[4], getStockcol()[2]), ylab = "negative log-likelihood", main = "Model Comparison",cex.lab = 1.5)

## BANDLE dirichlet
par(mfrow = c(1,1))
y <- 1 - probdir[order(probdir)]/ncol(resdyndir$allocOut[[1]])
plot(y, pch = 19, ylim = c(0,1), ylab = "Differential Localisation Probability", xlab = "Rank", 
     col = getStockcol()[2], main = "Rank plot from BANDLE", cex.lab = 1.5, cex.main = 2, cex.axis = 1.5)

par(mfrow = c(1,1))
bootresdir <- bootstrapdiffloc(object = tan2009rep[[1]], params = resdyndir, top = 30, Bootsample = 1000)
boxplot(t(1 - bootresdir/ncol(resdyndir$allocOut[[1]])), las = 2, col = getStockcol()[3])

require(reshape2)
df <- t(1 - bootresdir/ncol(resdyndir$allocOut[[1]]))
df_long <- melt(df)
colnames(df_long) <- c("bootsample", "Protein", "probability")
df_long$diff.loc <- rep(0, nrow(df_long))
df_long$diff.loc[df_long$Protein %in% perm1_names] <- 1
  
fill.col <- colnames(df) %in% perm1_names
fill.col[fill.col == TRUE] <- getStockcol()[4]
fill.col[fill.col == FALSE] <- getStockcol()[2]

require(ggplot2)
p <- ggplot(df_long, aes(x = Protein, y = probability, fill = Protein)) + geom_boxplot() + scale_fill_manual(values = fill.col)
p <- p + scale_y_continuous(limits = c(0.6, 1)) + theme(legend.position = "none")
p <- p + theme_classic() + theme(axis.text.x = element_text(angle = 90, colour = fill.col, size = 16), axis.text.y = element_text(size = 16),
                                 axis.title=element_text(size=20), plot.title = element_text(size = 22))  + theme(legend.position = "none")
p <- p + labs(title = "Rank plot of top 30 differentially localised proteins",
              x = "Protein",
              y = "Differential Localisation Probability")
p 

yloc <- y*0
yloc[perm1_names] <- 1 

plot(y, yloc, ylab = "Differentially localised", xlab = "probability")

require(vioplot)
par(mfrow = c(1,3))
vioplot(y[yloc == 0 ], y[yloc == 1], col = getStockcol()[2], main = "Differential localisation Probabilities",
        cex.main = 2, names = c("stable","movers"), cex.names = 2)
vioplot(mrres$Mscore[names(yloc)][yloc == 0], mrres$Mscore[names(yloc)][yloc == 1], col = getStockcol()[3], main = "M Score",
        cex.main = 2, names = c("stable","movers"), cex.names = 2)
vioplot(mrres$Rscore[names(yloc)][yloc == 0], mrres$Rscore[names(yloc)][yloc == 1], col = getStockcol()[4], main ="R Score",
        cex.main = 2, names = c("stable","movers"), cex.names = 2)

cor(x = y, yloc, method = "pearson")
cor(x = mrres$Mscore[names(yloc)], yloc, method = "pearson")
cor(x = mrres$Rscore[names(yloc)], yloc, method = "pearson")
```
