## Supplementary Code 1 for "Inferring differential subcellular localisation in comparative spatial proteomics using BANDLE": daviesprocessb.R

```
source("bandleFUN.R")

load("data/davies2018wt.rda")
load("data/davies2018ap4b1.rda")
load("data/davies2018ap4e1.rda")

davies2018wt <- filterNA(davies2018wt)
davies2018ap4b1 <- filterNA(davies2018ap4b1)
davies2018ap4e1 <- filterNA(davies2018ap4e1)

exprs(davies2018wt) <- log2(exprs(davies2018wt))
exprs(davies2018ap4b1) <- log2(exprs(davies2018ap4b1))
exprs(davies2018ap4e1) <- log2(exprs(davies2018ap4e1))

par(mfrow = c(2,3))
plot2D(davies2018wt[,1:5], dims = c(1,2))
plot2D(davies2018wt[,6:10], dims = c(1,2))
plot2D(davies2018ap4b1, dims = c(1,2))
plot2D(davies2018ap4e1[,1:5], dims = c(1,2))
getMarkerClasses(davies2018wt, fcol = "markers")

par(mfrow = c(1,1))
pRoloc::plotDist(davies2018wt, fcol = "markers")

exprs(davies2018wt)

K <- length(getMarkerClasses(davies2018wt))
dir_prior = diag(rep(1, K)) + matrix(0.02, nrow = K, ncol = K)
prior_pred_dir(object = davies2018wt, dir_prior = dir_prior)

pc_prior <- matrix(NA, ncol = 3, K)
pc_prior[seq.int(1:K), ] <- matrix(rep(c(0.05, 60, 100), each = K), ncol = 3)

load("bandleres_daviesb1.rda")

dim(bandleres_daviesb1[[1]]$allocprob$cond11)

par(mfrow = c(3,4))

mycols <- c("blue", "darkgreen", "orange", "purple", "pink", "brown", "gray", "cyan")
mycols <- sapply(mycols, function(x) alpha(x, 0.2))

for (j in seq_along(getMarkerClasses(davies2018ap4b1))) {
  for (i in seq_along(bandleres_daviesb1)){
    if (i ==  1) {
      M <- max(bandleres_daviesb1[[i]]$weights[j,j,]) + 0.05
      plot(bandleres_daviesb1[[i]]$weights[j,j,], col = mycols[[i]], type = "b", pch = 19, ylim = c(0, M), xlab = "mixing proportions")
    } else {
      points(bandleres_daviesb1[[i]]$weights[j,j,], col = mycols[[i]], type = "b", pch = 19)
    }  
  }
}

probdir_daviesb1 <- probsameorganelle(object = davies2018ap4b1, params = bandleres_daviesb1[[2]])

## BANDLE dirichlet
par(mfrow = c(1,1))
y <- 1 - probdir_daviesb1[order(probdir_daviesb1)]/ncol(bandleres_daviesb1[[2]]$allocOut$cond11)
plot(y, pch = 19, ylim = c(0,1), ylab = "Differential Localisation Probability", xlab = "Rank", 
     col = getStockcol()[2], main = "Rank plot from BANDLE", cex.lab = 1.5, cex.main = 2, cex.axis = 1.5)

par(mfrow = c(2,1))
.pca <- plot2D(object = davies2018wt, dims = c(1,4))
addLegend(where = "topleft", object = davies2018wt, ncol = 2)
highlightOnPlot(object = .pca, foi = names(y[1:25]), col = "purple3", cex = 1.5, pch = 19, labels = TRUE)
.pca <- plot2D(object = davies2018ap4b1, dims = c(1,2))
highlightOnPlot(object = .pca, foi = names(y[1:25]), col = "purple3", cex = 1.5, pch = 19, labels = TRUE)

movers <- names(y[1:50])

write.csv(movers, file = "daviesb_movers.csv")

rownames(bandleres_daviesb1[[2]]$alloc$cond11) <- rownames(unknownMSnSet(davies2018wt))
rownames(bandleres_daviesb1[[2]]$alloc$cond21) <- rownames(unknownMSnSet(davies2018wt))

alloc1 <- apply(bandleres_daviesb1[[2]]$alloc$cond11[names(y[1:25]),],1, table)
alloc2 <- apply(bandleres_daviesb1[[2]]$alloc$cond21[names(y[1:25]),],1, table)

qsepres <- QSep(davies2018wt)

par(mfrow = c(3,4))
hypres <- fitGPmaternPC(object = davies2018wt[,1:5], fcol = "markers", materncov = TRUE, nu = 2, hyppar = c(10, 60, 200))

daviesdyn <- c(davies2018wt[ ,1:5],
               davies2018wt[, 6:10],
               davies2018ap4b1[, 1:5],
               davies2018ap4b1[, 6:10])

daviesdyn <- lapply(daviesdyn, function(x) normalise(x, methed = "center.median"))
  
par(mfrow = c(3,4))
pc_prior <- matrix(NA, ncol = 3, K)
pc_prior[seq.int(1:K), ] <- matrix(rep(c(10, 60, 250), each = K), ncol = 3)
hypres <- lapply(daviesdyn, function(x) fitGPmaternPC(x, fcol = "markers", nu = 2, materncov = TRUE, hyppar = pc_prior)) 


save(hypres, file = "davies_hypers2.rda")

load("bandleres_daviesb2.rda")

dim(bandleres_daviesb2[[1]]$allocprob$cond11)


subactin <- fData(davies2018wt)$markers == "Actin binding proteins"
fData(davies2018wt)[rownames(davies2018wt[subactin]), "markers"] <- "unknown"

subactin <- fData(davies2018ap4b1)$markers == "Actin binding proteins"
fData(davies2018ap4b1)[rownames(davies2018ap4b1[subactin]), "markers"] <- "unknown"

subnpc <- fData(davies2018ap4b1)$markers == "Nuclear pore complex"
fData(davies2018ap4b1)[rownames(davies2018ap4b1[subnpc]), "markers"] <- "unknown"

subnpc <- fData(davies2018wt)$markers == "Nuclear pore complex"
fData(davies2018wt)[rownames(davies2018wt[subnpc]), "markers"] <- "unknown"


par(mfrow = c(3,4))

mycols <- c("blue", "darkgreen", "orange", "purple", "pink", "brown", "gray", "cyan")
mycols <- sapply(mycols, function(x) alpha(x, 0.2))

for (j in seq_along(getMarkerClasses(davies2018ap4b1))) {
  for (i in seq_along(bandleres_daviesb2)){
    if (i ==  1) {
      M <- max(bandleres_daviesb2[[i]]$weights[j,j,]) + 0.05
      plot(bandleres_daviesb2[[i]]$weights[j,j,], col = mycols[[i]], type = "b", pch = 19, ylim = c(0, M), xlab = "mixing proportions")
    } else {
      points(bandleres_daviesb2[[i]]$weights[j,j,], col = mycols[[i]], type = "b", pch = 19)
    }  
  }
}

for (j in seq_along(getMarkerClasses(davies2018ap4b1))) {
  for (i in c(1,3,4)){
    if (i ==  1) {
      M <- max(bandleres_daviesb2[[i]]$weights[j,j,]) + 0.05
      plot(bandleres_daviesb2[[i]]$weights[j,j,], col = mycols[[i]], type = "b", pch = 19, ylim = c(0, M), xlab = "mixing proportions")
    } else {
      points(bandleres_daviesb2[[i]]$weights[j,j,], col = mycols[[i]], type = "b", pch = 19)
    }  
  }
}

require(coda)
r_hat <- vector(mode = "list", length = length(getMarkerClasses(davies2018ap4b1)))
for (j in seq_along(getMarkerClasses(davies2018ap4b1))) {
  r_hat[[j]] <- gelman.diag(mcmc.list(mcmc(bandleres_daviesb2[[1]]$weights[j,j,]),
                                      mcmc(bandleres_daviesb2[[3]]$weights[j,j,])), transform = TRUE)
}
unlist(r_hat)

par(mfrow = c(1,2))
rk <- rank(cbind(bandleres_daviesb2[[1]]$weights[j,j,], mcmc(bandleres_daviesb2[[3]]$weights[j,j,])))
hist(rk[1:667], col = alpha("blue", 0.2), breaks = seq.int(from = 1, to = 1334, length.out = 10), main = "rank plot")
hist(rk[667:1334], col = alpha("darkgreen", 0.2),breaks = seq.int(from = 1, to = 1334, length.out = 10), main = "rank plot")

probdir_daviesb2 <- probsameorganelle(object = davies2018ap4b1, params = bandleres_daviesb2[[1]])

## BANDLE dirichlet
par(mfrow = c(1,1))
y <- 1 - probdir_daviesb2[order(probdir_daviesb2)]/ncol(bandleres_daviesb2[[1]]$allocOut$cond11)
plot(y, pch = 19, ylim = c(0,1), ylab = "Differential Localisation Probability", xlab = "Rank", 
     col = getStockcol()[2], main = "Rank plot from BANDLE", cex.lab = 1.5, cex.main = 2, cex.axis = 1.5)

par(mfrow = c(2,1))
.pca <- plot2D(object = davies2018wt, dims = c(1,2), grid = FALSE)
addLegend(where = "bottomright", object = davies2018wt, ncol = 2)
highlightOnPlot(object = .pca, foi = names(y[1:25]), col = "purple3", cex = 1.5, pch = 19, labels = TRUE)
highlightOnPlot(object = .pca, foi = "P04920", col = "purple3", cex = 1.5, pch = 19, labels = TRUE)
.pca <- plot2D(object = davies2018ap4b1, dims = c(1,2))
highlightOnPlot(object = .pca, foi = names(y[1:25]), col = "purple3", cex = 1.5, pch = 19, labels = TRUE)

rownames(bandleres_daviesb2[[2]]$alloc$cond11) <- rownames(unknownMSnSet(davies2018wt))
rownames(bandleres_daviesb2[[2]]$alloc$cond21) <- rownames(unknownMSnSet(davies2018wt))

alloc1 <- apply(bandleres_daviesb2[[2]]$alloc$cond11[names(y[1:25]),],1, table)
alloc2 <- apply(bandleres_daviesb2[[2]]$alloc$cond21[names(y[1:25]),],1, table)

movers <- names(y[1:50])

write.csv(movers, file = "daviesb_movers.csv")

vesprof <- read.csv("davies_vescicleprof.csv")
rownames(vesprof) <- vesprof$Majority.protein.IDs

vescol <- rep(alpha("darkgreen", 0.4), length = nrow(vesprof))
names(vescol) <- rownames(vesprof)
vescol[names(vescol) %in% movers] <- alpha("orange", 0.5)
par(mfrow = c(2,3))
plot(vesprof[,5], vesprof[,6], col = vescol, pch = 19)
plot(vesprof[,6], vesprof[,7], col = vescol, pch = 19)
plot(vesprof[,8], vesprof[,9], col = vescol, pch = 19)
plot(vesprof[,10], vesprof[,11], col = vescol, pch = 19)
plot(vesprof[,12], vesprof[,13], col = vescol, pch = 19)

par(mfrow = c(1,1))
.pcaves <- prcomp(impute::impute.knn(as.matrix(vesprof[,5:13]))$data)
plot(.pcaves$x[,1], .pcaves$x[,3], col = vescol, pch = 19)

rownames(vesprof)[.pcaves$x[,1] > 2.5 & .pcaves$x[,3] < -1]

vesprof_sub <- vesprof[ rownames(vesprof) %in% names(y),]
vescol_sub <- vescol[names(vescol) %in% names(y)]

t.test(vesprof_sub[vescol_sub == vescol_sub[1],12], vesprof_sub[vescol_sub != vescol_sub[1],12])
hist(vesprof_sub[vescol_sub == vescol_sub[1],12], breaks = 30)
plot(vesprof_sub[vescol_sub == vescol_sub[1],12], col = vescol_sub, pch = 19)

## Load davies data
load("C:/Users/OllyC/Desktop/2019-DiffLoc/bandleres_daviesb3.rda")

par(mfrow = c(3,4))

mycols <- c("blue", "darkgreen", "orange", "purple", "pink", "brown", "gray", "cyan")
mycols <- sapply(mycols, function(x) alpha(x, 0.2))

for (j in seq_along(getMarkerClasses(davies2018ap4b1))) {
  for (i in seq_along(bandleres_daviesb3)){
    if (i ==  1) {
      M <- max(bandleres_daviesb3[[i]]$weights[j,j,]) + 0.05
      plot(bandleres_daviesb3[[i]]$weights[j,j,], col = mycols[[i]], type = "b", pch = 19, ylim = c(0, M), xlab = "mixing proportions")
    } else {
      points(bandleres_daviesb3[[i]]$weights[j,j,], col = mycols[[i]], type = "b", pch = 19)
    }  
  }
}

for (j in seq_along(getMarkerClasses(davies2018ap4b1))) {
  for (i in c(2,3,4,5,6)){
    if (i ==  2) {
      M <- max(bandleres_daviesb3[[i]]$weights[j,j,]) + 0.05
      plot(bandleres_daviesb3[[i]]$weights[j,j,], col = mycols[[i]], type = "b", pch = 19, ylim = c(0, M), xlab = "mixing proportions")
    } else {
      points(bandleres_daviesb3[[i]]$weights[j,j,], col = mycols[[i]], type = "b", pch = 19)
    }  
  }
}

require(coda)
r_hat <- vector(mode = "list", length = length(getMarkerClasses(davies2018ap4b1)))
for (j in seq_along(getMarkerClasses(davies2018ap4b1))) {
  r_hat[[j]] <- gelman.diag(mcmc.list(mcmc(bandleres_daviesb3[[2]]$weights[j,j,]),
                                      mcmc(bandleres_daviesb3[[3]]$weights[j,j,])), transform = TRUE)
}
unlist(r_hat)

par(mfrow = c(2,3))
rk <- rank(cbind(bandleres_daviesb3[[2]]$weights[j,j,],
                 bandleres_daviesb3[[3]]$weights[j,j,],
                 bandleres_daviesb3[[4]]$weights[j,j,],
                 bandleres_daviesb3[[5]]$weights[j,j,],
                 bandleres_daviesb3[[6]]$weights[j,j,]))

hist(rk[1:667], col = alpha("blue", 0.2), breaks = seq.int(from = 1, to = length(rk), length.out = 10), main = "rank plot")
hist(rk[668:1334], col = alpha("darkgreen", 0.2),breaks = seq.int(from = 1, to = length(rk), length.out = 10), main = "rank plot")
hist(rk[1335:2001], col = mycols[3],breaks = seq.int(from = 1, to = length(rk), length.out = 10), main = "rank plot")
hist(rk[2001:2668], col = mycols[4],breaks = seq.int(from = 1, to = length(rk), length.out = 10), main = "rank plot")
hist(rk[2668:3335], col = mycols[5],breaks = seq.int(from = 1, to = length(rk), length.out = 10), main = "rank plot")

rk <- rank(cbind(bandleres_daviesb3[[2]]$weights[j,j,],
                 bandleres_daviesb3[[4]]$weights[j,j,],
                 bandleres_daviesb3[[5]]$weights[j,j,]))

par(mfrow = c(1,3))
hist(rk[1:667], col = alpha("blue", 0.2), breaks = seq.int(from = 1, to = length(rk), length.out = 10), main = "rank plot", xlab = "rank")
hist(rk[668:1334], col = alpha("darkgreen", 0.2),breaks = seq.int(from = 1, to = length(rk), length.out = 10), main = "rank plot", xlab = "rank")
hist(rk[1335:2001], col = mycols[3],breaks = seq.int(from = 1, to = length(rk), length.out = 10), main = "rank plot", xlab = "rank")

probdir_daviesb3 <- probsameorganelle(object = davies2018ap4b1, params = bandleres_daviesb3[[2]])

## BANDLE dirichlet
par(mfrow = c(1,1))
y <- 1 - probdir_daviesb3[order(probdir_daviesb3)]/ncol(bandleres_daviesb3[[1]]$allocOut$cond11)
plot(y, pch = 19, ylim = c(0,1), ylab = "Differential Localisation Probability", xlab = "Rank", 
     col = getStockcol()[2], main = "Rank plot from BANDLE", cex.lab = 1.5, cex.main = 2, cex.axis = 1.5)

par(mfrow = c(2,1))
.pca <- plot2D(object = davies2018wt, dims = c(1,2), grid = FALSE)
addLegend(where = "bottomright", object = davies2018wt, ncol = 2)
highlightOnPlot(object = .pca, foi = names(y[1:25]), col = "purple3", cex = 1.5, pch = 19, labels = TRUE)
.pca <- plot2D(object = davies2018ap4b1, dims = c(1,2))
highlightOnPlot(object = .pca, foi = names(y[1:25]), col = "purple3", cex = 1.5, pch = 19, labels = TRUE)

rownames(bandleres_daviesb3[[2]]$alloc$cond11) <- rownames(unknownMSnSet(davies2018wt))
rownames(bandleres_daviesb3[[2]]$alloc$cond21) <- rownames(unknownMSnSet(davies2018wt))

alloc1 <- apply(bandleres_daviesb3[[2]]$alloc$cond11[names(y[1:50]),],1, table)
alloc2 <- apply(bandleres_daviesb3[[2]]$alloc$cond21[names(y[1:50]),],1, table)

movers <- names(y[1:50])

write.csv(movers, file = "daviesb_movers.csv")


candmoves <- substr(names(y[1:50]), start = 1, stop = 6)
library(UniProt.ws)

species <- 9606
columns <- c("SEQUENCE", "SUBCELLULAR-LOCATIONS", "PROTEIN-NAMES")

up <- UniProt.ws(taxId=species)

keytype <- "UNIPROTKB"

# example keys
keys <- str_sub(rownames(davies2018wt), start = 1, end = 6)

protein_seq <- UniProt.ws::select(up, keys, columns, keytype) %>%
  mutate(motif=grepl(pattern='Y..[AVILMPFYW]', SEQUENCE))  

uniprotlocal <- protein_seq[,c(1,3)]
which.nuc <- grepl(uniprotlocal$`SUBCELLULAR-LOCATIONS`, pattern = " nucle")
which.act <- grepl(uniprotlocal$`SUBCELLULAR-LOCATIONS`, pattern = " actin")
names(which.nuc) <- uniprotlocal[,1]
names(which.act) <- uniprotlocal[,1]
to.filteruni <- names(which.nuc)[which.nuc]
to.filteract <- names(which.act)[which.act]


df <- read.csv(file = 'davies_hits.tsv', sep = '\t', header = FALSE, skip = 1)
write.csv(df, file = "davies_hits.csv")

write.csv(y, file = "davies_all.csv")

par(mfrow = c(2,3))
vesprof <- read.csv("davies_vescicleprof.csv")
rownames(vesprof) <- vesprof$Majority.protein.IDs

vescol <- rep(alpha("darkgreen", 0.4), length = nrow(vesprof))
names(vescol) <- rownames(vesprof)
vescol[names(vescol) %in% movers] <- alpha("orange", 0.5)
par(mfrow = c(2,3))
plot(vesprof[,5], vesprof[,6], col = vescol, pch = 19)
plot(vesprof[,6], vesprof[,7], col = vescol, pch = 19)
plot(vesprof[,8], vesprof[,9], col = vescol, pch = 19)
plot(vesprof[,10], vesprof[,11], col = vescol, pch = 19)
plot(vesprof[,12], vesprof[,13], col = vescol, pch = 19)

par(mfrow = c(1,1))
.pcaves <- prcomp(impute::impute.knn(as.matrix(vesprof[,5:13]))$data)
plot(.pcaves$x[,1], .pcaves$x[,3], col = vescol, pch = 19)

rownames(vesprof)[.pcaves$x[,1] > 2.5 & .pcaves$x[,3] < -1]

vesprof_sub <- vesprof[ rownames(vesprof) %in% names(y),]
vescol_sub <- vescol[names(vescol) %in% names(y)]

t.test(vesprof_sub[vescol_sub == vescol_sub[1],12], vesprof_sub[vescol_sub != vescol_sub[1],12])
boxplot(vesprof_sub[vescol_sub == vescol_sub[1],12], vesprof_sub[vescol_sub != vescol_sub[1],12])
hist(vesprof_sub[vescol_sub == vescol_sub[1],12], breaks = 30)
plot(vesprof_sub[vescol_sub == vescol_sub[1],12], col = vescol_sub, pch = 19)

par(mfrow = c(1,2))
#knockout 2
plot(vesprof_sub[,10], vesprof_sub[,11], col = vescol_sub, pch = 19)
abline(lm(vesprof_sub[,11] ~ vesprof_sub[,10]), col = "purple3", lwd = 4)

#knockout 1
plot(vesprof_sub[,8], vesprof_sub[,9], col = vescol_sub, pch = 19)
abline(lm(vesprof_sub[,9] ~ vesprof_sub[,8]), col = "purple3", lwd = 4)

par(mfrow = c(1,2))
#knockdowns
plot(vesprof_sub[,5], vesprof_sub[,6], col = vescol_sub, pch = 19)
abline(lm(vesprof_sub[,6] ~ vesprof_sub[,5]), col = "purple3", lwd = 4)

#knockdowns
plot(vesprof_sub[,6], vesprof_sub[,7], col = vescol_sub, pch = 19)
abline(lm(vesprof_sub[,7] ~ vesprof_sub[,6]), col = "purple3", lwd = 4)

## movers
classalloc <- sapply(alloc1, names)
classmostlike <- sapply(alloc1, which.max)

class <- vector(length = length(classalloc))
for (j in seq_along(classalloc)){
  class[j] <- classalloc[[j]][classmostlike[j]]
}
names(class) <- names(y[1:50])

# filter out proteins allocated to the large protein complex compartment
filteredmovers <- class[class != "9"]

par(mfrow = c(5,5))
for (j in seq_along(filteredmovers)) {
mydata <- t(rbind(exprs(davies2018wt[names(filteredmovers[j]),1:5]),
                  exprs(davies2018wt[names(filteredmovers[j]),6:10])))
M <- max(mydata) + 0.5
m <- min(mydata) - 0.5
matplot(mydata, col = mycols[1], pch = 19, lwd = 3, lty = 1,
        ylab = "abundance", type = "b", ylim = c(m, M), main = paste0(names(filteredmovers[j])))
matplot(t(rbind(exprs(davies2018ap4b1[names(filteredmovers[j]),1:5]),
                exprs(davies2018ap4b1[names(filteredmovers[j]),6:10]))), col = mycols[2], pch = 19, lwd = 3, lty = 1,
        ylab = "abundance", type = "b", add = TRUE)
}

write.csv(filteredmovers, file = "filteredmoversdavies.csv")

# candidate proteins remove all isoform annotations

rownames(hpa) <- hpa[,1]
candmoves <- substr(names(y[1:50]), start = 1, stop = 6)
nuclikeannots <- levels(hpa[,2])[c(18:24)]
is.nuc <- hpa[hpa[,1] %in% candmoves,2] %in% nuclikeannots
tofilter <- hpa[hpa[,1] %in% candmoves,1][is.nuc]

filteredmovershpa <- y[1:50][!(candmoves %in% tofilter)]

par(mfrow = c(5,5))
for (j in seq_along(filteredmovershpa)) {
  mydata <- t(rbind(exprs(davies2018wt[names(filteredmovershpa[j]),1:5]),
                    exprs(davies2018wt[names(filteredmovershpa[j]),6:10])))
  M <- max(mydata) + 0.5
  m <- min(mydata) - 0.5
  matplot(mydata, col = mycols[1], pch = 19, lwd = 3, lty = 1,
          ylab = "abundance", type = "b", ylim = c(m, M), main = paste0(names(filteredmovershpa[j])))
  matplot(t(rbind(exprs(davies2018ap4b1[names(filteredmovershpa[j]),1:5]),
                  exprs(davies2018ap4b1[names(filteredmovershpa[j]),6:10]))), col = mycols[2], pch = 19, lwd = 3, lty = 1,
          ylab = "abundance", type = "b", add = TRUE)
}

write.csv(filteredmovershpa, file = "filteredmoversdavieshpa.csv")


par(mfrow = c(5,5))
for (j in seq_along(y[1:50])) {
  mydata <- t(rbind(exprs(davies2018wt[names(y[j]),1:5]),
                    exprs(davies2018wt[names(y[j]),6:10])))
  M <- max(mydata) + 0.5
  m <- min(mydata) - 0.5
  matplot(mydata, col = mycols[1], pch = 19, lwd = 3, lty = 1,
          ylab = "abundance", type = "b", ylim = c(m, M), main = paste0(names(y[j])))
  matplot(t(rbind(exprs(davies2018ap4b1[names(y[j]),1:5]),
                  exprs(davies2018ap4b1[names(y[j]),6:10]))), col = mycols[2], pch = 19, lwd = 3, lty = 1,
          ylab = "abundance", type = "b", add = TRUE)
}


library(UniProt.ws)

species <- 9606
columns <- c("SEQUENCE", "SUBCELLULAR-LOCATIONS", "PROTEIN-NAMES")

up <- UniProt.ws(taxId=species)

keytype <- "UNIPROTKB"

# example keys
keys <- str_sub(rownames(davies2018wt), start = 1, end = 6)

protein_seq <- UniProt.ws::select(up, keys, columns, keytype) %>%
  mutate(motif=grepl(pattern='Y..[AVILMPFYW]', SEQUENCE))  

uniprotlocal <- protein_seq[,c(1,3)]
which.nuc <- grepl(uniprotlocal$`SUBCELLULAR-LOCATIONS`, pattern = " nucle")
which.act <- grepl(uniprotlocal$`SUBCELLULAR-LOCATIONS`, pattern = " actin")
names(which.nuc) <- uniprotlocal[,1]
names(which.act) <- uniprotlocal[,1]
to.filteruni <- names(which.nuc)[which.nuc]
to.filteract <- names(which.act)[which.act]

filteredmovers <- names(y[1:50][!(candmoves %in% c(to.filteruni, to.filteract))])
write.csv(candmoves[!(candmoves %in% to.filteruni)], file = "filtermoversdaviesuniprot.csv")


par(mfrow = c(5,5))
for (j in seq_along(filteredmovers)) {
  mydata <- t(rbind(exprs(davies2018wt[filteredmovers[j],1:5]),
                    exprs(davies2018wt[filteredmovers[j],6:10])))
  M <- max(mydata) + 0.5
  m <- min(mydata) - 0.5
  matplot(mydata, col = mycols[1], pch = 19, lwd = 3, lty = 1,
          ylab = "abundance", type = "b", ylim = c(m, M), main = paste0(filteredmovers[j]))
  matplot(t(rbind(exprs(davies2018ap4b1[filteredmovers[j],1:5]),
                  exprs(davies2018ap4b1[filteredmovers[j],6:10]))), col = mycols[2], pch = 19, lwd = 3, lty = 1,
          ylab = "abundance", type = "b", add = TRUE)
}

par(mfrow = c(3,3))
for (j in seq_along(filteredmovers)[c(1,2,17,15,23,24,9,7,13)]) {
  mydata <- t(rbind(exprs(davies2018wt[filteredmovers[j],1:5]),
                    exprs(davies2018wt[filteredmovers[j],6:10])))
  M <- max(mydata) + 0.5
  m <- min(mydata) - 0.5
  matplot(mydata, col = mycols[1], pch = 19, lwd = 3, lty = 1,
          ylab = "abundance", type = "b", ylim = c(m, M), main = paste0(filteredmovers[j]))
  matplot(t(rbind(exprs(davies2018ap4b1[filteredmovers[j],1:5]),
                  exprs(davies2018ap4b1[filteredmovers[j],6:10]))), col = mycols[2], pch = 19, lwd = 3, lty = 1,
          ylab = "abundance", type = "b", add = TRUE)
}

Q676U5 - ATG16L1
Q86YS7 - C2CD5
Q9H1H9 - KIF13A
O95857 - TSPAN13
Q8N129 - CNPY4
Q10559 - BST2
Q8N511 - TMEM199
Q9NRX5 - Serinc1
P51159


## analysis of dataset 4

## Load davies data
load("C:/Users/OllyC/Desktop/2019-DiffLoc/bandleres_daviesb4.rda")

par(mfrow = c(3,4))

mycols <- c("blue", "darkgreen", "orange", "purple", "pink", "brown", "gray", "cyan")
mycols <- sapply(mycols, function(x) alpha(x, 0.2))

for (j in seq_along(getMarkerClasses(davies2018ap4b1))) {
  for (i in seq_along(bandleres_daviesb4)){
    if (i ==  1) {
      M <- max(bandleres_daviesb4[[i]]$weights[j,j,]) + 0.05
      plot(bandleres_daviesb4[[i]]$weights[j,j,], col = mycols[[i]], type = "b", pch = 19, ylim = c(0, M), xlab = "mixing proportions")
    } else {
      points(bandleres_daviesb4[[i]]$weights[j,j,], col = mycols[[i]], type = "b", pch = 19)
    }  
  }
}

for (j in seq_along(getMarkerClasses(davies2018ap4b1))) {
  for (i in c(2,3,4,5,6)){
    if (i ==  2) {
      M <- max(bandleres_daviesb4[[i]]$weights[j,j,]) + 0.05
      plot(bandleres_daviesb4[[i]]$weights[j,j,], col = mycols[[i]], type = "b", pch = 19, ylim = c(0, M), xlab = "mixing proportions")
    } else {
      points(bandleres_daviesb4[[i]]$weights[j,j,], col = mycols[[i]], type = "b", pch = 19)
    }  
  }
}

require(coda)
r_hat <- vector(mode = "list", length = length(getMarkerClasses(davies2018ap4b1)))
for (j in seq_along(getMarkerClasses(davies2018ap4b1))) {
  r_hat[[j]] <- gelman.diag(mcmc.list(mcmc(bandleres_daviesb4[[1]]$weights[j,j,]),
                                      mcmc(bandleres_daviesb4[[3]]$weights[j,j,])), transform = TRUE)
}
unlist(r_hat)

par(mfrow = c(2,3))
rk <- rank(cbind(bandleres_daviesb4[[1]]$weights[j,j,],
                 bandleres_daviesb4[[3]]$weights[j,j,],
                 bandleres_daviesb4[[4]]$weights[j,j,],
                 bandleres_daviesb4[[5]]$weights[j,j,],
                 bandleres_daviesb4[[6]]$weights[j,j,]))

hist(rk[1:667], col = alpha("blue", 0.2), breaks = seq.int(from = 1, to = length(rk), length.out = 10), main = "rank plot")
hist(rk[668:1334], col = alpha("darkgreen", 0.2),breaks = seq.int(from = 1, to = length(rk), length.out = 10), main = "rank plot")
hist(rk[1335:2001], col = mycols[3],breaks = seq.int(from = 1, to = length(rk), length.out = 10), main = "rank plot")
hist(rk[2001:2668], col = mycols[4],breaks = seq.int(from = 1, to = length(rk), length.out = 10), main = "rank plot")
hist(rk[2668:3335], col = mycols[5],breaks = seq.int(from = 1, to = length(rk), length.out = 10), main = "rank plot")

rk <- rank(cbind(bandleres_daviesb4[[1]]$weights[j,j,],
                 bandleres_daviesb4[[3]]$weights[j,j,],
                 bandleres_daviesb4[[6]]$weights[j,j,]))

par(mfrow = c(1,3))
hist(rk[1:667], col = alpha("blue", 0.2), breaks = seq.int(from = 1, to = length(rk), length.out = 10), main = "rank plot", xlab = "rank")
hist(rk[668:1334], col = alpha("darkgreen", 0.2),breaks = seq.int(from = 1, to = length(rk), length.out = 10), main = "rank plot", xlab = "rank")
hist(rk[1335:2001], col = mycols[3],breaks = seq.int(from = 1, to = length(rk), length.out = 10), main = "rank plot", xlab = "rank")

probdir_daviesb4 <- probsameorganelle(object = davies2018ap4b1, params = bandleres_daviesb4[[1]])

## BANDLE dirichlet
par(mfrow = c(1,1))
y <- 1 - probdir_daviesb4[order(probdir_daviesb4)]/ncol(bandleres_daviesb4[[1]]$allocOut$cond11)
plot(y, pch = 19, ylim = c(0,1), ylab = "Differential Localisation Probability", xlab = "Rank", 
     col = getStockcol()[2], main = "Rank plot from BANDLE", cex.lab = 1.5, cex.main = 2, cex.axis = 1.5)

par(mfrow = c(2,1))
.pca <- plot2D(object = davies2018wt, dims = c(1,2), grid = FALSE)
addLegend(where = "bottomright", object = davies2018wt, ncol = 2)
highlightOnPlot(object = .pca, foi = names(y[1:25]), col = "purple3", cex = 1.5, pch = 19, labels = TRUE)
.pca <- plot2D(object = davies2018ap4b1, dims = c(1,2))
highlightOnPlot(object = .pca, foi = names(y[1:25]), col = "purple3", cex = 1.5, pch = 19, labels = TRUE)

rownames(bandleres_daviesb4[[3]]$alloc$cond11) <- rownames(unknownMSnSet(davies2018wt))
rownames(bandleres_daviesb4[[3]]$alloc$cond21) <- rownames(unknownMSnSet(davies2018wt))

alloc1 <- apply(bandleres_daviesb4[[3]]$alloc$cond11[names(y[1:50]),],1, table)
alloc2 <- apply(bandleres_daviesb4[[3]]$alloc$cond21[names(y[1:50]),],1, table)

movers <- names(y[1:50])

write.csv(y, file = "davies_all2.csv")

candmoves <- substr(names(y[1:80]), start = 1, stop = 6)

write.csv(candmoves[!(candmoves %in% c(to.filteruni, to.filteract))], file = "filtermoversdaviesuniprot.csv")

filteredmovers <- names(y[1:80][!(candmoves %in% c(to.filteruni, to.filteract))])
par(mfrow = c(5,5))
for (j in seq_along(filteredmovers)) {
  mydata <- t(rbind(exprs(davies2018wt[filteredmovers[j],1:5]),
                    exprs(davies2018wt[filteredmovers[j],6:10])))
  M <- max(mydata) + 0.5
  m <- min(mydata) - 0.5
  matplot(mydata, col = mycols[1], pch = 19, lwd = 3, lty = 1,
          ylab = "abundance", type = "b", ylim = c(m, M), main = paste0(filteredmovers[j]))
  matplot(t(rbind(exprs(davies2018ap4b1[filteredmovers[j],1:5]),
                  exprs(davies2018ap4b1[filteredmovers[j],6:10]))), col = mycols[2], pch = 19, lwd = 3, lty = 1,
          ylab = "abundance", type = "b", add = TRUE)
}
```
