## Supplementary Code 1 for "Inferring differential subcellular localisation in comparative spatial proteomics using BANDLE": daviesprocesse.R

```
load("data/davies2018wt.rda")
load("data/davies2018ap4b1.rda")
load("data/davies2018ap4e1.rda")
load("data/davies_hypers.rda")


davies2018wt <- filterNA(davies2018wt)
davies2018ap4b1 <- filterNA(davies2018ap4b1)
davies2018ap4e1 <- filterNA(davies2018ap4e1)

exprs(davies2018wt) <- log2(exprs(davies2018wt))
exprs(davies2018ap4b1) <- log2(exprs(davies2018ap4b1))
exprs(davies2018ap4e1) <- log2(exprs(davies2018ap4e1))

par(mfrow = c(2,3))
plot2D(davies2018wt[,1:5], dims = c(1,2))
plot2D(davies2018wt[,6:10], dims = c(1,2))
plot2D(davies2018ap4b1, dims = c(1,2))
plot2D(davies2018ap4e1[,1:5], dims = c(1,2))
getMarkerClasses(davies2018wt, fcol = "markers")

par(mfrow = c(1,1))
pRoloc::plotDist(davies2018wt, fcol = "markers")

exprs(davies2018wt)

subactin <- fData(davies2018wt)$markers == "Actin binding proteins"
fData(davies2018wt)[rownames(davies2018wt[subactin]), "markers"] <- "unknown"


subactin <- fData(davies2018ap4e1)$markers == "Actin binding proteins"
fData(davies2018ap4e1)[rownames(davies2018ap4e1[subactin]), "markers"] <- "unknown"

K <- length(getMarkerClasses(davies2018wt))
dir_prior = diag(rep(1, K)) + matrix(0.02, nrow = K, ncol = K)
prior_pred_dir(object = davies2018wt, dir_prior = dir_prior)

daviesdynE <- c(davies2018wt[ ,1:5],
               davies2018wt[, 6:10],
               davies2018ap4e1[, 1:5],
               davies2018ap4e1[, 6:10])

par(mfrow = c(3,4))
pc_prior <- matrix(NA, ncol = 3, K)
pc_prior[seq.int(1:K), ] <- matrix(rep(c(10, 60, 200), each = K), ncol = 3)
daviese_hypres <- lapply(daviesdynE, function(x) fitGPmaternPC(x, fcol = "markers", nu = 2, hyppar = pc_prior))


save(daviese_hypres, file = "daviese_hypers.rda")

## load data
load("C:/Users/OllyC/Desktop/2019-DiffLoc/bandleres_daviese1.rda")

par(mfrow = c(3,4))

mycols <- c("blue", "darkgreen", "orange", "purple", "pink", "brown", "gray", "cyan")
mycols <- sapply(mycols, function(x) alpha(x, 0.2))

for (j in seq_along(getMarkerClasses(davies2018ap4e1))) {
  for (i in seq_along(bandleres_daviese1)){
    if (i ==  1) {
      M <- max(bandleres_daviese1[[i]]$weights[j,j,]) + 0.05
      plot(bandleres_daviese1[[i]]$weights[j,j,], col = mycols[[i]], type = "b", pch = 19, ylim = c(0, M), xlab = "mixing proportions")
    } else {
      points(bandleres_daviese1[[i]]$weights[j,j,], col = mycols[[i]], type = "b", pch = 19)
    }  
  }
}

for (j in seq_along(getMarkerClasses(davies2018ap4e1))) {
  for (i in c(2,3,4,5,6)){
    if (i ==  2) {
      M <- max(bandleres_daviese1[[i]]$weights[j,j,]) + 0.05
      plot(bandleres_daviese1[[i]]$weights[j,j,], col = mycols[[i]], type = "b", pch = 19, ylim = c(0, M), xlab = "mixing proportions")
    } else {
      points(bandleres_daviese1[[i]]$weights[j,j,], col = mycols[[i]], type = "b", pch = 19)
    }  
  }
}

require(coda)
r_hat <- vector(mode = "list", length = length(getMarkerClasses(davies2018ap4e1)))
for (j in seq_along(getMarkerClasses(davies2018ap4e1))) {
  r_hat[[j]] <- gelman.diag(mcmc.list(mcmc(bandleres_daviese1[[1]]$weights[j,j,]),
                                      mcmc(bandleres_daviese1[[3]]$weights[j,j,])), transform = TRUE)
}
unlist(r_hat)

par(mfrow = c(2,3))
rk <- rank(cbind(bandleres_daviese1[[1]]$weights[j,j,],
                 bandleres_daviese1[[2]]$weights[j,j,],
                 bandleres_daviese1[[3]]$weights[j,j,],
                 bandleres_daviese1[[4]]$weights[j,j,],
                 bandleres_daviese1[[5]]$weights[j,j,],
                 bandleres_daviese1[[6]]$weights[j,j,]))

hist(rk[1:667], col = alpha("blue", 0.2), breaks = seq.int(from = 1, to = length(rk), length.out = 10), main = "rank plot")
hist(rk[668:1334], col = alpha("darkgreen", 0.2),breaks = seq.int(from = 1, to = length(rk), length.out = 10), main = "rank plot")
hist(rk[1335:2001], col = mycols[3],breaks = seq.int(from = 1, to = length(rk), length.out = 10), main = "rank plot")
hist(rk[2001:2668], col = mycols[4],breaks = seq.int(from = 1, to = length(rk), length.out = 10), main = "rank plot")
hist(rk[2668:3335], col = mycols[5],breaks = seq.int(from = 1, to = length(rk), length.out = 10), main = "rank plot")
hist(rk[3336:length(rk)], col = mycols[6],breaks = seq.int(from = 1, to = length(rk), length.out = 10), main = "rank plot")

rk <- rank(cbind(bandleres_daviese1[[1]]$weights[j,j,],
                 bandleres_daviese1[[4]]$weights[j,j,],
                 bandleres_daviese1[[6]]$weights[j,j,]))

par(mfrow = c(1,3))
hist(rk[1:667], col = alpha("blue", 0.2), breaks = seq.int(from = 1, to = length(rk), length.out = 10), main = "rank plot", xlab = "rank")
hist(rk[668:1334], col = alpha("darkgreen", 0.2),breaks = seq.int(from = 1, to = length(rk), length.out = 10), main = "rank plot", xlab = "rank")
hist(rk[1335:2001], col = mycols[3],breaks = seq.int(from = 1, to = length(rk), length.out = 10), main = "rank plot", xlab = "rank")


probdir_daviese1 <- probsameorganelle(object = davies2018ap4e1, params = bandleres_daviese1[[1]])

## BANDLE dirichlet
par(mfrow = c(1,1))
y <- 1 - probdir_daviese1[order(probdir_daviese1)]/ncol(bandleres_daviese1[[1]]$allocOut$cond11)
plot(y, pch = 19, ylim = c(0,1), ylab = "Differential Localisation Probability", xlab = "Rank", 
     col = getStockcol()[2], main = "Rank plot from BANDLE", cex.lab = 1.5, cex.main = 2, cex.axis = 1.5)

par(mfrow = c(2,1))
.pca <- plot2D(object = davies2018wt, dims = c(1,2), grid = FALSE)
addLegend(where = "bottomright", object = davies2018wt, ncol = 2)
highlightOnPlot(object = .pca, foi = names(y[1:25]), col = "purple3", cex = 1.5, pch = 19, labels = TRUE)
.pca <- plot2D(object = davies2018ap4e1, dims = c(1,2))
highlightOnPlot(object = .pca, foi = names(y[1:25]), col = "purple3", cex = 1.5, pch = 19, labels = TRUE)

rownames(bandleres_daviese1[[1]]$alloc$cond11) <- rownames(unknownMSnSet(davies2018wt))
rownames(bandleres_daviese1[[1]]$alloc$cond21) <- rownames(unknownMSnSet(davies2018wt))

alloc1 <- apply(bandleres_daviese1[[1]]$alloc$cond11[names(y[1:50]),],1, table)
alloc2 <- apply(bandleres_daviese1[[1]]$alloc$cond21[names(y[1:50]),],1, table)

movers <- names(y[1:50])

write.csv(movers, file = "daviese_movers.csv")

bmovers <- substr(bmovers[,2], start = 1, stop = 6)
emovers <- substr(movers, start = 1, stop = 6)

davies_hits <- read.csv(file = "davies_all.csv")

davies_reducednames <- substr(davies_hits[,1], start = 1, stop = 6)


bothbe <- davies_hits[,2] *y[as.character(davies_hits[,1])]
dbhits <- bothbe[order(bothbe, decreasing = TRUE)]

write.csv(names(dbhits), file = "daviesbothhits.csv")


par(mfrow = c(5,5))
for (j in seq_along(y[1:50])) {
  mydata <- t(rbind(exprs(davies2018wt[names(y[j]),1:5]),
                    exprs(davies2018wt[names(y[j]),6:10])))
  M <- max(mydata) + 0.5
  m <- min(mydata) - 0.5
  matplot(mydata, col = mycols[1], pch = 19, lwd = 3, lty = 1,
          ylab = "abundance", type = "b", ylim = c(m, M), main = paste0(names(y[j])))
  matplot(t(rbind(exprs(davies2018ap4e1[names(y[j]),1:5]),
                  exprs(davies2018ap4e1[names(y[j]),6:10]))), col = mycols[2], pch = 19, lwd = 3, lty = 1,
          ylab = "abundance", type = "b", add = TRUE)
}
```
