## Supplementary Code 1 for "Inferring differential subcellular localisation in comparative spatial proteomics using BANDLE": diffloc_sim_wild_batch_sys.R

```
## Results simulations
setwd("C:/Users/OllyC/Desktop/2019-DiffLoc")
require(pRoloc)
require(pRolocdata)


# functions requried
source("diffLoc.R")
source("diffLocUtils.R")
source("fitGPmatern.R")
source("fitGPmaternPC.R")
source("simDynamic.R")
source("metropolisGP.R")
source("MRmethod.R")
data("tan2009r1")

## 
require(pROC)
ROCresBayes <- list()
ROCresDir <- list()
ROCresmr <- list()
probpgres <- list()
probdirres <- list()
mrreslist <- list()
perm1_nameslist <- list()

pc_prior <- matrix(NA, ncol = 3, 11)
pc_prior[seq.int(1:11),] <- matrix(rep(c(0.05,60,100), each = 11), ncol = 3)

for (l in seq.int(10)) {
  # Make required data
  
  simres <- sim_dynamic(object = tan2009r1, method = "wild", batch = "systematic")
  
  tan2009rep <- simres$lopitrep
  perm1_names <- simres$perm1_names
  
  # computational methods
  mrres <- mrMethod(c(tan2009rep))
  par(mfrow = c(3,4))
  resdynpg <- diffLoc(object_cond1 = c(tan2009rep[1:3]), object_cond2 = tan2009rep[4:6], numIter = 5000, burnin = 1000,
                      pg = TRUE, hypLearn = "LBFGS", tau = 0.3, maternCov = TRUE, nu = 2, pc_prior = pc_prior)
  par(mfrow = c(3,4))
  resdyndir <- diffLoc(object_cond1 = c(tan2009rep[1:3]), object_cond2 = c(tan2009rep[4:6]),
                       numIter = 5000, burnin = 1000, pg = FALSE, hypLearn = "LFBGS", 
                       maternCov = TRUE, nu = 2, pc_prior = pc_prior)
  
  probpg <- probsameorganelle(object = tan2009r1, params = resdynpg)
  probdir <- probsameorganelle(object = tan2009r1, params = resdyndir)
  
  outcome <- matrix(0, ncol = 1, nrow = length(probpg))
  names(outcome) <- names(probpg)
  outcome[perm1_names] <- 1
  prediction <- probpg/ncol(resdynpg$allocOut[[1]])
  par(mfrow = c(1,1))
  
  ROC_bayes <- roc(outcome, 1 - prediction)
  ROC_mr <- roc(outcome, mrres$MRscore[names(outcome)], direction = "<")
  plot(ROC_bayes, type = "b", col = "darkgreen", lwd = 1, pch = 19, asp = NA, legacy.axes = T)
  lines(ROC_mr, type = "b", col = "blue", lwd = 1, pch = 19, asp = NA)
  
  outcome2 <- matrix(0, ncol = 1, nrow = length(probdir))
  names(outcome2) <- names(probdir)
  outcome2[perm1_names] <- 1
  prediction2 <- probdir/ncol(resdyndir$allocOut[[1]])
  ROC_dir <- roc(outcome2, 1 - prediction2)
  lines(ROC_dir,type = "b", col = "orange", lwd = 1, pch = 19, asp = NA)
  
  legend("bottomright", c("MR", "Dirichlet", "Polya-Gamma"), col = c("blue", "orange", "darkgreen"), pch = 19)
  
  
  ROCresBayes[[l]] <- ROC_bayes
  ROCresDir[[l]] <- ROC_dir
  ROCresmr[[l]] <- ROC_mr
  probpgres[[l]] <- probpg
  probdirres[[l]] <- probdir
  mrreslist[[l]] <- mrres
  perm1_nameslist[[l]] <- perm1_names
  
}

sim_results <- list(ROCresBayes = ROCresBayes,
                    ROCresDir = ROCresDir,
                    ROCresmr = ROCresmr,
                    probpgres = probpgres,
                    probdirres = probdirres,
                    mrreslist = mrreslist,
                    perm1_nameslist = perm1_nameslist)

save(sim_results, file = "diffloc_sim_wild_batch_sys.rda")
```
