## Supplementary methods for "Inferring differential subcellular localisation in comparative spatial proteomics using BANDLE"

### 1 Supplementary

#### 1.1 Appendix 1: BANDLE plate diagram

The parameter symbols are described by the following. The index $i = 1, .., N$ is over proteins, the index $r = 1, ..., R$ over replicates, the index $d = 1, 2$ over datasets, the index $k = 1, ..., K$ over subcellular niches. First, $x_{ird}, d = 1, 2$ is the abundance of protein $i$ in replicate $r$ for dataset $d$. Secondly, $z_{i,d}$ the the latent allocation of protein $i$ in dataset $d$. Then $\phi_{ir}$ is the outlier indicator for protein $i$ in replicate $r$; $\mu_{krd}$ is the regression function for subcellular niche $k$, in replicate $r$ for dataset $d$. $\lambda$ is a vector of hyperparameters for the penalised complexity priors. $e_d$ is the prior outlier probability in dataset $d$. $u$ and $v$ are the hyperameters of the prior beta distribution on the outlier probability. $\pi$ is a matrix where the $(i, j)^{th}$ entry corresponds to the prior probability of localising to subcellular niche $i$ in dataset 1 and subcellular niche $j$ in dataset 2. $\alpha$ denotes the hyperparameter of the matrix Dirichlet prior on $\pi$. $M$ and $V$ are the mean and variance of the data, respectively.

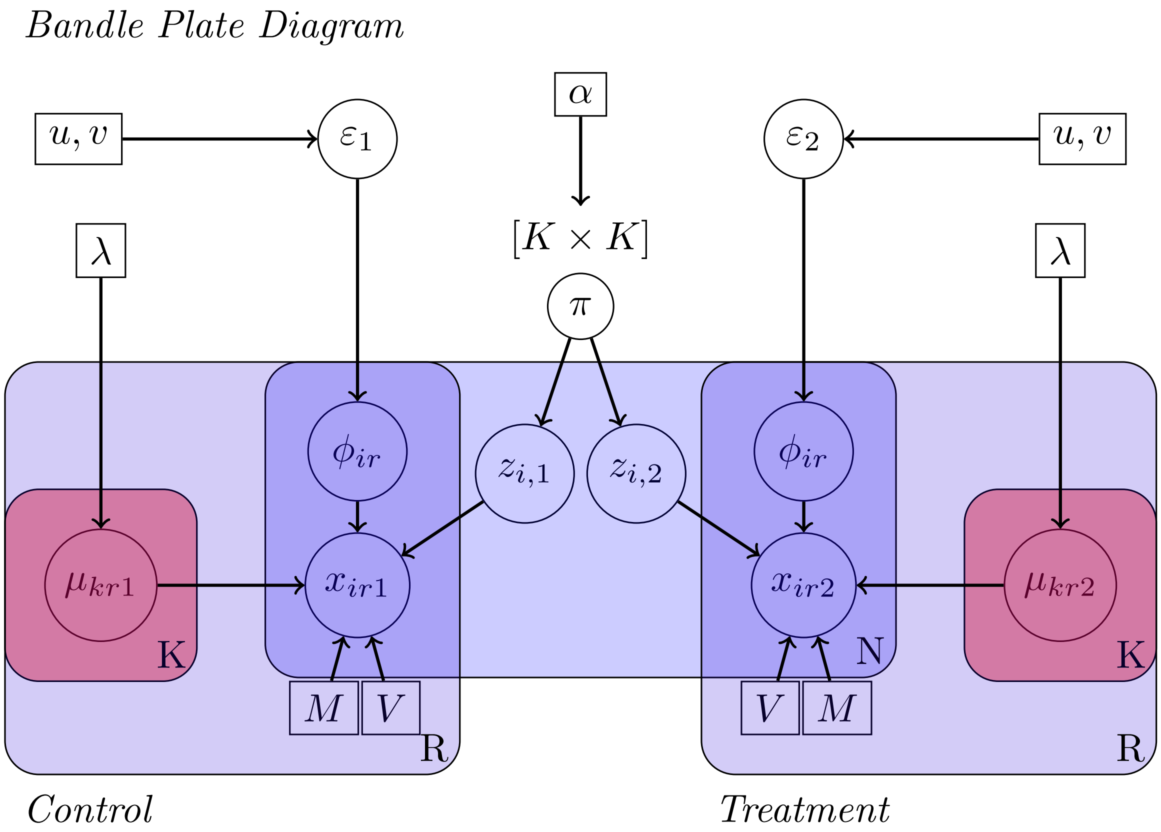

Figure 1: The BANDLE model in plate diagram notation

#### 1.2 Appendix 2: Additional Simulations

We perform additional simulations comparing the MR approach (2016 and 2017) to BANDLE. The simulation scenarios are the same as performed in the main text. However, we start from the LOPIT-DC dataset of Geladaki *et al.* (2019), instead. The conclusion are as for the main text that BANDLE significantly outperforms the MR method.

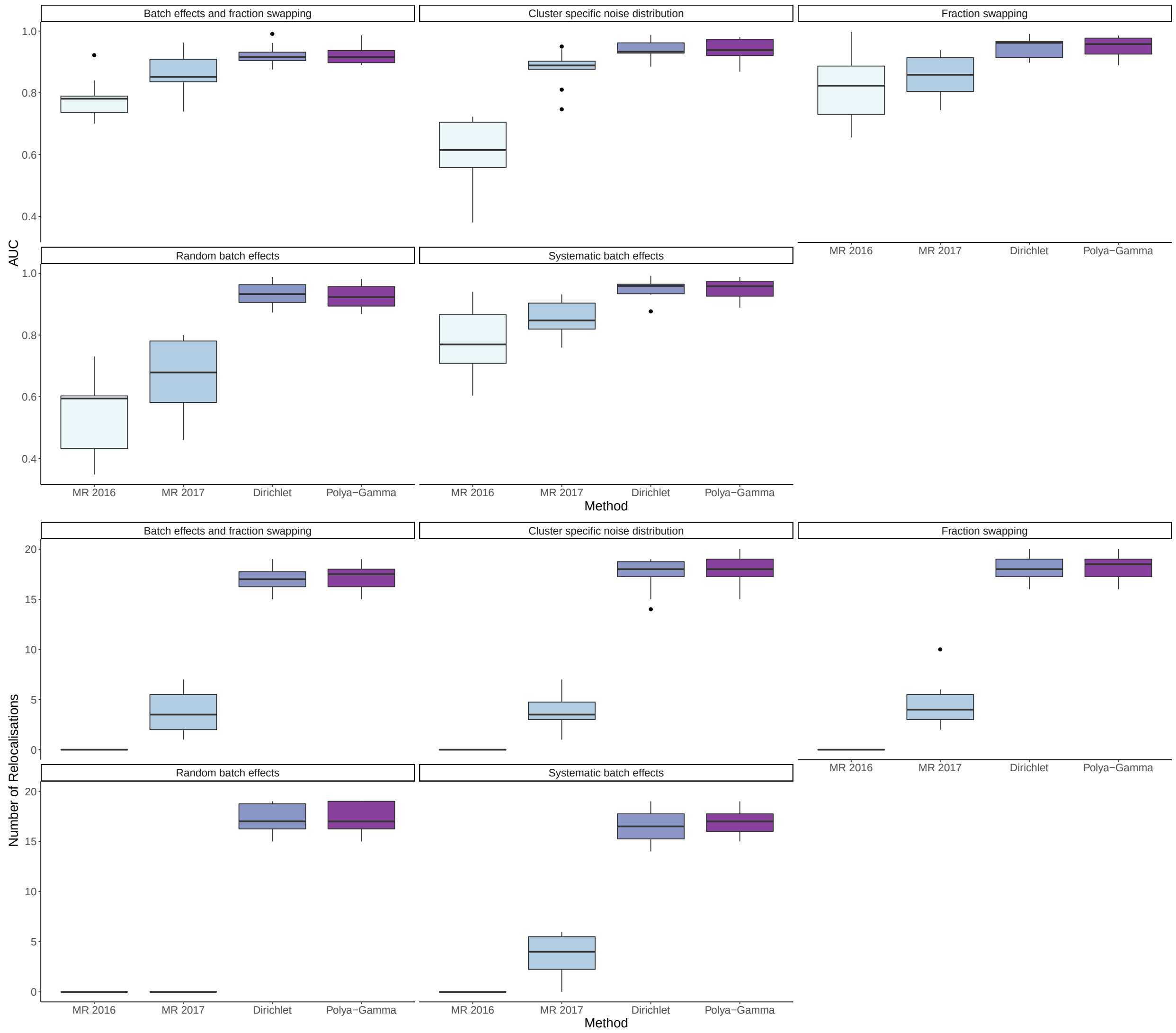

Figure 2: The first 5 boxplots compare MR to BANDLE with two different prior settings, using the area under curve (AUC). Distributions are over new simulated datasets. The second set of boxplots demonstrate how these AUCs translate into confident differential localisation events.

#### 1.3 Appendix 3: Posterior predictive distribution of subcellular niche

Here, we plot example posterior predictive distributions for the underlying abundance distributions of different organelles and subcellular niches from the BANDLE model. We observe close fits in all cases.

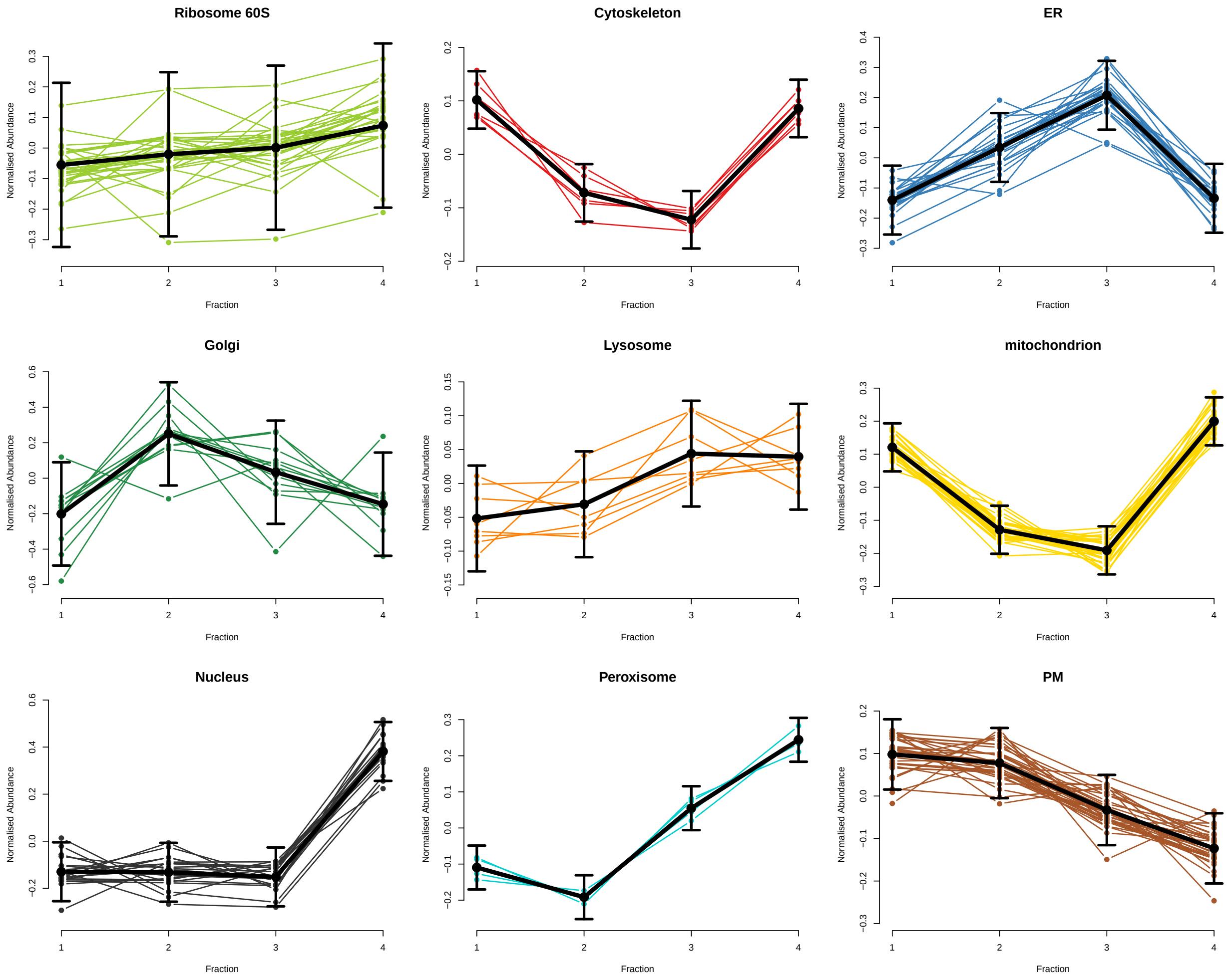

Figure 3: Posterior predictive fits for 9 organelles from the simulated example in the main text. Distributions fit the underlying data well.

##### 1.4 Appendix 4: MR-method p-value histogram

P-value histogram from the distance statistic underlying the MR method. P-values are clearly not uniformly distributed. Uniform distribution marked in purple

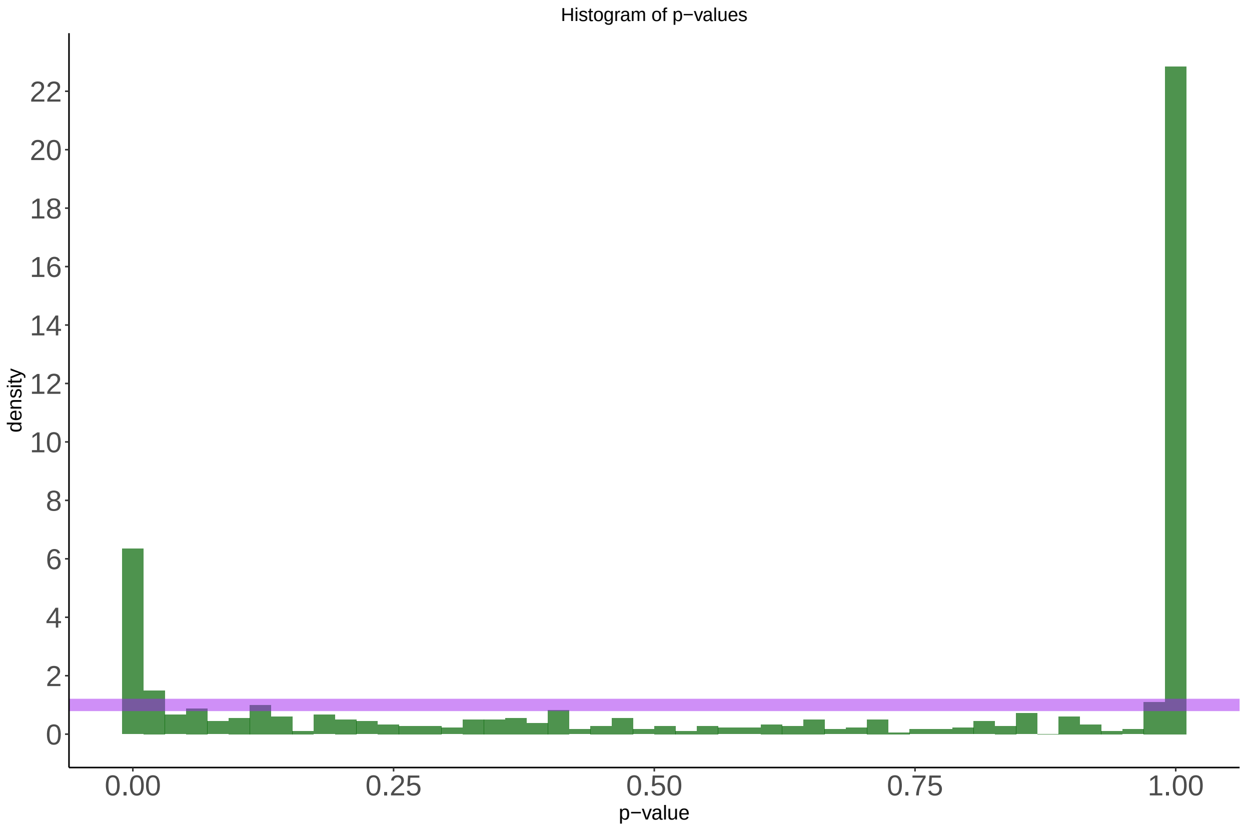

Figure 4: P-value histogram for the MR method

##### 1.5 Appendix 5: Example changes in localisation dynamics from BANDLE

BANDLE samples from the posterior distribution of subcellular localisation in each experimental condition. Hence, we can visualise this posterior distribution in each condition as a violin plot across the possible subcellular localisations. Here, we plot example changes in steady-state localisation from our simulated example in the main text. We capture direct changes in localisation, changes where the localisation was uncertain in one condition and changes where the localisation was uncertain in both conditions.

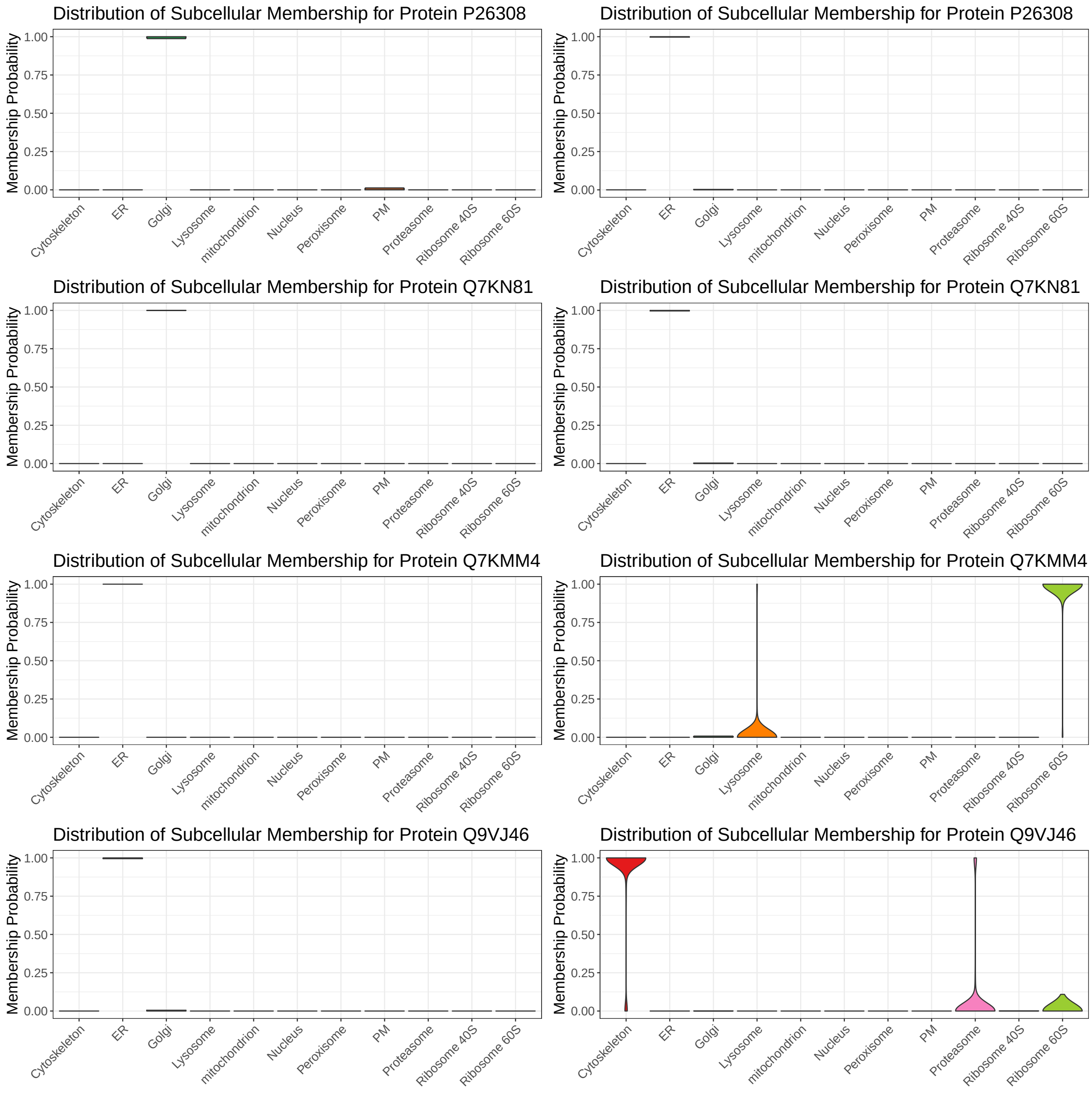

Figure 5: Posterior distribution of localisation probabilities for simulated control (left) and treatment (right)

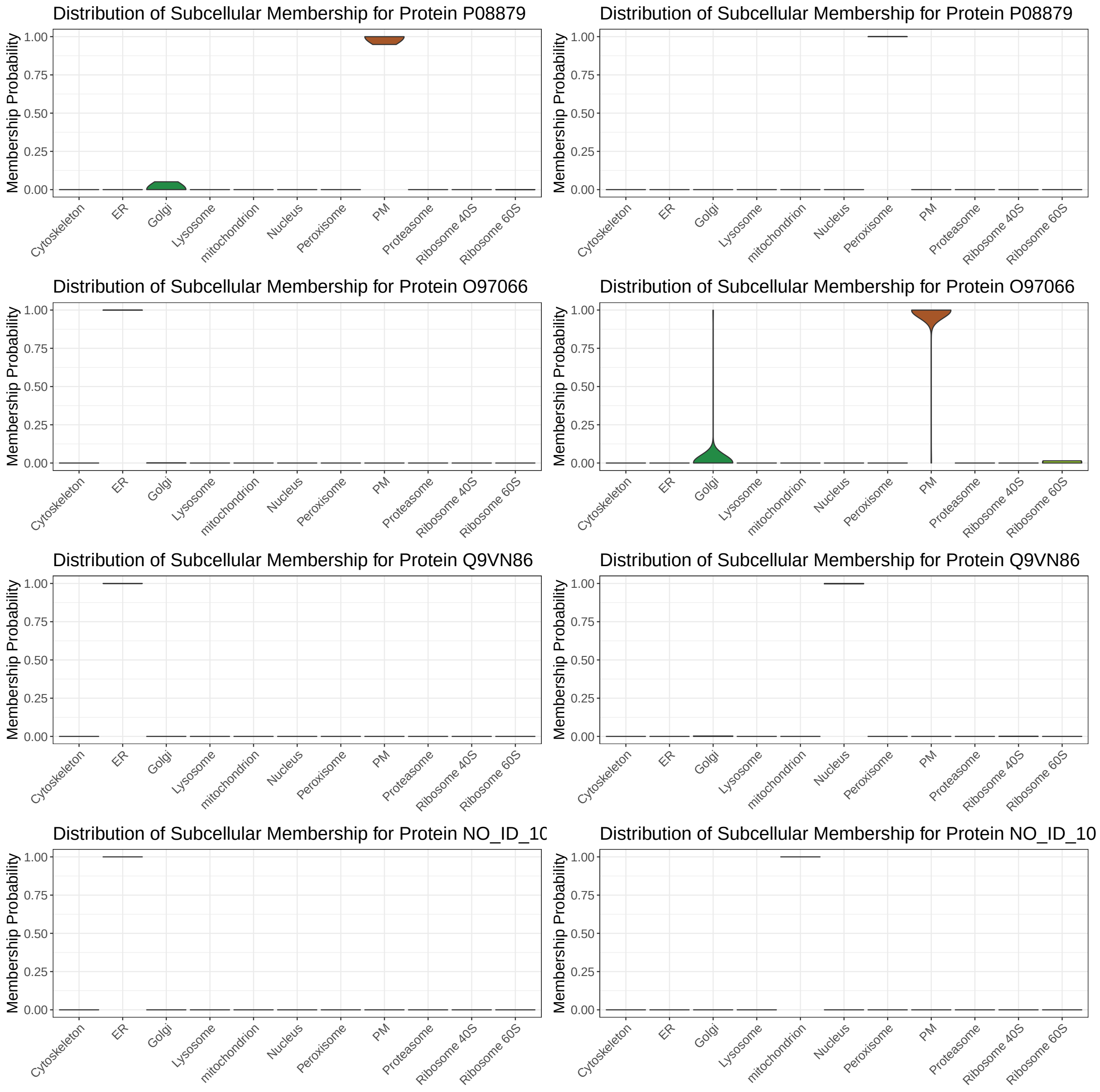

Figure 6: Posterior distribution of localisation probabilities for simulated control (left) and treatment (right)

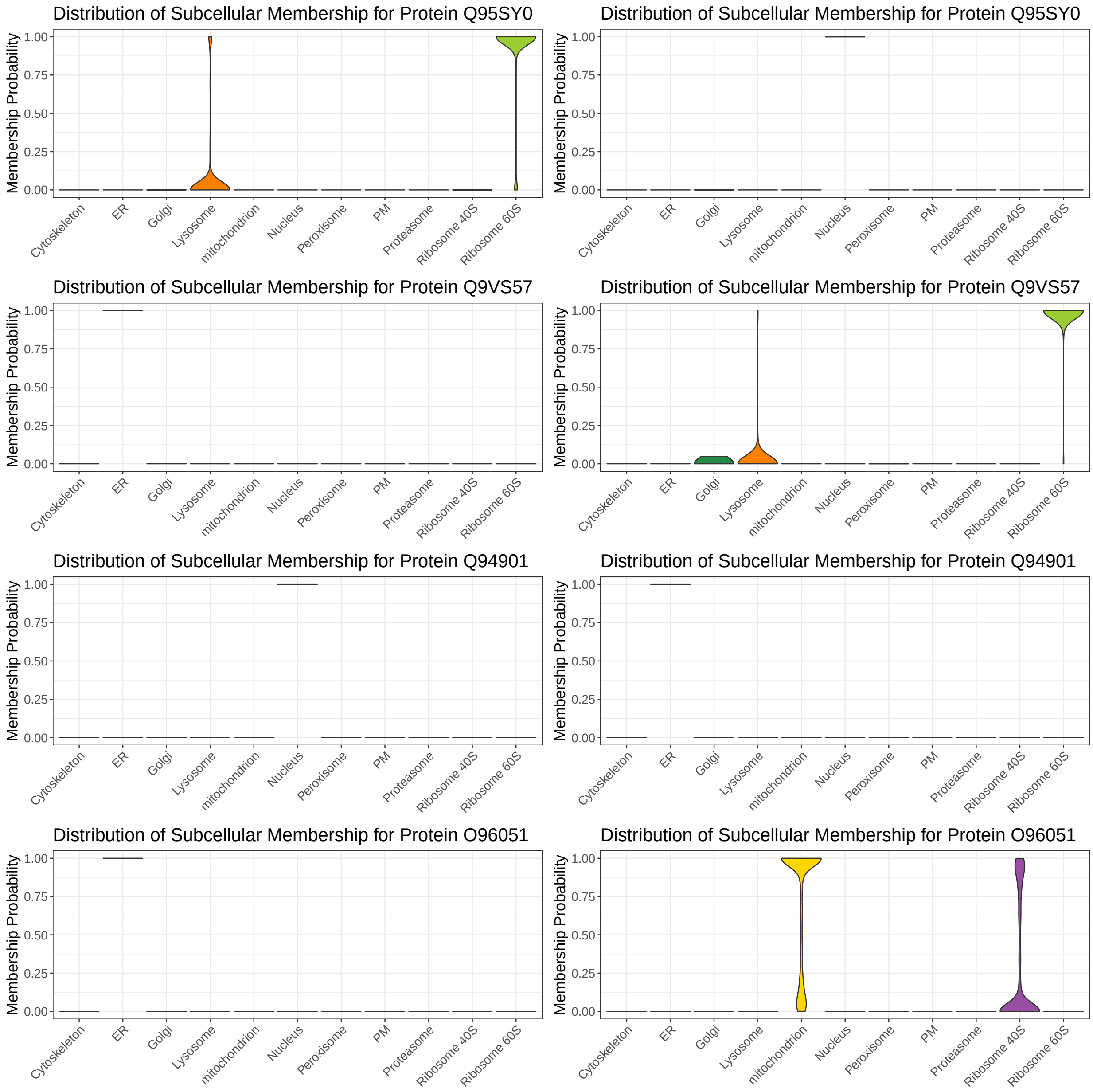

Figure 7: Posterior distribution of localisation probabilities for simulated control (left) and treatment (right)

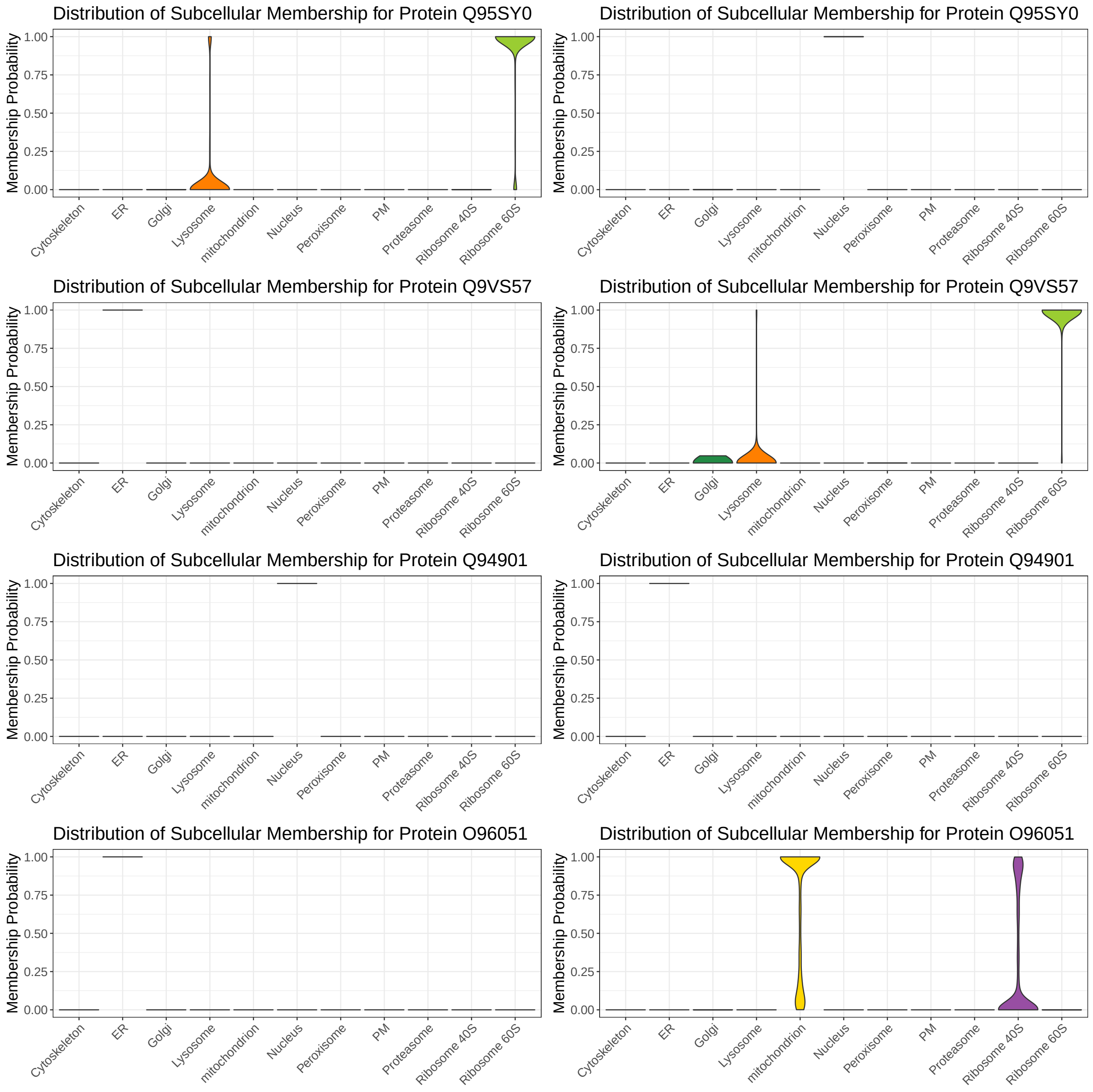

Figure 8: Posterior distribution of localisation probabilities for simulated control (left) and treatment (right)

#### 1.6 Appendix 6: Convergence Analysis EGF stimulation

We ran our MCMC sampler for 20,000 iterations, where we discarded 10,000 iterations for burn-in and retained every $10^{th}$ iteration for thinning to reduce autocorrelation. 8 chains were run in parallel and two were discarded for lack convergence by visual inspection. Example trace plots are plotted below. We further assessed convergence by computing $\hat{R}$ for parallel chains of the mixing weights and confirmed that they were less than 1.01 indicating that are chains are well-mixed. Finally, we concatenated the 6 remaining chains and computed the rank of each sample. These ranks are the plotted in separate histograms for each chain separately. Departures from uniformity of these histograms indicates non-convergence and we observe well-behaved rank plots.

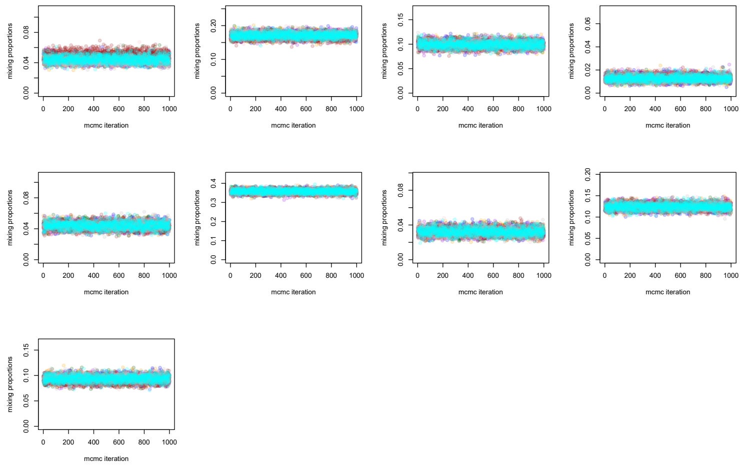

Figure 9: MCMC traceplot for EGF data. Colours correspond to independently run parallel MCMC chains.

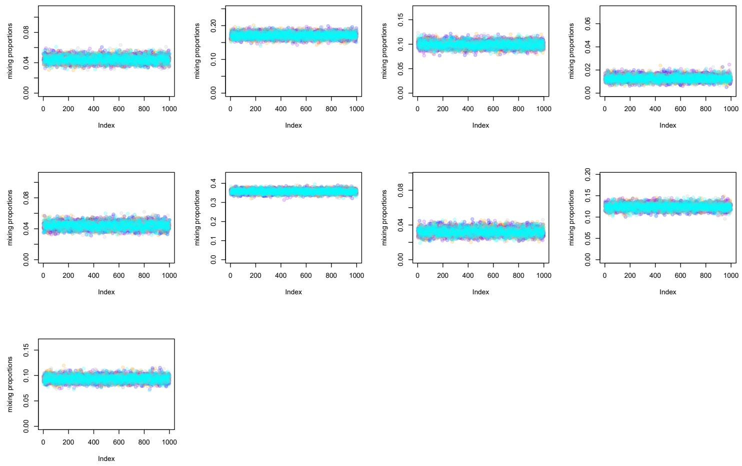

Figure 10: MCMC traceplot for EGF data, Colours correspond to independently run parallel MCMC chains.

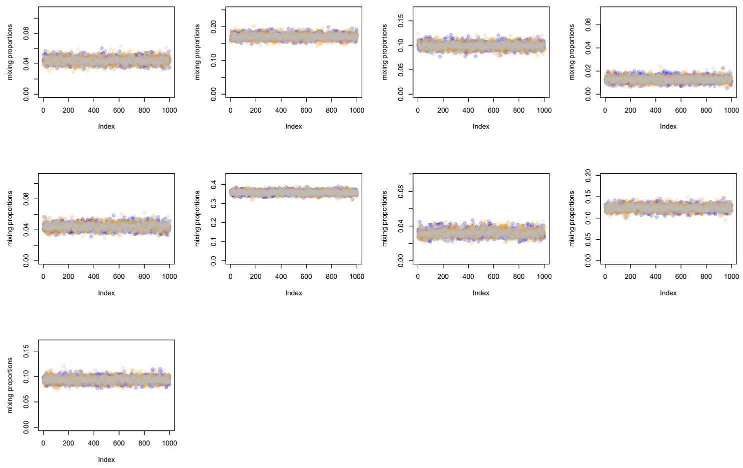

Figure 11: MCMC traceplot for EGF data. Colours correspond to independently run parallel MCMC chains.

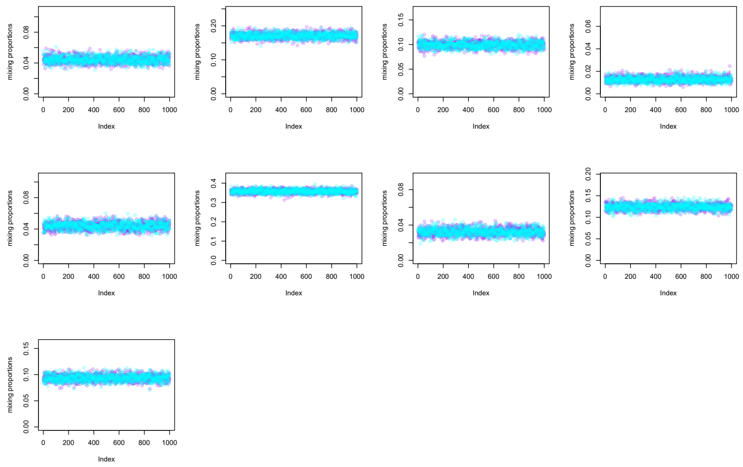

Figure 12: MCMC traceplot for EGF data. Colours correspond to independently run parallel MCMC chains.

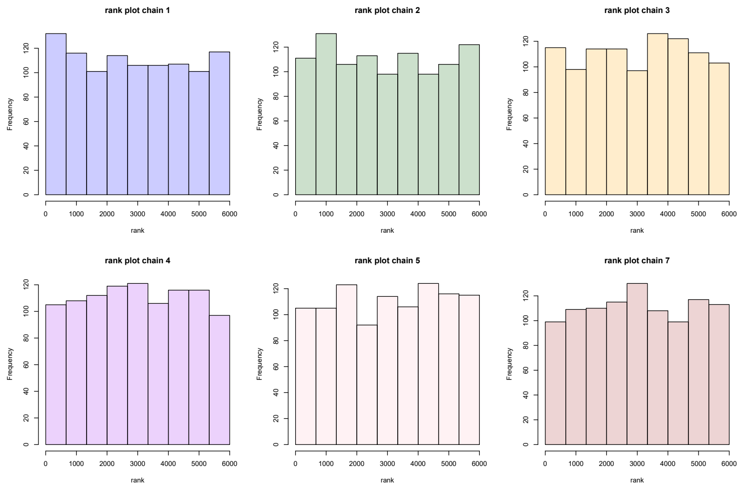

Figure 13: MCMC rank plot for EGF data. Colours correspond to independently run parallel MCMC chains.

#### 1.7 Appendix 7: EGF stimulation Phosphoproteomics time course

Example abundance changes for the phosphoproteomic time course experiment.

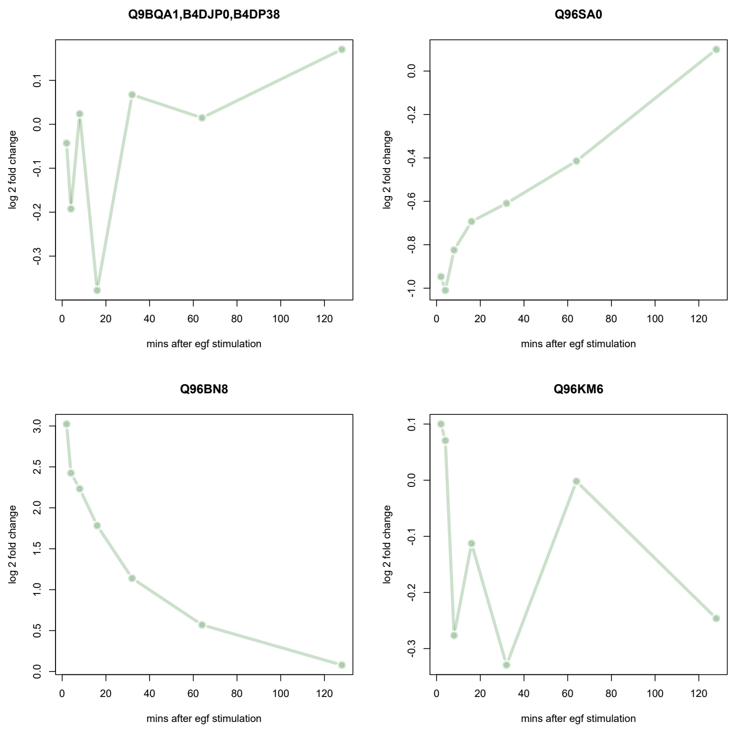

Figure 14: Example trajectories from the timecourse phosphoproteomics experiment

#### 1.8 Appendix 8: Convergence Analysis AP-4 Knockout

We ran our MCMC sampler for $20,000$ iterations, where we discarded $10,000$ iterations for burn-in and retained every $50^{th}$ iteration for thinning to reduce autocorrelation. 6 chains were run in parallel and one was discarded for lack convergence by visual inspection. Example trace plots are plotted below. We further assessed convergence by computing $\hat{R}$ for parallel chains of the mixing weights and confirmed that they were less than 1.01 indicating that are chains are well-mixed. Finally, we concatenated the 5 remaining chains and computed the rank of each sample. These ranks are the plotted in separate histograms for each chain separately. Departures from uniformity of these histograms indicates non-convergence and we observe well-behaved rank plots.

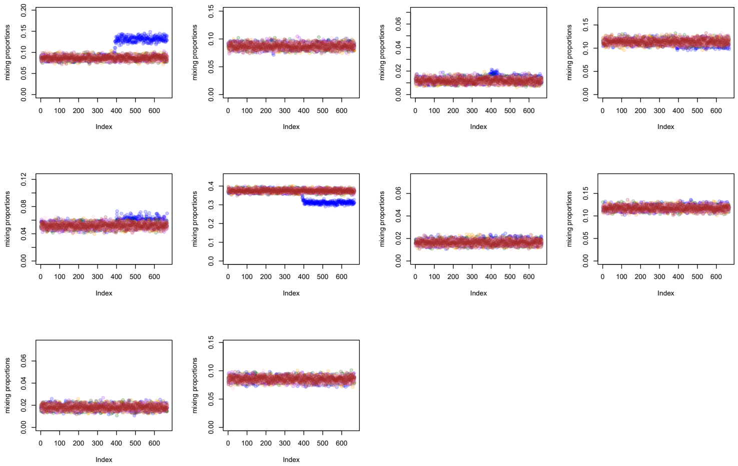

Figure 15: MCMC trace plot for AP-4 dataset. Colours correspond to independently run parallel MCMC chains.

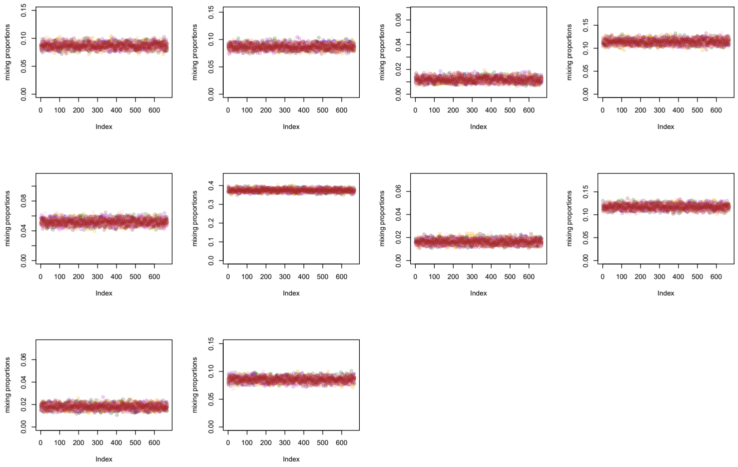

Figure 16: MCMC traceplot for AP-4 dataset. Colours correspond to independently run parallel MCMC chains.

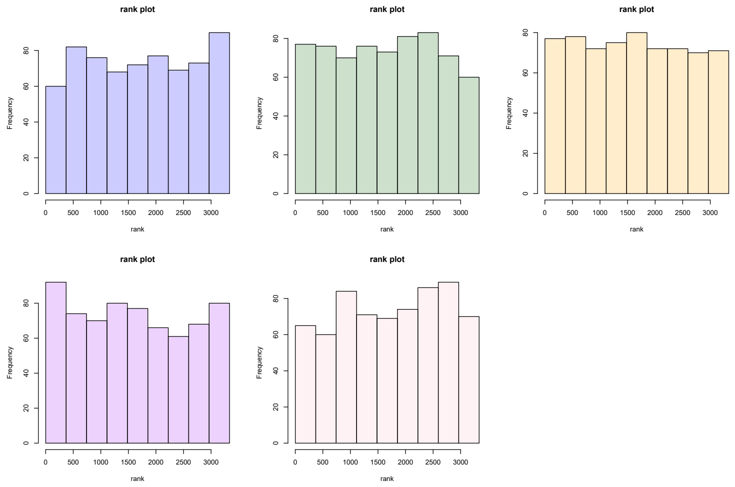

Figure 17: MCMC rank plot for AP-4 dataset

#### 1.9 Appendix 9: Convergence Analysis for HCMV datasets

We ran our MCMC sampler for 20, 000 iterations, where we discarded 10, 000 iterations for burn-in and retained every $50^{th}$ iteration for thinning to reduce autocorrelation. 6 chains were run in parallel and convergence was analysed by visual inspection, and one chains was discarded. Example trace plots are plotted below. We further assessed convergence by computing $\hat{R}$ for parallel chains of the mixing weights and confirmed that they were less than 1.01 indicating that are chains are well-mixed. Finally, we concatenated the 5 remaining chains and computed the rank of each sample. These ranks are the plotted in separate histograms for each chain separately. Departures from uniformity of these histograms indicates non-convergence and we observe well-behaved rank plots.

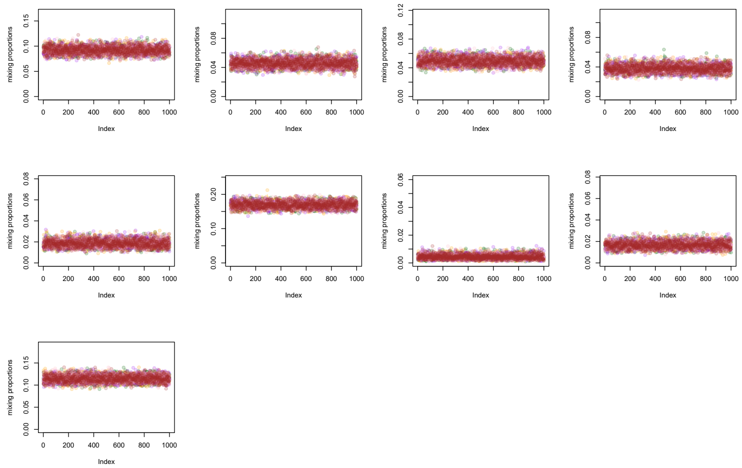

Figure 18: MCMC trace plot for HCMV dataset 24 hpi

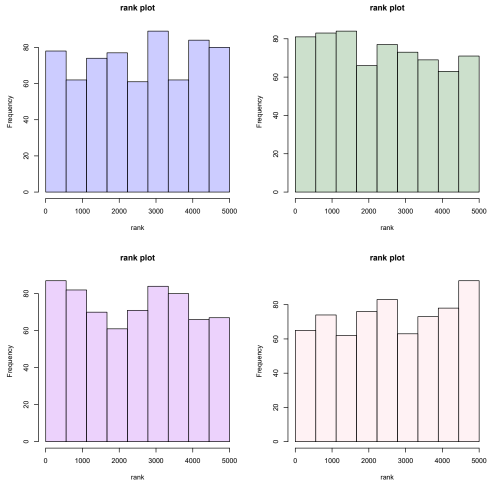

Figure 19: MCMC rank plot for HCMV dataset 24 hpi

##### 1.10 Appendix 10: Prior settings and sensitivity analysis

The hyperparameter $\alpha$ is the prior on the mixing weights $\boldsymbol{\pi}$, where $\pi_{ij}$ is the prior probability that a protein belongs to the $i^{th}$ niche in the control dataset and niche $j$ in the treatment dataset. The entries of $\alpha$ can be interpreted as the prior relative proportions of protein allocations. Let $J$ be the matrix of all ones, it is typical in Bayesian mixture modelling to set $\alpha = 0.5J$ or $\alpha = J$, corresponding to the Jeffreys' prior and the symmetric prior, respectively. However, in our scenario the diagonal and off-diagonal terms have different meanings. The diagonal terms correspond to proteins allocated to the same niche in both datasets and the off-diagonal terms correspond to differential localised proteins. However, there are far more off diagonal terms than diagonal terms. Hence, the Jeffreys' and symmetric priors implicitly assume that the there are more differentially localised proteins that spatial stable. Of course, this is at odds with our expectations and thus we opt for a more sensible weakly informative prior as a default. We set $\alpha_{jj} = 1$ and $\alpha_{jk} = 0.01$ for $k \neq j$. This assume that there are roughly an order of magnitude fewer differentially localised proteins that spatially stable ones. This default is used in all simulations and application except the EGF simulation dataset. In that case, we have prior knowledge of a differentially localisation between the Plasma membrane and the Endosome and so we set the corresponding entry of $\alpha$ to 1.

In general, we do not find that our analysis is very sensitive to the prior choice. To demonstrate, we perform a sensitivity analysis for the results using the Jeffreys' prior, the symmetric prior and our weakly informative prior. In the context of the simulation examples in the main text, we apply the different prior choice and examine the results. Since the primary quantity of interest is the prediction of differentially localised proteins, we examine this quantity. The following ROC curve demonstrate that the results are almost identical across the different prior choices.

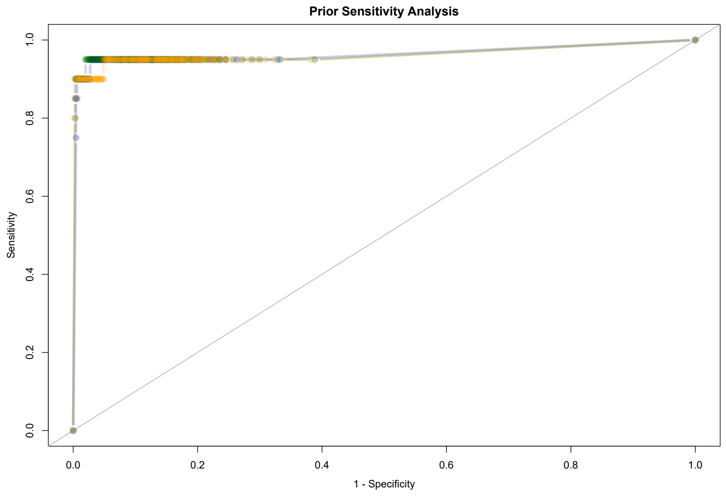

Figure 20: ROC curve for examining prior sensitivity. Blue corresponds to default, dark green to the Jeffreys' prior and orange to the symmetric prior. The curves are essentially indistinguishable.

Prior information is carefully encoded using domain knowledge and previous analysis. In brief, for the EGF application, we encode that the most likely transition is from plasma membrane to Lysosome. To evaluate the coherence of our prior we perform a *prior predictive check* (Gelman *et al.*, 1995). The summary statistic of interest is the number of differential localisation and the expected number of differential localisation, given our prior, is roughly 4.6. Furthermore, the prior probability that there are more than 15 differential localisation is less than 0.01.

For the AP-4 Application, we first performed a prior predictive check and find that our prior configuration leads to a 5.3 proteins *a priori* differential localisation in expectation and the probability that more than 15 proteins are differential localised is $\approx 0.03$.

For the HCMV application, priors are set such that the expected prior number of differential localisation is roughly 3.

We interpret these values, in all situations, in that the prior will induce shrinkage towards a small number of differentially localised proteins, avoiding spurious results that are not strongly supported by the data.

##### 1.11 Appendix 11: Selecting $\tau$

One hyperparameter that we haven not yet discussed in the choice of $\tau$, when using the empirical strategy to select the prior for the Pólya-Gamma based prior (see supplementary methods). One possible way to select $\tau$ is to first perform a prior predictive check. However, this can be arduous if a reasonable value is not known in advance. We suggest one strategy for generating appropriate values of $\tau$. The first is to select a weakly informative Dirichlet prior, for example, the default we have suggested in the previous section. We then compute the standardised Kullback-Leibler (KL) divergence between this weakly informative Dirichlet prior and the a range of possible Pólya-Gamma based priors. If we believe that our Dirichlet prior is sensible then a sensible Pólya-Gamma prior will have low KL divergence. In figure 21, we vary the value of $\tau$ (on the log scale) for different values of the mean for the Pólya-Gamma prior. There is a clear elbow in this plot. We do not advise purely selecting the value of $\tau$ which minimises this KL divergence, rather choose $\tau$ roughly in the that region and perform a prior predictive check to ensure that it leads to sensible prior inferences. A default value of $\tau = 0.3$ appears to work well, in practice.

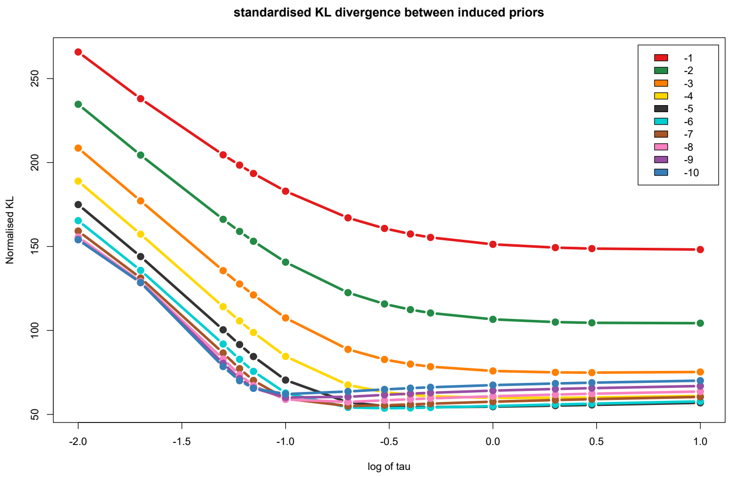

Figure 21: KL divergence plot show KL divergence between the Polya-Gamma prior and the weakly informative default Dirichlet prior for vary values of $\tau$. Each colour indicates a different choice of mean for the Polya-Gamma based prior.

#### 1.12 Appendix 12: EGF stimulation figures

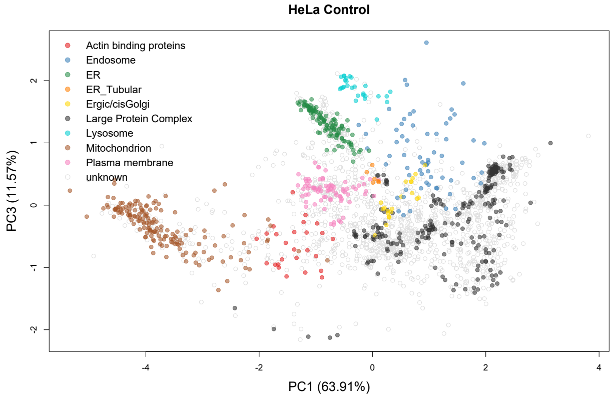

Figure 22: A PCA plot of the control HeLA dataset from (Itzhak *et al.*, 2016). Each pointer corresponds to a protein and marker proteins are highlighted according to their subcellular niche.

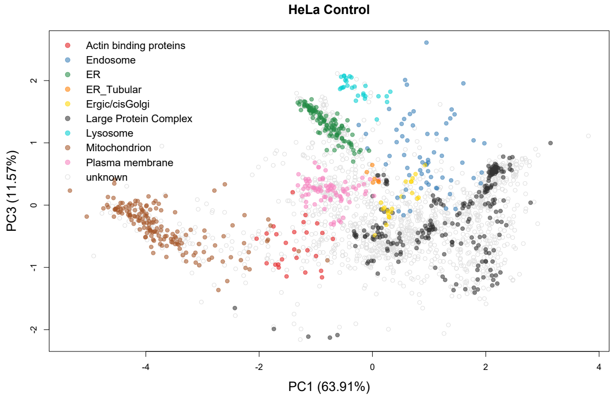

Figure 23: A PCA plot of the EGF stimulated HeLA dataset from (Itzhak *et al.*, 2016). Each pointer corresponds to a protein and marker proteins are highlighted according to their subcellular niche.

#### 1.13 Appendix 13: AP-4 knockout figures

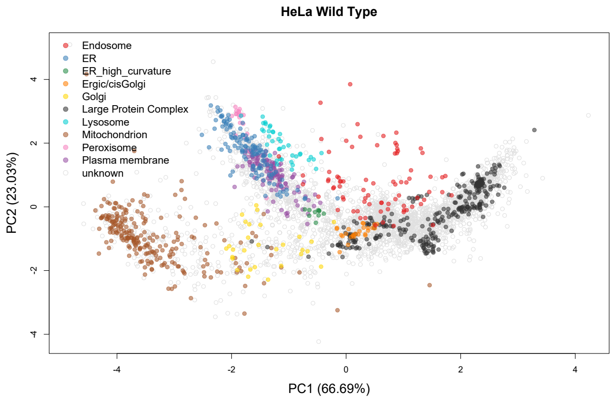

Figure 24: A PCA plot of the control HeLA dataset from (Davies *et al.*, 2018). Each pointer corresponds to a protein and marker proteins are highlighted according to their subcellular niche.

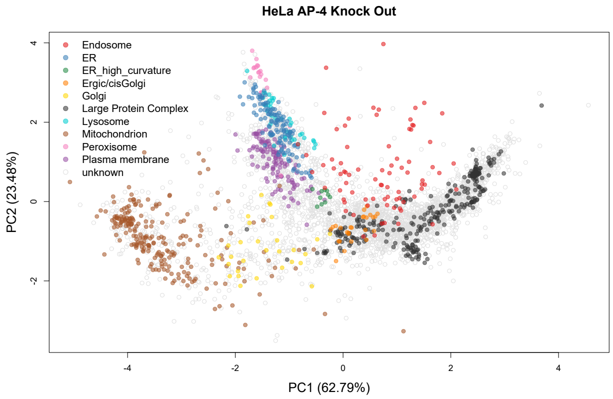

Figure 25: A PCA plot of the AP-4 knockout HeLA dataset from (Davies *et al.*, 2018). Each pointer corresponds to a protein and marker proteins are highlighted according to their subcellular niche.

#### 1.14 Appendix 14: HCMV PCA plots

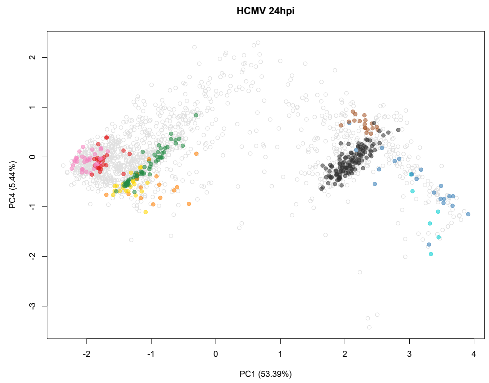

Figure 26: A PCA plot of control fibroblast cells dataset from (Beltran *et al.*, 2016). Each pointer corresponds to a protein and marker proteins are highlighted according to their subcellular niche.

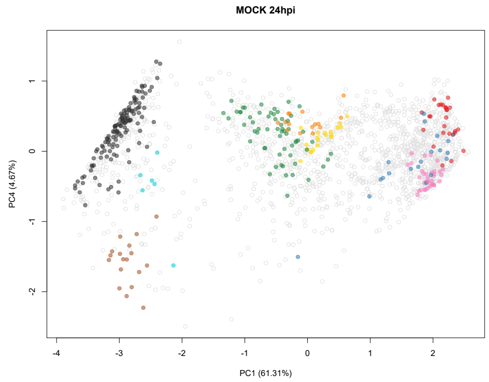

Figure 27: A PCA plot of HCMV infected fibroblast cells dataset from (Beltran *et al.*, 2016). Each pointer corresponds to a protein and marker proteins are highlighted according to their subcellular niche.

#### 1.15 Appendix 15: GO enrichment analysis HCMV dataset

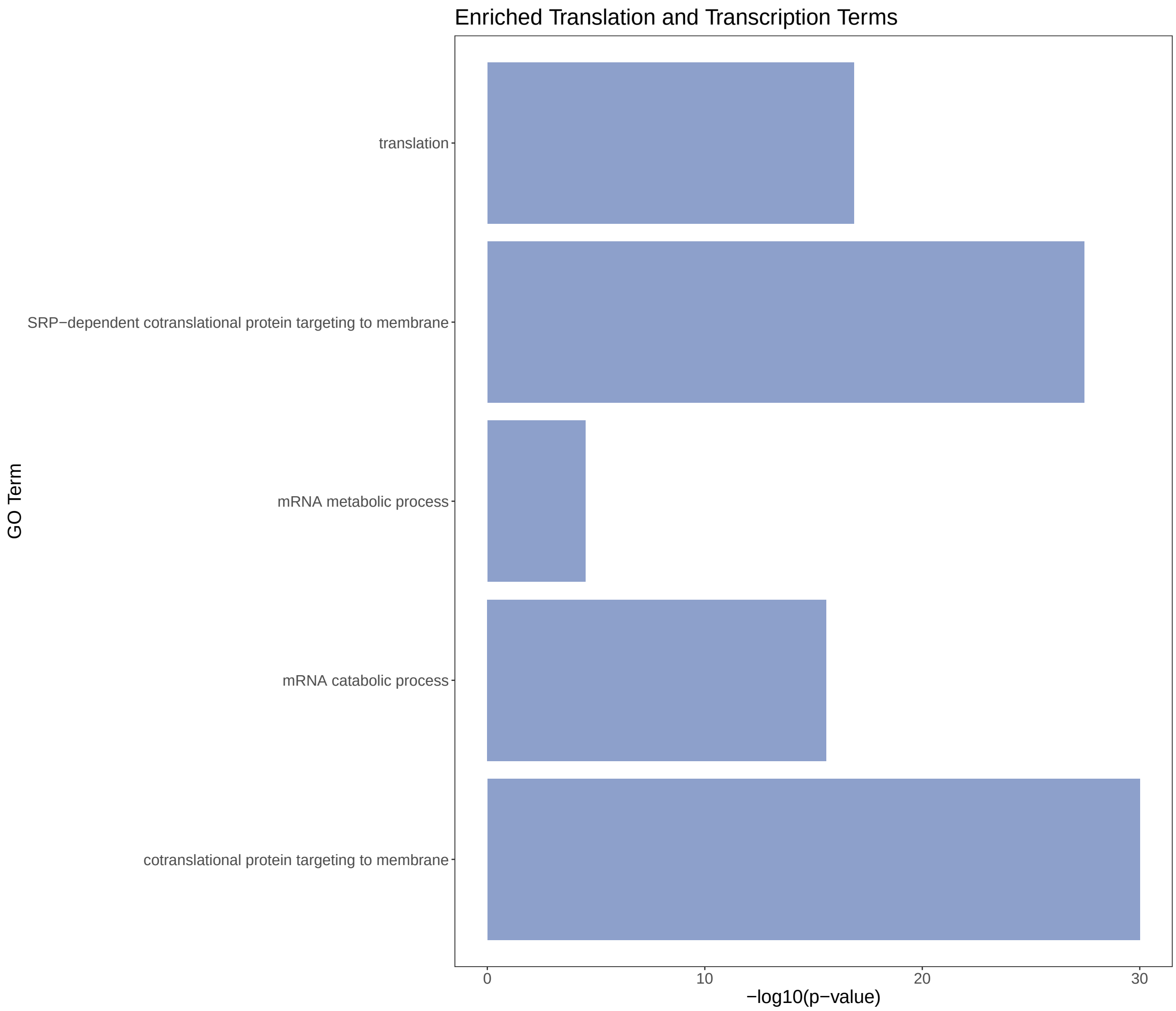

Figure 28: GO enrichment results (Translation and Transcription terms)

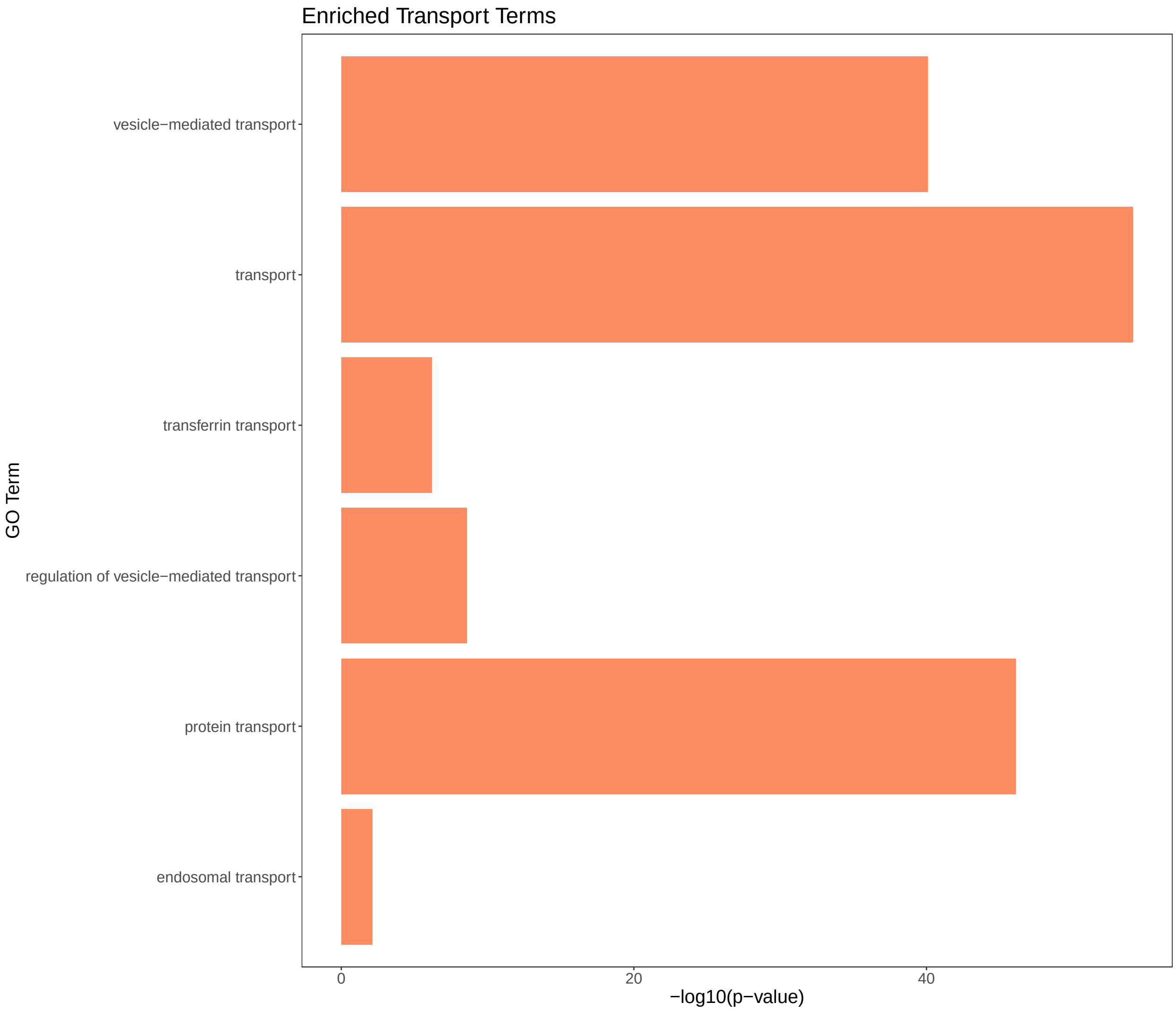

Figure 29: GO enrichment results (Transport terms)

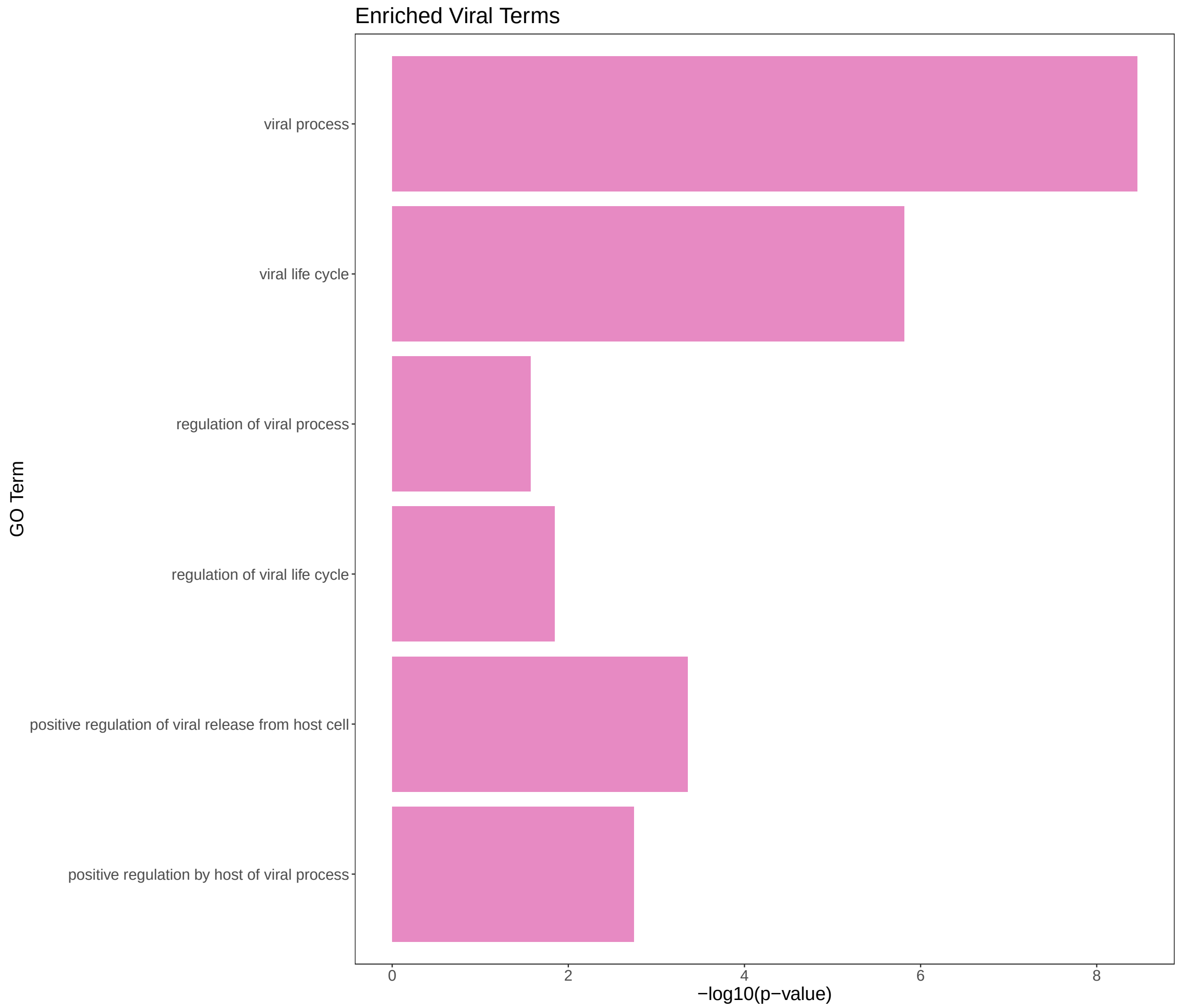

Figure 30: GO enrichment results (Viral processes)

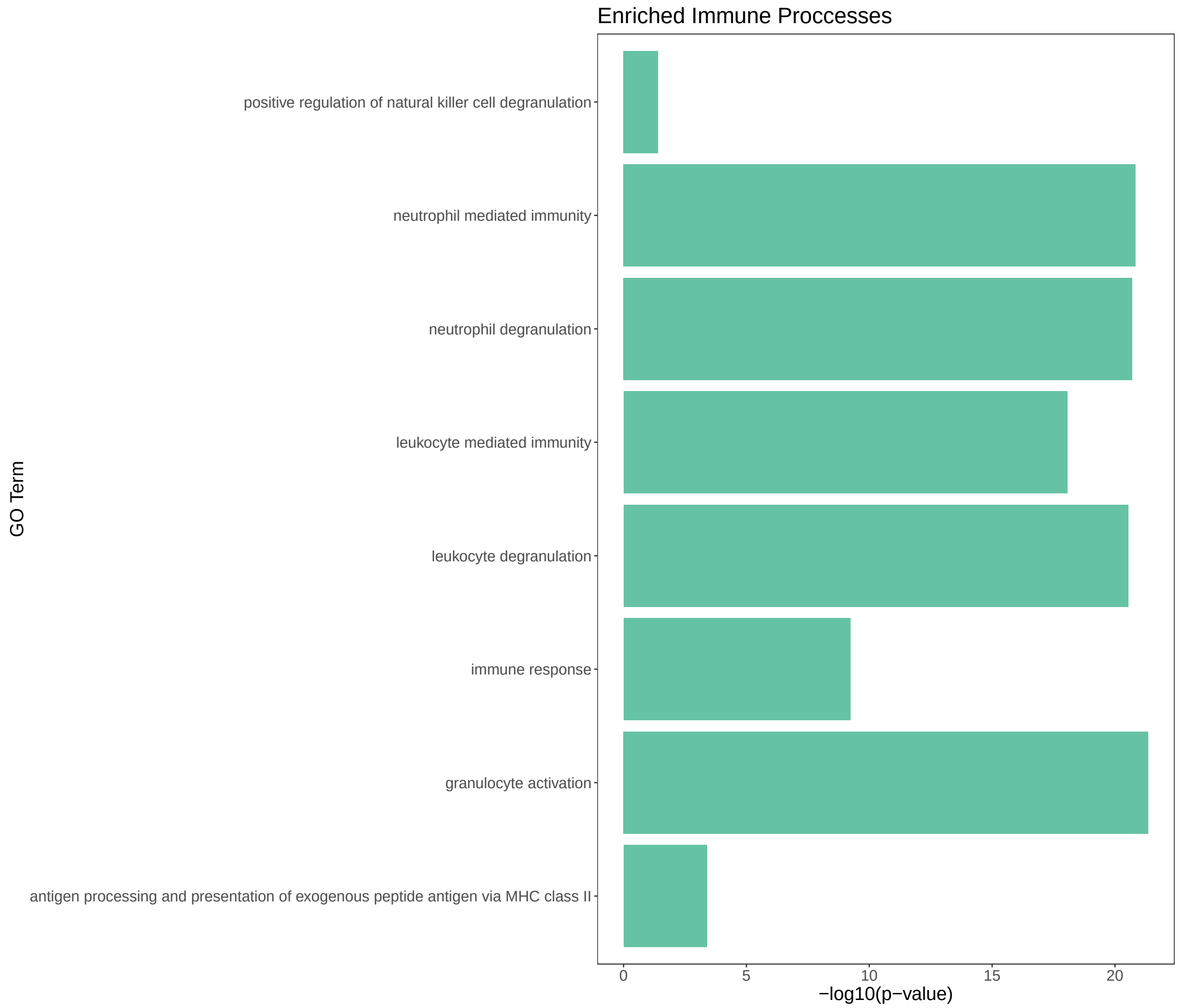

Figure 31: GO enrichment results (Immune processes)

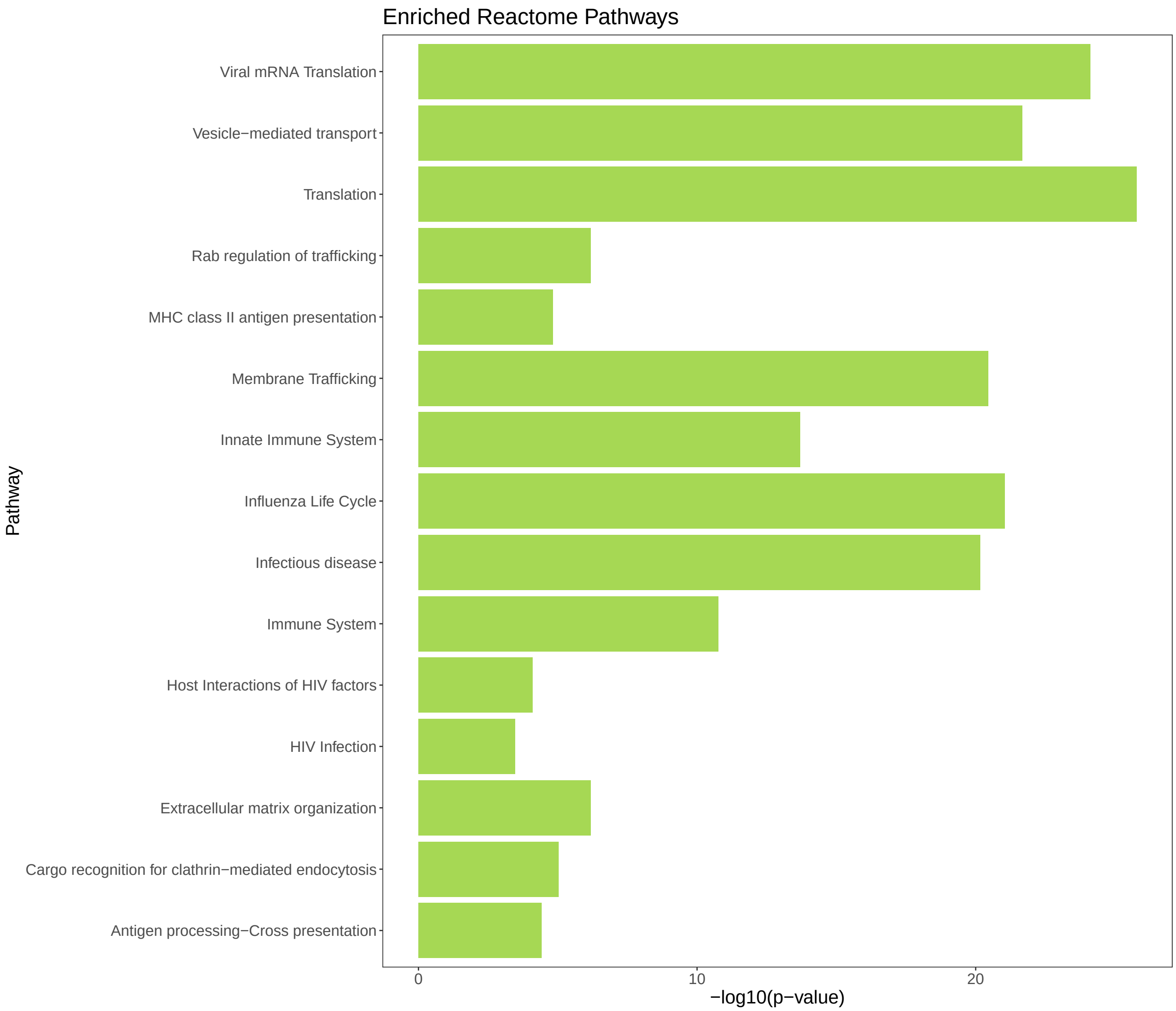

Figure 32: Reactome pathway enrichment results

#### 1.16 Appendix 16: HCMV additional figures abundance and degradation assays

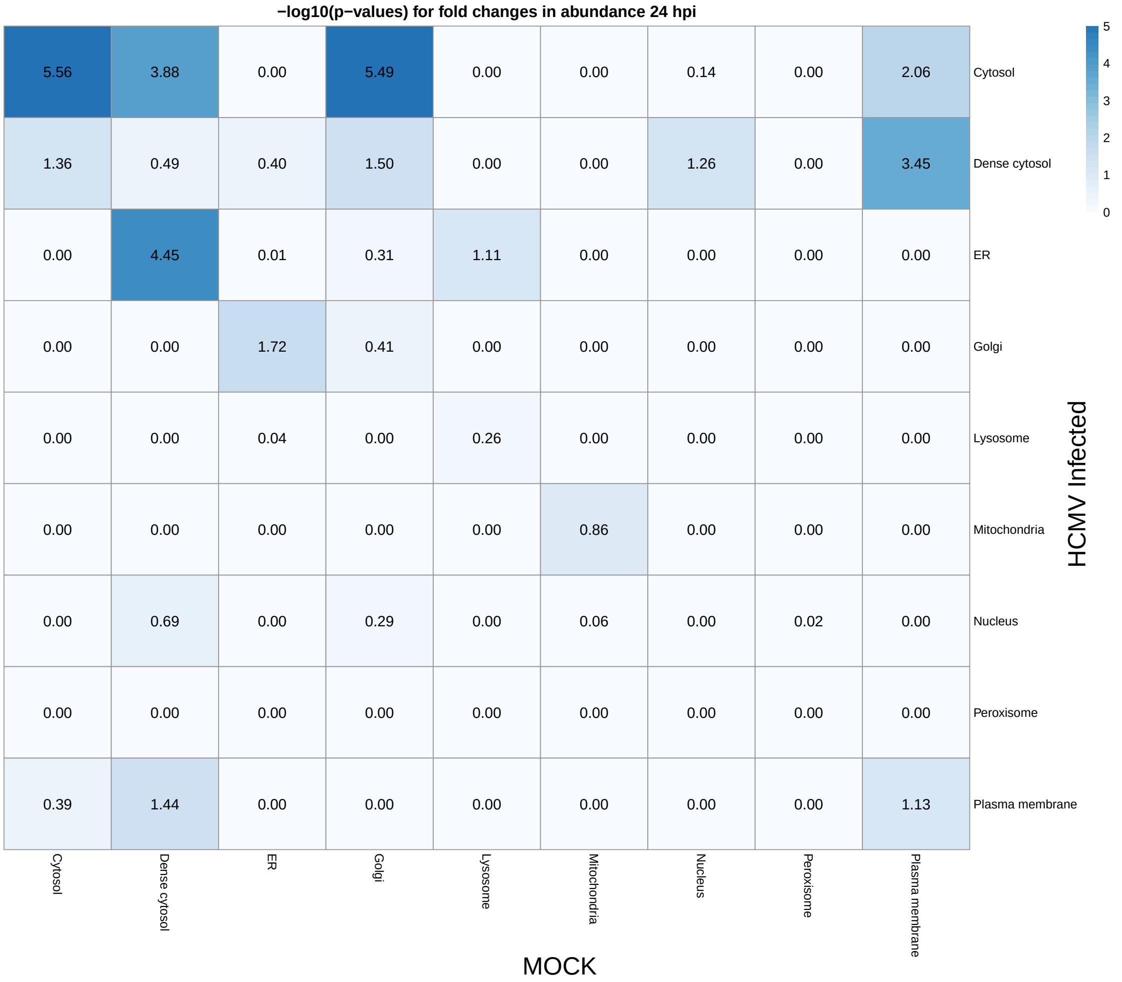

Figure 33: $-\log_{10}(p\text{-values})$ for the fold changes in abundance stratified by localisation dynamics.

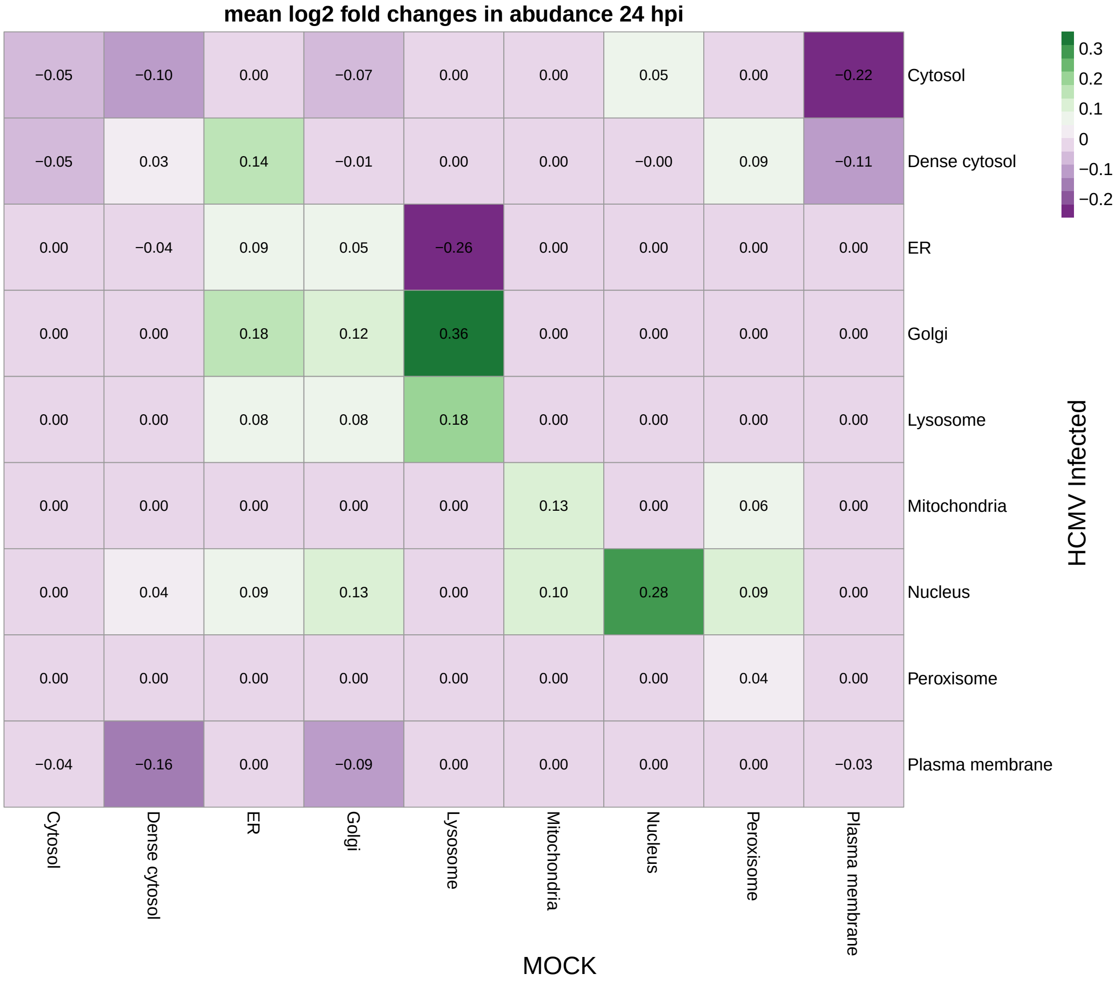

Figure 34: Fold changes in abundance stratified by localisation dynamics.

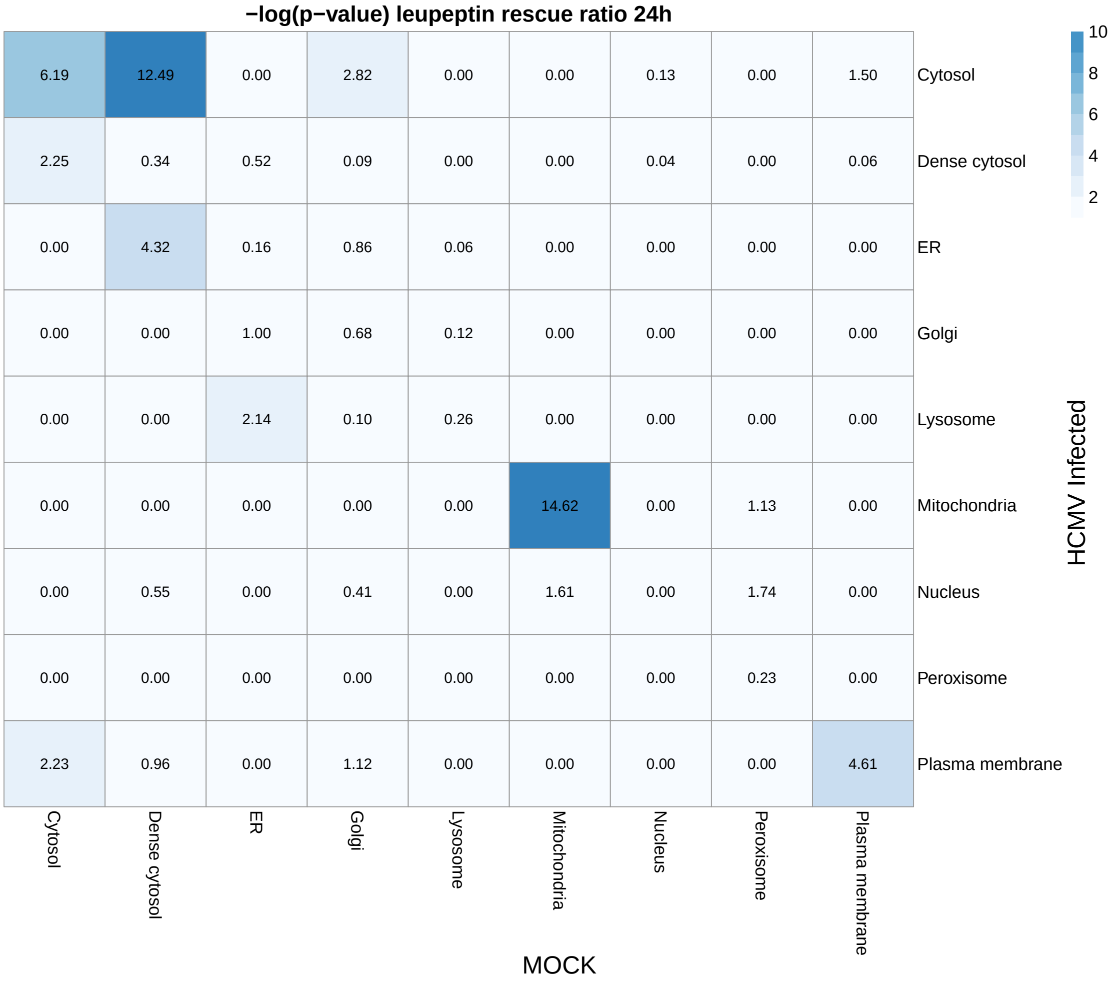

Figure 35: $-\log_{10}(p\text{-values})$ for rescue ratios for leupeptin degradation assays stratified by localisation dynamics. Values of the diagonal represent ratio for protein with no change in localisation.

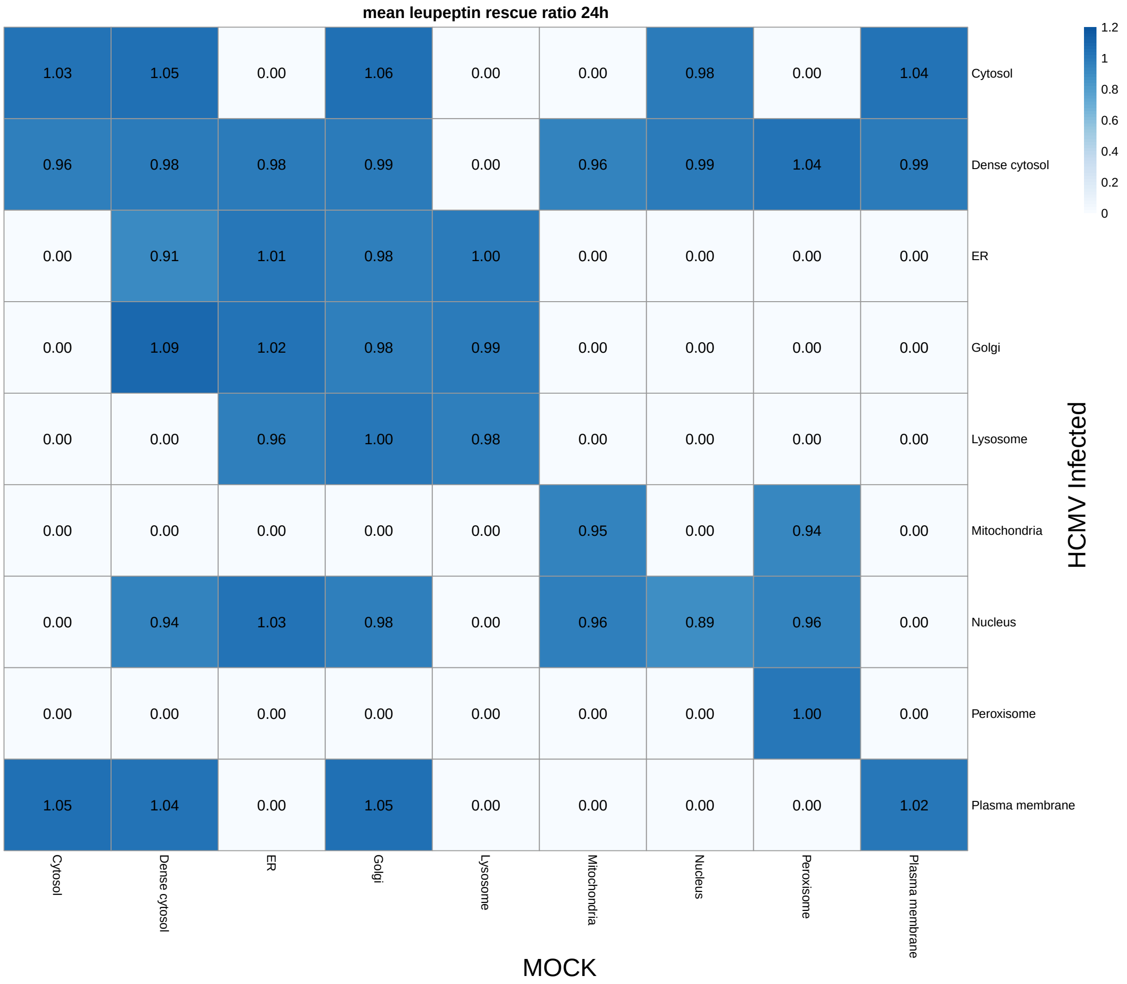

Figure 36: Rescue ratios for leupeptin degradation assays stratified by localisation dynamics

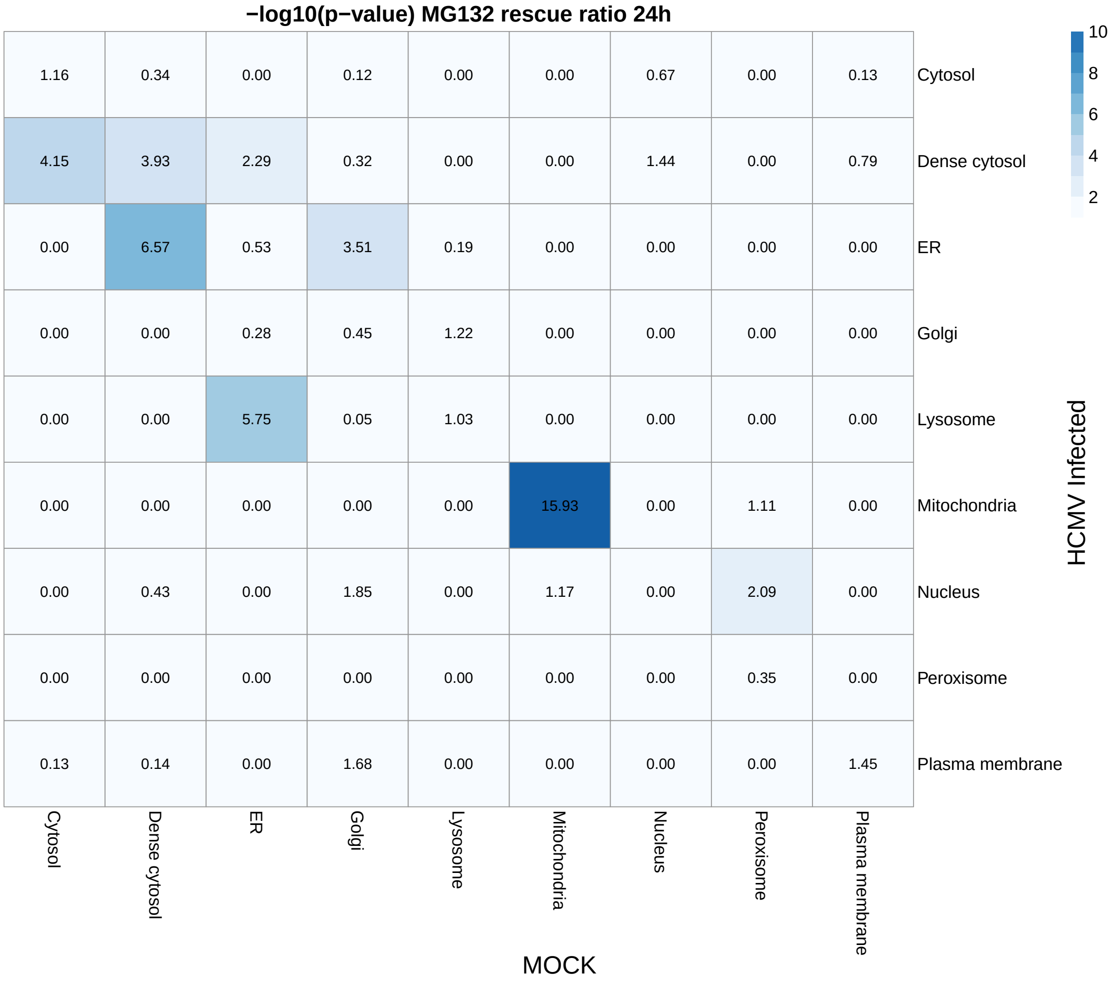

Figure 37: $-\log_{10}(p\text{-values})$ for rescue ratios for MG132 degradation assays stratified by localisation dynamics. Values of the diagonal represent ratio for protein with no change in localisation.

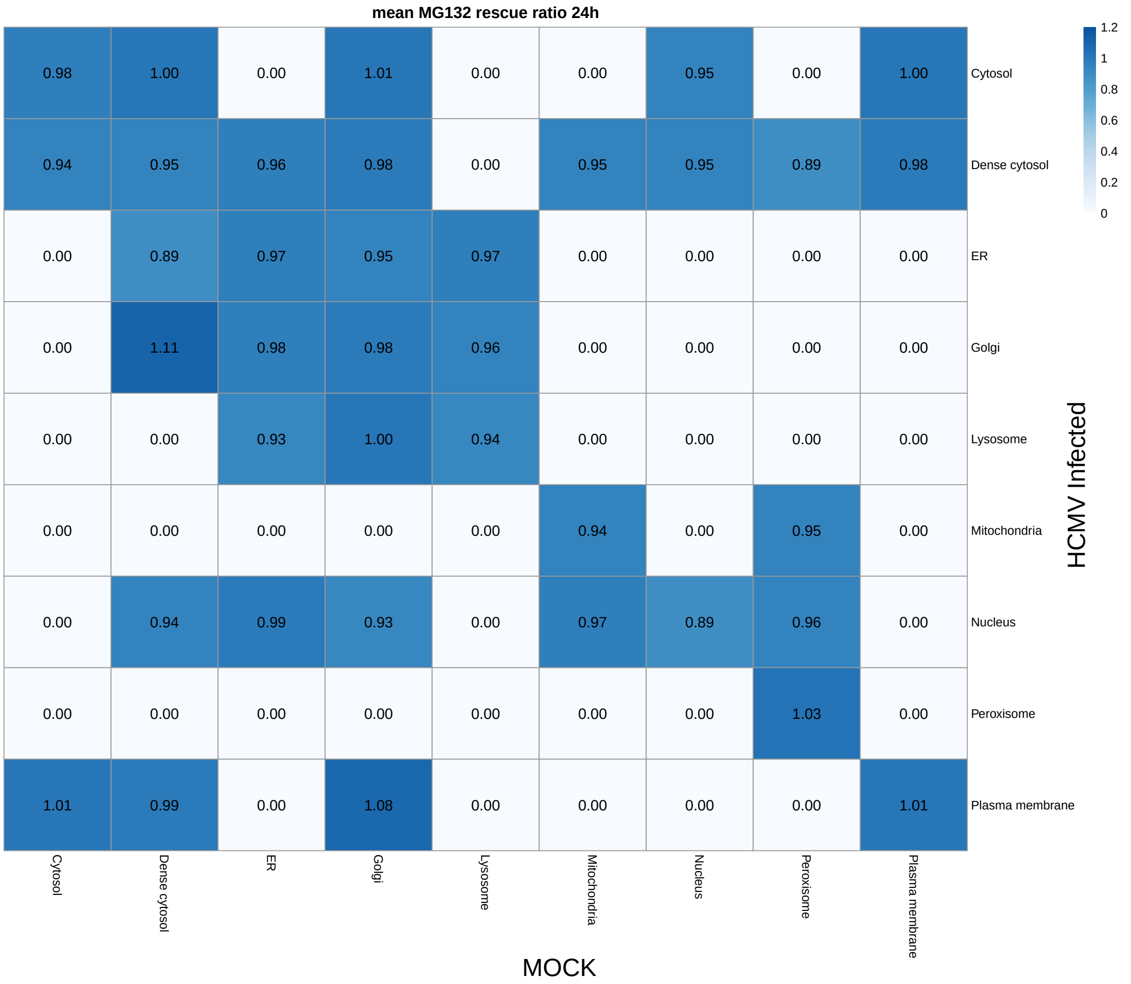

Figure 38: Rescue ratios for MG132 degradation assays stratified by localisation dynamics

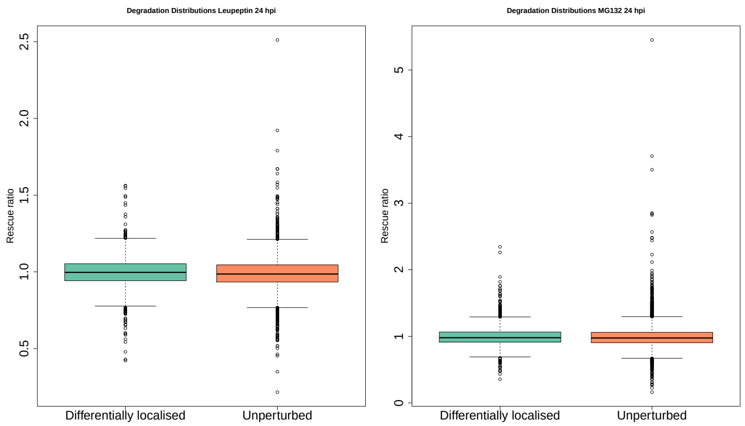

Figure 39: Boxplots of the global degradation distributions (Rescue Ratio) for MG132 and leupeptin. Separate distributions are plotted for differentially localised proteins are those are not. No difference is observed using a t-test with threshold 0.05.

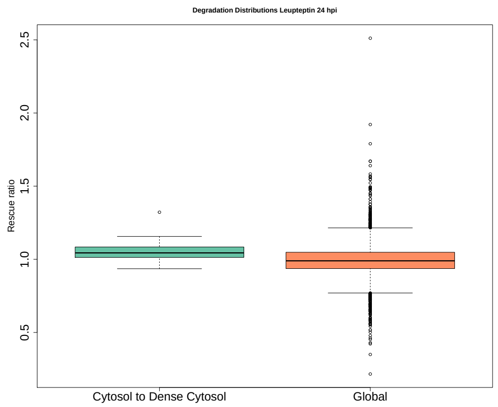

Figure 40: Leupeptin distributions of protein recruited from the cytosol to the dense cytosol, showing increased proteins targeted for degradation. A t-test was applied to test for differences $p < 0.05$.

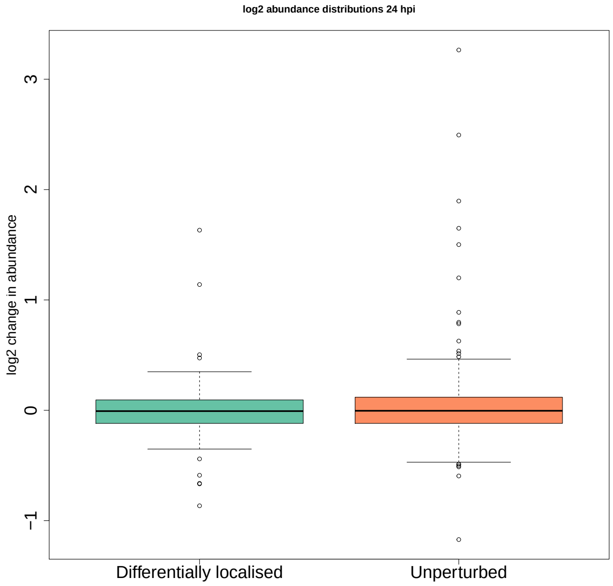

Figure 41: Global abundance distributions for proteins 24 hpi separated into differentially localised or not. There is no difference between differentially proteins and those that are not. No difference is observed using a t-test with threshold 0.05

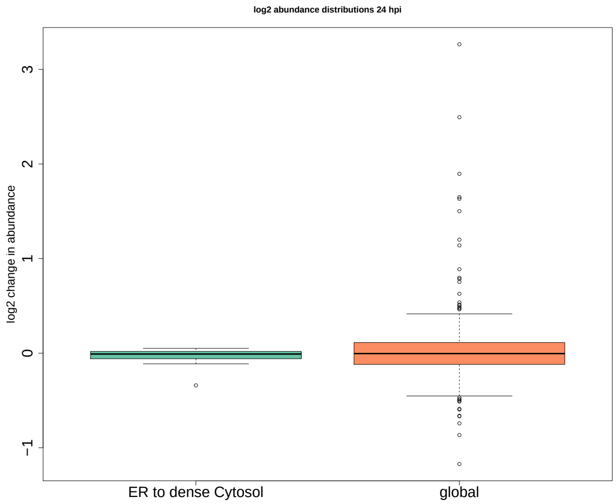

Figure 42: Boxplots for $\log_2$ normalised abundance distributions. Proteins recruited from the ER to dense cytosol show a decrease in abundance when compared to the global distribution. A t-test was applied to test for differences $p < 0.05$.

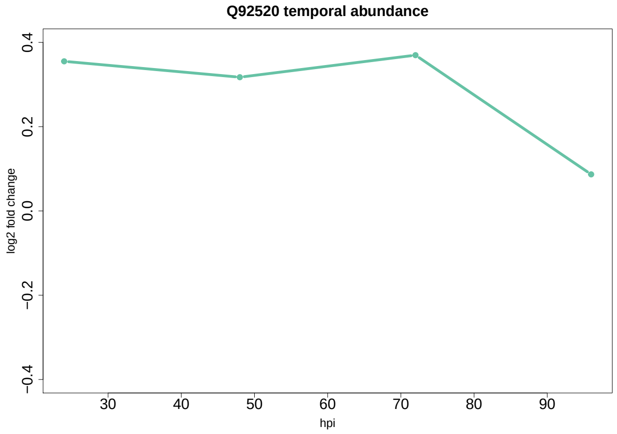

Figure 43: The temporal protein abundance of Q92520 at 24 hour intervals over a 4 day period. Clearly Q92520 is upregulated until 92 hpi.

#### 1.17 Appendix 17: HCMV additional figures acetylation data

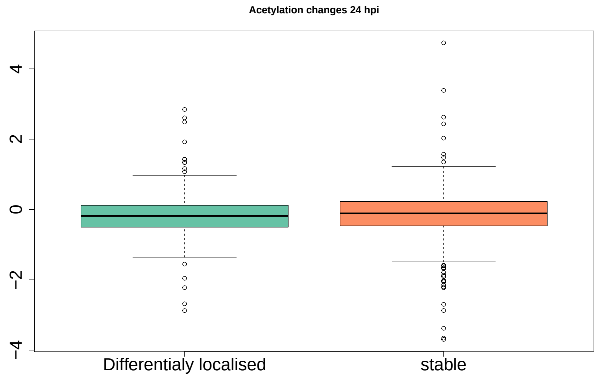

Figure 44: Global distributions for acetylation changes for HCMV 24 hpi compare to MOCK. We do not observe any correlations between differential localisation and acetylation changes; that is, no difference is observed using a t-test with threshold $p = 0.05$.

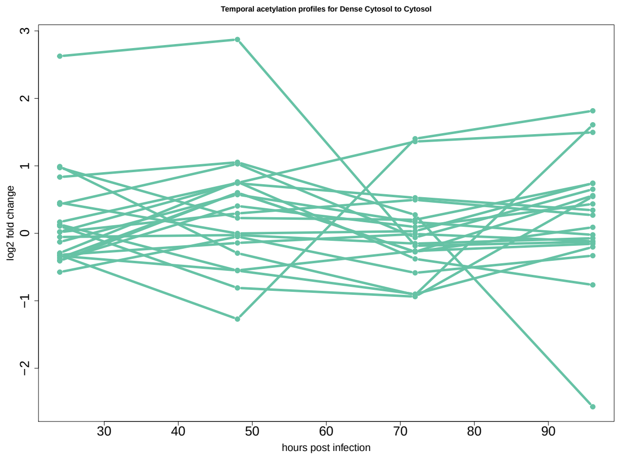

Figure 45: Temporal acetylation profiles for HCMV infected cells which relocalise from dense Cytosol to the Cytosol. Skp1 has a 2.5 fold increase in acetylation at 24 hpi. No test was performed.

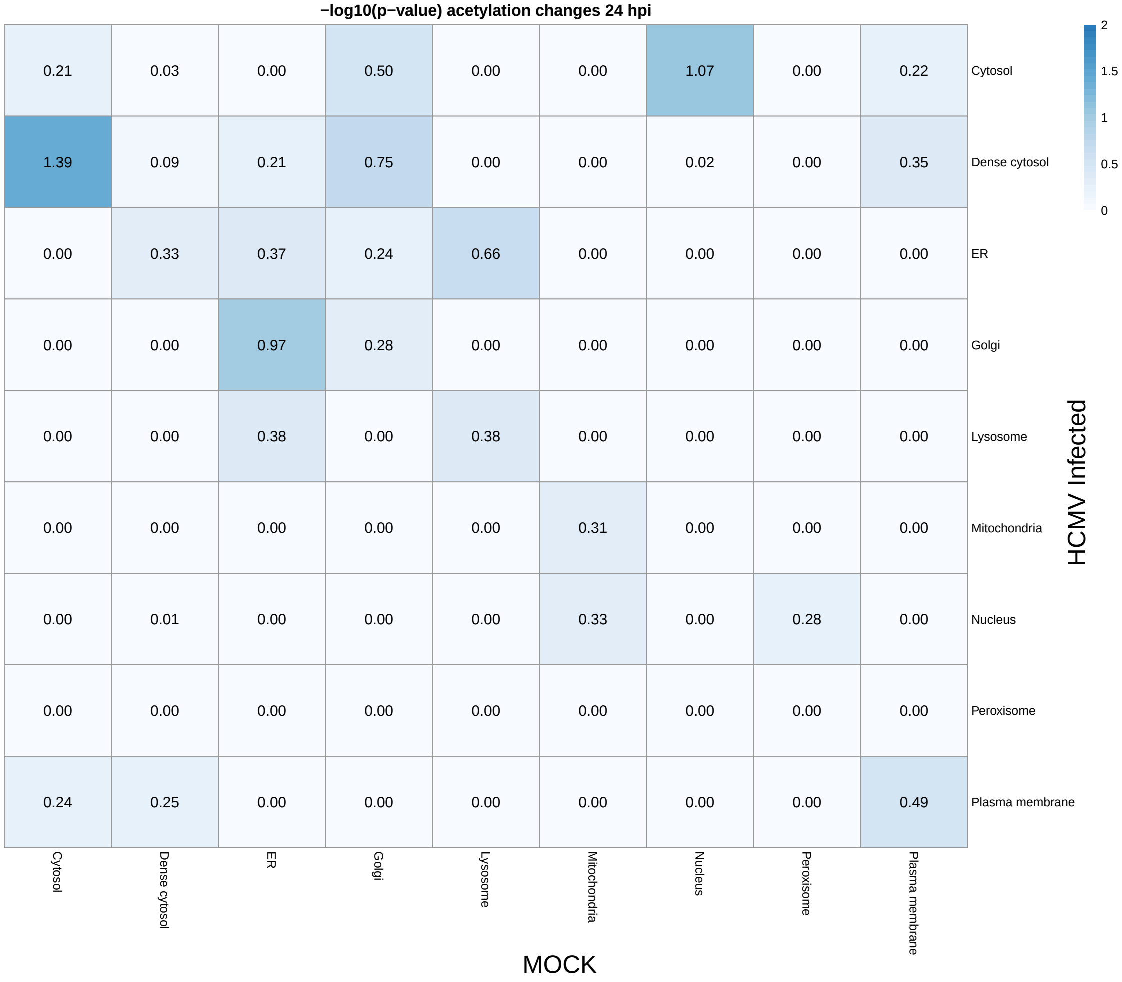

Figure 46: $-\log_{10}(p\text{-values})$ for the fold changes in acetylation stratified by localisation dynamics.

#### 1.18 Appendix 18: HCMV Interactome figures

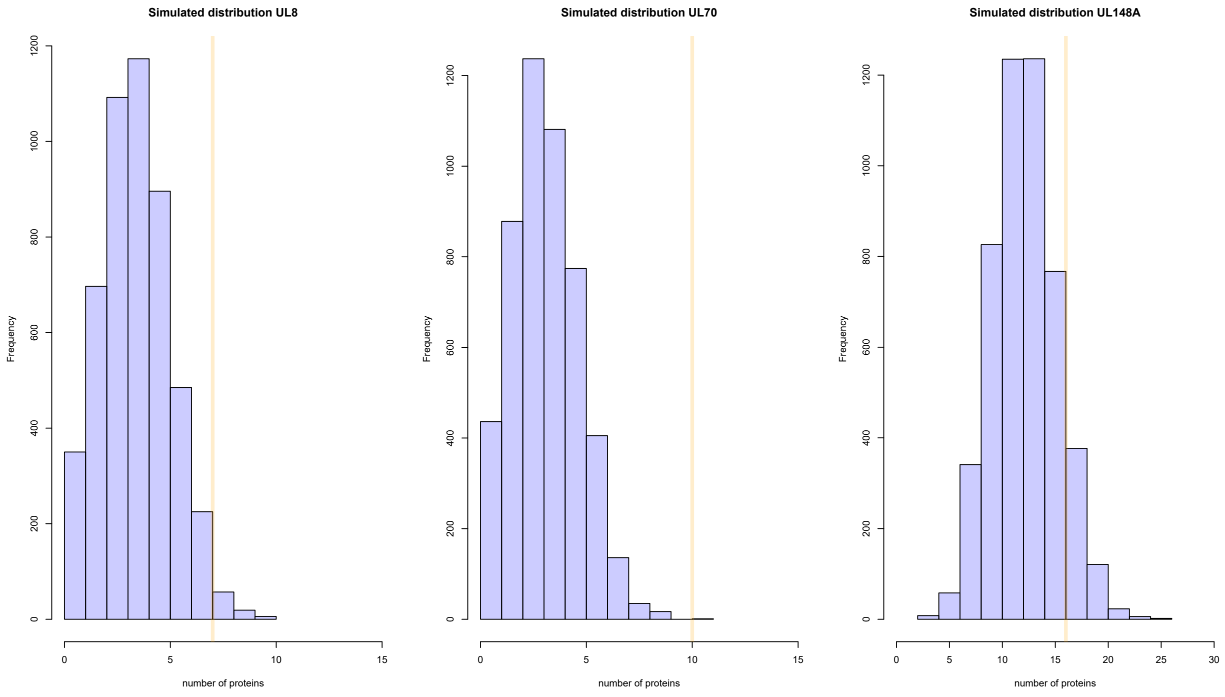

Figure 47: Distributions for predicted number of proteins to be in the same localisation give the spatial pattern observed in A for each of the viral interactomes. The observed statistic is marked in orange, UL8 and UL70 have more proteins in the same location than would be expected at random. This is not true for UL148A.

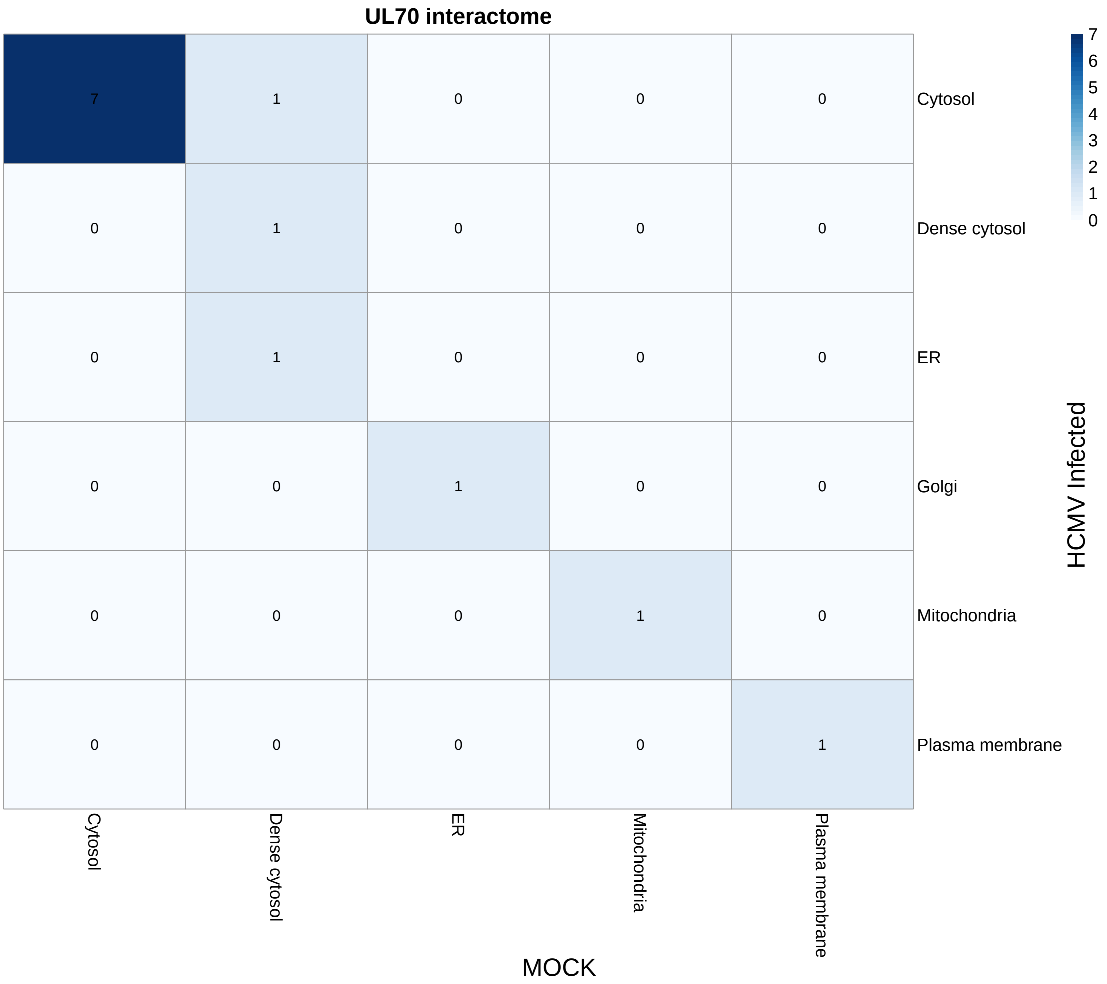

Figure 48: Localisation dynamics for the UL70 interactome

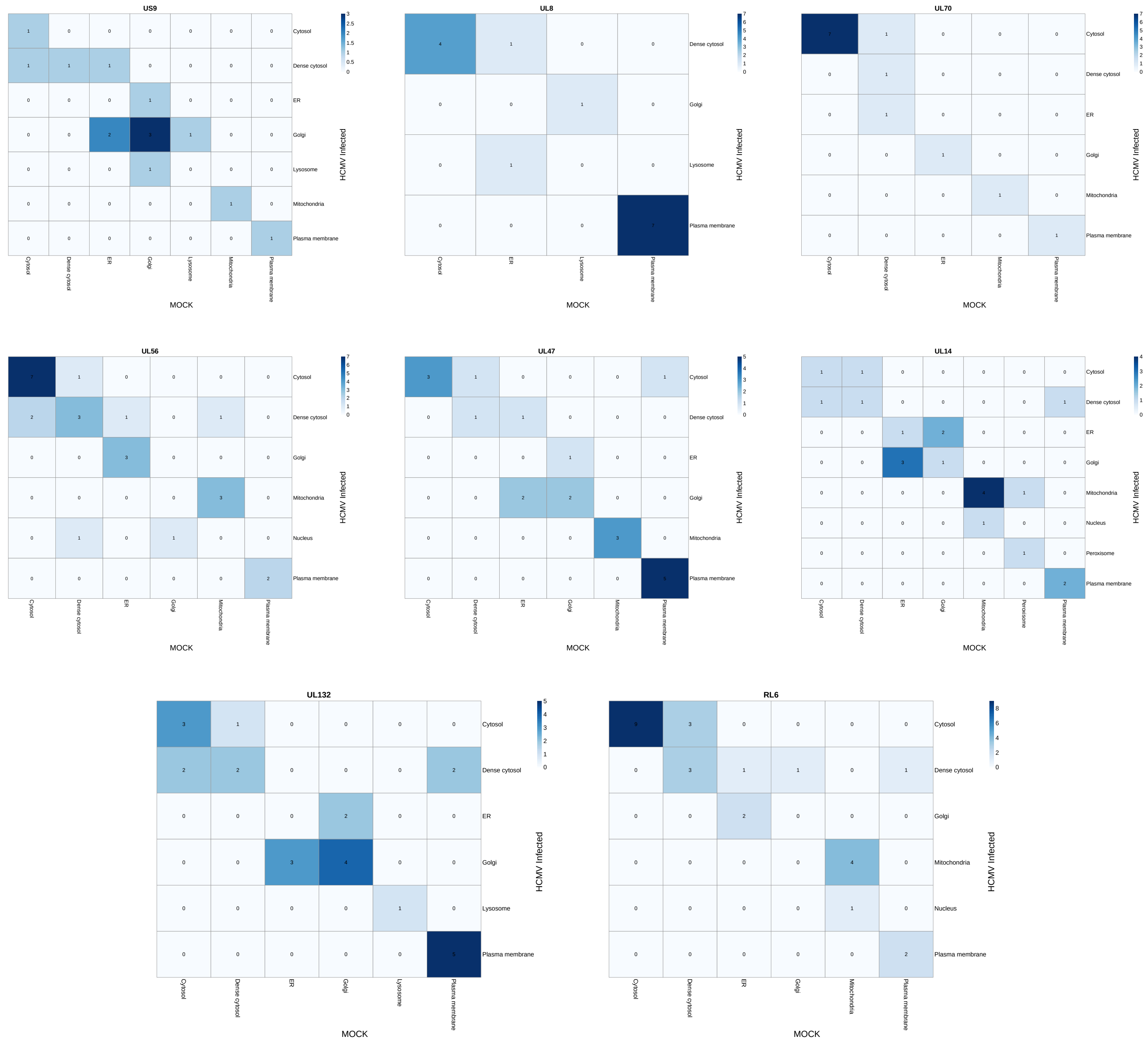

Figure 49: Protein localisation distribution and relocalisation for viral interactomes, the viral bait is indicated in the heading of each heatmap.

#### 1.19 Appendix 19: Example changes in multi-localisation in the HCMV dataset

Sampling from the posterior distribution of localisation probabilities and using a probabilistic framework for analysis allows us to use BANDLE to examine proteins with uncertain localisations. One possible explanation for proteins with uncertain localisation to one or more subcellular niches is that they are genuine residents of multiple organelles. This could either be simultaneously in the same cells or due to single-cell heterogeneity. Spatial proteomics via mass-spectrometry cannot distinguish the two scenarios. Here we plot the posterior localisation probabilities for some proteins from the HCMV dataset. We are able to simultaneously obtain information about mixed localisations and also differential localisation. Thus, we obtain possible candidates for changes in mixed localisation. This analysis would be extremely challenging without a Bayesian approach. These changes could be explained by differential recruitment, synthesis or degradation of proteins at either localisation or genuine perturbation to trafficking and localisation. Complex experiments would be required to distinguish the possibilities.

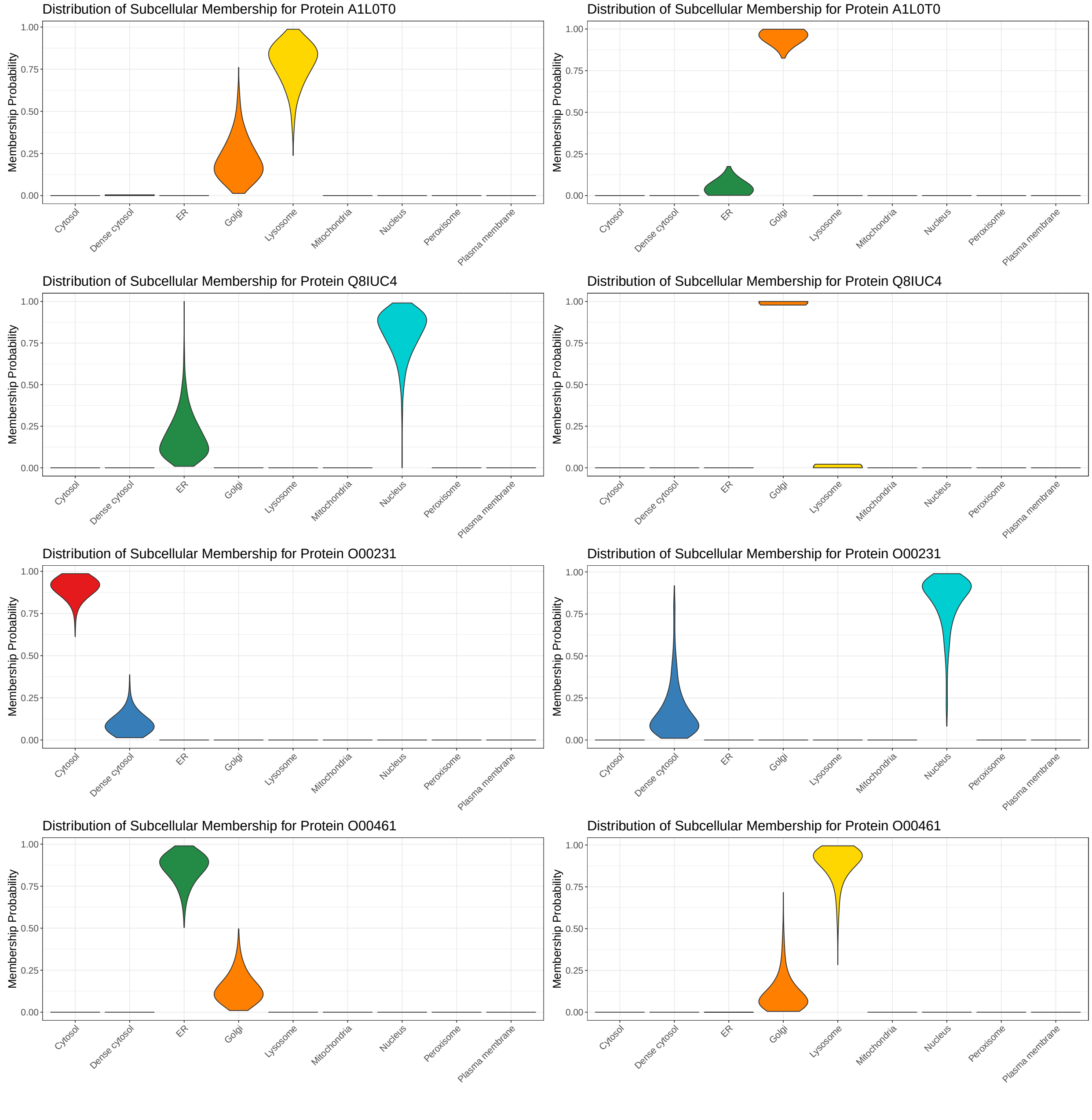

Figure 50: Distributions of posterior localisation probabilities for some differentially localised proteins from the HCMV dataset. All examples show mixed localisation in each condition.

##### 1.20 Appendix 20: Supplementary methods

###### 1.20.1 Penalised Complexity Priors

The penalised-complexity (PC) prior framework introduced by Simpson *et al.* (2017) and Fuglstad *et al.* (2019) allows priors to be specified in modular fashion. In brief, the PC prior framework considers model components, such as GRFs, as a flexible extension of some *base model*. Priors are then constructed such that they shrink the more flexible model towards the base model. The first consideration is an appropriate distance from the base model, $P_0$, to the flexible model, $P$. Simpson *et al.* (2017) choose the distance as $\sqrt{2\text{KL}(P||P_0)}$, where $\text{KL}(P||P_0)$ denotes the *Kullback-Leibler* divergence from $P_0$ to $P$. The square root and the factor 2 puts the distance on the appropriate scale (Simpson *et al.*, 2017). A constant-rate penalisation principles is then used to derive the following condition

$$\frac{\pi(t+\delta)}{\pi(t)} = r^{\delta}\ t, \delta > 0, \tag{1}$$

where $0 < r < 1$ is the constant rate decay. This construction means that the strength of the penalty on the model increases as we depart from the base model. The only continuous distribution that satisfies this property is the exponential distribution $\pi(t) = \lambda \exp(-\lambda t)$ for $t > 0$. The hyperparameter $\lambda$ allows expert information to be expressed, once the prior has been transformed onto the quantity of interest.

###### 1.20.2 Penalised Complexity Priors for GRFs

To derive PC priors for GRF with a Matern covariance requires some complex considerations. Firstly, it is required to parametrise the Matern covariance to clarify which parameters are identifiable under in-fill asymptotics (Fuglstad *et al.*, 2019). This parametrisation loses physical interpretation but is more amenable to theoretical considerations:

$$\kappa = \frac{\sqrt{8\nu}}{\rho} \text{ and } \tau = a\kappa^{\nu}\sqrt{\frac{\Gamma(\nu+1/2)(4\pi)^{1/2}}{\Gamma(\nu)}}. \tag{2}$$

Thus, since $\tau$ can be inferred under in-fill asymptotics and $\kappa$ cannot, a joint PC prior $\pi(\kappa,\tau) = \pi(\tau|\kappa)\pi(\kappa)$ is constructed in two stages. Fuglstad *et al.* (2019) argue that the appropriate base model in the this scenario are GRFs with infinite length-scales and zero marginal variance. Using spectral representations of the GRFs Fuglstad *et al.* (2019) derive the required joint PC prior for $\kappa$ and $\tau$. It is then simple to parametrise the prior for the parameters $\rho$ and $a$. The joint PC prior is stated below (Fuglstad *et al.*, 2019):

**Theorem 1.** *Joint PC prior for GRFs*
*Let $u$ be a GRF defined on $\mathbb{R}$, with Matern covariance with parameters $a, \rho$ and $\nu$. Then the joint PC prior $\pi(a,\rho)$ corresponding to a base model with infinite range and zero variance is*

$$\pi(a,\rho) = \frac{\lambda_1\lambda_2}{2}\rho^{-3/2}\exp(-\lambda_1\rho^{-1/2} - \lambda_2 a), \tag{3}$$

*where $P(\rho < \rho_0) = \alpha_1$ and $P(a > a_0) = \alpha_2$ are achieved by*

$$\lambda_1 = -\log(\alpha_1)\rho^{1/2} \textit{ and } \lambda_2 = \frac{-\log(\alpha_2)}{a_0}. \tag{4}$$

###### 1.20.3 Penalised complexity prior for the noise model

The noise effect is distributed according to $\varepsilon_{ij} \sim \mathcal{N}(0, \sigma_k^2)$ for $k = 1, ..., K$. We additionally choose a PC prior in this scenario, first we reparametrize in terms of a precision $\tau_k = 1/\sigma_k^2$ for $k = 1, ..., K$. Then appealing to Simpson *et al.* (2017) the PC prior is a type-2 Gumbel distribution:

$$\pi(\tau) = \frac{\lambda_3}{2} \tau^{-3/2} \exp(-\lambda_3 \tau^{-1/2}). \tag{5}$$

The penalised complexity prior in this case shrinks towards zero variance. The hyperparameter $\lambda_3$ can be set using the following tail probability $p(\sigma_k > U) = \alpha$ results in $\lambda_3 = \frac{-\log(\alpha)}{U}$.

###### 1.20.4 Semi-supervised inference

Some proteins are deemed markers of their subcellular localisation; that is, they have *a priori* known locations. We use these markers to determine hyperparameters for GRFs for each of the niches. Suppose we are considering the labelled data $X_k^{(L)}$ for organelle $k$ (suppressing notational dependence on the replicate and dataset). Let $n_k$ be the number of proteins in $X_k^{(L)}$ and $D$ the number of locations at which we evaluate the GRF. Finally, let $\tau$ index these locations, denote $C_k$ by the covariance and $\sigma_k^2$ by the variance of the residual noise. We recall that the log-marginal likelihood of a GRF for these data is given by the following (Rasmussen and Williams, 2006):

$$\begin{aligned} &\log p(X|\tau, \boldsymbol{\theta_k}) \\ &\quad = -\frac{1}{2} X_k(\tau) \left(C_k + \sigma_k^2 I_{n_k D}\right)^{-1} X_k(\tau)^T - \frac{1}{2} \log|C_k + \sigma_k^2 I_{n_k D}| - \frac{n_k D}{2} \log 2\pi. \end{aligned} \tag{6}$$

Then the log-posterior is then given by

$$\log p(\boldsymbol{\theta_k}|X, \tau) \propto \log p(X|\tau, \boldsymbol{\theta_k}) + \log p_0(\boldsymbol{\theta_k}), \tag{7}$$

where $p_0(\boldsymbol{\theta_k})$ denotes the (hyper)priors for the parameters, as described in the previous sections, and the constant of proportionality does not depend on $\boldsymbol{\theta_k}$. This posterior distribution could, in principle, be sampled from using MCMC methods, as well as being informed by labelled and unlabelled data (see Crook *et al.* (2019)); however, these considerations add additional computational burden. Here, we choose to fix $\boldsymbol{\theta_k}$ by learning these parameters using *maximum a posteriori* estimation using the labelled data only. This requires optimisation of equation 7 using L-BFGS (Liu and Nocedal, 1989). This approach does not quantify uncertainty in these parameters but reduces the computational cost of our approach. However, the unlabelled data still contribute to the computations of the component specific means and covariances. The L-BFGS can only find a local optimum and so we initialise over a grid of values. We terminate the algorithm when successive iterations of the gradient are less than $10^{-8}$. To further accelerate computations we make extensive use of high performance R packages to interface with C++ (Eddelbuettel and Francois, 2011; Eddelbuettel and Sanderson, 2014).

###### 1.20.5 Modelling Outliers

As shown in previous work some proteins are not well described by any of the annotated components (Crook *et al.*, 2018). This could because of undiscovered biological novelty, poor protein quantitation, the protein could reside in a yet describe sub-cellular component or in multiple annotated compartments. To alleviate this issue we augment our model with an additional outlier component(s). Let us introduce a latent variable indicator $\phi_{i,d}$ to denote whether protein $i$ is better modelled by a subcellular niche or a disperse outlier component in dataset $d$. Since an indicator can only take two values, it has a Bernoulli distribution and so we write $p_0(\phi_{i,d} = 0) = \epsilon_d$. Then we can write the following conditional probability:

$$p(x_{i,1}|z_{i,1} = k, z_{i,2} = j, \phi_{i,1}) \propto \prod_{r=1}^{R} F(x_{i,1}^{(r)}|\boldsymbol{\theta_k^{(r)}})^{\mathbb{1}(\phi_{i,1}=1)} G(x_{i,1}^{(r)}|\Phi^{(r)})^{\mathbb{1}(\phi_{i,1}=0)}, \tag{8}$$

where $\Phi^{(r)}$ denotes the parameters of the outlier component for replicates $r$ and $F$ and $G$ are the densities of the niche specific component and outlier component respectively. Furthermore, we can marginalise $\phi$ in the above equation to obtain:

$$p(x_{i,1}|z_{i,1} = k, z_{i,2} = j, \phi_{i,1}) \propto (1 - \epsilon_1) \prod_{r=1}^{R} F(x_{i,1}^{(r)}|\boldsymbol{\theta_k^{(r)}}) + \epsilon_1 \prod_{r=1}^{R} G(x_{i,1}^{(r)}|\Phi^{(r)}), \tag{9}$$

where we note that the mixed-product terms disappear because the product of two opposing indicators is 0. As in previous work, we let $G$ be a student's t-distribution with degrees of freedom 4, mean equal to the empirical mean and the covariance to be covariance of the empirical covariance of the data. We also assume the covariance matrix to be diagonal. Corresponding equations are available to the second dataset. Finally, we place a Beta prior on $\epsilon_d \sim B(u_d, v_d)$ allowing us to specify the a prior number of outliers. We opt for a weakly-informative prior $u_d = 2$ and $v_d = 10$ for $d = 1, 2$.

###### 1.20.6 Accelerated likelihood computations

Recall, the marginal likelihood associated with the GRF priors in equation 6. The burden of this computation is inversion of an $n_k D \times n_k D$ covariance matrix. Our covariance matrix has a particularly simple structure allowing us to exploit extended Trench and Durbin algorithms for fast matrix computations (Durbin, 1960; Trench, 1964; Zhang *et al.*, 2005; Crook *et al.*, 2019). The structure is given by

$$\hat{C} = \sigma^2 I_{nD} + B, \tag{10}$$

where

$$B = J_n \otimes A, \tag{11}$$

and $\otimes$ denotes the Kronecker (tensor) product. Naive inversion would would require computation on the order of $O((n_k D)^3)$, whereas the methods describe in Crook *et al.* (2019) reduce this to $O(D^2)$.

###### 1.20.7 A non-conjugate prior

Thus far, we have been using a conjugate Dirichlet prior for the a priori mixing proportions $\boldsymbol{\pi}$. Our model assumes no correlation across $\boldsymbol{\pi}$ due to the use of the Dirichlet distribution. However, we describe how we can extend the model to include correlations. Firstly, the joint prior of the allocation probabilities is

$$(z_{i,1}, z_{i,2}) \sim cat(f(\boldsymbol{\pi})), \tag{12}$$

where $f(\boldsymbol{\pi}) = \frac{\exp(\boldsymbol{\pi})}{\sum_{j,k}\exp(\pi_{jk})}$. Thus the prior correlations between organelles can be included using a multivariate Gaussian

$$vec(\boldsymbol{\pi})|\mu, \Sigma \sim \mathcal{N}(\mu, \Sigma). \tag{13}$$

Given $\boldsymbol{\pi}$ the underlying conditional posterior allocation probabilities are the same as before. The conditional posterior of $\boldsymbol{\pi}$ now changes and because of loss of conjugacy a metropolis-hastings step is required. In the next section, we develop a prior using stick-breaking Pólya-Gamma augmentation to facilitate Gibbs sampling.

##### 1.20.8 Pólya-Gamma augmentation

A random variable $X$ has a Pólya-Gamma distribution with parameters $b > 0$ and $c \in \mathbb{R}$, denoted $X \sim \mathcal{PG}(b, c)$ if (Polson *et al.*, 2013)

$$X =_d \frac{1}{2\pi^2}\sum_{k=1}^{\infty}\frac{g_k}{(k-\frac{1}{2})^2 + \frac{c^2}{4\pi^2}}, \tag{14}$$

where $g_k \sim_{iid} \mathcal{G}(b, 1)$. The fundamental equation that renders Pólya-Gamma augmentation useful is the following

$$\frac{(e^{\phi})^a}{(1+e^{\phi})^b} = 2^{-b}e^{\kappa\phi}\int_0^{\infty} e^{-\omega\phi^2/2}p(\omega)\,d\omega, \tag{15}$$

where $\kappa = a - b/2$ and $w \sim \mathcal{PG}(b, 0)$. This is advantageous because of the following construction. Consider the binomial regression problem

$$y_i = \text{Binom}\left(n_i, \frac{1}{1+e^{-x_i^T\beta}}\right), \tag{16}$$

where $\beta$ has the Gaussian prior $\mathcal{N}(\mu, \Sigma)$. To sample from the posterior using Pólya-Gamma augmentation, we first introduce strategic variables $w$ and then sample according to

$$w_i|\beta \sim \mathcal{PG}(n_i, x_i^T\beta) \tag{17}$$

$$\beta|y, w \sim \mathcal{N}(\tilde{\mu}, \tilde{\Sigma}), \tag{18}$$

where

$$\tilde{\Sigma} = (X^T\Omega X + \Sigma^{-1})^{-1} \tag{19}$$

$$\tilde{\mu} = \tilde{\Sigma}(X^T\kappa + \Sigma^{-1}\mu), \tag{20}$$

and

$$\kappa = (y_1 - n_1/2, ..., y_N - n_N/2) \tag{21}$$

$$\Omega = diag(\omega_1, ..., \omega_N). \tag{22}$$

Thus, a simple tuning free two step auxiliary variable sampler is required rather than Metropolis-Hastings move. To adapt this method to our situation we consider a slightly different construction in the following section.

##### 1.20.9 Stick-breaking Pólya-Gamma augmentation

In this section, we extended the Pólya-Gamma augmentation of the previous section to multinomial variables using a stick-breaking approach (Linderman *et al.*, 2015). Consider a likelihood of the form

$$p(x|\phi) = c(x)\frac{(e^{\phi})^{a(x)}}{(1+e^{\phi})^{b(x)}}. \tag{23}$$

The joint probability distribution can then be written as

$$p(\phi, x) = p(\phi)c(x)\frac{(e^{\phi})^{a(x)}}{(1+e^{\phi})^{b(x)}} = p(\phi)c(x)2^{-b(x)}e^{\kappa(x)\phi}\int_0^{\infty} e^{-\omega\phi^2/2}p(\omega)\,d\omega \tag{24}$$

Thus, the conditional distribution can be written as

$$p(\phi|x,\omega) \propto p(\phi)e^{\kappa(x)\phi}e^{-\omega\phi^2/2}, \tag{25}$$

which is Gaussian if $p(\phi)$ is Gaussian. Furthermore, by the exponential tilting property of the Pólya-Gamma distribution (Polson *et al.*, 2013) it follows that

$$\omega|\phi, x \sim \mathcal{PG}(b(x), \phi). \tag{26}$$

Now consider a multinomial model on $K$ categories with $N$ trials with probability vector $\pi$. It can be written as a stick-breaking construction of binomials as follows

$$\text{Multi}(x|N,\pi) = \prod_{k=1}^{K-1}\text{Binom}(x_k|N_k, \tilde{\pi}_k), \tag{27}$$

where

$$N_k = N - \sum_{j<k} x_j \tag{28}$$

$$\tilde{\pi}_k = \frac{\pi_k}{1-\sum_{j<k}\pi_j}. \tag{29}$$

Let $\sigma(\phi_k) = \exp(\phi_k)/(1+\exp(\phi_k))$ and $\tilde{\pi}_k = \sigma(\phi_k)$. Now Substituting into the stick-breaking model

$$\begin{aligned}\text{Multi}(x|N,\pi) &= \prod_{k=1}^{K-1}\text{Binom}(x_k|N_k, \sigma(\phi_k)) && (30)\\ &= \prod_{k=1}^{K-1}\binom{N_k}{x_k}\sigma(\phi_k)^{x_k}(1-\sigma(\phi_k))^{N_k-x_k} && (31)\\ &= \prod_{k=1}^{K-1}\binom{N_k}{x_k}\frac{(e^{\phi_k})^{x_k}}{(1+e^{\phi_k})^{N_k}}. && (32)\end{aligned}$$

Thus, we can set $a_k(x) = x_k$ and $b_k(x) = N_k$ and introduce Pólya-Gamma variables $w_k$. Then

$$p(x,w|\phi) \propto \prod_{k=1}^{K-1}\exp\left[\left(x_k - \frac{N_k}{2}\right)\phi_k - \frac{w_k\phi_k^2}{2}\right] \propto N(\phi|\Omega^{-1}\kappa(x), \Omega^{-1}), \tag{33}$$

where $\Omega = diag(\omega_1, ..., \omega_K)$ and $\kappa(x_k) = x_k - N_k/2$.

###### 1.20.10 A correlated model for differential localisation

The above schema allows us to construct a correlated differential localisation model, using stick-breaking Pólya-Gamma augmentation. Suppose that there are $K$ organelles to which a protein could localises. Then we specify a joint model on the allocation probabilities

$$vec(\boldsymbol{\pi})|\mu, \Sigma \sim \mathcal{N}(\mu, \Sigma) \tag{34}$$
$$(z_{i,1}, z_{i,2}) \sim cat(f(\boldsymbol{\pi})) \tag{35}$$
$$\omega \sim \mathcal{PG}(1, 0), \tag{36}$$

For easy of notation let $\psi = vec(\boldsymbol{\pi})$ and $f$ is the stick-breaking map. Then it follows from the previous sections

$$p(\psi|Z_1, Z_2, \omega) \propto N(\psi|\Omega^{-1}\kappa, \Omega^{-1})N(\psi|\mu, \Sigma) \propto N(\psi|\tilde{\mu}, \tilde{\Sigma}), \tag{37}$$

where

$$\tilde{\mu} = \tilde{\Sigma}\left(\kappa + \Sigma^{-1}\mu\right) \tag{38}$$
$$\tilde{\Sigma} = \left(\Omega + \Sigma^{-1}\right)^{-1}. \tag{39}$$

To compute $\kappa$, first let $n_{j,k} = \sum_i \mathbb{1}(z_{i1} = j, z_{i2} = k)$ and let $\boldsymbol{n} = vec(n)$. Then $\kappa_l = \boldsymbol{n}_l - \frac{1}{2}$ for $l = 1, ..., K^2$. Finally, we can sample the conditional posterior of the Pólya-Gamma variables

$$\omega_l|Z_1, Z_2, \psi \sim \mathcal{PG}(1, \psi_l). \tag{40}$$

###### 1.20.11 Calibration of Polya-Gamma prior

The Polya-Gamma augmentation method was used to take advantage of the knowledge that some classes were known to be correlated *a priori*. The Polya-Gamma prior admits are more flexible prior to be placed on the prior allocation probabilities. Recall that the following prior on the allocation probabiltiies

$$p(z_{i,1} = k, z_{i,2} = k'|\boldsymbol{\pi}) = f(\pi_{kk'}). \tag{41}$$

This prior is then expanded hierarchically in the following fasion:

$$vec(\boldsymbol{\pi})|\mu, \Sigma \sim \mathcal{N}(\mu, \Sigma) \tag{42}$$
$$(z_{i,1}, z_{i,2}) \sim cat(f(\boldsymbol{\pi})) \tag{43}$$
$$\omega \sim \mathcal{PG}(1, 0), \tag{44}$$

where,

$$f(\pi_{kk'}) = \sigma(\pi_{kk'})(1 - \sum_{j<k, j'<k'} f(\pi_{jj'})). \tag{45}$$

There are no analytic formula for the moments of the logit-normal distribution and thus analysing the behaviour of the above prior above is challenging. The implied distribution on $f(\boldsymbol{\pi})$ can be computed by standard transformations:

$$p(f(\boldsymbol{\pi})|\mu, \Sigma) = \mathcal{N}(vec(\boldsymbol{\pi})|\mu, \Sigma) \cdot \prod_{k,k'} \left[\frac{1 - \sum_{j<k, j'<k'} f(\pi_{jj'})}{f(\pi_{kk'})\left(1 - \sum_{j\leq k, j'\leq k'} f(\pi_{jj'})\right)}\right]. \tag{46}$$

This equation clearly demonstrate the complexity of the prior. Recall we are interested in the quantity

$$p(z_{i,1} \neq z_{i,2}|\boldsymbol{\pi}) =: \rho_{pg} = \sum_{j,k;j\neq k} f(\pi_{jk}). \tag{47}$$

The prior expectation of the above and the following prior quantile can be used to calibrate the prior:

$$p(N_U \rho_{pg} > q) = p\left(N_U \sum_{j,k;j\neq k} f(\pi_{jk}) > q\right) = \delta. \tag{48}$$

This computation can be performed via Monte-Carlo simulation and corresponding quantiles as for the Dirichlet prior can be calibrated:

$$p\left(N_U \sum_{j,k;j\neq k} f(\pi_{jk}) > q\right) \approx \frac{1}{T}\sum_{t=1}^{T} \mathbb{1}\left(N_U \sum_{j,k;j\neq k} f(\pi_{jk}^{(t)}) > q\right). \tag{49}$$

However, this is impractical in general for user to specify such a complex prior, since it requires the specification of a full covariance matrix. To alleviate this we suggest using prior data to set this prior. We suggest computing $\Sigma_1$ ,the covariance between the classes using the marker data from the first dataset, and likewise $\Sigma_2$, the covariance between the classes from the second dataset. We then set the prior covariance

$$\Sigma = \tau^{-1} \cdot (\Sigma_1 + \lambda I_K \otimes \Sigma_2 + \lambda I_K) \tag{50}$$

or the precision

$$\Sigma^{-1} = \tau \cdot \left((\Sigma_1 + \lambda I_K)^{-1} \otimes (\Sigma_2 + \lambda I_K)^{-1}\right), \tag{51}$$

where $\tau$ is a tuning parameter that is user specified and $\lambda I_K$ is constant multiple of the identity to provide stability.

##### 1.20.12 Prior Coherence Analysis

The previous sections have constructed two different priors, that capture prior beliefs in different ways. The Dirichlet prior is considerably easier to specify and illicit for domain expertise; however, the stick-breaking Pólya-Gamma prior is much more flexible and can encode more complex prior beliefs - with the task that the prior is more challenging to specify. This section elaborates on the differences between the priors.

Suppose we are given a Dirichlet prior on $\boldsymbol{\pi}$, we can compute the corresponding distribution on $\psi = g_{SB}^{-1}(\boldsymbol{\pi})$, where $g_{SB}$ denotes the stick-breaking map. Recall the matrix Dirichlet prior:

$$q(\boldsymbol{\pi}|\alpha) = \prod_{k=1}^{K} \frac{1}{\mathcal{B}(\alpha_k)} \prod_{j=1}^{K} \pi_{jk}^{\alpha_{jk}-1}. \tag{52}$$

The induced prior on $\psi$, computed from a change of variables, is the following

$$q(\psi|\alpha) = \frac{1}{\mathcal{B}(\alpha)} \prod_{k,k'} \sigma(\psi_{k,k'})^{\alpha_{kk'}} \sigma(-\psi_{k,k'})^{\sum_{j>k,j>k'} \alpha_{jj'}}. \tag{53}$$

As well as looking at the induced priors on the corresponding parameter spaces, we can compute how far about these priors are from each other. Aitchison demonstrated that the the Dirichlet distribution and logit-Normal are never equal for any choice of parameters; however, there are parameters choices that minimise the *Kullback-Leibler* (KL) divergence between them (Aitchison and Shen, 1980; Aitchison, 1982). The stick-breaking Polya-Gamma prior in less straightforward to work with than the Logit-Normal, but facilitates Gibbs sampling. Furthermore, the Logit-Normal transform preserves permutation symmetry in the density; while the stick-breaking transform does not preserve symmetry.

In light of similar analysis, we compute the KL divergence, defined below, between the two priors: the Gaussian Prior and the prior induced on this space by the inverse stick-breaking map from the Dirichlet prior. The KL divergence is

$$KL(P||Q) = \int_{\mathcal{X}} \log\left(\frac{dP}{dQ}\right) dP, \tag{54}$$

for probability measures $P$ and $Q$ defined on measurable space $\mathcal{X}$ and $\frac{dP}{dQ}$ the Radon-Nikodym derivative of $P$ with respect to $Q$. Thus, we compute as follows, where, for ease of notation, we re-label the indexes, such that $vec(\alpha) = [\alpha_1, ..., \alpha_D]$ and likewise for $\psi$ (with abuse of notation).

$$\begin{aligned} KL(p(\psi|\mu,\Sigma)||q(\psi|\alpha)) &= \int p(\psi|\mu,\Sigma)\log\frac{p(\psi|\mu,\Sigma)}{q(\psi|\alpha)} d\psi \\ &= \mathbb{E}[\log\mathcal{N}(\psi|\mu,\Sigma)] - \mathbb{E}[\log q(\psi|\alpha)] \\ &= (A) - (B), \end{aligned} \tag{55}$$

where the expectations are computed with respect to $p$. Continuing the computation

$$\begin{aligned} (A) &= \mathbb{E}\left[\log\left((2\pi)^{-D/2}|\Sigma|^{-1/2}\exp\left(\frac{1}{2}(\psi-\mu)^T\Sigma^{-1}(\psi-\mu)\right)\right)\right] \\ &= -\frac{D}{2}\log(2\pi) - \frac{1}{2}\log|\Sigma| - \frac{1}{2}\mathbb{E}\left[tr\left((\psi-\mu)^T\Sigma^{-1}(\psi-\mu)\right)\right] \\ &= -\frac{D}{2}\log(2\pi) - \frac{1}{2}\log|\Sigma| - \frac{1}{2}\Sigma^{-1}tr\left(\mathbb{E}\left[(\psi-\mu)(\psi-\mu)^T\right]\right) \\ &= -\frac{D}{2}\log(2\pi) - \frac{1}{2}\log|\Sigma| - \frac{1}{2}D \\ &= -\frac{1}{2}\log((2\pi e)^D|\Sigma|), \end{aligned} \tag{56}$$

where in the second line we employed the trace trick and in the third line the linearity of the expectation. For part (B), we write

$$
\begin{aligned}
(B) &= \mathbb{E}\left[\log q(\psi|\alpha)\right] \\
&= \mathbb{E}\left[\log\left(\frac{1}{\mathcal{B}(\alpha)}\prod_{k=1}^{D-1}\sigma(\psi_k)^{\alpha_k}\sigma(-\psi_k)^{\sum_{j=k+1}^{D}\alpha_j}\right)\right] \\
&= -\log\mathcal{B}(\alpha) + \sum_{k=1}^{D-1}\mathbb{E}\left[\alpha_k\log(\sigma(\psi_k)) + \sum_{j=k+1}^{D}\alpha_j\log(\sigma(-\psi_k))\right] \\
&= -\log\mathcal{B}(\alpha) + \sum_{k=1}^{D-1}\alpha_k\mathbb{E}\left[\log(\sigma(\psi_k))\right] + \sum_{k=1}^{D-1}\sum_{j=k+1}^{D}\alpha_j\mathbb{E}\left[\log(\sigma(-\psi_k))\right] \\
&= -\log\mathcal{B}(\alpha) + \sum_{k=1}^{D-1}\alpha_k\mathbb{E}\left[\log(\sigma(\psi_k))\right] + \sum_{k=2}^{D}(k-1)\alpha_k\mathbb{E}\left[\log(\sigma(-\psi_k))\right].
\end{aligned}
\tag{57}
$$

To compute the first summand, we expand the logistic function and then make a second order Taylor approximation about $x_0 = \mathbb{E}[x]$.

$$
\begin{aligned}
\sum_{k=1}^{D-1}\alpha_k\mathbb{E}\left[\log(\sigma(\psi_k))\right] &= -\sum_{k=1}^{D-1}\alpha_k\mathbb{E}\left[\log(1+e^{-\psi_k})\right] \\
&\approx -\sum_{k=1}^{D-1}\alpha_k\left(\log(1+e^{-\mathbb{E}[\psi_k]}) + \frac{e^{\mathbb{E}[\psi_k]}}{(1+e^{\mathbb{E}[\psi_k]})^2}\cdot\mathbb{V}(\psi_k)\right) \\
&= -\sum_{k=1}^{D-1}\alpha_k\left(\log(1+e^{-\mu_k}) + \frac{e^{\mu_k}}{(1+e^{\mu_k})^2}\cdot\Sigma_{kk}\right).
\end{aligned}
\tag{58}
$$

Then, likewise for the second summand

$$
\begin{aligned}
\sum_{k=2}^{D}(k-1)\alpha_k\mathbb{E}\left[\log(\sigma(-\psi_k))\right] &= -\sum_{k=2}^{D}(k-1)\alpha_k\mathbb{E}\left[\log(1+e^{\psi_k})\right] \\
&\approx -\sum_{k=2}^{D}(k-1)\alpha_k\left(\log(1+e^{\mathbb{E}[\psi_k]}) - \frac{e^{\mathbb{E}[\psi_k]}}{(1+e^{\mathbb{E}[\psi_k]})^2}\cdot\mathbb{V}(\psi_k)\right) \\
&= -\sum_{k=2}^{D}(k-1)\alpha_k\left(\log(1+e^{\mu_k}) - \frac{e^{\mu_k}}{(1+e^{\mu_k})^2}\cdot\Sigma_{kk}\right).
\end{aligned}
\tag{59}
$$

Hence,

$$
\begin{aligned}
KL(p(\psi|\mu,\Sigma)||q(\psi|\alpha)) \approx &-\frac{1}{2}\log((2\pi e)^D|\Sigma|) + \log\mathcal{B}(\alpha) \\
&+\sum_{k=1}^{D-1}\alpha_k\left(\log(1+e^{-\mu_k}) + \frac{e^{\mu_k}}{(1+e^{\mu_k})^2}\cdot\Sigma_{kk}\right) \\
&+\sum_{k=2}^{D}(k-1)\alpha_k\left(\log(1+e^{\mu_k}) - \frac{e^{\mu_k}}{(1+e^{\mu_k})^2}\cdot\Sigma_{kk}\right).
\end{aligned}
\tag{60}
$$

To obtain a reasonable scale for the above result, we state the KL divergence between two Dirichlet distributions and two Gaussian distributions. Let us note that the KL divergence between two Dirichlet distribution is the following

$$\begin{aligned} KL(Dir(\pi|\alpha)||Dir(\pi|\alpha')) = \log\Gamma(\alpha_0) - \sum_{k=1}^{K}\log\Gamma(\alpha_k) - \log\Gamma(\alpha'_0) \\ + \sum_{k=1}^{K}\log\Gamma(\alpha'_k) + \sum_{k=1}^{K}(\alpha_k - \alpha'_k)(\psi(\alpha_k) - \psi(\alpha_0)), \end{aligned} \tag{61}$$

where $\psi$ denotes the digamma function. Likewise the KL divergence between two Gaussian distributions is the following

$$KL(p(x|\mu,\Sigma)||q(x|\mu',\Sigma')) = \frac{1}{2}\left(tr(\Sigma'^{-1}\Sigma) + (\mu' - \mu)\Sigma'^{-1}(\mu' - \mu) - K + \log\frac{|\Sigma'|}{|\Sigma|}\right). \tag{62}$$

##### 1.20.13 Priors in integrative mixture models

For completeness, we include a discussion on other priors used in integrative mixture models. The first example we consider is the multiple dataset integration (MDI) method of (Kirk *et al.*, 2012), where the joint prior for allocations (in the two dataset scenario) is given by:

$$\begin{aligned} \phi &\sim \mathcal{G}(a,b) \\ \pi_1 &\sim Dir(\frac{\alpha_1}{K_1},....,\frac{\alpha_1}{K_1}) \\ \pi_2 &\sim Dir(\frac{\alpha_2}{K_2},...,\frac{\alpha_2}{K_2}) \\ p(z_{i1},z_{i2}|\phi) &\sim \pi_{z_i1}\pi_{z_i2}(1+\phi\mathbb{1}(z_{i1}=z_{i2})). \end{aligned} \tag{63}$$

Meanwhile for clusternomics (Gabasova *et al.*, 2017) the prior is

$$\begin{aligned} \rho &\sim Dir(\gamma\text{vec}(\pi_1 \otimes \pi_2)) \\ \pi_1 &\sim Dir(\frac{\alpha_1}{K_1},....,\frac{\alpha_1}{K_1}) \\ \pi_2 &\sim Dir(\frac{\alpha_2}{K_2},...,\frac{\alpha_2}{K_2}) \\ p(z_{i1}=k,z_{i2}=j) &= \rho_{kj}. \end{aligned} \tag{64}$$

The model for Bayesian Consensus Clustering (BCC) is the following (Lock and Dunson, 2013). First, define a global latent allocation $C = \{c_1, ..., c_n\}$, to one of $K$ possible clusters. Then, for the $d^{th}$ dataset define the local latent allocation $z_{id}$, where the conditional probability is given by

$$p(z_{id}=k|c_i) = \alpha_d\mathbb{1}(z_{id}=c_i) + \frac{1-\alpha_d}{K-1}(1-\mathbb{1}(z_{id}=c_i)). \tag{65}$$

The key idea of BCC is that first a global latent allocation (or clustering) is defined and then local clusterings are defined conditional on the global clustering. The concentration parameter $\alpha_d$ controls the level of association between the global and local allocation. Importantly, note that if the $k^{th}$ global component is empty, then corresponding local component probability is 0 if and only if $\alpha_d = 1$. Hence, in general, there will be more local components than global components. The approach is most similar to MDI, which allows the clusters to vary arbitrary between the datasets. In the language of BCC and clusternomics, each dataset is allowed its own set of local clusters. Then the parameter $\phi$ up weights the prior probability that observations are allocated to the corresponding local components in each dataset. Note if $\phi = 0$ then we are in the independent case and so, in general, there is some up weighting of the joint allocation probabilities as $p_0(\phi > 0) > 0$ and more up weighting if the datasets are more similar. Clusternomics is, somewhat, the reverse of BCC. In contrast, allocations are defined first at the local level and then information is shared via a global allocations. As the number of local clusters increase so does the number of global clusters. Furthermore, the global concentration parameter $\gamma$ controls the level of sharing across the datasets. The difference between the prior used in our approach, BANDLE, and clusternomics can be seen in multiple ways. Firstly, we defined mixture proportions across datasets rather than explicit local/global clusters. Secondly, if there are $K$ components then clusternomics has $2K + 1$ hyperparameters whilst BANDLE has $K^2$ parameters. Finally, it is clearest to see the differences via the marginal probabilities. Rewriting the BANDLE priors in the notation of of clusternomics for clarity, we write the marinal probabilities fo clusternomics:

$$p(z_{i1} = k, z_{i2} = j) = \rho_{kj}, \;\; \rho_{kj} \sim \mathcal{B}(\gamma\pi_{kk}\pi_{jj}, \pi_0 - \gamma\pi_{kk}\pi_{jj}), \tag{66}$$

where $\pi_0 = \sum_{j,k} \pi_{jk}$. Likewise for BANDLE

$$p(z_{i1} = k, z_{i2} = j) = \rho_{kj}, \;\; \rho_{kj} \sim \mathcal{B}(\pi_{kj}, \pi_0 - \pi_{kj}), \tag{67}$$

which we see allows for a much more flexible prior specification.

###### 1.20.14 Simulating dynamic spatial proteomics experiments

We describe the ways in which we produce synthetic dynamic spatial proteomics experiments from real datasets. The expression value for each protein can be written as follows:

$$y_i = f_k + \varepsilon_i \tag{68}$$

for some value $k = 1, ..., K$ which index the $K$ possible subcellular niches. The value of $f_k$ is unknown so we estimate it from the data. We use K-NN classification, with the number of nearest neighbours $\widehat{K} = 10$, to assign every protein to an organelle. That is, the probability the $i^{th}$ protein belongs to the $j^{th}$ organelle is approximated by:

$$P(z_i = j|Y = y_i) \approx \frac{1}{\widehat{K}} \sum_{l \in \mathcal{N}_i} \mathbb{1}(y_l = j), \tag{69}$$

where $\mathcal{N}_i$ is the set of $\widehat{K}$ closest labelled points to $y_i$. We then assign proteins to their most probable subcellular niche. We proceed to estimate $f_k$ for $k = 1, ..., K$ by the mean of expression values of all the proteins allocated to that niche:

$$\hat{f}_k \approx \frac{1}{|n_k|} \sum_{i \in n_k} y_i, \tag{70}$$

where $n_k$ indexes the proteins assigned to the $k^{th}$ subcellular niche. We then use the residual bootstrap to generate synthetic data. To be precise, we first compute the residuals

$$\hat{\varepsilon}_i = y_i - \hat{f}_k \; i = 1, ..., N, \tag{71}$$

where $k$ is the organelle to which protein $i$ was assigned by K-NN classification. We then obtain $\mathcal{E} = \{\hat{\varepsilon}_{i,g}\}_{g=1}^{G}$, where $G$ is length of the vector $y_i$. The we use a nonparamateric bootstrap (uniform sampling with replacement) to obtain $\mathcal{E}_B = \{\hat{\varepsilon}^*_{i,g}\}_{g=1}^{G}$. Replicates of the data are then obtained as follows

$$y_i^{rep} = \hat{f}_k + \hat{\varepsilon}^*_i \; i = 1, ..., N. \tag{72}$$

We further propose to use

$$y_i^{rep} = \hat{f}_k + \nu\hat{\varepsilon}^*_i \; i = 1, ..., N, \tag{73}$$

where $\nu$ is some deterministic or random value. In addition, we consider organelle specific multiplicative noise:

$$y_i^{rep} = \hat{f}_k + \nu_k\hat{\varepsilon}^*_i \; i = 1, ..., N, \tag{74}$$

where $\nu_k$ are different random values for $k = 1, ..., K$.

The above process produce replicates without any translocation events. To simulate translocation events we randomly select, with equal probability, $L$ proteins. Then for each of these $l$ proteins we randomly select, with equal probability, one of the $K$ possible organelles to which we translocate the protein. Then we replace the quantitative value for $l^{th}$ protein with a sample from the following distribution

$$y_l \sim \mathcal{N}(\hat{f}_k, \hat{\sigma}^2_{f_k}), \tag{75}$$

where $k$ is the newly assigned organelle and $\hat{\sigma}^2_{f_k}$ is an unbiased estimator of the population variance of $\hat{f}_k$:

$$\hat{\sigma}^2_{f_k} = \frac{1}{|n_k| - 1} \sum_{i \in n_k} (y_i - \hat{f}_k). \tag{76}$$

Different spatial proteomics experiments are usually run on different mass-spectrometry runs and thus both random and systematic batch effects can occur. Furthermore, differences in the labelling efficiency of each tag, as well as slight differences in the amount of protein labelled and how well ions fly in the mass-spectrometer can lead to systematic difference between experiments. Furthermore, there is inherent technical variability in the apparatus and sample handling; for example, density-gradients or differential centrifugation speeds are never precisely the same. We propose three approaches to test the robustness of the available methods to these effects.

*Random batch effects.* After the replicates have been produced and translocation events simulated. We propose to generate random batch effects through the following process. For each replicate in turn, we sample a fraction with equal probability from $S_G = \{1, ..., G\}$. For the sampled fraction, say $g$, we add a random biased effect, $\mu_{batch}$, to that fraction; such that,

$$y_{i,g}^{rep,batch} = y_{i,g}^{rep} + \mu_{batch} \tag{77}$$

*Systematic batch effects.* Systematic batch effects are produce in identical manner to random batch effects, but instead the fraction is sampled first and the effect is added to same fraction across the experiments. The magnitude of the effect is allow to differ across experiments.

*Fraction permutations* We permute the fractions in different experiments, which is designed to reflect the inherent technical variabilities of the procedure. Let $\sigma : S_G \rightarrow S_G$ be a permutation such that $\sigma(S_G) = \{\sigma(1), ..., \sigma(G)\}$. We then replace each fraction with its permuted value, as follows:

$$y_{i,g}^{rep,perm} = y_{i,\sigma(g)}^{rep}. \tag{78}$$

In the five possible simulation scenarios, which are all repeated 10 times, the following setting are used.

- $\nu_k \sim U[1, 2]$

For the systematic and random batch effects we take

- $\mu_{batch} = 0.3$

##### 1.20.15 Bandle in Hierarchical model notation

The Bandle model can be summarised in the following Bayesian Hierarchical model:

$$\begin{aligned}
x_{i,d}^{(r)}|z_{i,d} = k, \theta, \phi_{i,d} = 1 &\sim \mu_k^{(r)}(s_j) + \epsilon_{kj}^{(r)} \\
x_{i,d}^{(r)}|z_{i,d} = k, \theta, \phi_{i,d} = 0 &\sim \mathcal{T}(4, M, V) \\
\mu_{k,d}^{(r)} &\sim GRF(m_{k,d}^{(r)}(\boldsymbol{s}), C_{k,d}^{(r)}(\boldsymbol{s}, \boldsymbol{s'})) \\
\epsilon_{kj}^{(r)} &\sim \mathcal{N}(0, \sigma_{r,k}^2) \\
1/\sigma_{r,k}^2 &\sim \text{Type-2 Gumbel}(\lambda_3) \\
(z_{i,1}, z_{i,2}) &\sim cat(\boldsymbol{\pi}) \\
\pi|\alpha &\sim \mathcal{M}Dir(\alpha, K) \\
\phi_{i,d} &\sim Ber(\epsilon_d) \\
\epsilon_d &\sim B(u_d, v_d) \\
C_v(\delta) &= a^2 \frac{2^{1-\nu}}{\Gamma(\nu)} \left(\sqrt{8\nu}\frac{\delta}{\rho}\right)^\nu \mathcal{K}_v \left(\sqrt{8\nu}\frac{\delta}{\rho}\right) \\
\kappa, \rho &\sim PC(\lambda_1, \lambda_2)
\end{aligned} \tag{79}$$

##### 1.20.16 Major algorithmic steps of Bandle

Here, we summarise the major steps of bandle in algorithmic steps.

1. First, for each subcellular niche $k$ in each dataset $d$ and replicate $r$, learn the GRFs and corresponding hyperparameters by *maximum a posteriori* estimation for a pre-specified $\lambda_1, \lambda_2$ and $\lambda_3$.

2. Select $\alpha$, potentially using a prior predictive check.

3. For $T$ Monte-Carlo iterations perform:

   (a) Compute the likelihood of each protein $i$ belonging to subcellular niche $k$ in replicate $r$ for dataset $d$.

   (b) Sample from the conditional posterior distribution of $\pi$.

   (c) Compute the conditional posterior of each protein $i$ belonging to each subcellular niche $k$ in each dataset $d$.

   (d) Sample $(z_{i,1}, z_{i,2})$ from the computed conditional posterior.

   (e) Compute the likelihood of belonging to the outlier component.

   (f) Sample $\epsilon_d$ from the conditional posterior distribution.

   (g) Sample $\phi_{i,d}$ for all $i$ and all $d$ from the conditional posterior distribution.

   (h) Optional: Sample new GRF hyperparameters for the conditional posterior distribution using Metropolis-Hastings or Hamiltonian Monte-Carlo. Otherwise use precomputed values.

   (i) Update the GRF distributions for each subcellular niche $k$ in replicate $r$ for dataset $d$.

##### 1.20.17 Comparison of normalisation approaches

In this section, we compare how different normalisation approaches affect the performance of the studied methods (BANDLE and MR 2017). We use the same simulation schema in the main text where the MR approach performed best. We compare the normalisation approach in Itzhak *et al.* (2016), where the weighted ratios of the fraction are summed across the fractions and then each ratio is divided by the total this protein. We also compare some popular normalisation approaches, such as row median normalisation and variance-stabilisation normalisation (VSN). The performance of the BANDLE approaches using different normalisation approaches is unchanged, likely as a result of the non-parametric modelling used. Curiously, the MR method performed substantially better if a different normalisation procedure other than that recommend by the original authors (Itzhak *et al.*, 2016, 2017). These normalisation approaches help the underlying normality assumptions of the MR approach.

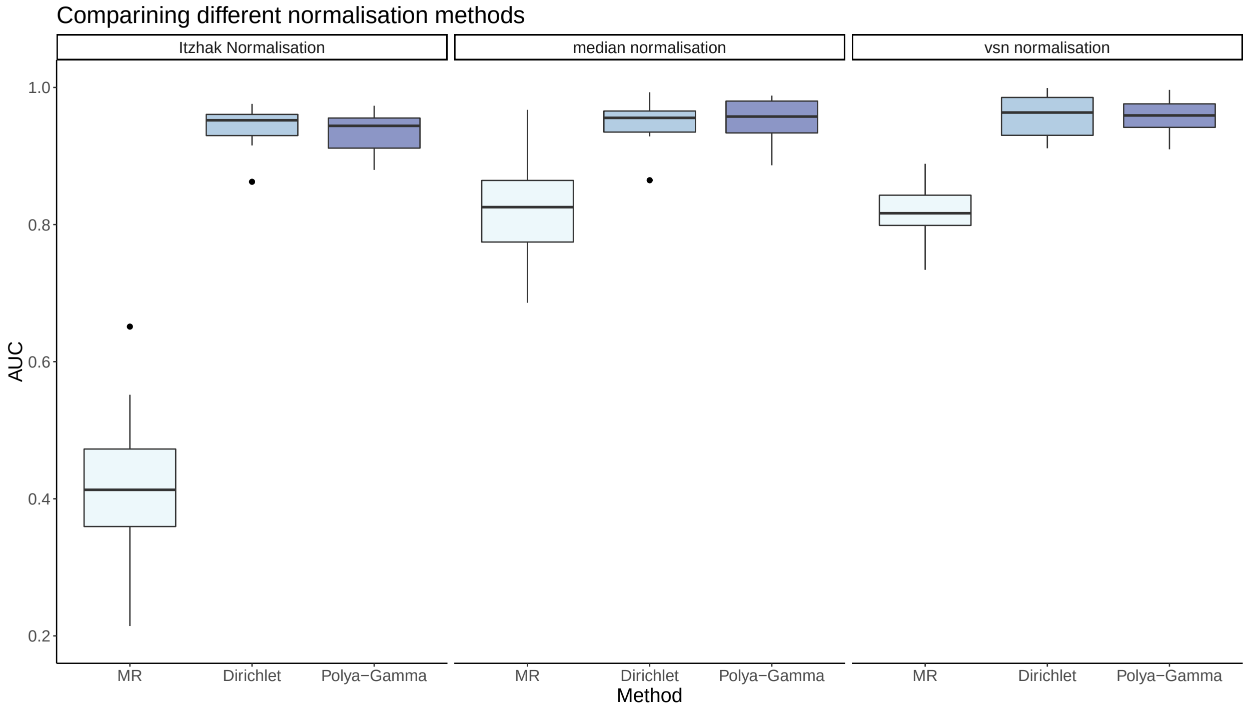

Figure 51: Area under the curve (AUC) for the MR 2017 method and BANDLE with different prior choices. BANDLE performs consistently with different normalisation approaches, whilst the MR 2017 method performs better using median normalisation and VSN normalisation when compared to the normalisation in (Itzhak *et al.*, 2016). Boxplots are over new simulated datasets and runs of the methods.

##### 1.20.18 Frequentist Calibration of BANDLE

One of the major outputs of BANDLE is the differential localisation probability. Scientist will use this probability, additional experiments and supporting literature to decide whether a differential localisation has occurred. To help way up different sources of evidence it is useful to ask what is the frequentist calibration of these probabilities. To perform this analysis, we compute the differential localisation probabilities in a simulation scenario (random batch effects) and compute the expected calibration error (ECE) (Guo *et al.*, 2017). That is, let $B_m$ index samples whose prediction probability falls into the interval $I_m = \left(\frac{m-1}{M}, \frac{m}{M}\right]$. The accuracy of $B_m$ is defined as

$$\mathrm{acc}(B_m) = \frac{1}{|B_m|} \sum_{B_m} \mathbb{1}(\hat{y}_i = y_i), \tag{80}$$

where $\hat{y}_i$ and $y_i$ are the prediction and true label for protein $i$. Meanwhile, the average confidence for bin $B_m$ is

$$\mathrm{conf}(B_m) = \frac{1}{|B_m|} \sum_{B_m} \hat{p}_i, \tag{81}$$

where $\hat{p}_i$ is the predicted confidence for sample $i$. The ECE is then given by

$$\mathrm{ECE} = \sum_{m=1}^{M} \frac{|B_m|}{n} |\mathrm{acc}(B_m) - \mathrm{conf}(B_m)|. \tag{82}$$

Here $n = \sum_{m=1}^{M} B_m$. We find that using the Dirichlet prior the ECE is roughly 4%, whilst the Polya-Gamma prior this rises to 10% (see figure). This is not unexpected from the Polya-Gamma prior, since a subjective Bayesian probability is not designed to have frequenstist properties. For this prior, additional information has been included which improves the performance of the method (in terms of identify differentially localised proteins), but is less well calibrated. Furthermore, these ECEs are small for an approach that does not have a holdout dataset on which to calibrate and it is competitive with values where post-hoc calibration has been performed using a holdout dataset (Guo *et al.*, 2017).

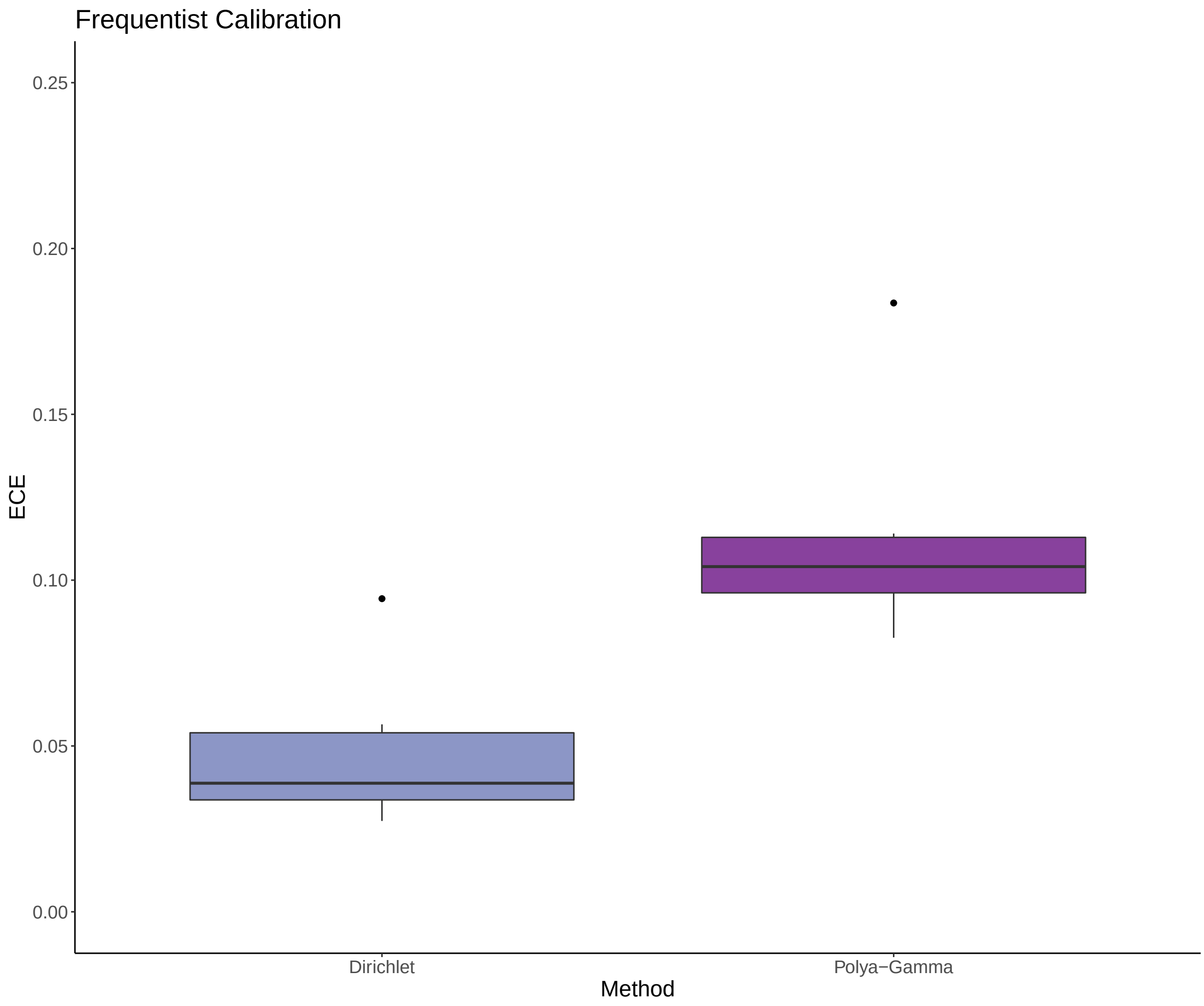

Figure 52: ECE for BANDLE using the Dirichlet and Polya-Gamma priors using a simulation scenerio. Boxplots are over new simulated dataset and runs of the method.

#### References

Aitchison, J. (1982). The statistical analysis of compositional data. *Journal of the Royal Statistical Society: Series B (Methodological)*, **44**(2), 139–160.

Aitchison, J. et al. (1980). Logistic-normal distributions: Some properties and uses. *Biometrika*, **67**(2), 261–272.

Beltran, P. M. J. et al. (2016). A portrait of the human organelle proteome in space and time during cytomegalovirus infection. *Cell systems*, **3**(4), 361–373.

Crook, O. M. et al. (2018). A bayesian mixture modelling approach for spatial proteomics. *PLOS Computational Biology*, **14**(11), 1–29.

Crook, O. M. et al. (2019). Semi-supervised non-parametric bayesian modelling of spatial proteomics. *arXiv preprint arXiv:1903.02909*.

Davies, A. K. et al. (2018). Ap-4 vesicles contribute to spatial control of autophagy via rusc-dependent peripheral delivery of atg9a. *Nature communications*, **9**(1), 3958.

Durbin, J. (1960). The fitting of time-series models. *Revue de l'Institut International de Statistique*, pages 233–244.

Eddelbuettel, D. et al. (2011). Rcpp: Seamless r and c++ integration. *Journal of Statistical Software, Articles*, **40**(8), 1–18.

Eddelbuettel, D. et al. (2014). Rcpparmadillo: Accelerating r with high-performance c++ linear algebra. *Comput. Stat. Data Anal.*, **71**, 1054–1063.

Fuglstad, G.-A. et al. (2019). Constructing priors that penalize the complexity of gaussian random fields. *Journal of the American Statistical Association*, **114**(525), 445–452.

Gabasova, E. et al. (2017). Clusternomics: Integrative context-dependent clustering for heterogeneous datasets. *PLoS computational biology*, **13**(10), e1005781.

Geladaki, A. et al. (2019). Combining lopit with differential ultracentrifugation for high-resolution spatial proteomics. *Nature Communications*, **10**, 331.

Gelman, A. et al. (1995). *Bayesian Data Analysis*. Chapman & Hall, London.

Guo, C. et al. (2017). On calibration of modern neural networks. In *International Conference on Machine Learning*, pages 1321–1330. PMLR.

Itzhak, D. N. et al. (2016). Global, quantitative and dynamic mapping of protein subcellular localization. *Elife*, **5**, e16950.

Itzhak, D. N. et al. (2017). A mass spectrometry-based approach for mapping protein subcellular localization reveals the spatial proteome of mouse primary neurons. *Cell reports*, **20**(11), 2706–2718.

Kirk, P. et al. (2012). Bayesian correlated clustering to integrate multiple datasets. *Bioinformatics*, **28**(24), 3290–3297.

Linderman, S. et al. (2015). Dependent multinomial models made easy: Stick-breaking with the polya-gamma augmentation. In *Advances in Neural Information Processing Systems*, pages 3456–3464.

Liu, D. C. et al. (1989). On the limited memory bfgs method for large scale optimization. *Mathematical programming*, **45**(1-3), 503–528.

Lock, E. F. et al. (2013). Bayesian consensus clustering. *Bioinformatics*, **29**(20), 2610–2616.

Polson, N. G. et al. (2013). Bayesian inference for logistic models using pólya–gamma latent variables. *Journal of the American statistical Association*, **108**(504), 1339–1349.

Rasmussen, C. E. et al. (2006). *Gaussian processes for machine learning*. MIT Press.

Simpson, D. et al. (2017). Penalising model component complexity: A principled, practical approach to constructing priors. *Statistical science*, **32**(1), 1–28.

Trench, W. F. (1964). An algorithm for the inversion of finite toeplitz matrices. *Journal of the Society for Industrial and Applied Mathematics*, **12**(3), 515–522.

Zhang, Y. et al. (2005). Time-series gaussian process regression based on toeplitz computation of o (n 2) operations and o (n)-level storage. In *Decision and Control, 2005 and 2005 European Control Conference. CDC-ECC'05. 44th IEEE Conference on*, pages 3711–3716. IEEE.
